# Prostate tumor-induced stromal reprogramming generates Tenascin C that promotes prostate cancer metastasis through YAP/TAZ inhibition

**DOI:** 10.1101/2021.11.08.467778

**Authors:** Yu-Chen Lee, Song-Chang Lin, Guoyu Yu, Ming Zhu, Jian H Song, Keith Rivera, Darryl Pappin, Christopher J. Logothetis, Theocharis Panaretakis, Guocan Wang, Li-Yuan Yu-Lee, Sue-Hwa Lin

## Abstract

Metastatic prostate cancer (PCa) in bone induces bone-forming lesions that enhance PCa progression. How tumor-induced bone formation enhances PCa progression is not known. We have previously shown that PCa-induced bone originates from endothelial cells (EC) that have undergone endothelial-to-osteoblast (EC-to-OSB) transition by tumor-secreted BMP4. Here, we show that EC-to-OSB transition leads to changes in the tumor microenvironment that increases the metastatic potential of PCa cells. We found that conditioned medium (CM) from EC-OSB hybrid cells increases the migration, invasion and survival of PC3-mm2 and C4-2B4 PCa cells. Quantitative mass spectrometry (iTRAQ) identified Tenascin C (TNC) as one of the major proteins secreted from EC-OSB hybrid cells. TNC expression in tumor-induced osteoblasts was confirmed by immunohistochemistry of MDA-PCa118b xenograft and human bone metastasis specimens. Mechanistically, BMP4 increases TNC expression in EC-OSB cells through the Smad1-Notch/Hey1 pathway. How TNC promotes PCa metastasis was next interrogated by in vitro and in vivo studies. In vitro studies showed that a TNC neutralizing antibody inhibits EC-OSB-CM-mediated PCa cell migration and survival. TNC knockdown decreased, while addition of recombinant TNC or TNC overexpression increased migration and anchorage-independent growth of PC3 or C4-2b cells. When injected orthotopically, PC3-mm2-shTNC clones decreased metastasis to bone, while C4-2b-TNC overexpressing cells increased metastasis to lymph nodes. TNC enhances PCa cell migration through α5β1 integrin-mediated YAP/TAZ inhibition. These studies elucidate that tumor-induced stromal reprogramming generates TNC that enhances PCa metastasis and suggest that TNC may be a target for PCa therapy.

## Introduction

Lethal prostate cancer (PCa) is dominated by complications arising from bone metastasis. One unique feature of PCa bone metastasis is the induction of aberrant bone formation [1, 2]. This contrasts with other cancers, e.g., renal carcinoma and multiple myeloma, which induce osteolytic bone lesions. We and others have shown that PCa-induced aberrant bone formation is partly due to PCa-secreted bone morphogenetic protein (BMP) family proteins, including BMP4 [3] and BMP6 [4], which increase osteoblast differentiation. Such a stromal reprogramming by PCa was shown to enhance PCa progression in bone [3, 5]. However, how tumor-induced bone enhances PCa progression is not known.

Elucidating mechanisms by which PCa induces aberrant bone formation will allow us to identify factors that are involved in such a pathological event. Tumors are known to recruit cells, e.g., endothelial cells, from the host microenvironment to support their growth. Using an osteogenic patient-derived xenograft (PDX) MDA PCa-118b, we found that prostate tumors recruited host (mouse) stromal cells and converted them into osteoblasts (OSB) [5]. We further showed that BMP4 induces tumor associated-endothelial cells (ECs) to undergo endothelial-to-osteoblast (EC-to-OSB) transition, resulting in ectopic bone formation [5]. During BMP4-mediated EC-to-OSB transition, Tie2^+^ ECs undergo transition to become osteocalcin^+^ (Ocl^+^) OSBs, generating EC-OSB hybrid cells that express both Tie2 and Ocl [5]. EC-OSB hybrid cells were found to form a rim along tumor-induced bone and are located in close proximity with tumor cells [5]. We recapitulated the bone forming process in vitro by showing that EC-OSB hybrid cells can further differentiate into mature OSBs when cultured in OSB differentiation medium [5].

The signaling pathways mediating EC-to-OSB transition was examined using an EC line 2H11 [6]. We found that multiple pathways, including Smad1-Notch/Hey1, GSK3β-βcatenin-Slug, and Smad1-Dlx2 pathways are involved in BMP4-mediated EC-to-OSB transition. Activation of these pathways results in upregulation of several transcription factors, including Osx, Dlx2, Slug and Hey1, which play critical roles in EC transition into OSB, OSB differentiation, and bone matrix mineralization [6]. Together, these findings uncovered the mechanism of PCa-induced aberrant bone formation.

The mechanistic understanding of PCa-induced aberrant bone formation allows us to address how tumor-induced bone enhances PCa progression. Because EC-OSB hybrid cells generated during EC-to-OSB transition are a unique cell type before they become mature osteoblasts [5], factors that are secreted by EC-OSB hybrid cells may elicit paracrine effects on tumor cells to modulate their properties. In this study, we identified factors secreted from EC-OSB hybrid cells and found Tenascin C, an extracellular matrix glycoprotein, to be one of the major factors. Using in vitro and in vivo approaches, we showed that Tenascin C increases the metastatic potential of PCa cells, providing a mechanism by which PCa-mediated stromal cell reprogramming enhances PCa progression in bone.

## Results

### Conditioned media from EC-OSB hybrid cells increase the migration, invasion and anchorage-independent growth of prostate cancer cells

We previously showed that treatment of 2H11 endothelial cells (EC) with BMP4 (100 ng/ml) for 48 h leads EC to transition into osteoblasts (OSB). During EC-to-OSB transition, cells underwent a change in cell morphology, from a cuboidal shape to a more elongated shape (Fig. 1A), that was accompanied by a significant increase in the expression of OSB markers alkaline phosphatase and osteocalcin (Fig. 1B). We designated the cells that are undergoing transition as EC-OSB hybrid cells, and examined whether conditioned media (CM) from EC-OSB hybrid cells can modulate PCa cell activities (Fig. 1C). EC-OSB-CM was found to significantly increase the migratory (Fig. 1C) and invasive (Fig. 1D) activities of C4-2B4 and PC3-mm2 PCa cells relative to control 2H11-CM. EC-OSB-CM was also found to significantly increase tumor cell survival in anchorage-independent growth in soft agar assay relative to control 2H11-CM (Fig. 1E), without having an effect on PCa cell proliferation (Supplementary Fig. 1A, B). Further, adding BMP4 directly to PCa cells as a control did not have effects on these activities (Fig. 1C – E). These results suggest that EC-OSB hybrid cells secrete factors that stimulate the migration, invasion and anchorage-independent growth of PCa cells (Fig. 1F).

**Figure 1.**
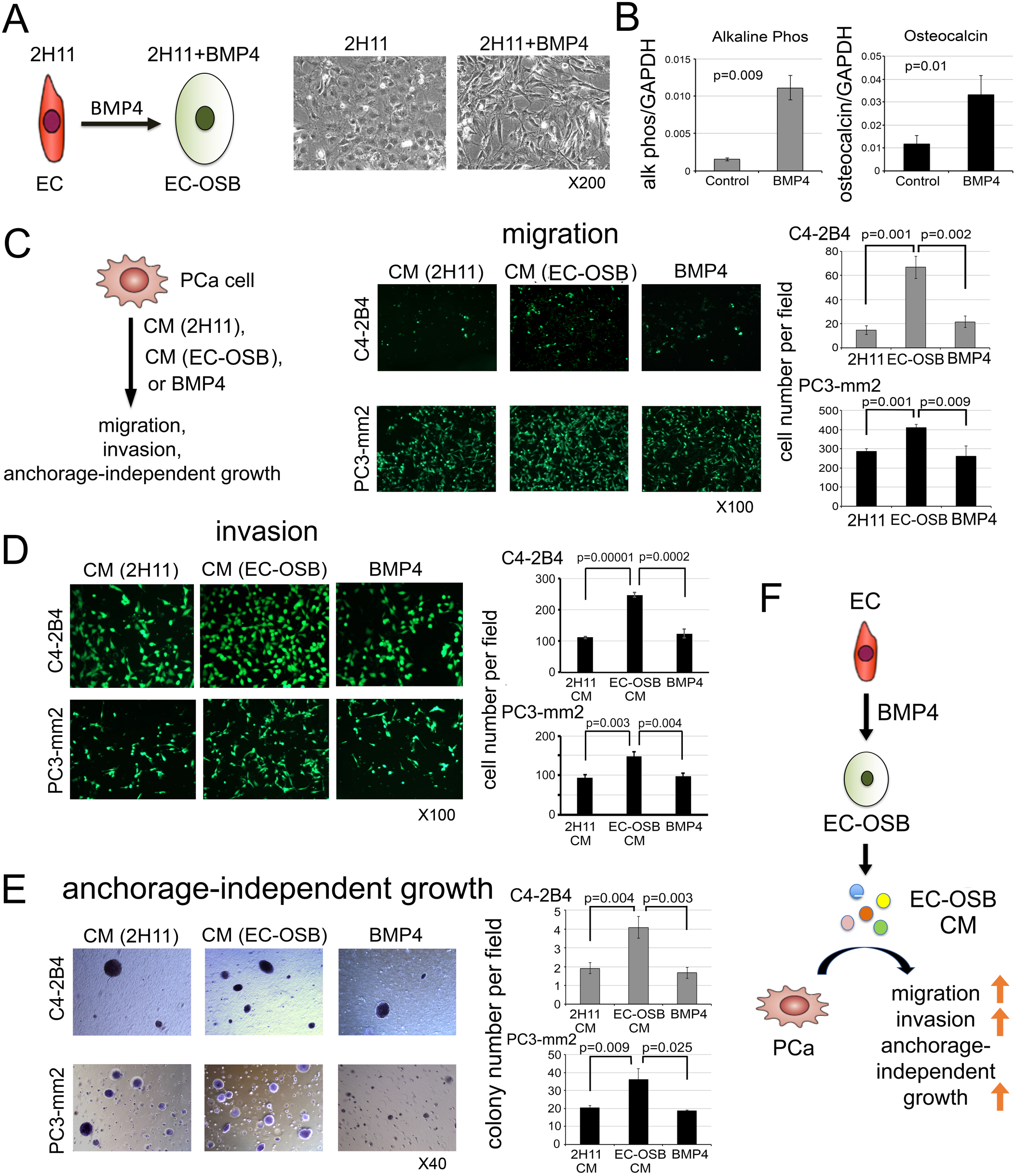
EC-OSB conditioned medium increases the migration, invasion and anchorage-independent growth of PCa cells. (A) Morphological change in 2H11 endothelial cells from EC to EC-OSB hybrid cells following BMP4 (100 ng/mL) treatment for 48 h. In this and subsequent experiments, 2H11 cells treated for 48 h with BMP4 are designated as EC-OSB hybrid cells. (B) Alkaline phosphatase activity and osteocalcin mRNA levels in 2H11 cells treated as in (A). (C-E) C4-2B4 or PC3-mm2 PCa cells were treated with conditioned medium (CM) from 2H11 or EC-OSB hybrid cells, or directly with BMP4 alone. The effects of these treatments on (C) migration, (D) invasion, and (E) anchorage-independent growth were measured. P values are from Student’s t-test. (F) Graphical summary. BMP4-mediated EC-to-OSB transition generates EC-OSB hybrid cells that secrete factors, which increase the migration, invasion and anchorage-independent growth of PCa cells. BMP4 alone does not elicit these activities in PCa cells.

### iTRAQ analysis identifies Tenascin C in EC-OSB-CM

We employed Isobaric Tags for Relative and Absolute Quantitation (iTRAQ) to compare the levels of proteins in EC-OSB-CM versus 2H11-CM (Fig. 2A) in triplicate samples. A total of 1414 proteins with p values of less than 0.017, corresponding to a 1% false discovery rate in Mascot search, were identified. Based on the iTRAQ ratio between two samples, 133 proteins were found to be upregulated (with changes in protein ratios of >1.22-fold) and 663 proteins downregulated (with changes <0.81-fold) in EC-OSB-CM versus control 2H11-CM (Supplementary Table 1). The numbers for significance (1.22 and 0.81) were calculated by taking all the numbers in the BMP4 average column and using the Easyfit program to find the distribution that best fit the data. From this we calculated a 2-sigma up/down number for the whole distribution. The top 15 proteins with the highest fold change were extracellular matrix proteins, including Tenascin C (TNC), CTGF, Versican and other secreted factors (Fig. 2B). TNC and TNC isoform 2 are the most promising candidate proteins due to a high number of significant peptides, 174 and 172 peptides, respectively, identified for isoform 1 and 2, respectively (Supplementary Table 1). Mouse TNC is a secreted protein of 2110 amino acids, containing 15 EGF-like domains in N-terminus (aa 174-621), 14 fibronectin type III domains (aa 625-1886), and a C-terminus fibrinogen domain. TNC isoform 2 differs from TNC in its absence of domain #10 of fibronectin type III domain (aa 1436-1526), resulting in a protein of 2019 amino acids, which is about 10 kDa less in molecular weight than TNC. Based on the comparable numbers of peptides identified in mass spectrometry during iTRAQ analysis, the levels of TNC and TNC isoform 2 secreted from 2H11 cells during BMP4 stimulation are predicted to be similar.

**Figure 2.**
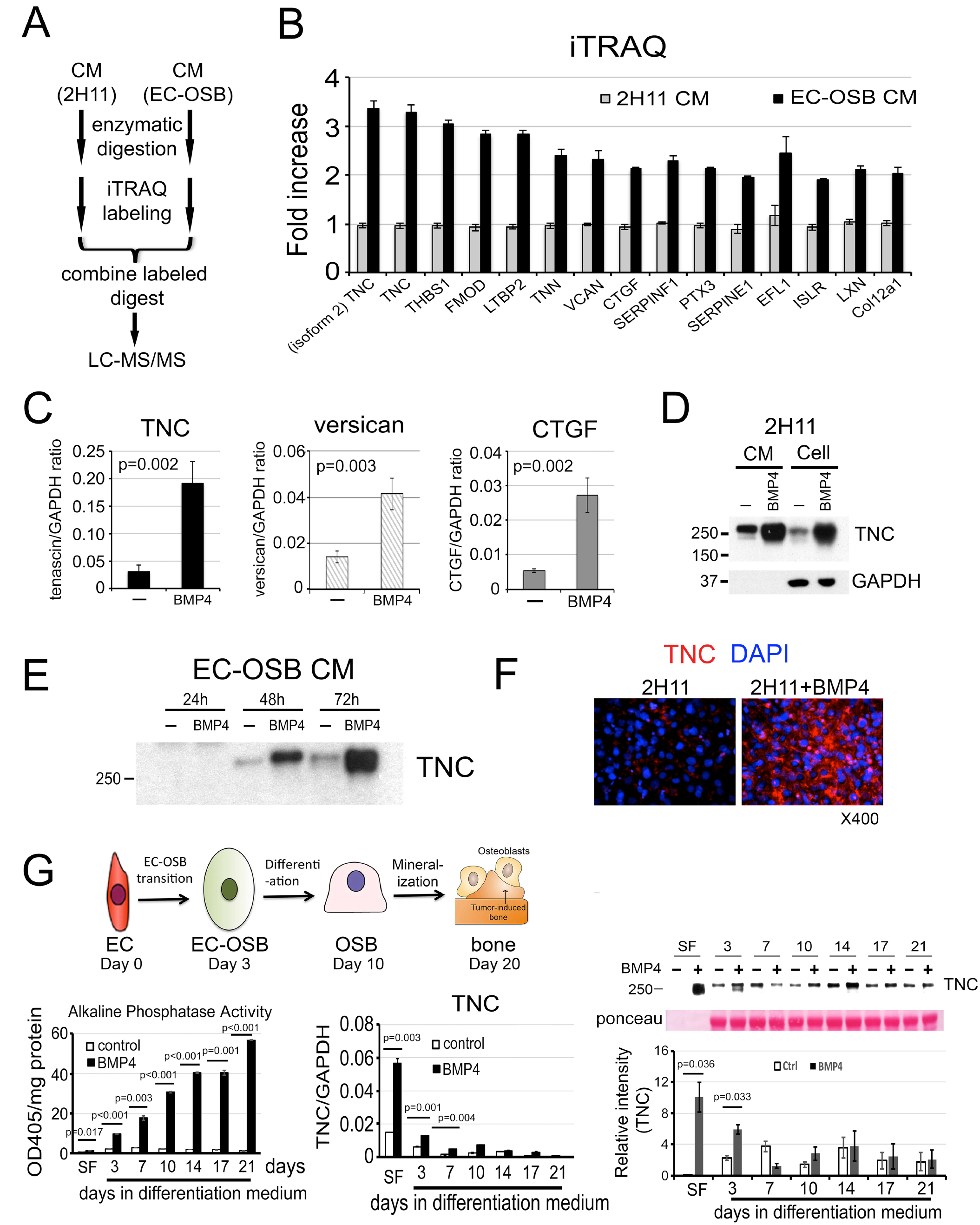
iTRAQ analysis identifies Tenascin C in EC-OSB conditioned medium. (A) Flow chart for iTRAQ procedure. (B) The top 15 proteins with the highest fold change in EC-OSB-CM compared to 2H11-CM using iTRAQ. (C) qRT-PCR for mRNA of Tenascin C (TNC), Versican and CTGF in 2H11 cells treated with or without BMP4 for 48 h. P values were by Student’s t-test. (D) TNC protein levels in CM or corresponding cells treated as in C. (E) Time course of TNC secretion into CM after BMP4 treatment. CM (10 microliter each) was used in each lane. (F) Immunofluorescence for the expression of TNC in 2H11 cells treated with or without BMP4 for 48 h. Nuclei were stained with DAPI. (G) Upper left, three phases of EC-to-OSB transition in 2H11 cells after treatment with BMP4. Lower left, time course of alkaline phosphatase protein activity in cell lysates. Lower middle, time course of TNC mRNA by qRT-PCR. Lower right, TNC protein levels in CM. Ponceau S, loading control. SF, serum-free.

Real-time RT-PCR confirmed the significant upregulation of TNC, Versican and CTGF mRNAs in BMP4-treated 2H11 cells (Fig. 2C). TNC also showed the highest abundance in mRNA levels, around 5-fold higher compared to those of Versican and CTGF, based on mRNA to GAPDH ratio (Fig. 2C). Because TNC was the most upregulated and abundantly-expressed protein, we selected TNC for further study. TNC is a large extracellular matrix protein with an apparent molecular mass of ∼250 kDa [7], which may correspond to the elevated 250 kDa protein band seen on SDS-PAGE of EC-OSB CM (Supplementary Fig. 1C, arrow). TNC protein was upregulated not only in the cell pellet of EC-OSB hybrid cells compared to 2H11 cells (Fig. 2D) but also accumulated in EC-OSB-CM from 48 to 72 h (Fig. 2E). Elevated expression of TNC was also observed in EC-OSB hybrid cells by immunofluorescence relative to control 2H11 cells (Fig. 2F).

BMP4-induced EC-to-OSB transition occurs in several stages [5]. Upon BMP4 treatment, 2H11 cells transition into OSB by day 3 in serum-free medium (SF) (Days 0-3, Transition Phase). EC-OSB hybrid cells then undergo OSB differentiation in differentiation medium and express bone matrix proteins (Days 3-10, Differentiation Phase), and finally mineralized by Day 20 (Mineralization Phase) (Fig. 2G, upper). Consistently, the osteogenesis marker alkaline phosphatase showed a time-dependent increase of enzyme activity during the Differentiation and Mineralization Phases (Fig. 2G, lower left). qRT-PCR of BMP4-treated 2H11 cells (Fig. 2G, lower middle) and Western blot of the corresponding CM (Fig. 2G, lower right) showed that TNC was highly upregulated during the Transition Phase. Note, depending on the gel running condition, TNC may appear as a broad single band or be resolved into a 250 kDa TNC doublet of distinct bands in western blot analysis. These results suggest that TNC is mainly expressed during the early phase of EC-to-OSB transition.

### Tenascin C is expressed in tumor-induced osteoblasts in PDX and human bone metastasis specimens

We examined whether TNC is induced in the EC-OSB hybrid cells of osteogenic tumors by immunohistochemical analysis. In the osteogenic MDA PCa-118b xenograft tumors, which is a PDX generated from the osteoblastic bone lesions of a patient with bone metastasis [8], TNC is highly expressed in tumor-induced OSBs (Fig. 3A). Similarly, in another osteogenic tumor model in which BMP4 was overexpressed in C4-2b cells [5], TNC is also expressed in the OSBs rimming the tumor-induced bone in the C4-2b-BMP4 tumors (Supplementary Fig. 2A). Further, we observed that TNC is mainly localized in the tumor-induced bone matrix that surrounds the metastatic PCa cells in human bone metastasis specimens (Fig. 3A and Supplementary Fig. 2B-C).

**Figure 3.**
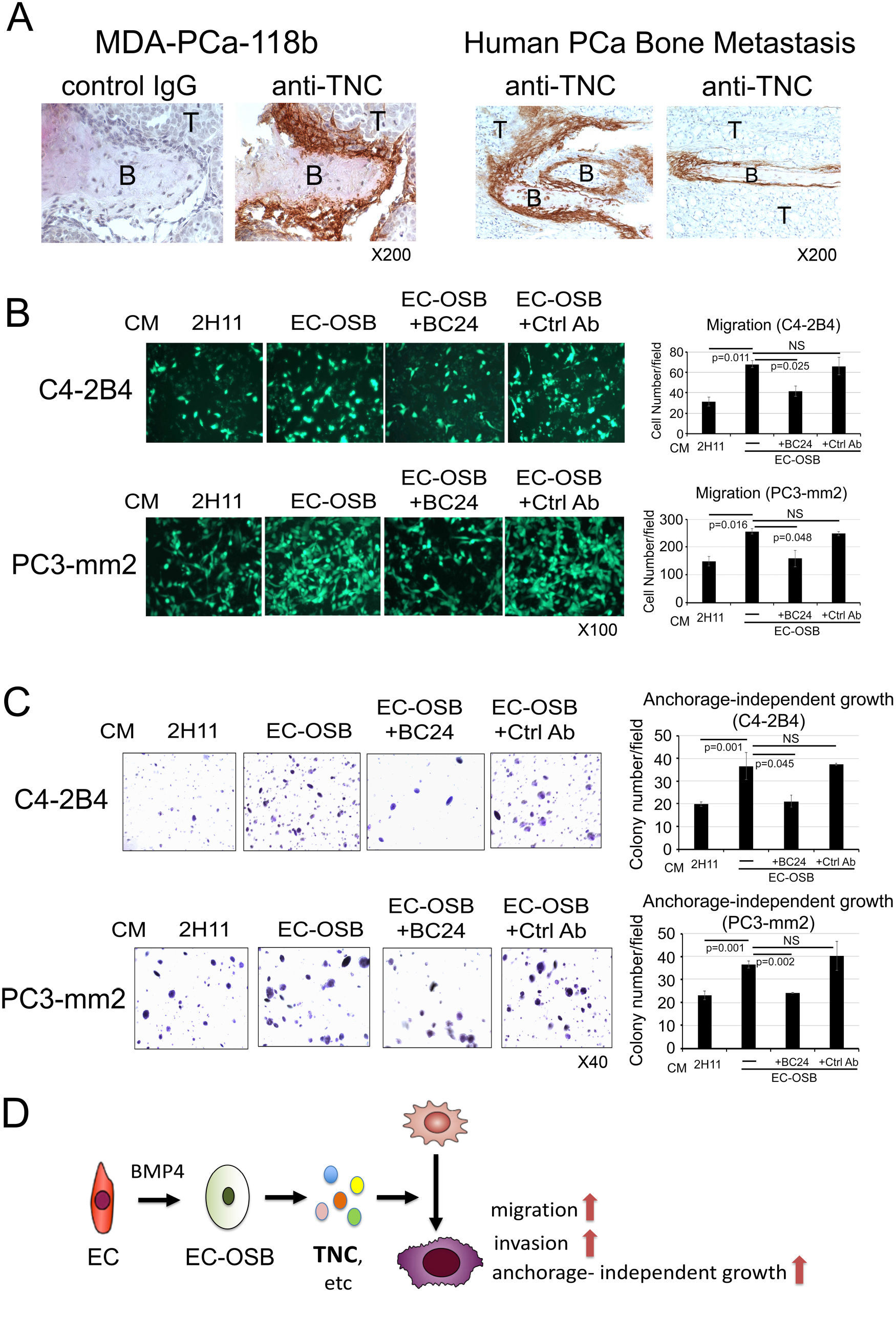
Tenascin C is expressed in tumor-induced osteoblasts in human bone metastasis specimens and Tenascin C neutralizing antibody reduces EC-OSB-CM-mediated migratory activity in PCa cells. (A) IHC of TNC in osteogenic tumors. TNC is found in EC-OSB hybrid cells rimming tumor-induced bone in MDA-PCa-118b PDX (left) and human PCa bone metastasis (right). B, bone; T, tumor. Magnification X200. Effects of TNC neutralizing antibody BC-24 on (B) migratory activity and (C) Anchorage-independent growth of EC-OSB-CM on C4-2B4 and PC3-mm2 cells. EC-OSB-CM were pre-incubated with anti-TNC mAb BC-24 (200 ng/ml) or control anti-AE1/AE3 mAb (200 ng/mL) overnight at 4°C prior to use in migration or anchorage-independent growth assays. P values were by Student’s t-test. NS, not significant. (D) Graphical summary. One of the factors secreted by EC-OSB hybrid cells is TNC, which promotes PCa migration, invasion and anchorage-independent growth.

### Tenascin C neutralizing antibody reduces migration stimulatory activity of EC-OSB-CM

Next, we incubated EC-OSB-CM with monoclonal antibody BC-24 [9], a TNC neutralizing antibody that recognizes the N-terminal epidermal growth factor-like sequence that is present in both TNC and TNC isoform 2. Antibody AE1/AE3 that recognizes cytokeratins was used as a control. Treatment with BC-24 antibody, but not control antibody, abrogated the migration stimulating activity (Fig. 3B) as well as anchorage-independent growth stimulating activity (Fig. 3C) in EC-OSB-CM for both C4-2B4 and PC3-mm2 cells. These results suggest that EC-OSB hybrid cells secrete TNC that promotes PCa migration, invasion and anchorage-independent growth (Fig. 3D).

### BMP4 increases Tenascin C in EC-OSB hybrid cells through Smad1-Notch/Hey1 pathway

Next, we examined the mechanism by which BMP4 increases TNC expression. We found that inhibition of BMP receptor kinase activity with 100 nM LDN193189 decreased BMP4-mediated phosphorylation of Smad1 and TNC protein levels in the CM (Fig. 4A). Although both Smad1 and Smad5 can be phosphorylated by BMP4 stimulation, Smad1 is expressed at a high level while Smad5 is low to undetectable in 2H11 cells (Supplementary Fig. 2D). Incubation of 2H11 cells with LDN alone did not have an impact on total Smad1 levels as they were comparable to those in control cells (Fig. 4A). Knockdown of Smad1 in 2H11 cells by shRNA in 2H11-shSmad1#1, #3, and #5 clones (Supplementary Fig. 2D) significantly reduced BMP4-stimulated pSmad1 (Fig. 4B) and led to a significant reduction in BMP4-stimulated TNC expression at both the TNC mRNA and TNC protein levels in the corresponding CM (Fig. 4C). These results suggest that TNC expression in 2H11 cells is mediated through BMP4-activated Smad1 signaling.

**Figure 4.**
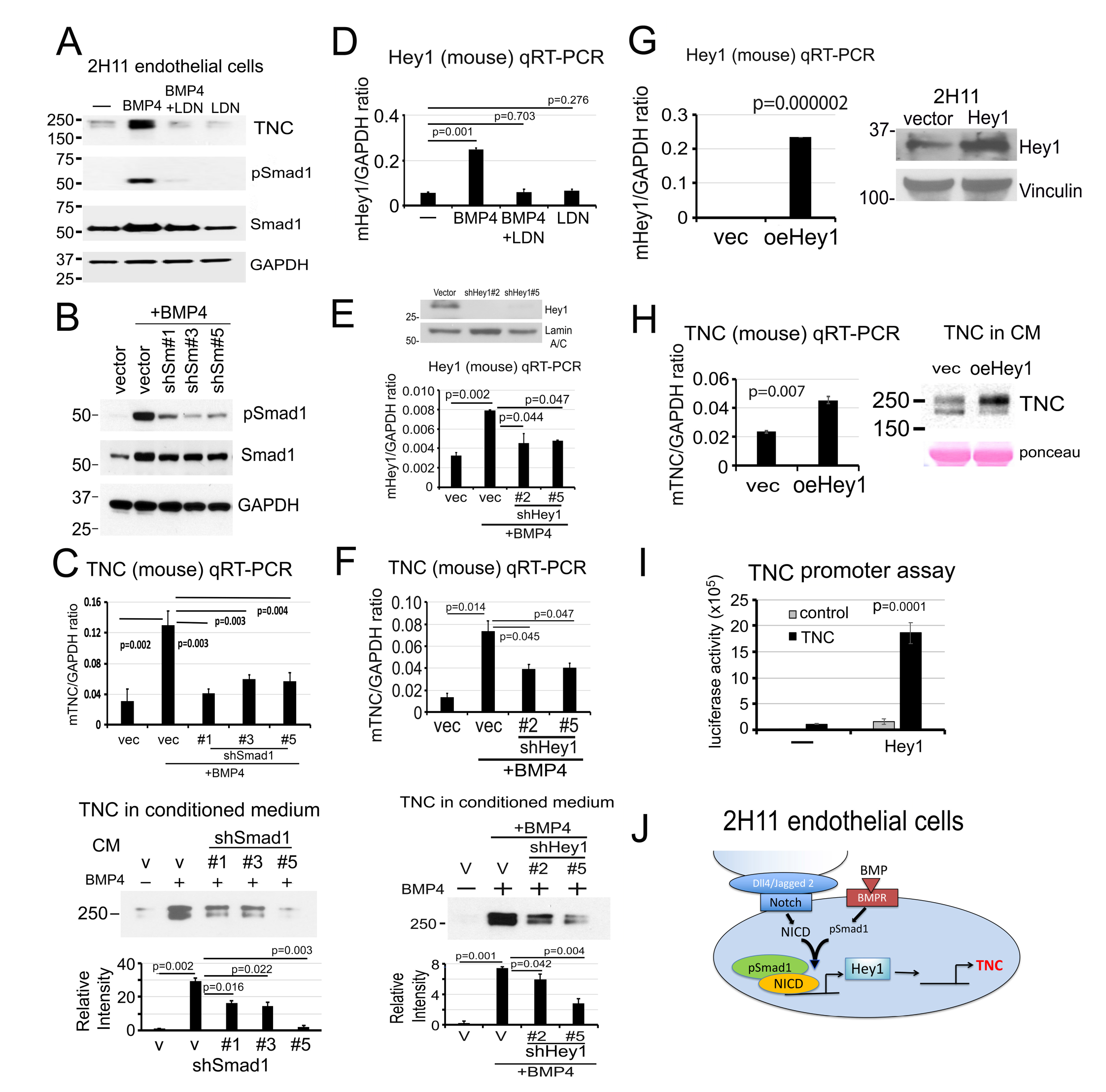
BMP4 stimulates Tenascin C expression in 2H11 cells through Smad1-Notch/Hey1 pathway. (A) TNC protein levels in the CM and pSmad1 levels in 2H11 cells treated as indicated. LDN193189, inhibitor of BMP receptor kinase activity. GAPDH, loading control. (B) pSmad1 levels in Smad1 knockdown clones 2H11-shSmad1#1, #3, and #5 treated as indicated. (C) TNC mRNA in cells (upper) and TNC protein in CM (lower) in Smad1 knockdown clones in (B). P values were by Student’s t-test. (D) Hey1 mRNA levels in 2H11 cells treated as indicated. (E) Hey1 protein (upper) and mRNA (lower) levels in 2H11-shHey1#2 and #5 clones treated as indicated. (F) TNC mRNA in cells (upper) and TNC protein in CM (lower) in Hey1 knockdown clones in (E). (G) Hey1 mRNA (left) and protein (right) levels in 2H11-oeHey1 cells. Hey1 protein was extracted from cells by immunoprecipitation followed with western blot. (H) TNC mRNA (left) and protein in CM (right) in 2H11-oeHey1 cells. (I) TNC promoter-luciferase construct was co-transfected with Hey1 expression vector in 2H11 cells, and the luciferase reporter activity was measured. (J) Graphical summary. BMP4-stimulates TNC expression through pSmad1-Notch/Hey1 pathway in 2H11 endothelial cells.

We previously showed that pSmad1 forms a complex with Notch intracellular domain (NICD) to upregulate Hey1, one of the most highly-expressed transcription factors during EC-to-OSB transition [6]. LDN193189 treatment abrogated BMP4-induced Hey1 mRNA expression (Fig. 4D), consistent with a role of pSmad1 in Hey1 regulation. Knockdown of Hey1 in 2H11-shHey1#2 and #5 clones (Fig. 4E) significantly reduced BMP4-stimulated TNC expression at both the mRNA and protein levels (Fig. 4F). These results suggest that upregulation of Hey1 is necessary for BMP4-mediated TNC expression. Next, we overexpressed Hey1 in 2H11 cells (oeHey1) (Fig. 4G) and showed that TNC mRNA and protein levels were significantly increased in 2H11-oeHey1 cells compared to control vector-transfected cells (Fig. 4H). These results suggest that Hey1 is necessary and sufficient for TNC expression in 2H11 cells. Co-transfection of a TNC promoter-luciferase construct together with an Hey1 expression vector into 293T cells resulted in a significant increase in TNC promoter activity only in the presence of Hey1 (Fig. 4I), suggesting that Hey1 has a direct effect on TNC expression in 2H11 cells. These results support that BMP4-induced TNC expression during EC-to-OSB transition is largely mediated through a pSmad1-Notch/Hey1 signaling pathway (Fig. 4J).

### Tenascin C is expressed in prostate cancer cells with high metastatic potential

Although TNC is mainly secreted by stromal EC-OSB hybrid cells, TNC is also reported to be expressed in highly metastatic breast cancer cells [10], possibly due to epithelial-to-mesenchymal transition. We found that the highly metastatic PCa cells, i.e., PC3, PC3-mm2 and DU145 cells, express high levels of TNC mRNA (Fig. 5A, upper) and secrete high levels of TNC in their CMs (Fig. 5A, lower). The less aggressive PCa cells, i.e., LNCaP, C4-2B, C4-2B4, 22RV1 and VCaP cells, express low levels of TNC (Fig. 5A and Supplementary Fig. 9A-B).

**Figure 5.**
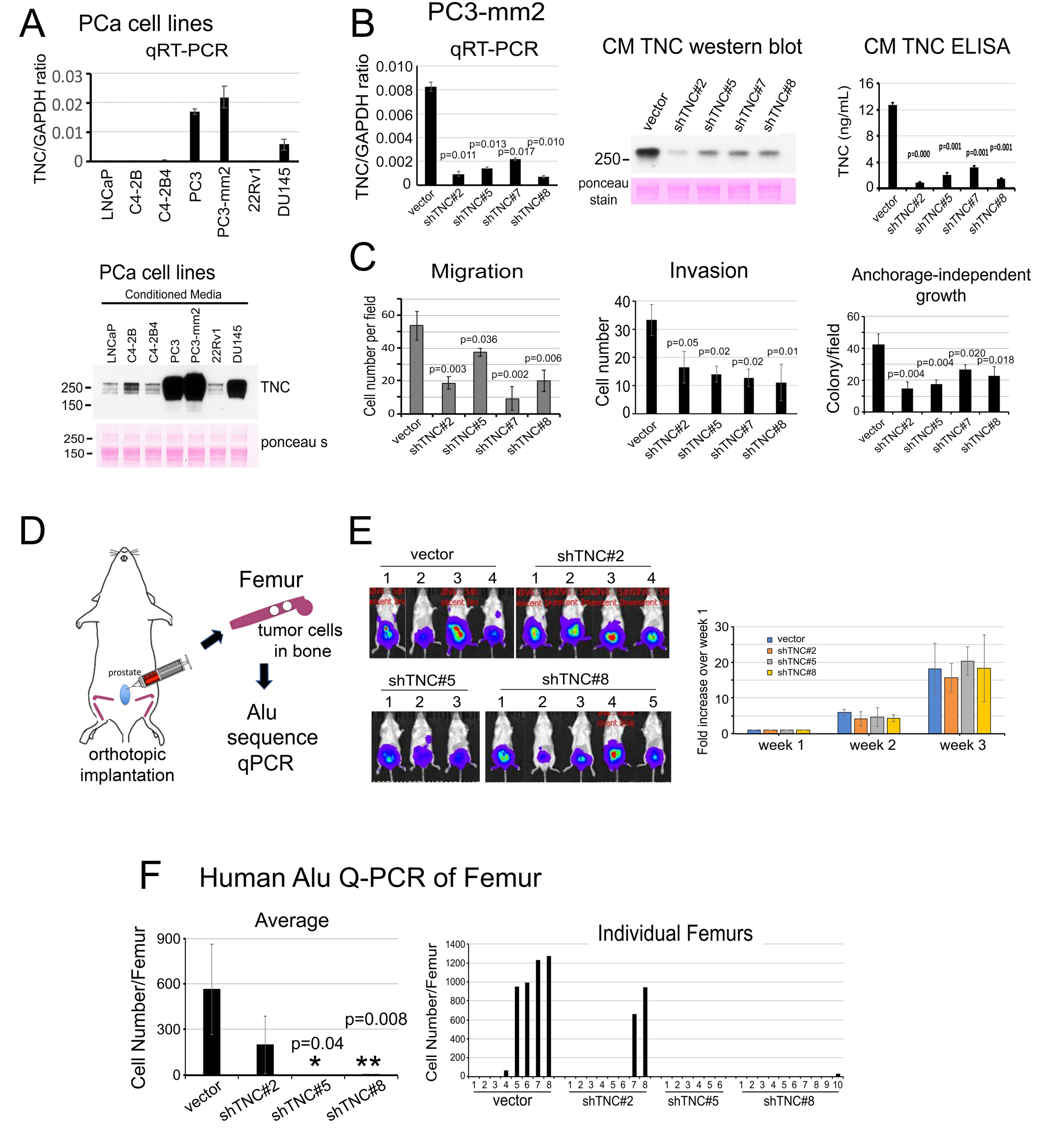
Knockdown of Tenascin C decreases the migration, invasion, anchorage-independent growth of PC3-mm2 cells in vitro and the metastasis of PC3-mm2 cells to bone in vivo. (A) qRT-PCR for TNC mRNA in PCa cell lines (upper). Western blot of TNC protein in CM from PCa cell lines (lower). (B) qRT-PCR (left) for TNC mRNA in PC3-mm2-shTNC#2, #5, #7 and #8 clones. CMs from PC3-mm2-shTNC clones were analyzed by Western blot (middle) and ELISA (right). (C) Migration, invasion, and anchorage-independent growth of PC3-mm2-shTNC clones in (B). (D) Experimental scheme. PC3-vector, shTNC#2, #5, or #8 cells were injected orthotopically into mouse prostate. After 3 weeks, femurs were dissected and total DNA prepared. Tumor cells that metastasized to bone were determined by human *Alu* sequence qPCR. (E) Bioluminescence of tumors in the various mouse groups at 3 weeks post-injection. Time course of tumor growth based on bioluminescence is shown. (F) Quantification of tumor cells that have metastasized to bone. Left, average number of tumor cells metastasized from prostate to bone. Right, number of tumor cells detected in individual legs in control mice and PC3-mm2-shTNC#2, shTNC#5, or shTNC#8 injected mice. P values were by Student’s t-test.

Interestingly, as also observed in EC-OSB conditioned media (Fig. 2 and 4), TNC and TNC isoform 2 appeared in concert in the 250 kDa TNC doublet, and their levels were similarly altered in the various PCa cells (Fig. 5A).

### Knockdown of Tenascin C decreases the migration, invasion and anchorage-independent growth of PC3-mm2 cells in vitro

Next, we employed shRNA to knockdown TNC in PC3-mm2 cells. In PC3-shTNC#2, #5, #7 and #8 clones, TNC levels were reduced at the mRNA (Fig. 5B, left) and protein levels in the corresponding CM by western blot (Fig. 5B, middle) and ELISA (Fig. 5B, right).

Knockdown of TNC in these PC3-shTNC clones was found to significantly decrease migration, invasion, and anchorage-independent growth (Fig. 5C and Supplementary Fig. 3A-C) relative to the vector control PC3-mm2 cells. Similarly, knockdown of TNC in C4-2b cells decreased migration, invasion and anchorage-independent growth but not proliferation of C4-2b-shTNC clones (Supplementary Fig. 4A-E).

### Knockdown of Tenascin C reduces the metastasis of PC3-mm2 cells to bone in vivo

To examine whether knockdown of TNC in PC3-mm2 cells has an impact on the metastasis of PC3-mm2 cells to bone, PC3-vector, PC3-shTNC#2, #5, or #8 cells were injected orthotopically into the mouse prostate (Fig. 5D). We found that knockdown of TNC had little effect on tumor growth in the prostate as monitored by bioluminescence (Fig. 5E). Because the number of tumor cells that metastasized to bone were too low to be detected by bioluminescence, we quantified the tumor cells that have metastasized to bone using human-specific *Alu*-qPCR [11]. We first established an *Alu* PCR standard curve by using DNA from PC3-mm2 cells.

Then, we compared the *Alu* PCR signals obtained from each femur against the *Alu* standard curve to determine the number of PC3-mm2 cells present in each femur. We found that knockdown of TNC in PC3-mm2 cells decreased the number of PC3-shTNC cells that metastasized from the prostate to bone compared to vector-transfected control cells (Fig. 5F, left). We detected tumor cells in 5 of 8 legs analyzed in vector control mice, 2 of 8 legs in PC3-shTNC#2, 0 of 6 legs in PC3-shTNC#5, and 1 of 10 legs in PC3-shTNC#8 mice (Fig. 5F, right). These results suggest that TNC increases the metastasis of PC3-mm2 from the prostate to the bone. Similar results were obtained when the experiment was repeated in a separate group of mice (Supplementary Fig. 3D-H). Together, these results suggest that TNC plays a role in the metastatic potential of PC3-mm2 cells.

### Tenascin C increases the migration and anchorage-independent growth of C4-2B4 and PC3 cells in vitro

Next, we examined whether TNC is sufficient to increase the metastatic potential of PCa cells. We examined the effects of TNC on C4-2B4 cells by incubation with recombinant human TNC, containing amino acid Gly23-Pro623 (the EGF-like domains) [12]. The addition of TNC led to a significant increase in the migration (Fig. 6A) and anchorage-independent growth (Fig. 6B) of C4-2B4 cells compared to medium only or BSA control. We next expressed the first 625 amino acids of TNC in C4-2b cells (C4-2b-TNC), and showed that the exogenously expressed TNC was detected at both the mRNA and protein levels (Fig. 6C). C4-2b-TNC cells exhibited a significant increase in migration, invasion and anchorage-independent growth compared to C4-2b-vector control cells (Fig. 6D – F). Overexpression of TNC had no effect on C4-2b-TNC cell proliferation in vitro compared to C4-2b-vector cells (Supplementary Fig. 5A). Similarly, overexpression of TNC in PC3 cells increased migration, invasion and anchorage-independent growth, but not cell proliferation in PC3-TNC cells (Supplementary Fig. 6A-E). These results suggest that TNC confers migratory and survival properties to both C4-2B4 and C4-2b PCa cells, which are sublines derived from LNCaP cells [13-16], as well as to PC3 cells.

**Figure 6.**
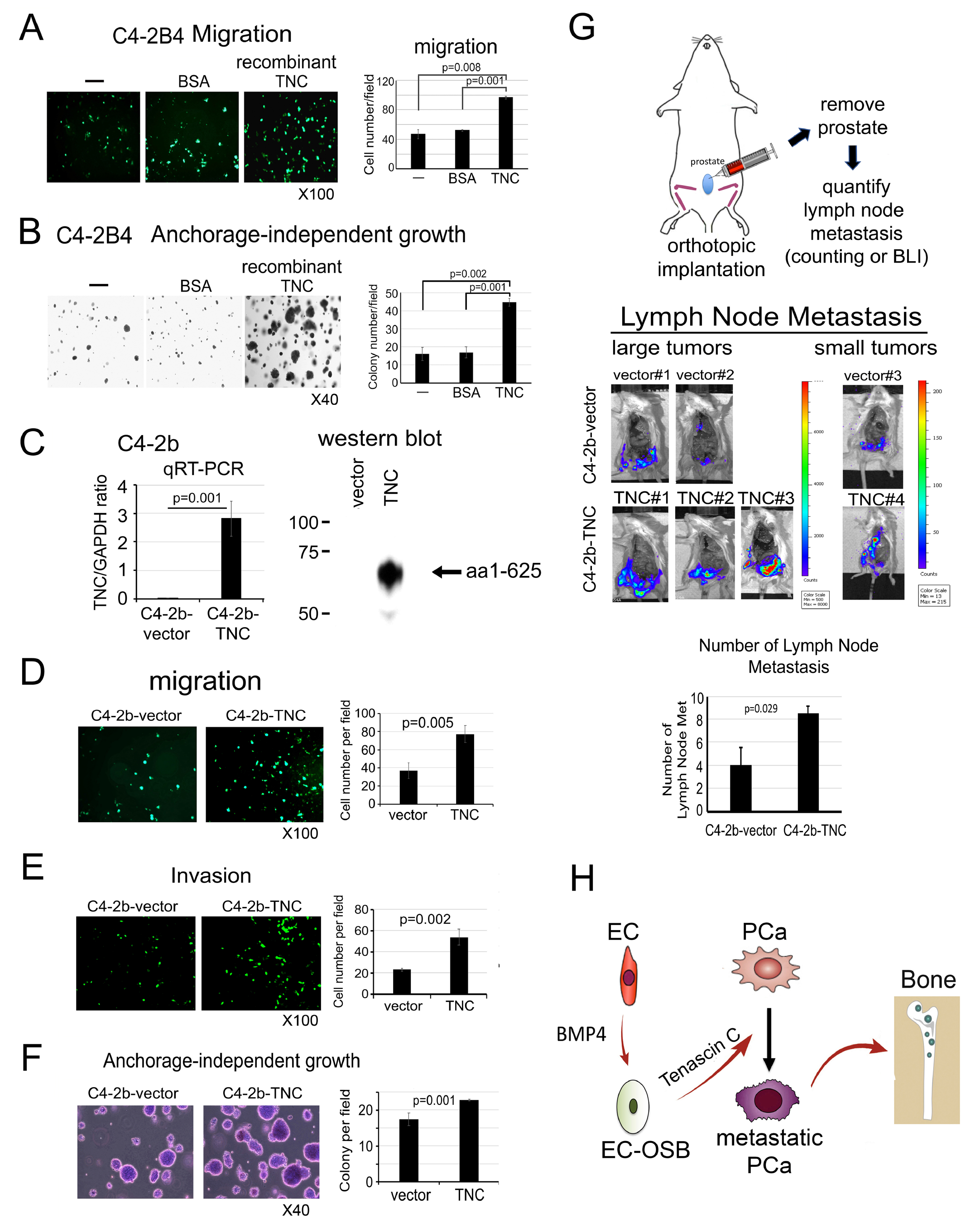
Tenascin C increases the migration and anchorage-independent growth of C4-2b cells in vitro and metastasis to lymph nodes in vivo. C4-2B4 cell migration (A) and anchorage-independent growth in vitro (B) in response to recombinant human TNC (5 µg/m), BSA (5 µg/m) or media alone. (C) Overexpression of N-terminal domain (1 – 625 aa) of TNC in C4-2b cells by qRT-PCR and western blot. (D-F) Migration, invasion and anchorage-independent growth of cells in (C). (G) C4-2b-vector or C4-2b-TNC cells were injected orthotopically into mouse prostate (upper). Tumor growth was monitored by bioluminescence. When tumors reached certain sizes, they were removed and lymph node metastases were counted (middle). Average number of lymph node metastasis was significantly increased in C4-2b-TNC cells relative to C4-2b-vector control cells (lower). P values were by Student’s t-test. (H) Graphical summary. Endothelial cells (EC) undergo EC-to-OSB transition by BMP4. EC-OSB hybrid cells then secrete TNC that increases metastatic potential of PCa cells.

### Tenascin C increases the metastasis of C4-2b cells to lymph nodes in vivo

To examine whether overexpression of TNC in C4-2b cells has an impact on the metastasis of C4-2b cells, C4-2b-vector and C4-2b-TNC cells were injected orthotopically into the prostate of intact mouse (Fig. 6G). Because C4-2b is a cell line with low metastasis potential [16], we examined the effect of TNC on the metastasis of C4-2b from prostate to lymph nodes.

Although tumors were inoculated in 10 mice each, 2 large and 1 small tumors were detected in C4-2b-vector and 3 large and 1 small tumors were detected in C4-2b-TNC injected mice. There was no significant difference in C4-2b-TNC tumor growth in the prostate compared to C4-2b-vector as measured by bioluminescence imaging and tumor weight (Supplementary Fig. 5B-E). After removing the tumors, the number of lymph node metastases from the three tumors from C4-2b-vector and four tumors from C4-2b-TNC were compared. We found that overexpression of TNC in C4-2b cells led to an increase in the number of lymph node metastasis compared to vector-transfected cells (Fig. 6G, Supplementary Fig. 5F). These results suggest that TNC increases the metastasis of C4-2b cells from prostate to lymph nodes. Together, these results support that EC-to-OSB transition generates a tumor microenvironment enriched with TNC that enhances metastatic progression of PCa (Fig. 6H).

### Tenascin C promotes prostate cancer cell migration through α5β1-YAP/TAZ pathway

Next, we examined the mechanism by which TNC promotes PCa cell migration. TNC signals through integrins, including α2β1 [17], α5β1 [18], α8β1 [19], α9β1 [20, 21], and αv [22-24]. Using human integrin primers, we found that the mRNA levels of α5, α11, αv, β1, and β5 are expressed at higher levels than other integrins in C4-2b cells (Fig. 7A). We then performed qRT-PCR for the mRNA levels of those candidate TNC receptors in C4-2b-vector and C4-2b-TNC cells. Interestingly, expression of TNC in C4-2b cells led to a decrease in the mRNA levels of α5 and β1 (Fig. 7B), suggesting that α5 and β1 may be the major integrins for TNC in C4-2b cells. Indeed, when C4-2b-TNC cells were treated with integrin α5β1 neutralizing antibody (Volociximab), we found that blocking integrin α5β1 significantly decreased TNC-mediated migration (Fig. 7C).

**Figure 7.**
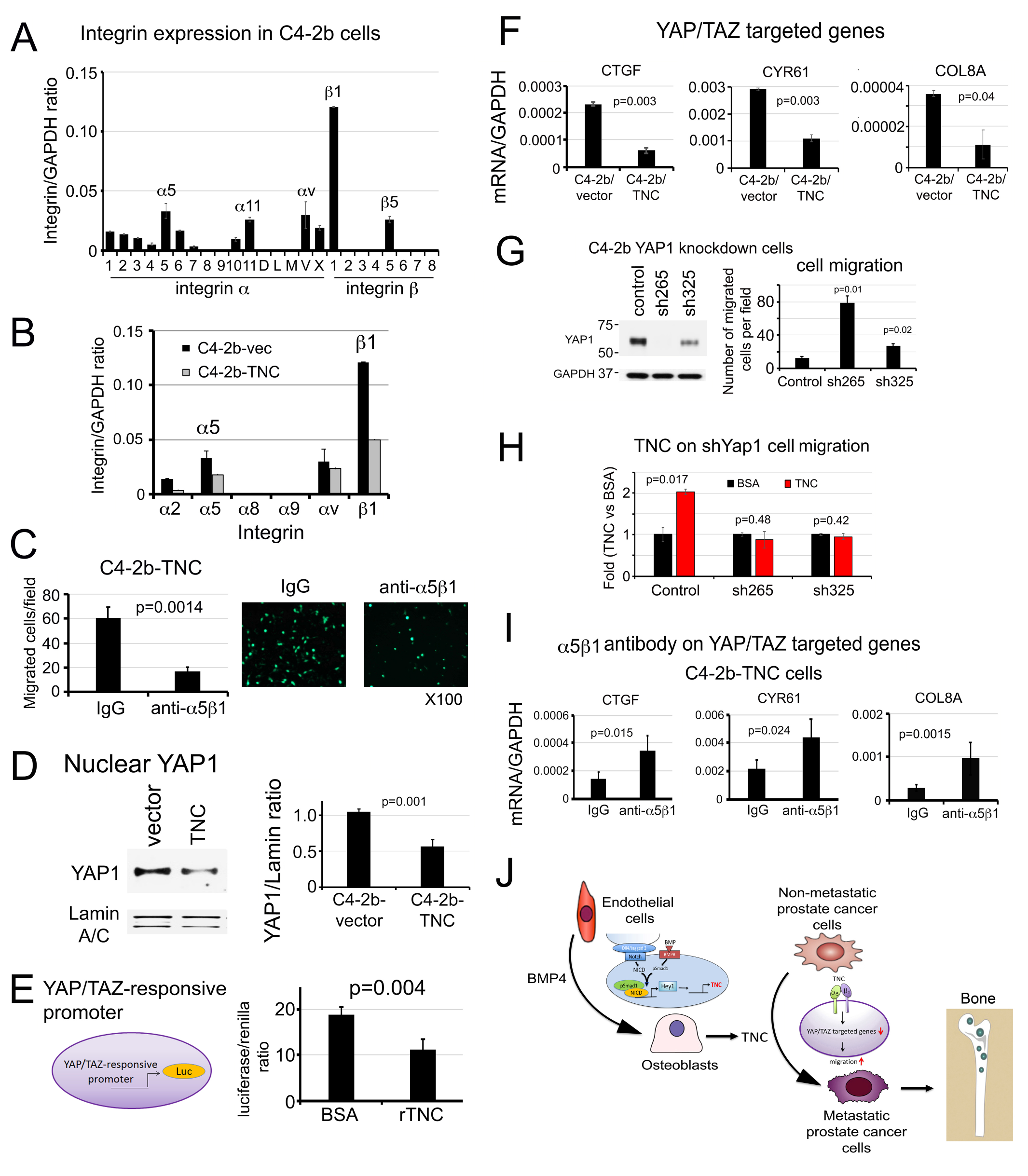
Tenascin C promotes prostate cancer cell migration through α5β1-YAP1 pathway. (A) Integrin expression in C4-2b cells. (B) Expression of TNC binding integrins in C4-2b-vector and C4-2b-TNC cells. (C) Cell migration in C4-2b-TNC cells incubated with anti-α5β1 integrin neutralizing antibody or control IgG. (D) Nuclear YAP1 protein levels in cells in (B) by Western blot. Lamin A/C, loading control. (E) YAP/TAZ responsive promoter-driven luciferase activity in C4-2b cells incubated with recombinant TNC (rTNC) or BSA control. (F) qRT-PCR for mRNAs of YAP/TAZ target genes in C4-2b-vector or C4-2b-TNC cells. (G) Cell migration in YAP1 knockdown C4-2b-shYAP1-265 or -325 clones. (H) Cell migration in YAP1 knockdown clones in response to recombinant TNC or BSA control. (I) YAP/TAZ target gene expression in C4-2b-TNC cells following incubation with α5β1 integrin antibody or IgG control. P values were by Student’s t-test. (J) Graphical summary. PCa secreted BMP4 induces endothelial cells to undergo EC-to-OSB transition as part of stromal reprogramming, during which TNC is expressed through a Smad1-Notch/Hey1 signaling pathway. TNC in turn enhances PCa cell migration and invasion through α5β1integrin-mediated inhibition of the YAP1 pathway, leading to metastatic progression of PCa in bone.

Various mechanisms have been reported for the effect of TNC on cell migration, including YAP inhibition [21], Src activation and FAK phosphorylation [22], and activation of Notch and Wnt signaling [10, 17]. Among these possible mechanisms, we found that YAP/TAZ inhibition is involved in TNC-mediated cell migration as follows. Exogenous expression of TNC in C4-2b cells led to a decrease in the levels of nuclear YAP1 protein (Fig. 7D). Interestingly, treatment of C4-2b cells with recombinant TNC also led to a decrease in YAP/TAZ-responsive promoter activity (Fig. 7E). Consistent with lowered nuclear YAP1 levels in C4-2b-TNC cells, there was a significant decrease in the expression of several YAP/TAZ target genes, including CTGF, CYR61, and COL8A (Fig. 7F), thus supporting that TNC downregulates YAP1 nuclear levels and its activity in the nucleus. Knockdown of YAP1 in C4-2b cells by shRNA led to an increase in cell migration in C4-2b-shYAP1-265 and C4-2b-shYAP1-325 clones (Fig. 7G). Treatment of C4-2b-shYAP1 cells with recombinant TNC did not increase migration relative to BSA (Fig. 7H), suggesting that TNC-mediated migration is through YAP1. Furthermore, when C4-2b-TNC cells were incubated with anti-α5β1 neutralizing antibody, the inhibitory effects of TNC on CTGF, CYR61, and COL8A expression were de-repressed (Fig. 7I). Similar studies were performed in PC3 cells. We found that α5β1 integrins are also expressed in PC3 cells (Supplementary Fig. 7A). We examined whether α5β1 also plays a role in TNC-mediated YAP1 regulation by treating PC3 cells with α5β1 neutralizing antibody. We found that α5β1 neutralizing antibody blocked TNC-mediated migration (Supplementary Fig. 7B) and increased the expression of YAP/TAZ target genes CTGF, CYR61 and COL8A in PC3 cells (Supplementary Fig. 7C). Knockdown of YAP1 in PC3 cells led to a significant increase in cell migration (Supplementary Fig. 7D) and abrogated TNC’s effect on cell migration in the PC3-shYAP1 clones (Supplementary Fig. 7E). Together, these results suggest that TNC promotes PCa cell migration, at least in part, through integrin α5β1-YAP/TAZ pathway (Fig. 7J).

## Discussion

We have identified a novel mechanism whereby PCa-induced EC-to-OSB transition leads to changes in the extracellular milieu that enhances the metastatic potential of PCa (Fig. 7J). We showed that BMP4-mediated Smad1-Notch/Hey1 signaling pathway leads to TNC secretion from 2H11 endothelial cells. TNC in turn enhances PCa cell migration through α5β1integrin-mediated inhibition of the YAP1 pathway. These insights into stromal reprogramming-mediated metastatic progression may provide strategies to improve therapies for bone metastasis.

Previous studies by Sung et al. [25] showed that prostate stromal fibroblasts co-evolve with cancer cells through reciprocal cancer-stroma interaction in an in vitro co-culture system, suggesting that tumor-stromal interaction is a dynamic process. Recently, tumor-induced stromal reprogramming was shown to occur in many different cancer types, including pancreatic [26], liver [27], breast cancers [28], and tumor draining lymph nodes [29]. Tumor cells were also found to reprogram their immune microenvironment [30]. Knowledge of such tumor-induced stromal changes will lead to therapies targeting the altered stromal components to improve therapy outcomes.

In this study, we used BMP4 to stimulate EC-to-OSB transition. While BMP6 [4] and BMP7 [31] were also shown to contribute to the bone-forming phenotype of PCa bone metastasis, we focused on BMP4 for the following reasons. First, we identified BMP4 as a factor that is secreted by the osteoblastic PDX MDA PCa 118b, and showed that BMP4 promotes tumor growth through osteogenesis [3]. Second, we validated this observation in humans, demonstrating high BMP4 expression in bone metastases but not primary tumors [5]. Third, Nordstand et al. [32] showed that BMP4 expression in human PCa bone metastases correlated with increased bone formation. It is likely that other BMP family proteins may also stimulate EC-to-OSB transition.

Our mechanistic studies revealed that BMP4 induced TNC expression during EC-to-OSB transition is mediated through the Smad1-Hey1 pathway. Although Hey1 transfection in HEK293 cells increases Tnc promoter activity, Hey1 may not directly bind and regulate expression of TNC as TNC promoter does not contain a Hey1 binding site and Hey1 is a transcriptional repressor [33]. TNC promoter is reported to contain GATA4 or GATA6 binding sites and Hey1 has been shown to interact with GATA6 [33]. Thus, it is possible that Hey1 increases TNC promoter activity through interaction with GATA6. The precise mechanism remains to be determined.

Previous studies in human GBM cells by Miroshnikova et al. [34] have shown that HIF1α binds directly to a putative hypoxia-response element in the TNC promoter, resulting in increased expression of TNC that increases ECM stiffness and mechanosignalling. HIF1α is also known to activate the Notch pathway [35]. Under hypoxic conditions, HIF1α is recruited to the Notch-responsive promoter through binding with Notch intracellular domain [36].

Interestingly, Notch signaling is also shown to affect the hypoxic response via regulation of HIF2α [37]. However, in our recent RNAseq analysis of BMP4 treated 2H11 cells during EC-to-OSB transition [6], we did not find upregulation of HIF1α or HIF2α during this process. Thus, the HIF1 pathway is not involved in the regulation of TNC expression during EC-to-OSB transition.

TNC has been shown to be an organ-specific driver of breast cancer metastasis. Oskarsson et al. [10] showed that TNC initiates and sustains lung colonization of breast tumor cells. They also showed that TNC knockdown does not hinder mammary tumor growth but rather restricts the progression of lung micro-metastases [10]. We also found that TNC does not affect PCa proliferation but endows PCa cells with migratory and survival ability in vitro and increases their metastasis to lymph nodes and bone in vivo. In support of our observation, San Martin et al. [20] showed that TNC enhances PCa cell migration towards bone tissues in vitro. TNC is upregulated in reactive stroma in primary PCa [38] and is also shown to correlate with PCa progression [39]. Together, these observations strongly support a role of stromal TNC in PCa progression.

We found that TNC signals through α5β1 integrins in C4-2b and PC3 PCa cells. Integrins are composed of a large family of 18 α subunits and 8 β subunits, which are able to generate 24 different integrins [40]. Integrins are widely expressed in many different cell types with various subunit combinations. The regulation of the expression of various integrin subunits in different cells are unknown. In previous studies, we found that C4-2B4, PC3-mm2 and MDA PCa-118b xenografts have different integrin expression profiles [41]. In our current studies, we found that the expression of integrins in C4-2B and PC3 cells are also somewhat different from those reported for C4-2B4 and PC3-mm2 cells, respectively. TNC has been shown to signal through various integrins, including α2β1 [17], α5β1 [18], α8β1 [19], α9β1 [20, 21], and αv [22-24]. In studies by San Martin et al. [20], TNC was shown to signal through α9β1 integrin, which is expressed in VCaP but not LNCaP, C4-2b, PC3 or 22RV1 cells. Thus, it is likely that TNC may signal through different integrins expressed in different PCa cells. Because PCa is heterogeneous, the integrins expressed in human bone metastasis specimens may be different in different PCa patients and remain to be determined.

Our studies showed that TNC promotes cell migration through inhibition of YAP/TAZ signaling pathway. YAP/TAZ are known as the nuclear transducers of the Hippo pathway during development [42]. Interestingly, Dupont et al. [43] showed that YAP/TAZ has a unique role as nuclear relays of mechanical signals from cellular microenvironment, acting independently from the Hippo/LATS cascade [43]. Cells respond to soft matrices or reduce cell spreading by inhibiting YAP/TAZ activity [43]. We found that TNC inhibits YAP/TAZ activity, providing a mechanism by which TNC enhances cancer metastasis. How signaling information generated by TNC reduces YAP/TAZ activity is not clear. YAP/TAZ activities are regulated by multiple mechanisms [44]. Recently, it was reported that post-translational modification of YAP1 by proline hydroxylation also plays a role in YAP/TAZ activity in PCa cells [45]. Whether TNC-mediated inhibition of YAP/TAZ activity is through proline hydroxylation is unknown. The signal transduction pathway linking TNC/integrin-mediated signaling to YAP/TAZ inhibition requires further investigation. Although TNC has been shown to increase FAK and Src phosphorylation in breast cancer cells [22], we did not find a consistent regulation of FAK and Src phosphorylation by TNC in PCa cells. Our results showed that overexpression of TNC in C4-2b cells did not increase levels of pY397-FAK and pY416-Src (Supplementary Fig. 8A), although downregulation of TNC in PC3 cells showed a trend of decrease in FAK and Src phosphorylation (Supplementary Fig. 8B). Thus, TNC likely uses different signaling pathways in different tumor cells to regulate cellular activity.

Because androgen receptor (AR) plays a critical role in PCa, it is interesting to consider whether AR has an impact on TNC levels in PCa cells. We found that TNC levels of AR-negative PC3 and PC3-mm2 cells were higher as compared to those in AR-expressing LNCaP, C4-2b, 22RV1 and VCAP cells (Supplementary Fig. 9). However, the TNC levels in another AR-negative line DU145 were found to be similar to those in AR-expressing C4-2b and C4-2B4 cells (Supplementary Fig. 9B). We then examined whether TNC expression correlated with levels of the AR V7 splice variant in these cells. A high level of AR V7 was detected in 22RV1 cells, followed by lower levels in VCaP cells, and low to undetectable levels in LNCaP, C4-2b and C4-2B4 cells (Supplementary Fig. 9B). Notably, we found that a similarly low level of TNC was found in 22RV1 (high AR V7) versus LNCaP (undetectable AR V7) cells (Supplementary Fig. 9B). Thus, neither AR nor AR V7 levels appear to impact TNC levels in PCa cells.

That tumor cells are able to modify tissue stroma to further promote their malignant progression suggests that targeting the tumor-induced changes in the tumor microenvironment should be included as an integral part of treatment strategies. TNC is downregulated in adult tissues but is re-expressed in tumors and the surrounding microenvironment [46], and TNC re-expression has been postulated to contribute to the acquisition of a migratory or stem cell-like phenotype by tumor cells [47]. As mice with TNC knockout develop normally [48], it is possible to consider using TNC neutralizing antibody for metastasis prevention. Another possibility is to use BMP receptor kinase inhibitor to inhibit EC-to-OSB transition. Several BMP receptor kinase inhibitors have been developed for skeletal related disease [49]. Our findings raise the interesting possibility of applying BMP receptor inhibitors for PCa bone metastasis treatments. Together, our studies on the molecular mechanisms leading to the metastatic progression of PCa in bone open up the possibility for future development of strategies to prevent and treat bone metastases more effectively.

## Materials and Methods

### Cell lines, antibodies and reagents

Cell lines, antibodies and reagents are as listed (Supplementary Table 2). C4-2B4-LT (Luciferase-Tomato), C4-2b-LT and PC3-mm2-LT cells were generated previously [50]. Cell lines were authenticated using short tandem repeat (STR) profiling and regularly tested for mycoplasma.

### Differentiation of 2H11 cells to EC-OSB hybrid cells

2H11 cells were incubated overnight in serum-free media and treated with BMP4 (100 ng/mL) for 48-72 h to generate EC-OSB hybrid cells. For differentiation and mineralization, EC-OSB cells were further cultured in StemPro osteoblast differentiation medium (Gibco™) for 21 days. Alkaline phosphatase activity was measured in cell lysates using alkaline phosphatase kit.

### Real-time RT-PCR

Total RNA was isolated using Qiagen RNAeasy kit. 20 ng cDNA was used for qRT-PCR with primers as listed (Supplementary Table 3). Human integrin primers (HINT-1) were purchased from RealTimePrimers. C4-2b-TNC cells were starved overnight, treated with 150 µg/mL integrin α5β1 antibody or control IgG for 48 h before RNA isolation.

### Migration assay and invasion assay

PCa cells (3 × 10^5^) in serum-free medium were seeded into FluoroBlock TM Cell Culture insert (BD Falcon). The lower chamber contained conditioned media (CM) with 10% FBS. Migrated cells were labeled with calcein AM [50], and five or more fields were counted for quantification. The same procedure was used for Matrigel-coated invasion chamber (BD Falcon). For TNC neutralization, EC-OSB-CM was incubated with TNC neutralizing antibody BC-24 at 4°C overnight. Anti-cytokeratin antibody AE1/AE3 was used as control. For anti-integrin α5β1 study, C4-2b-TNC cells were treated with M200 (Volociximab) or IgG for 20 min before assay.

### Soft agar colony assay

C4-2B4, C4-2b or PC3-mm2 cells (3 × 10^4^) were plated on soft agar as described [51], and incubated in 5% FBS medium containing test proteins or CM for 10 to 14 days. Five or more fields were counted for quantification.

### Isobaric Tag for Relative and Absolute Quantitation (iTRAQ)

iTRAQ analysis of CM (10X concentrated) from 2H11 or EC-OSB hybrid cells was performed by Cold Spring Harbor Laboratory’s Mass Spectrometry Shared Resource. Samples were digested with trypsin, labeled with 4-plex iTRAQ reagent, and analyzed on an Orbitrap Fusion Lumos mass spectrometer, equipped with a nano-ion spray source coupled to an EASY-nLC 1200 system (Thermo Scientific) as previously described [52].

### Immunohistochemistry and immunocytochemistry

Formalin-fixed, paraffin-embedded tissues or methanol-fixed cells were analyzed for TNC as described [53, 54]. Human PCa bone metastasis specimens were obtained from Institutional Tissue Bank at M.D. Anderson Cancer Center through IRB approved protocol.

### Overexpression of TNC in C4-2b and PC3 cells or overexpression of Hey1 in 2H11 cells

Human TNC cDNA encoding 1-625 aa was inserted into retroviral vector pBMN-I-GFP to generate pBMN-hTNC-GFP. C4-2b-TNC and PC3-TNC cells were generated by transduction with bicistronic retroviral particles and selected by FACS sorting for GFP. pBMN-I-GFP was used as control. 2H11-oeHey1 cells were generated as previously described [6].

### Knockdown of transcription factors in prostate cancer or 2H11 cells

GIPZ lentiviral shRNA (Horizon) was used to knockdown TNC in PC3-mm2 and C4-2b cells. MISSION pLKO.1 lentiviral shRNA (MilliPORE Sigma) was used to knockdown Smad1, Hey1 in 2H11 cells or YAP in C4-2b cells. Cells transduced with empty pGIPZ or pLKO.1 lentiviral vector were used as controls.

### Intra-prostate injection of prostate cancer cells

PCa cells (1 × 10^6^) were injected into the mouse prostate [55]. Tumor growth was monitored by bioluminescence imaging. DNA was isolated from individual femurs using DNeasy kit. Human-specific *Alu* qPCR was performed to determine the number of tumor cells in each femur as previously described [11].

### Promoter-luciferase reporter assay

LightSwitch TNC promoter GoClone expression plasmid was co-transfected with Hey1 expression plasmid or control plasmid into 293T cells using Lipofectamine 2000. TNC promoter-luciferase activity was measured by the LightSwitch Luciferase Assay Kit. 293T cells were transfected with 8xTEAD synthetic YAP/TAZ-responsive promoter-luciferase reporter and treated with BSA or recombinant TNC (5 µg/mL) for 48 h to measure YAP1 signaling activity using Dual-luciferase Reporter Assay Kit (Promega).

### Statistical Analysis

Data were expressed as the mean ± S.D. p < 0.05 by Student’s *t*-test was considered statistically significant.

## Acknowledgements

This work was supported by grants from the NIH R01CA174798 (S.-H. Lin, L.-Y. Yu-Lee), NIH 5P50CA140388 (C. Logothetis, S.-H. Lin), NIH P30CA16672 Core grant to M.D. Anderson Cancer Center; and Cancer Prevention Research Institute of Texas grants RP150179, RP190252 (S.-H. Lin and L.-Y. Yu-Lee).

