## Supplementary Figures and Tables for "Prostate tumor-induced stromal reprogramming generates Tenascin C that promotes prostate cancer metastasis through YAP/TAZ inhibition"

**Supplementary Figures 1 - 9**

### Supplementary Figure 1

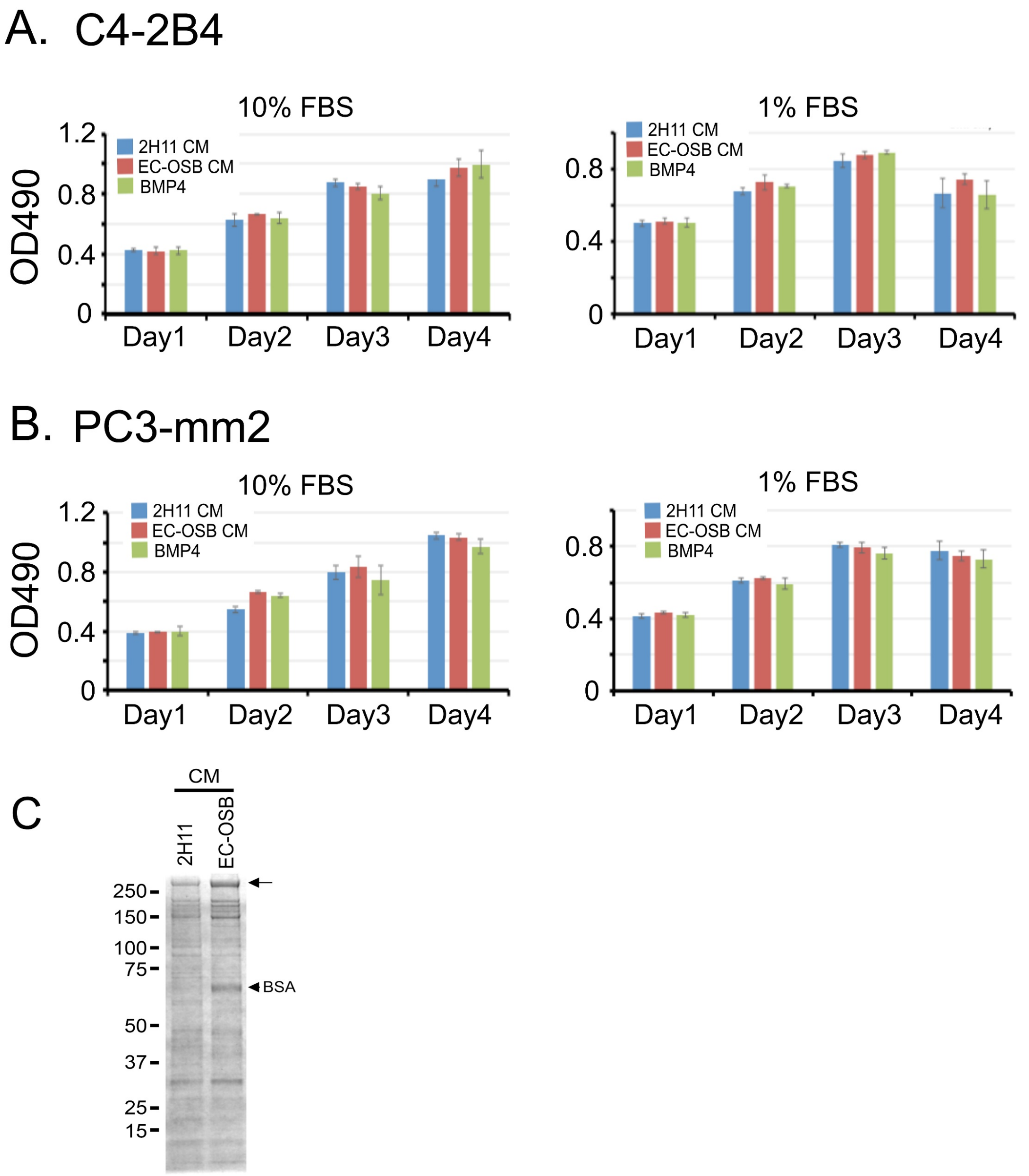

**Supplementary Figure 1**. **Conditioned medium (CM) from 2H11 cells or EC-OSB hybrid cells do not affect C4-2b4 or PC3-mm2 proliferation.** (A, B) C4-2B4 or PC3-mm2 cells were incubated with conditioned media (CM) from 2H11 cells or EC-OSB hybrid cells, or treated directly with BMP4 (100 ng/mL) as a control. Cell proliferation in either 10% or 1% FBS was measured by MTS assay. (C) SDS-PAGE protein profile of 2H11-CM or EC-OSB-CM from serum-free media. Arrow, potential 250 kDa Tenascin C.

Supplementary Figure 2

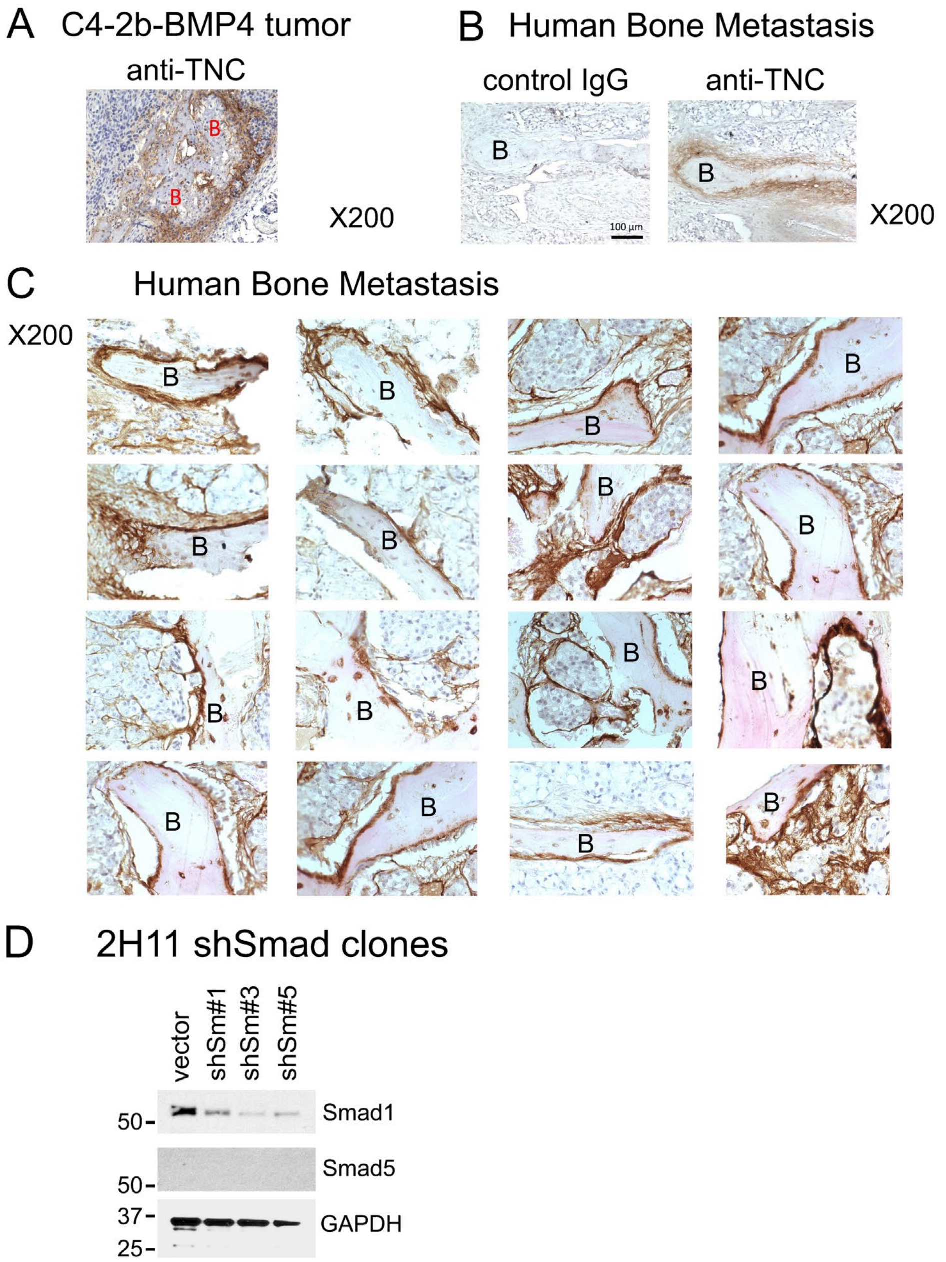

**Supplementary Figure 2**. **Immunohistochemical analyses of Tenascin C in human bone metastasis specimens and western blot of Smad1 and Smad5 levels in 2H11 shSmad clones.** IHC of TNC in osteogenic C4-2b-BMP4 tumor (A) and human bone metastasis specimens (B, C). Anti-TNC antibody, but not control IgG, showed TNC expression in EC-OSB hybrid cells rimming tumor-induced bone in human PCa bone metastasis. TNC staining (brown) is localized in newly formed bone, where EC-OSB hybrid cells are located. B, bone. (D) Western blot of Smad1 and Smad5 levels in 2H11 shSmad1 clones. Smad5 was undetectable in 2H11 cells with or without Smad1 knockdown.

### Supplementary Figure 3

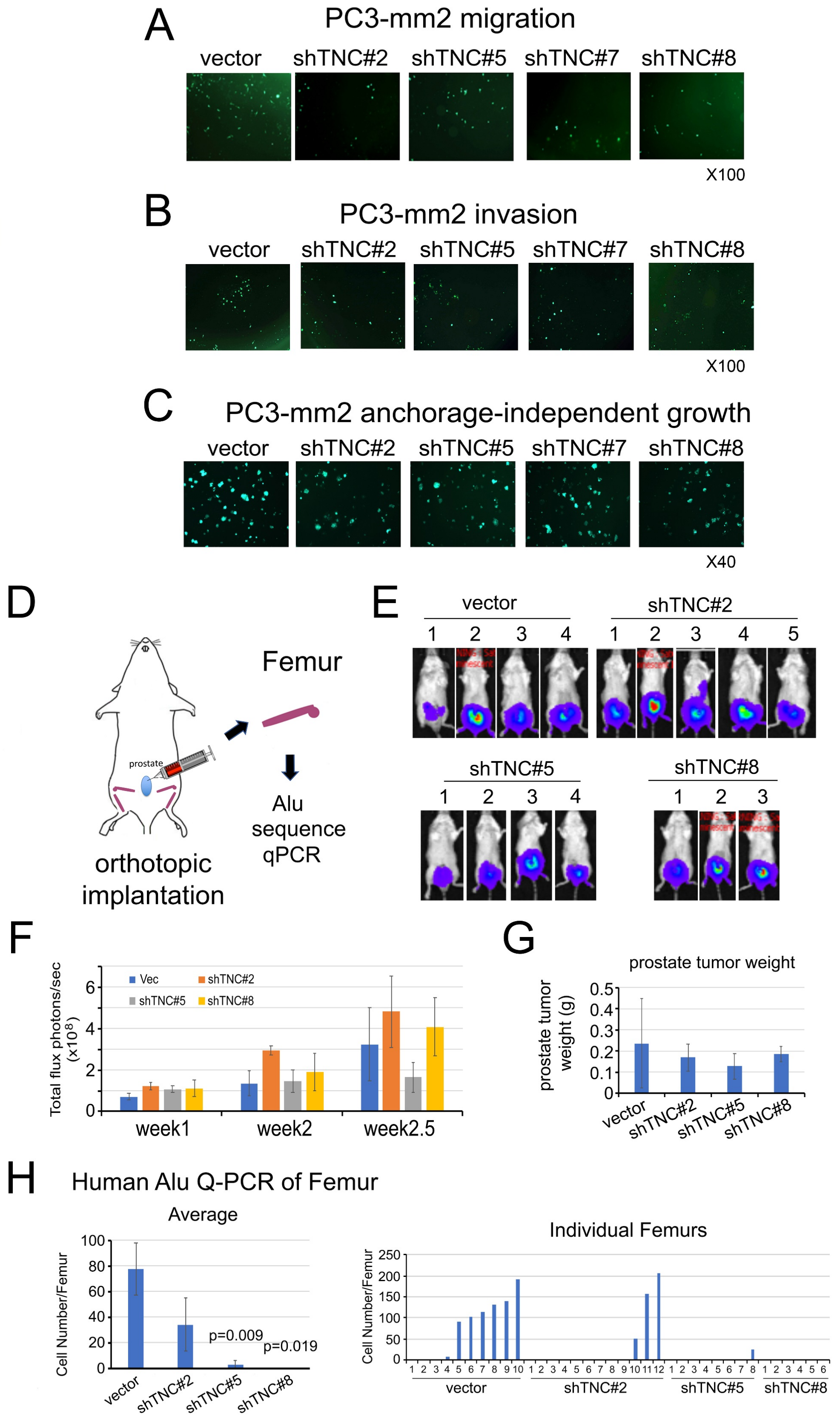

**Supplementary Figure 3**. **Knockdown of Tenascin C in PC3-mm2 cells reduces metastasis of PC3-mm2 cells to bone in vivo.** PC3-mm2-vector and four PC3-mm2-shTNCmm2 clones were examined for (A) migration, (B) invasion, and (C) anchorage-independent growth. (D) Cells in A were injected into mouse prostate. (E) Bioluminescence of tumors at 3 weeks post-injection. (F) Tumor growth rates. (G) Average tumor weights. (H) Quantification of the number of tumor cells that have metastasized from the prostate to bone using human-specific *Alu*-qPCR of DNA from femur. Left, average number of tumor cells metastasized from prostate to bone. P values were by Student's t-test. Right, tumor cell numbers in individual legs.

### Supplementary Figure 4

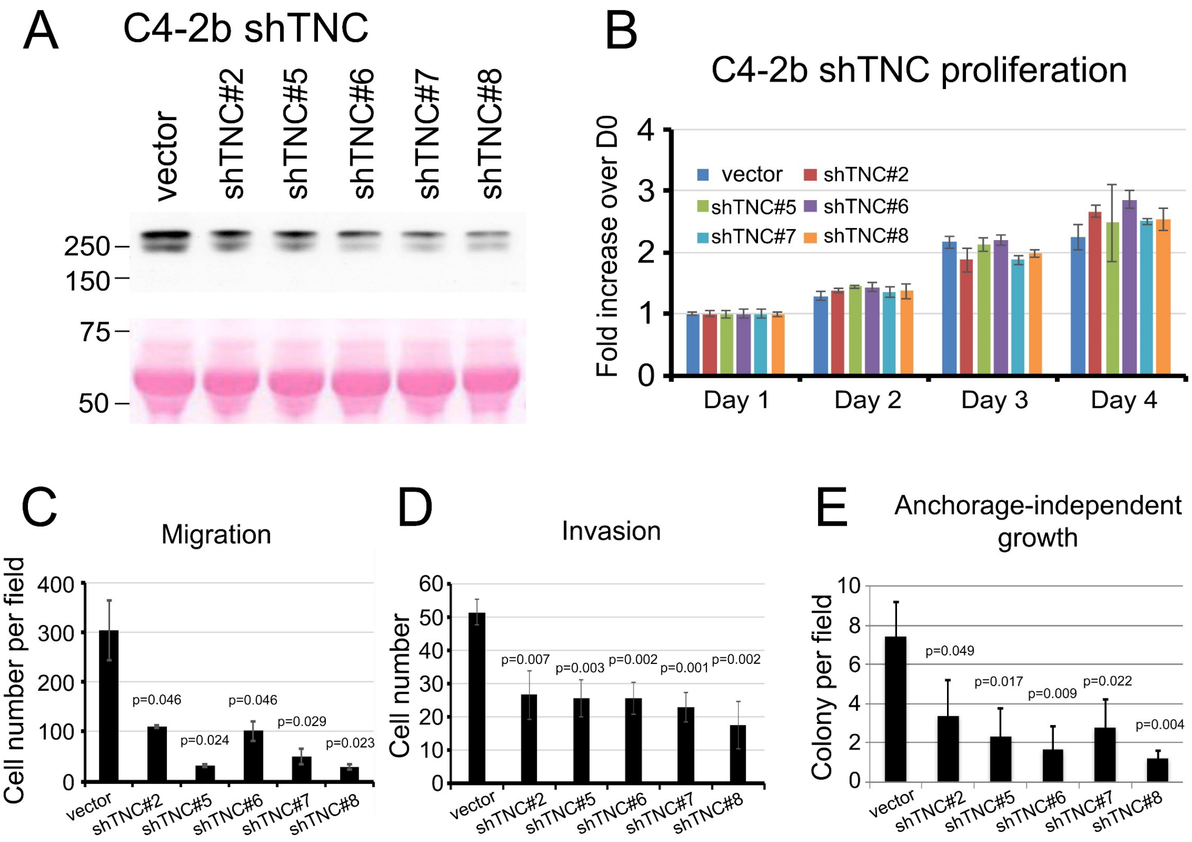

**Supplementary Figure 4**. **Knockdown of Tenascin C in C4-2b cells reduces migration, invasion, and anchorage-independent growth of C4-2b cells in vitro.** C4-2b-vector and five C4-2b-shTNC clones were examined for (A) TNC in conditioned media by western blot, (B) proliferation by MTS assay, (C) migration, (D) invasion, and (E) anchorage-independent growth.

#### Supplementary Figure 5

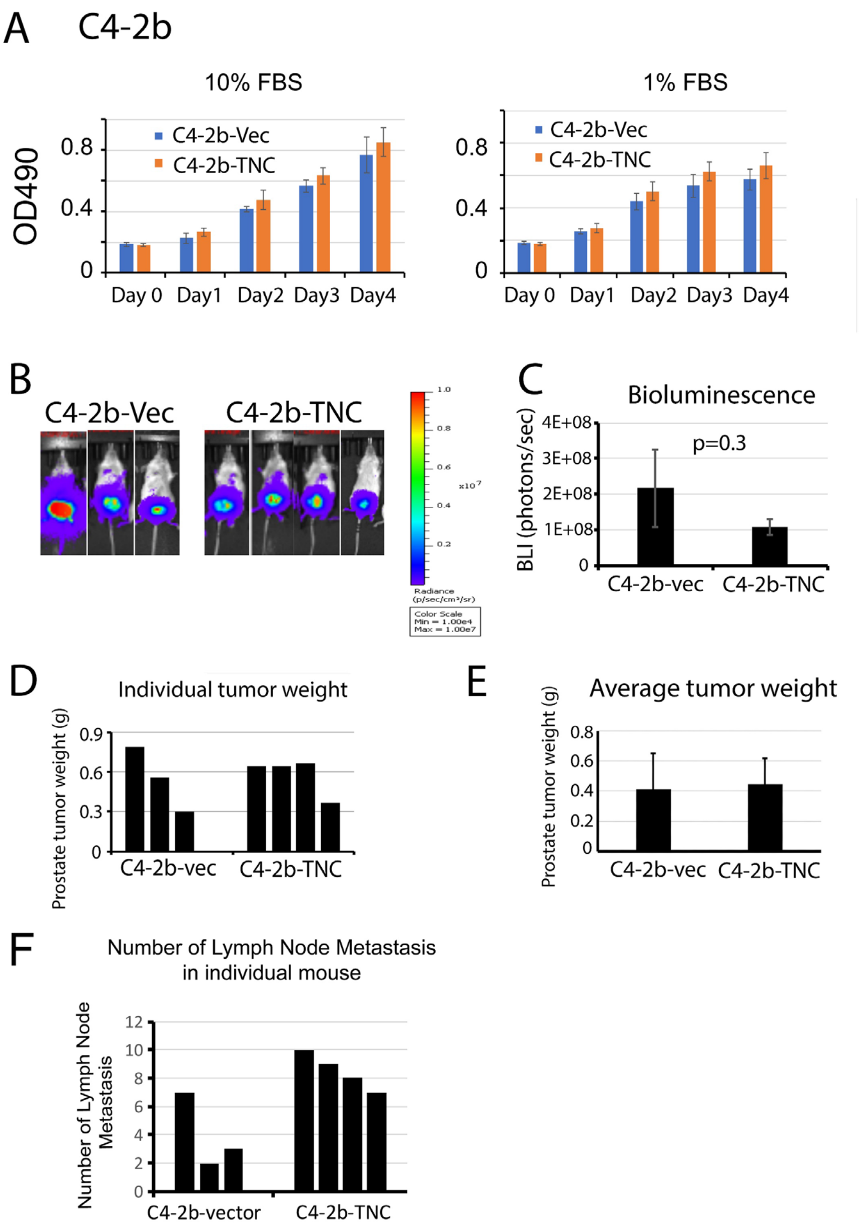

**Supplementary Figure 5**. **Overexpression of Tenascin C in C4-2b cells does not affect C4-2b cell proliferation and tumor growth in mouse prostate, but increases C4-2b-TNC metastasis to lymph nodes.** (A) Proliferation of C4-2b-vector and C4-2b-TNC cells by MTS assay. (B) C4-2b-vector or C4-2b-TNC cells were injected into mouse prostate. Tumor growth was monitored by bioluminescence. (C) No significant difference in tumor sizes by bioluminescence. (D) Individual tumor weight. (E) No significant difference in average tumor weight. (F) Number of lymph node metastasis in individual mouse.

Supplementary Figure 6

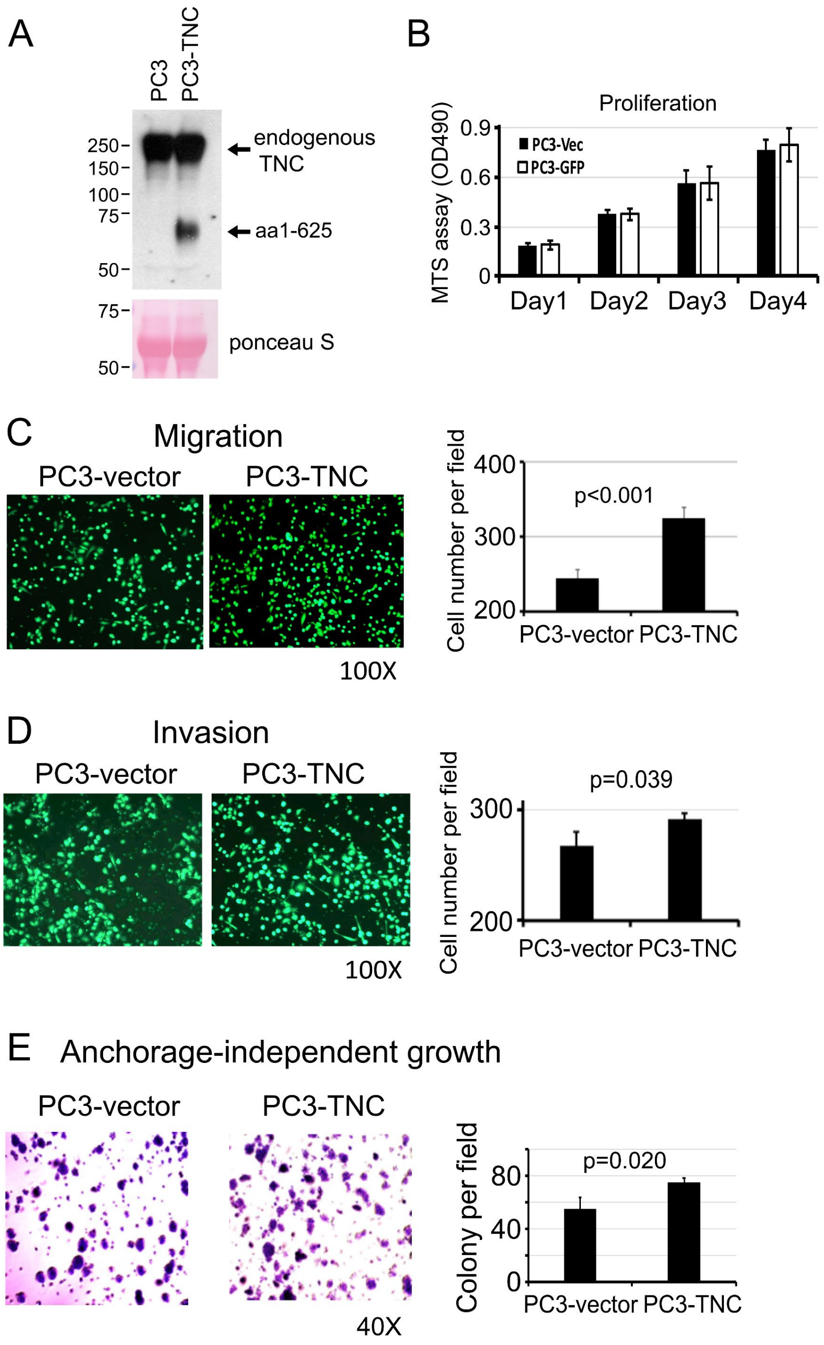

**Supplementary Figure 6**. **Overexpression of N-terminal domain (1 – 625 aa) of TNC in PC3 cells.** (A) Western blot for endogenously-expressed TNC (~250 kDa) and exogenously-expressed TNC (~70 kDa) in PC3-TNC cells. (B) Cell proliferation for PC3-vector and PC3-TNC cells by MTS assay. (C-E) Migration, invasion and anchorage-independent growth of cells in (B). P values are from Student's *t*-test. Overexpression of TNC in PC3 cells increased migration, invasion and anchorage-independent growth in PC3-TNC cells.

### Supplementary Figure 7

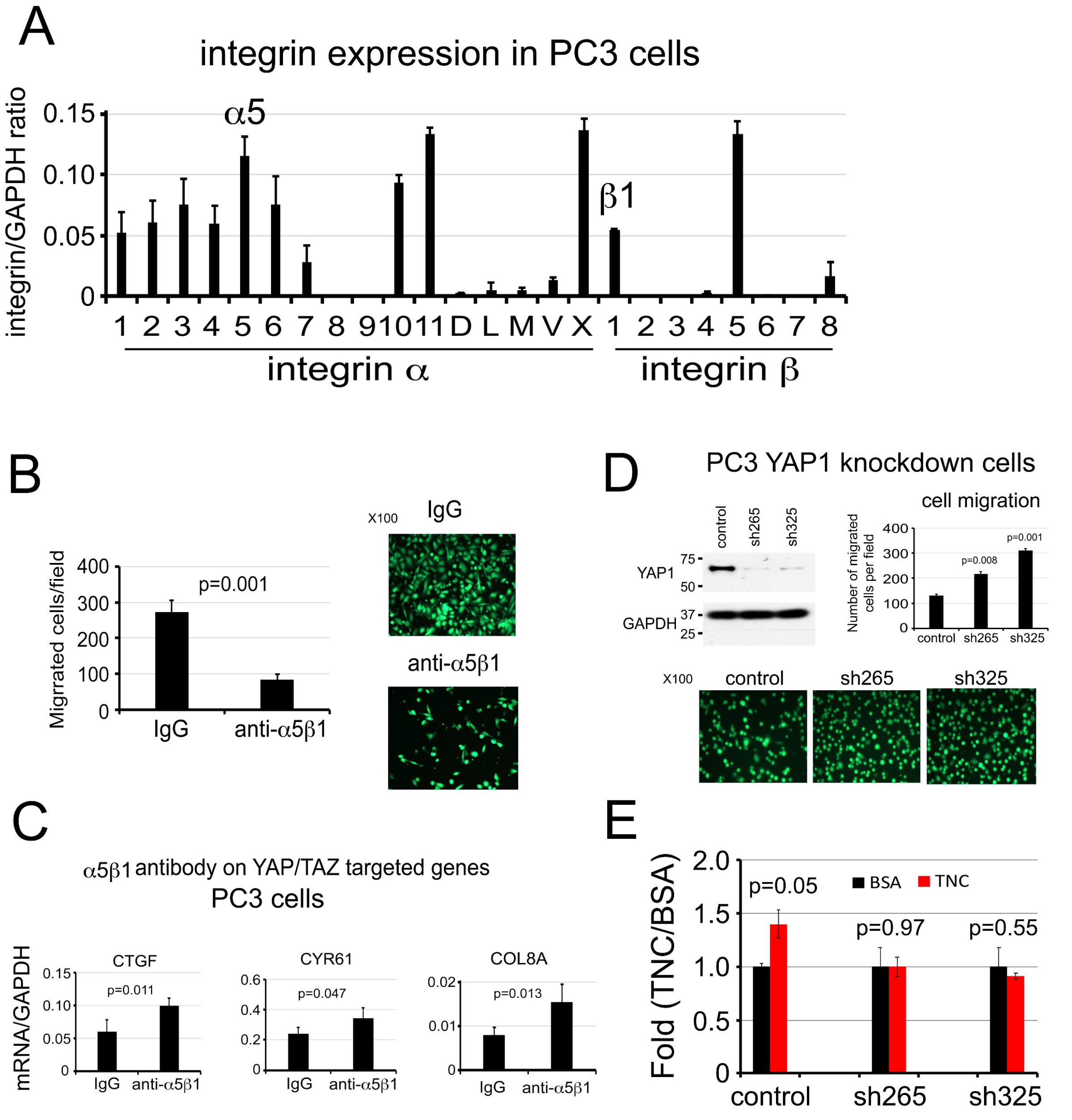

**Supplementary Figure 7. Tenascin C promotes PC3 cell migration through** α5β1**-YAP1 pathway.** (A) Integrin expression in PC3 cells by qRT-PCR. (B) Cell migration in PC3 cells incubated with anti-α5β1 integrin neutralizing antibody or control IgG. (C) YAP/TAZ target gene expression in cells following incubation with α5β1 integrin antibody or IgG control. (D) Cell migration in YAP1 knockdown PC3-shYAP1clones. (E) Cell migration in PC3-shYAP1 clones in response to 5 μg/mL recombinant TNC or control BSA.

Supplementary Figure 8

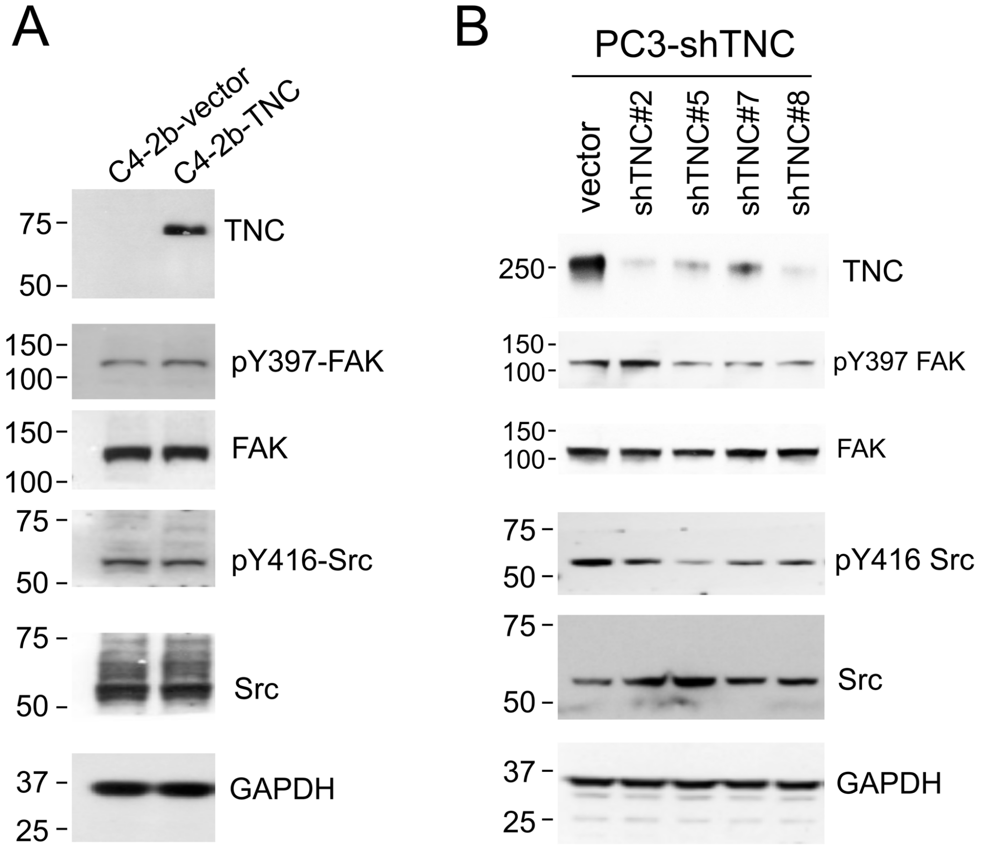

**Supplementary Figure 8. TNC expression does not correlate with FAK and Src phosphorylation in prostate cancer cell lines.** (A) FAK and Src phosphorylation in C4-2b-vector and C4-2b-TNC cells by western blot. (B) FAK and Src phosphorylation in PC3-shTNC clones. No consistent correlation between TNC expression and pY397-FAK and pY416-Src phosphorylation was observed in different PCa cells.

Supplementary Figure 9

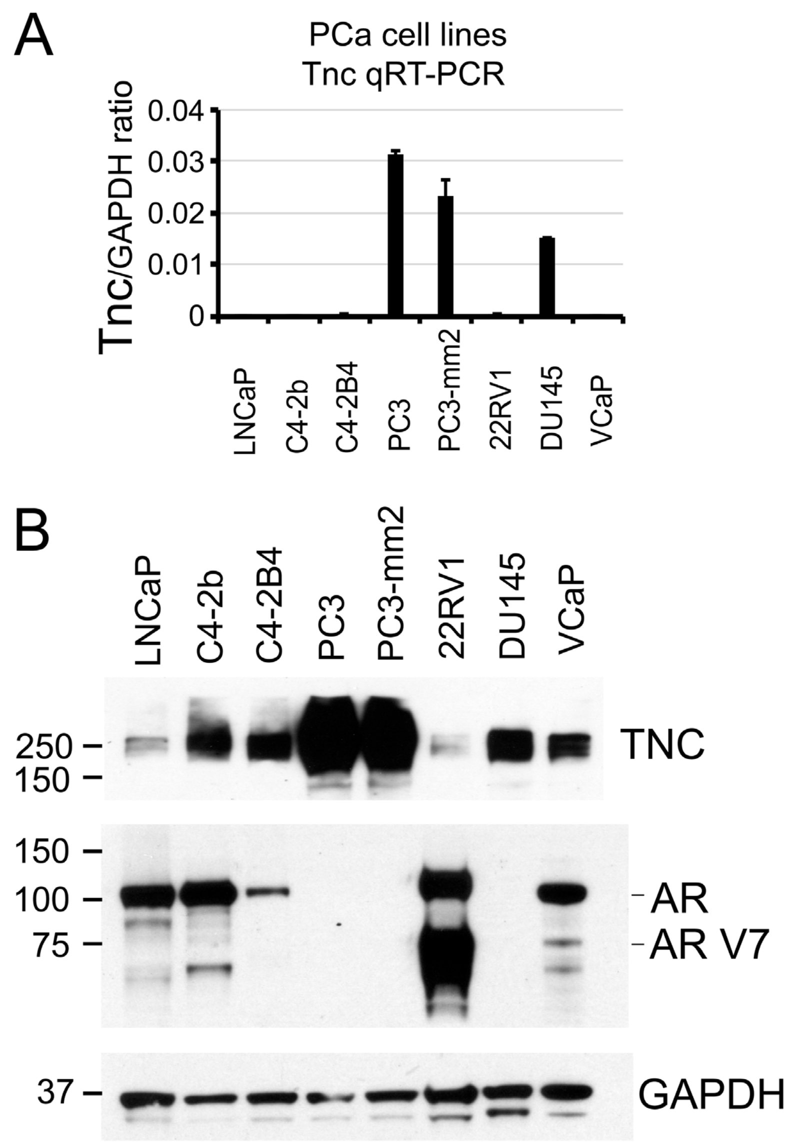

**Supplementary Figure 9. Tenascin C expression does not correlate with androgen receptor and androgen receptor variant expression in prostate cancer cell lines.** (A) qRT-PCR for TNC mRNA in PCa cell lines. (B) Western blot of protein expression of TNC, androgen receptor (AR) and androgen receptor variant V7 (AR V7) in PCa cell lines.

User admin
E-mail admin@localhost
Search titl YuChen iTRAQ2
MS data fi E:\Shared Storage\MGF\3255 YuChen\mascot daemon merge.mgf
Database UniProt mouse 20170829
Timestam 17 Sep 2017 at 02:52:19 GMT
Type of se MS/MS Ion Search
Quantitat YuChen (Applied Biosystems iTRAQ(TM) 8-plex)
Enzyme Trypsin
Fixed moc iTRAQ8plex (N-term),iTRAQ8plex (K),Methylthio (C)
Variable n Deamidated (NQ),Oxidation (M)
Mass valu Monoisotopic
Protein m Unrestricted
Peptide m 30 ppm
Fragment 0.2 Da
Max misse 2
Instrumer ESI-QUAD
Number c 163,401

Significan 0.01734
Max. num AUTO
Ions score -1
Preferred All entries
Show Perc no

Quantitation parameters:
Protein ra weighted
Normalisa "Summed intensities"
Min. prec 1
Outlier re none
Min. num 1
Peptide th at least homology
Unique pe no

Significant changes (<0.81, >1.22)

| Family | Member | Database | Accession | Score | Mass | Num. of n | Num. of s | Num. of s | Num. of si | emPAI | C1 | C2 | C3 | BMP4-1 | BMP4-2 | BMP4-3 | C Average | BMP4 Average | Description |
|---|---|---|---|---|---|---|---|---|---|---|---|---|---|---|---|---|---|---|---|
| 88 | 1 | UniProt n | P07724 | 1122 | 86122 | 24 | 24 | 2 | 2 | 0.16 | 1 | 0.915 | 0.798 | 8.049 | 7.767 | 8.814 | 0.900 | 8.198 | Serum albumin OS=Mus musculus GN=Alb PE=1 SV=3 |
| 1149 | 1 | UniProt n | Q8BKT8 | 32 | 49137 | 2 | 2 | 1 | 1 | 0.09 | 1 | 1.04 | 1.105 | 4.265 | 4.627 | 3.965 | 1.047 | 4.277 | HAUS augmin-like complex subunit 7 OS=Mus musculus GN=Haus7 PE=1 SV=2 |
| 1071 | 1 | UniProt n | F6ZIS7 | 39 | 36563 | 2 | 2 | 1 | 1 | 0.12 | 1 | 0.705 | 0.878 | 3.322 | 4.695 | 4.354 | 0.852 | 4.080 | Retinoic acid receptor responder (tazarotene-induced) 1 OS=Mus musculus GN=Rarres1 PE=4 SV=1 |
| 13 | 2 | UniProt n | Q80YX1-2 | 5038 | 247888 | 172 | 172 | 59 | 59 | 2.32 | 1 | 0.917 | 0.988 | 3.476 | 3.192 | 3.426 | 0.968 | 3.362 | Isoform 2 of Tenascin OS=Mus musculus GN=Tnc |
| 13 | 1 | UniProt n | Q80YX1 | 5096 | 258723 | 174 | 174 | 60 | 60 | 2.21 | 1 | 0.908 | 0.98 | 3.391 | 3.121 | 3.35 | 0.962 | 3.285 | Tenascin OS=Mus musculus GN=Tnc PE=1 SV=1 |
| 28 | 1 | UniProt n | P35441 | 3360 | 150427 | 134 | 134 | 42 | 42 | 3.46 | 1 | 0.916 | 0.97 | 3.033 | 2.931 | 3.128 | 0.961 | 3.030 | Thrombospondin-1 OS=Mus musculus GN=Thbs1 PE=1 SV=1 |
| 1175 | 1 | UniProt n | A0A1D5RI | 30 | 182787 | 1 | 1 | 1 | 1 | 0.02 | 1 | 0.966 | 1.076 | 2.456 | 3.371 | 3.357 | 1.013 | 3.029 | Golgin subfamily A member 4 (Fragment) OS=Mus musculus GN=Golga4 PE=1 SV=1 |
| 1036 | 1 | UniProt n | E9PVD7 | 42 | 191077 | 7 | 7 | 1 | 1 | 0.02 | 1 | 0.767 | 0.877 | 3.345 | 3.015 | 2.594 | 0.876 | 2.969 | Cystic fibrosis transmembrane conductance regulator OS=Mus musculus GN=Cftr PE=1 SV=1 |
| 181 | 1 | UniProt n | E9QNQ3 | 516 | 211992 | 25 | 25 | 16 | 16 | 0.38 | 1 | 0.907 | 0.937 | 2.892 | 2.744 | 2.851 | 0.947 | 2.828 | Latent-transforming growth factor beta-binding protein 2 OS=Mus musculus GN=Ltbp2 PE=1 SV=1 |
| 50 | 1 | UniProt n | P50608 | 1944 | 46391 | 58 | 58 | 9 | 9 | 1.98 | 1 | 0.915 | 0.87 | 2.786 | 2.749 | 2.929 | 0.927 | 2.820 | Fibromodulin OS=Mus musculus GN=Fmod PE=2 SV=1 |
| 1134 | 1 | UniProt n | Q8C0D5 | 33 | 150575 | 1 | 1 | 1 | 1 | 0.03 | 1 | 1.159 | 1.417 | 2.304 | 2.252 | 2.835 | 1.180 | 2.450 | Elongation factor-like GTPase 1 OS=Mus musculus GN=Efl1 PE=1 SV=1 |
| 636 | 1 | UniProt n | E9Q1F6 | 94 | 171037 | 3 | 3 | 2 | 2 | 0.05 | 1 | 0.9 | 0.988 | 2.434 | 2.231 | 2.492 | 0.962 | 2.383 | Tenascin-N OS=Mus musculus GN=Tnn PE=1 SV=1 |
| 360 | 1 | UniProt n | G3XA35 | 234 | 296756 | 12 | 12 | 7 | 7 | 0.11 | 1 | 0.999 | 0.952 | 2.334 | 2.104 | 2.489 | 0.983 | 2.304 | MCG116562, isoform CRA a OS=Mus musculus GN=Vcan PE=1 SV=1 |
| 1249 | 1 | UniProt n | E0CYH5 | 23 | 17558 | 1 | 1 | 1 | 1 | 0.27 | 1 | 1.411 | 1.907 | 2.339 | 2.226 | 2.333 | 1.391 | 2.299 | Collagen, type XXII, alpha 1 (Fragment) OS=Mus musculus GN=Col22a1 PE=1 SV=8 |
| 16 | 1 | UniProt n | P97298 | 4730 | 53552 | 138 | 138 | 20 | 20 | 6.2 | 1 | 1.045 | 1.027 | 2.312 | 2.21 | 2.376 | 1.024 | 2.298 | Pigment epithelium-derived factor OS=Mus musculus GN=Serpinf1 PE=1 SV=2 |
| 1163 | 1 | UniProt n | Q8BZW8 | 31 | 91543 | 1 | 1 | 1 | 1 | 0.05 | 1 | 2.41 | 2.376 | 3.277 | 1.636 | 2.106 | 1.789 | 2.243 | NHL repeat-containing protein 2 OS=Mus musculus GN=Nhlrc2 PE=1 SV=1 |
| 1166 | 1 | UniProt n | A0A1W2P | 30 | 15397 | 1 | 1 | 1 | 1 | 0.31 | 1 | 0.877 | 0.773 | 2.197 | 2.338 | 1.977 | 0.878 | 2.166 | RNA-binding protein Nova-1 (Fragment) OS=Mus musculus GN=Nova1 PE=1 SV=1 |
| 269 | 1 | UniProt n | P48759 | 328 | 45895 | 14 | 14 | 8 | 8 | 1.29 | 1 | 0.917 | 0.987 | 2.166 | 2.114 | 2.14 | 0.967 | 2.140 | Pentraxin-related protein PTX3 OS=Mus musculus GN=Ptx3 PE=1 SV=2 |
| 1066 | 1 | UniProt n | H3BJW0 | 39 | 37170 | 2 | 2 | 2 | 2 | 0.25 | 1 | 0.893 | 0.91 | 2.128 | 2.078 | 2.161 | 0.933 | 2.122 | Connective tissue growth factor OS=Mus musculus GN=Ctgf PE=1 SV=1 |
| 265 | 1 | UniProt n | P70202 | 329 | 30177 | 6 | 6 | 2 | 2 | 0.32 | 1 | 1.039 | 1.093 | 2.106 | 2.037 | 2.202 | 1.043 | 2.114 | Latexin OS=Mus musculus GN=Lxn PE=1 SV=2 |
| 1268 | 1 | UniProt n | G3X914 | 20 | 122558 | 1 | 1 | 1 | 1 | 0.04 | 1 | 3.646 | 3.933 | --- | 1.865 | 2.332 | 2.429 | 2.085 | Cullin-5 OS=Mus musculus GN=Cul5 PE=1 SV=1 |
| 5 | 1 | UniProt n | Q60847 | 10658 | 387818 | 387 | 387 | 105 | 105 | 3.07 | 1 | 0.979 | 1.077 | 2.063 | 1.888 | 2.155 | 1.018 | 2.032 | Collagen alpha-1(XII) chain OS=Mus musculus GN=Col12a1 PE=2 SV=3 |
| 443 | 1 | UniProt n | P22777 | 176 | 51575 | 10 | 10 | 5 | 5 | 0.51 | 1 | 0.804 | 0.89 | 1.906 | 1.968 | 1.967 | 0.894 | 1.947 | Plasminogen activator inhibitor 1 OS=Mus musculus GN=Serpine1 PE=1 SV=1 |
| 227 | 1 | UniProt n | Q6GU68 | 382 | 49226 | 17 | 17 | 7 | 7 | 0.99 | 1 | 0.895 | 0.903 | 1.916 | 1.882 | 1.919 | 0.931 | 1.906 | Immunoglobulin superfamily containing leucine-rich repeat protein OS=Mus musculus GN=Islr PE=1 SV=1 |
| 123 | 1 | UniProt n | F8WH23 | 819 | 35081 | 29 | 29 | 13 | 13 | 4.36 | 1 | 1.031 | 1 | 1.839 | 1.933 | 1.89 | 1.010 | 1.887 | Insulin-like growth factor-binding protein 7 OS=Mus musculus GN=Igfbp7 PE=1 SV=1 |
| 1235 | 1 | UniProt n | E9PUJ2 | 25 | 81977 | 1 | 1 | 1 | 1 | 0.05 | 1 | 2.562 | 2.56 | --- | 1.481 | 2.379 | 1.872 | 1.877 | DNA repair protein RAD50 OS=Mus musculus GN=Rad50 PE=1 SV=1 |
| 20 | 1 | UniProt n | Q62356 | 4499 | 44273 | 178 | 178 | 17 | 17 | 5.72 | 1 | 1.079 | 1.001 | 1.88 | 1.819 | 1.868 | 1.026 | 1.855 | Follistatin-related protein 1 OS=Mus musculus GN=Fstl1 PE=1 SV=2 |
| 1 | 2 | UniProt n | A0A087W | 29487 | 279220 | 786 | 786 | 95 | 95 | 8.24 | 1 | 1.028 | 1.01 | 1.901 | 1.716 | 1.925 | 1.013 | 1.845 | Fibronectin OS=Mus musculus GN=Fn1 PE=1 SV=1 |
| 1 | 1 | UniProt n | A0A087W | 30334 | 289653 | 806 | 806 | 98 | 98 | 7.78 | 1 | 1.026 | 1.009 | 1.896 | 1.712 | 1.923 | 1.012 | 1.841 | Fibronectin OS=Mus musculus GN=Fn1 PE=1 SV=1 |
| 1 | 3 | UniProt n | P11276 | 29079 | 299343 | 787 | 787 | 97 | 97 | 7.07 | 1 | 1.025 | 1.006 | 1.893 | 1.709 | 1.924 | 1.010 | 1.839 | Fibronectin OS=Mus musculus GN=Fn1 PE=1 SV=4 |
| 62 | 1 | UniProt n | P28653 | 1524 | 49586 | 50 | 50 | 14 | 14 | 3.63 | 1 | 1.049 | 1.039 | 1.797 | 1.739 | 1.919 | 1.029 | 1.817 | Biglycan OS=Mus musculus GN=Bgn PE=1 SV=1 |
| 1182 | 1 | UniProt n | P50228 | 29 | 18365 | 1 | 1 | 1 | 1 | 0.26 | 1 | 1.076 | 1.1 | 1.885 | 1.955 | 1.543 | 1.058 | 1.785 | C-X-C motif chemokine 5 OS=Mus musculus GN=Cxcl5 PE=1 SV=2 |
| 1 | 4 | UniProt n | A0A087W | 202 | 21585 | 19 | 19 | 6 | 6 | 4.73 | 1 | 1.041 | 0.967 | 1.835 | 1.609 | 1.907 | 1.002 | 1.779 | Fibronectin (Fragment) OS=Mus musculus GN=Fn1 PE=1 SV=1 |
| 179 | 1 | UniProt n | A0A0R4J0 | 519 | 50469 | 16 | 16 | 7 | 7 | 0.95 | 1 | 0.976 | 0.955 | 1.813 | 1.756 | 1.747 | 0.977 | 1.772 | Platelet-derived growth factor receptor-like protein OS=Mus musculus GN=Pdgfrl PE=4 SV=1 |
| 125 | 1 | UniProt n | P97299 | 818 | 42488 | 26 | 26 | 7 | 7 | 3.01 | 1 | 1.082 | 1.172 | 1.785 | 1.775 | 1.679 | 1.082 | 1.746 | Secreted frizzled-related protein 2 OS=Mus musculus GN=Sfrp2 PE=2 SV=2 |
| 319 | 1 | UniProt n | P12032 | 271 | 26511 | 8 | 8 | 4 | 4 | 1.2 | 1 | 0.986 | 0.909 | 1.726 | 1.727 | 1.78 | 0.964 | 1.744 | Metalloproteinase inhibitor 1 OS=Mus musculus GN=Timp1 PE=1 SV=2 |
| 28 | 2 | UniProt n | Q03350 | 2531 | 147832 | 82 | 82 | 29 | 29 | 1.65 | 1 | 0.934 | 0.981 | 1.774 | 1.644 | 1.783 | 0.971 | 1.732 | Thrombospondin-2 OS=Mus musculus GN=Thbs2 PE=1 SV=2 |
| 1270 | 1 | UniProt n | D3YU89 | 19 | 286327 | 1 | 1 | 1 | 1 | 0.01 | 1 | 1.161 | 1.028 | 1.549 | 1.792 | 1.799 | 1.061 | 1.709 | Citron Rho-interacting kinase OS=Mus musculus GN=Cit PE=1 SV=1 |
| 1076 | 1 | UniProt n | Q9DAU1 | 38 | 39967 | 1 | 1 | 1 | 1 | 0.11 | 1 | 0.873 | 0.946 | 0.955 | 5.993 | 0.81 | 0.938 | 1.667 | Protein canopy homolog 3 OS=Mus musculus GN=Cnpy3 PE=1 SV=1 |
| 849 | 1 | UniProt n | Q61592 | 60 | 86468 | 2 | 2 | 2 | 2 | 0.1 | 1 | 0.887 | 1.034 | 1.773 | 1.609 | 1.584 | 0.972 | 1.653 | Growth arrest-specific protein 6 OS=Mus musculus GN=Gas6 PE=2 SV=2 |
| 36 | 1 | UniProt n | A0A1L1SS | 2709 | 44724 | 135 | 135 | 11 | 11 | 2.74 | 1 | 1.074 | 0.95 | 1.611 | 1.709 | 1.538 | 1.007 | 1.618 | SPARC OS=Mus musculus GN=Sparc PE=1 SV=1 |
| 1226 | 1 | UniProt n | Q5SQX6 | 26 | 170202 | 1 | 1 | 1 | 1 | 0.03 | 1 | 1.381 | 1.082 | 1.259 | 1.653 | 2.028 | 1.143 | 1.616 | Cytoplasmic FMR1-interacting protein 2 OS=Mus musculus GN=Cyfip2 PE=1 SV=2 |
| 1097 | 1 | UniProt n | Q3UVI3 | 36 | 72828 | 1 | 1 | 1 | 1 | 0.06 | 1 | 0.749 | 0.724 | 1.667 | 1.444 | 1.731 | 0.815 | 1.609 | Transformation-related protein 63 OS=Mus musculus GN=Trp63 PE=2 SV=1 |
| 194 | 1 | UniProt n | P98064-2 | 463 | 93065 | 22 | 22 | 11 | 11 | 0.65 | 1 | 0.942 | 0.964 | 1.601 | 1.564 | 1.615 | 0.968 | 1.593 | Isoform 2 of Mannan-binding lectin serine protease 1 OS=Mus musculus GN=Masp1 |
| 691 | 1 | UniProt n | Q9JK53 | 84 | 49625 | 3 | 3 | 2 | 2 | 0.19 | 1 | 0.889 | 0.98 | 1.645 | 1.487 | 1.52 | 0.955 | 1.549 | Prolargin OS=Mus musculus GN=Prelp PE=1 SV=2 |
| 11 | 1 | UniProt n | Q640N1 | 5622 | 151089 | 179 | 179 | 29 | 29 | 2.44 | 1 | 0.994 | 0.981 | 1.495 | 1.451 | 1.632 | 0.992 | 1.524 | Adipocyte enhancer-binding protein 1 OS=Mus musculus GN=Aebp1 PE=1 SV=1 |
| 1264 | 1 | UniProt n | D3Z763 | 20 | 133419 | 1 | 1 | 1 | 1 | 0.03 | 1 | 0.998 | 0.992 | 1.515 | 1.512 | 1.479 | 0.997 | 1.502 | Brother of CDO OS=Mus musculus GN=Boc PE=1 SV=2 |
| 337 | 1 | UniProt n | Q9JHI0 | 252 | 67786 | 9 | 9 | 5 | 5 | 0.45 | 1 | 0.959 | 0.999 | 1.45 | 1.531 | 1.491 | 0.986 | 1.490 | Matrix metalloproteinase-19 OS=Mus musculus GN=Mmp19 PE=2 SV=1 |
| 1254 | 1 | UniProt n | D3YU40 | 22 | 21925 | 1 | 1 | 1 | 1 | 0.21 | 1 | 1.124 | 1.29 | 1.404 | 1.505 | 1.545 | 1.132 | 1.483 | Insulin-like growth factor-binding protein 2 OS=Mus musculus GN=Igfbp2 PE=1 SV=1 |
| 1118 | 1 | UniProt n | A0A0R4J0 | 34 | 47749 | 2 | 2 | 2 | 2 | 0.19 | 1 | 0.904 | 0.996 | 1.476 | 1.507 | 1.393 | 0.966 | 1.458 | Follistatin OS=Mus musculus GN=Fst PE=1 SV=1 |
| 1267 | 1 | UniProt n | P59222 | 20 | 100877 | 1 | 1 | 1 | 1 | 0.04 | 1 | 1.517 | 0.711 | 1.935 | 1.131 | 1.379 | 1.026 | 1.445 | Scavenger receptor class F member 2 OS=Mus musculus GN=Scarf2 PE=1 SV=1 |
| 489 | 1 | UniProt n | P04202 | 150 | 51222 | 5 | 5 | 2 | 2 | 0.18 | 1 | 1.021 | 1.115 | 1.503 | 1.39 | 1.443 | 1.044 | 1.445 | Transforming growth factor beta-1 OS=Mus musculus GN=Tgfb1 PE=1 SV=1 |
| 1199 | 1 | UniProt n | Q91XD7 | 28 | 52698 | 2 | 2 | 1 | 1 | 0.08 | 1 | 0.998 | 1.206 | 1.467 | 1.417 | 1.397 | 1.064 | 1.427 | Cysteine-rich with EGF-like domain protein 1 OS=Mus musculus GN=Creld1 PE=1 SV=1 |
| 709 | 1 | UniProt n | Q04998 | 80 | 58957 | 1 | 1 | 1 | 1 | 0.07 | 1 | 0.887 | 1.01 | 1.525 | 1.513 | 1.258 | 0.964 | 1.426 | Inhibin beta A chain OS=Mus musculus GN=Inhba PE=1 SV=1 |
| 64 | 1 | UniProt n | Q8BK62 | 1514 | 51819 | 53 | 53 | 14 | 14 | 3 | 1 | 0.947 | 0.957 | 1.423 | 1.399 | 1.447 | 0.968 | 1.423 | Olfactomedin-like protein 3 OS=Mus musculus GN=Olfml3 PE=2 SV=2 |
| 29 | 7 | UniProt n | P19001 | 434 | 50599 | 22 | 22 | 8 | 8 | 0.95 | 1 | 1.117 | 1.284 | 1.171 | 1.739 | 1.403 | 1.128 | 1.419 | Keratin, type I cytoskeletal 19 OS=Mus musculus GN=Krt19 PE=1 SV=1 |
| 894 | 1 | UniProt n | Q9QXP7 | 55 | 37528 | 2 | 2 | 1 | 1 | 0.12 | 1 | 1.022 | 1.225 | 1.297 | 1.501 | 1.464 | 1.078 | 1.418 | Complement C1q tumor necrosis factor-related protein 1 OS=Mus musculus GN=C1qtnf1 PE=2 SV=1 |
| 331 | 1 | UniProt n | Q03366 | 261 | 14872 | 9 | 9 | 3 | 3 | 2.05 | 1 | 1.078 | 1.005 | 1.351 | 1.469 | 1.416 | 1.027 | 1.411 | C-C motif chemokine 7 OS=Mus musculus GN=Ccl7 PE=3 SV=1 |
| 222 | 1 | UniProt n | P28301 | 393 | 49049 | 25 | 25 | 7 | 7 | 1.17 | 1 | 0.731 | 0.703 | 1.267 | 1.385 | 1.581 | 0.801 | 1.405 | Protein-lysine 6-oxidase OS=Mus musculus GN=Lox PE=1 SV=1 |
| 845 | 1 | UniProt n | P04117 | 60 | 19296 | 2 | 2 | 2 | 2 | 0.54 | 1 | 0.888 | 1.004 | 1.554 | 1.492 | 1.193 | 0.962 | 1.404 | Fatty acid-binding protein, adipocyte OS=Mus musculus GN=Fabp4 PE=1 SV=3 |
| 68 | 1 | UniProt n | P55288 | 1461 | 100244 | 39 | 39 | 17 | 17 | 1.54 | 1 | 0.999 | 1.019 | 1.438 | 1.403 | 1.361 | 1.006 | 1.400 | Cadherin-11 OS=Mus musculus GN=Cdh11 PE=1 SV=1 |
| 671 | 1 | UniProt n | P41245 | 88 | 91351 | 3 | 3 | 3 | 3 | 0.15 | 1 | 0.674 | 0.68 | 1.235 | 1.386 | 1.596 | 0.771 | 1.398 | Matrix metalloproteinase-9 OS=Mus musculus GN=Mmp9 PE=1 SV=2 |
| 132 | 2 | UniProt n | Q9QUK9 | 320 | 29854 | 18 | 18 | 2 | 2 | 0.52 | 1 | 0.905 | 2.079 | 2.189 | 1.922 | 0.648 | 1.235 | 1.397 | MCG15083 OS=Mus musculus GN=Try5 PE=1 SV=1 |
| 132 | 1 | UniProt n | Q9Z1R9 | 753 | 29712 | 38 | 38 | 2 | 2 | 1.03 | 1 | 0.887 | 1.85 | 2.086 | 1.865 | 0.7 | 1.180 | 1.396 | MCG124046 OS=Mus musculus GN=Prss1 PE=1 SV=1 |
| 124 | 1 | UniProt n | A2AVA0 | 819 | 443177 | 37 | 37 | 21 | 21 | 0.24 | 1 | 0.903 | 0.938 | 1.393 | 1.374 | 1.415 | 0.946 | 1.394 | Sushi, von Willebrand factor type A, EGF and pentraxin domain-containing protein 1 OS=Mus musculus GN=Svep1 PE=1 SV=1 |
| 29 | 2 | UniProt n | Q6IFX2 | 1556 | 55854 | 54 | 54 | 16 | 16 | 4.29 | 1 | 1.108 | 1.33 | 1.175 | 1.692 | 1.362 | 1.138 | 1.394 | Keratin, type I cytoskeletal 42 OS=Mus musculus GN=Krt42 PE=1 SV=1 |
| 28 | 3 | UniProt n | E9PWF0 | 400 | 109796 | 20 | 20 | 13 | 13 | 0.72 | 1 | 0.934 | 1.037 | 1.439 | 1.362 | 1.372 | 0.989 | 1.391 | Thrombospondin-3 OS=Mus musculus GN=Thbs3 PE=1 SV=1 |
| 132 | 3 | UniProt n | Q792Y8 | 269 | 29696 | 17 | 17 | 3 | 3 | 0.76 | 1 | 0.908 | 2.111 | 2.193 | 1.923 | 0.637 | 1.242 | 1.390 | MCG15081 OS=Mus musculus GN=Gm10334 PE=3 SV=1 |
| 758 | 1 | UniProt n | Q9D8C4 | 72 | 36603 | 1 | 1 | 1 | 1 | 0.12 | 1 | 2.246 | 1.268 | --- | 1.052 | 1.836 | 1.417 | 1.390 | Interferon-induced 35 kDa protein homolog OS=Mus musculus GN=Ifi35 PE=1 SV=3 |
| 1139 | 1 | UniProt n | P97464 | 33 | 101148 | 2 | 2 | 2 | 2 | 0.09 | 1 | 0.88 | 1.033 | 1.456 | 1.303 | 1.404 | 0.969 | 1.386 | Exostosin-1 OS=Mus musculus GN=Ext1 PE=1 SV=1 |
| 9 | 1 | UniProt n | P21460 | 5719 | 18443 | 161 | 161 | 8 | 8 | 36.19 | 1 | 0.971 | 0.966 | 1.389 | 1.39 | 1.368 | 0.979 | 1.382 | Cystatin-C OS=Mus musculus GN=Cst3 PE=1 SV=2 |
| 226 | 1 | UniProt n | P07141 | 384 | 68768 | 24 | 24 | 10 | 10 | 1.09 | 1 | 0.984 | 0.916 | 1.316 | 1.439 | 1.383 | 0.966 | 1.378 | Macrophage colony-stimulating factor 1 OS=Mus musculus GN=Csf1 PE=1 SV=2 |
| 384 | 1 | UniProt n | P17439 | 220 | 64600 | 8 | 8 | 4 | 4 | 0.39 | 1 | 0.991 | 0.97 | 1.335 | 1.41 | 1.383 | 0.987 | 1.376 | Glucosylceramidase OS=Mus musculus GN=Gba PE=1 SV=1 |
| 1043 | 1 | UniProt n | Q9DAW6 | 41 | 69532 | 1 | 1 | 1 | 1 | 0.06 | 1 | 0.947 | 0.931 | 1.316 | 1.226 | 1.572 | 0.959 | 1.364 | U4/U6 small nuclear ribonucleoprotein Prp4 OS=Mus musculus GN=Prpf4 PE=1 SV=1 |
| 19 | 13 | UniProt n | Q8VED5 | 291 | 67786 | 31 | 31 | 6 | 6 | 0.45 | 1 | 0.846 | 1.086 | 1.163 | 1.626 | 1.335 | 0.972 | 1.362 | Keratin, type II cytoskeletal 79 OS=Mus musculus GN=Krt79 PE=1 SV=2 |
| 985 | 1 | UniProt n | Q9DBH5 | 47 | 46156 | 3 | 3 | 1 | 1 | 0.1 | 1 | 0.938 | 1.048 | 1.26 | 1.529 | 1.308 | 0.994 | 1.361 | Vesicular integral-membrane protein VIP36 OS=Mus musculus GN=Lman2 PE=1 SV=2 |
| 3 | 6 | UniProt n | Q61245 | 363 | 213064 | 19 | 19 | 10 | 10 | 0.27 | 1 | 1.058 | 1.048 | 1.386 | 1.319 | 1.375 | 1.035 | 1.360 | Collagen alpha-1(XI) chain OS=Mus musculus GN=Col11a1 PE=1 SV=2 |
| 1152 | 1 | UniProt n | Q91UZ5 | 31 | 35715 | 1 | 1 | 1 | 1 | 0.12 | 1 | 0.901 | 0.981 | 1.207 | 1.357 | 1.496 | 0.960 | 1.348 | Inositol monophosphatase 2 OS=Mus musculus GN=Impa2 PE=1 SV=1 |
| 33 | 2 | UniProt n | F8WI14 | 2652 | 70096 | 99 | 99 | 19 | 19 | 2.56 | 1 | 0.918 | 0.849 | 1.39 | 1.291 | 1.347 | 0.920 | 1.342 | Extracellular matrix protein 1 OS=Mus musculus GN=Ecm1 PE=1 SV=1 |
| 29 | 5 | UniProt n | Q9QWL7 | 598 | 55359 | 41 | 41 | 18 | 18 | 3.96 | 1 | 1.161 | 1.212 | 1.111 | 1.572 | 1.378 | 1.121 | 1.340 | Keratin, type I cytoskeletal 17 OS=Mus musculus GN=Krt17 PE=1 SV=3 |
| 59 | 1 | UniProt n | Q62009 | 1557 | 113714 | 58 | 58 | 22 | 22 | 1.64 | 1 | 1.037 | 1.049 | 1.326 | 1.327 | 1.362 | 1.028 | 1.338 | Periostin OS=Mus musculus GN=Postn PE=1 SV=2 |
| 33 | 1 | UniProt n | Q61508 | 2827 | 70210 | 109 | 109 | 19 | 19 | 2.54 | 1 | 0.916 | 0.846 | 1.381 | 1.283 | 1.333 | 0.919 | 1.332 | Extracellular matrix protein 1 OS=Mus musculus GN=Ecm1 PE=1 SV=2 |
| 1068 | 1 | UniProt n | A0A0R4J1 | 39 | 69730 | 1 | 1 | 1 | 1 | 0.06 | 1 | 0.985 | 1.049 | 1.452 | 1.191 | 1.362 | 1.011 | 1.331 | Immunoglobulin superfamily member 8 OS=Mus musculus GN=Igsf8 PE=1 SV=1 |
| 451 | 1 | UniProt n | A2APM1 | 171 | 79196 | 6 | 6 | 2 | 2 | 0.11 | 1 | 0.989 | 0.923 | 1.31 | 1.3 | 1.377 | 0.970 | 1.329 | CD44 antigen OS=Mus musculus GN=Cd44 PE=1 SV=1 |
| 1049 | 1 | UniProt n | E9PUN6 | 40 | 251708 | 1 | 1 | 1 | 1 | 0.02 | 1 | 0.571 | 0.636 | 1.305 | 1.381 | 1.301 | 0.713 | 1.328 | A disintegrin-like and metallopeptidase (reprolysin type) with thrombospondin type 1 motif, 9 OS=Mus musculus GN=Adamts9 PE=1 SV=1 |
| 732 | 1 | UniProt n | Q9Z175 | 76 | 93693 | 3 | 3 | 3 | 3 | 0.15 | 1 | 0.958 | 1.08 | 1.344 | 1.316 | 1.312 | 1.011 | 1.324 | Lysyl oxidase homolog 3 OS=Mus musculus GN=Loxl3 PE=1 SV=2 |
| 961 | 1 | UniProt n | Q52KC3 | 48 | 97296 | 1 | 1 | 1 | 1 | 0.04 | 1 | 1.029 | 1.11 | 1.304 | 1.432 | 1.234 | 1.045 | 1.321 | DNA helicase OS=Mus musculus GN=Mcm5 PE=1 SV=1 |
| 1197 | 1 | UniProt n | Q8VDF2 | 28 | 107346 | 1 | 1 | 1 | 1 | 0.04 | 1 | 1.071 | 1.018 | 1.212 | 1.505 | 1.254 | 1.029 | 1.318 | E3 ubiquitin-protein ligase UHRF1 OS=Mus musculus GN=Uhrf1 PE=1 SV=2 |
| 569 | 1 | UniProt n | P18406 | 116 | 52250 | 4 | 4 | 2 | 2 | 0.18 | 1 | 0.966 | 0.938 | 1.175 | 1.415 | 1.356 | 0.968 | 1.311 | Protein CYR61 OS=Mus musculus GN=Cyr61 PE=1 SV=1 |
| 138 | 1 | UniProt n | P01887 | 713 | 16904 | 41 | 41 | 3 | 3 | 2.43 | 1 | 1.054 | 0.978 | 1.264 | 1.182 | 1.457 | 1.010 | 1.296 | Beta-2-microglobulin OS=Mus musculus GN=B2m PE=1 SV=2 |
| 1127 | 1 | UniProt n | P62889 | 33 | 17477 | 1 | 1 | 1 | 1 | 0.27 | 1 | 1.084 | 1.259 | 1.179 | 1.505 | 1.219 | 1.109 | 1.293 | 60S ribosomal protein L30 OS=Mus musculus GN=Rpl30 PE=1 SV=2 |
| 661 | 1 | UniProt n | Q3UZG4 | 89 | 47026 | 3 | 3 | 3 | 3 | 0.31 | 1 | 1.058 | 1.161 | 1.251 | 1.469 | 1.174 | 1.071 | 1.292 | Aminoacyl tRNA synthase complex-interacting multifunctional protein 1 OS=Mus musculus GN=Aimp1 PE=1 SV=1 |
| 346 | 1 | UniProt n | Q61805 | 246 | 61724 | 9 | 9 | 5 | 5 | 0.51 | 1 | 1.032 | 1.112 | 1.27 | 1.233 | 1.359 | 1.047 | 1.286 | Lipopolysaccharide-binding protein OS=Mus musculus GN=Lbp PE=1 SV=2 |
| 1214 | 1 | UniProt n | Q8K0C9 | 27 | 50752 | 1 | 1 | 1 | 1 | 0.09 | 1 | 1.082 | 1.149 | 1.352 | 1.324 | 1.187 | 1.075 | 1.286 | GDP-mannose 4,6 dehydratase OS=Mus musculus GN=Gmds PE=1 SV=1 |
| 994 | 1 | UniProt n | Q61477 | 45 | 21988 | 4 | 4 | 1 | 1 | 0.21 | 1 | 1.066 | 1.225 | 1.333 | 1.273 | 1.25 | 1.093 | 1.285 | Neuroblastoma suppressor of tumorigenicity 1 OS=Mus musculus GN=Nbl1 PE=1 SV=2 |
| 499 | 1 | UniProt n | Q8R2G6 | 146 | 132812 | 3 | 3 | 3 | 3 | 0.1 | 1 | 0.969 | 1.01 | 1.267 | 1.326 | 1.262 | 0.993 | 1.285 | Coiled-coil domain-containing protein 80 OS=Mus musculus GN=Ccdc80 PE=1 SV=2 |
| 19 | 8 | UniProt n | Q3TTY5 | 911 | 80982 | 68 | 68 | 10 | 10 | 0.87 | 1 | 0.741 | 1.057 | 1.13 | 1.615 | 1.155 | 0.922 | 1.282 | Keratin, type II cytoskeletal 2 epidermal OS=Mus musculus GN=Krt2 PE=1 SV=1 |
| 624 | 1 | UniProt n | A0A0A6Y) | 98 | 50411 | 2 | 2 | 1 | 1 | 0.18 | 1 | 0.753 | 0.782 | 1.366 | 1.198 | 1.285 | 0.838 | 1.281 | cAMP-dependent protein kinase type II-alpha regulatory subunit OS=Mus musculus GN=Prkar2a PE=1 SV=1 |
| 6 | 1 | UniProt n | Q61398 | 7189 | 57685 | 219 | 219 | 20 | 20 | 6.77 | 1 | 1.006 | 0.992 | 1.302 | 1.235 | 1.3 | 0.999 | 1.279 | Procollagen C-endopeptidase enhancer 1 OS=Mus musculus GN=Pcolce PE=1 SV=2 |
| 914 | 1 | UniProt n | Q9DBG9 | 53 | 16148 | 1 | 1 | 1 | 1 | 0.29 | 1 | 1.222 | 1.092 | 1.08 | 2.191 | 0.876 | 1.101 | 1.275 | Tax1-binding protein 3 OS=Mus musculus GN=Tax1bp3 PE=1 SV=1 |
| 593 | 1 | UniProt n | P15535 | 106 | 50485 | 3 | 3 | 3 | 3 | 0.29 | 1 | 0.938 | 0.936 | 1.322 | 1.246 | 1.257 | 0.958 | 1.275 | Beta-1,4-galactosyltransferase 1 OS=Mus musculus GN=B4galt1 PE=1 SV=1 |
| 759 | 1 | UniProt n | Q8K4Q8 | 72 | 97836 | 4 | 4 | 2 | 2 | 0.09 | 1 | 0.843 | 0.894 | 1.267 | 1.294 | 1.26 | 0.910 | 1.274 | Collectin-12 OS=Mus musculus GN=Colec12 PE=1 SV=1 |
| 93 | 1 | UniProt n | G5E8D6 | 1080 | 54463 | 30 | 30 | 12 | 12 | 1.74 | 1 | 1.009 | 1.032 | 1.322 | 1.219 | 1.281 | 1.014 | 1.273 | EGF-containing fibulin-like extracellular matrix protein 2 OS=Mus musculus GN=Efemp2 PE=1 SV=1 |
| 836 | 1 | UniProt n | Q9WV92 | 61 | 124632 | 1 | 1 | 1 | 1 | 0.03 | 1 | 0.814 | 0.855 | 1.27 | 1.261 | 1.281 | 0.886 | 1.271 | Band 4.1-like protein 3 OS=Mus musculus GN=Epb41l3 PE=1 SV=1 |
| 390 | 1 | UniProt n | Q9CYN9 | 214 | 43630 | 13 | 13 | 9 | 9 | 1.39 | 1 | 1.009 | 1.045 | 1.228 | 1.255 | 1.32 | 1.018 | 1.267 | Renin receptor OS=Mus musculus GN=Atp6ap2 PE=1 SV=2 |
| 638 | 1 | UniProt n | P10404 | 94 | 79614 | 2 | 2 | 2 | 2 | 0.11 | 1 | 1.027 | 0.921 | 1.261 | 1.236 | 1.305 | 0.982 | 1.267 | MLV-related proviral Env polyprotein OS=Mus musculus PE=1 SV=3 |
| 441 | 1 | UniProt n | Q8VE43 | 176 | 37097 | 6 | 6 | 4 | 4 | 0.76 | 1 | 1.01 | 1.021 | 1.343 | 1.242 | 1.217 | 1.010 | 1.266 | Meteorin-like protein OS=Mus musculus GN=Metrnl PE=1 SV=1 |
| 209 | 1 | UniProt n | Q61112 | 418 | 49384 | 18 | 18 | 8 | 8 | 0.98 | 1 | 1 | 0.995 | 1.349 | 1.244 | 1.206 | 0.998 | 1.265 | 45 kDa calcium-binding protein OS=Mus musculus GN=Sdf4 PE=1 SV=1 |
| 897 | 1 | UniProt n | Q8K1S3 | 55 | 115930 | 2 | 2 | 2 | 2 | 0.08 | 1 | 0.888 | 0.976 | 1.35 | 1.273 | 1.173 | 0.953 | 1.263 | Netrin receptor UNC5B OS=Mus musculus GN=Unc5b PE=1 SV=1 |
| 263 | 1 | UniProt n | A0A140LJI | 333 | 22035 | 10 | 10 | 5 | 5 | 1.58 | 1 | 0.858 | 0.877 | 1.337 | 1.2 | 1.255 | 0.910 | 1.263 | Serine protease 23 (Fragment) OS=Mus musculus GN=Prss23 PE=1 SV=1 |
| 3 | 4 | UniProt n | O88207 | 3212 | 213624 | 104 | 104 | 22 | 22 | 0.85 | 1 | 0.974 | 1.025 | 1.283 | 1.248 | 1.249 | 0.999 | 1.260 | Collagen alpha-1(V) chain OS=Mus musculus GN=Col5a1 PE=1 SV=2 |
| 98 | 7 | UniProt n | E9PX63 | 532 | 43947 | 18 | 18 | 5 | 5 | 0.96 | 1 | 0.943 | 0.962 | 1.325 | 1.271 | 1.187 | 0.968 | 1.260 | Histocompatibility 2, Q region locus 1 OS=Mus musculus GN=H2-Q1 PE=3 SV=1 |
| 19 | 5 | UniProt n | P50446 | 1069 | 69309 | 73 | 73 | 16 | 16 | 2 | 1 | 0.902 | 1.108 | 1.088 | 1.546 | 1.185 | 1.000 | 1.258 | Keratin, type II cytoskeletal 6A OS=Mus musculus GN=Krt6a PE=1 SV=3 |
| 19 | 3 | UniProt n | Q922U2 | 1641 | 70734 | 110 | 110 | 17 | 17 | 3.21 | 1 | 0.81 | 1.076 | 1.11 | 1.597 | 1.112 | 0.955 | 1.254 | Keratin, type II cytoskeletal 5 OS=Mus musculus GN=Krt5 PE=1 SV=1 |
| 980 | 1 | UniProt n | Q7TPC1 | 47 | 60399 | 4 | 4 | 3 | 3 | 0.23 | 1 | 0.855 | 0.874 | 1.239 | 1.197 | 1.318 | 0.907 | 1.250 | Corneodesmosin OS=Mus musculus GN=Cdsn PE=2 SV=2 |
| 19 | 4 | UniProt n | P04104 | 1117 | 76626 | 61 | 61 | 7 | 7 | 0.74 | 1 | 0.816 | 1.167 | 1.173 | 1.509 | 1.104 | 0.984 | 1.250 | Keratin, type II cytoskeletal 1 OS=Mus musculus GN=Krt1 PE=1 SV=4 |
| 1078 | 1 | UniProt n | V9GX81 | 38 | 80881 | 3 | 3 | 1 | 1 | 0.05 | 1 | 0.914 | 0.873 | 1.325 | 1.266 | 1.158 | 0.928 | 1.248 | Maestro heat-like repeat family member 6 OS=Mus musculus GN=Mroh6 PE=4 SV=1 |
| 1241 | 1 | UniProt n | A0A0U1RI | 24 | 27148 | 1 | 1 | 1 | 1 | 0.17 | 1 | 0.63 | 0.908 | 0.962 | 1.451 | 1.391 | 0.830 | 1.248 | Aminopeptidase N (Fragment) OS=Mus musculus GN=Anpep PE=1 SV=1 |
| 706 | 1 | UniProt n | F8VQA4 | 81 | 126794 | 3 | 3 | 3 | 3 | 0.11 | 1 | 0.971 | 1.041 | 1.301 | 1.226 | 1.205 | 1.004 | 1.243 | Peptidyl-glycine alpha-amidating monooxygenase OS=Mus musculus GN=Pam PE=1 SV=1 |
| 409 | 1 | UniProt n | A2ATR8 | 202 | 44525 | 4 | 4 | 3 | 3 | 0.33 | 1 | 1.055 | 1.046 | 1.213 | 1.236 | 1.275 | 1.033 | 1.241 | Plasma protease C1 inhibitor OS=Mus musculus GN=Serping1 PE=1 SV=1 |
| 1028 | 1 | UniProt n | O55029 | 42 | 120110 | 2 | 2 | 2 | 2 | 0.07 | 1 | 0.882 | 1.092 | 1.126 | 1.376 | 1.231 | 0.988 | 1.240 | Coatomer subunit beta' OS=Mus musculus GN=Copb2 PE=1 SV=2 |
| 975 | 1 | UniProt n | F8WIP8 | 47 | 38440 | 3 | 3 | 2 | 2 | 0.39 | 1 | 0.854 | 0.888 | 1.08 | 1.399 | 1.262 | 0.912 | 1.240 | Osteopontin OS=Mus musculus GN=Spp1 PE=1 SV=1 |
| 1208 | 1 | UniProt n | Q9QYB5 | 27 | 93910 | 1 | 1 | 1 | 1 | 0.05 | 1 | 1.046 | 1.077 | 1.164 | 1.399 | 1.169 | 1.041 | 1.239 | Gamma-adducin OS=Mus musculus GN=Add3 PE=1 SV=2 |
| 98 | 1 | UniProt n | Q8HWB2 | 1056 | 43169 | 40 | 40 | 10 | 10 | 2.56 | 1 | 1.002 | 1.045 | 1.266 | 1.21 | 1.219 | 1.015 | 1.231 | Histocompatibility 2, Q region locus 4 OS=Mus musculus GN=H2-Q4 PE=1 SV=1 |
| 1024 | 1 | UniProt n | O35405 | 43 | 59894 | 1 | 1 | 1 | 1 | 0.07 | 1 | 0.963 | 1.029 | 1.196 | 1.198 | 1.3 | 0.997 | 1.230 | Phospholipase D3 OS=Mus musculus GN=Pld3 PE=1 SV=1 |
| 647 | 1 | UniProt n | D3YVW2 | 91 | 91463 | 1 | 1 | 1 | 1 | 0.05 | 1 | 0.944 | 1.107 | 1.242 | 1.17 | 1.281 | 1.015 | 1.230 | Golgi integral membrane protein 4 OS=Mus musculus GN=Golim4 PE=1 SV=1 |
| 557 | 1 | UniProt n | O35988 | 120 | 24861 | 5 | 5 | 2 | 2 | 0.4 | 1 | 0.848 | 0.93 | 1.24 | 1.277 | 1.171 | 0.924 | 1.229 | Syndecan-4 OS=Mus musculus GN=Sdc4 PE=1 SV=1 |
| 1237 | 1 | UniProt n | Q9EQC7 | 24 | 30578 | 1 | 1 | 1 | 1 | 0.15 | 1 | 0.812 | 0.865 | 1.283 | 1.257 | 1.147 | 0.889 | 1.228 | Follistatin-related protein 3 OS=Mus musculus GN=Fstl3 PE=1 SV=1 |
| 1161 | 1 | UniProt n | Q8R555 | 31 | 75441 | 1 | 1 | 1 | 1 | 0.06 | 1 | 1.179 | 1.268 | 1.241 | 1.133 | 1.315 | 1.143 | 1.227 | Cartilage acidic protein 1 OS=Mus musculus GN=Crtac1 PE=2 SV=1 |
| 970 | 1 | UniProt n | Q8R550 | 40 | 98642 | 2 | 2 | 2 | 2 | 0.09 | 1 | 1.046 | 1.087 | 1.165 | 1.22 | 1.289 | 1.044 | 1.224 | SH3 domain-containing kinase-binding protein 1 OS=Mus musculus GN=Sh3kbp1 PE=1 SV=1 |
| 19 | 6 | UniProt n | Q8BGZ7 | 955 | 68101 | 62 | 62 | 12 | 12 | 1.24 | 1 | 0.838 | 1.072 | 1.08 | 1.503 | 1.126 | 0.965 | 1.223 | Keratin, type II cytoskeletal 75 OS=Mus musculus GN=Krt75 PE=1 SV=1 |

| | | | | | | | | | | | | | | | | | | | | |
|---|---|---|---|---|---|---|---|---|---|---|---|---|---|---|---|---|---|---|---|---|
| 1195 | 1 | UniProt | n | A0A0B4J1 | 29 | 222414 | 1 | 1 | 1 | 1 | 0.02 | 1 | 0.924 | 1.096 | 1.215 | 1.289 | 1.165 | 1.004 | 1.222 | Trichohyalin OS=Mus musculus GN=Tchh PE=4 SV=1 |
| 531 | 1 | UniProt | n | A0A0U1RI | 128 | 17267 | 6 | 6 | 3 | 3 | 1.06 | 1 | 0.944 | 1.026 | 1.252 | 1.162 | 1.248 | 0.989 | 1.220 | Complement C1q tumor necrosis factor-related protein 5 (Fragment) OS=Mus musculus GN=C1qtnf5 PE=1 SV=2 |
| 1171 | 1 | UniProt | n | Q03145 | 30 | 125188 | 1 | 1 | 1 | 1 | 0.03 | 1 | 1.267 | 1.251 | 1.294 | 1.193 | 1.146 | 1.166 | 1.209 | Ephrin type-A receptor 2 OS=Mus musculus GN=Epha2 PE=1 SV=3 |
| 799 | 1 | UniProt | n | Q8CIB5 | 65 | 97163 | 2 | 2 | 2 | 2 | 0.09 | 1 | 0.926 | 1.136 | 1.259 | 1.274 | 1.098 | 1.017 | 1.208 | Fermitin family homolog 2 OS=Mus musculus GN=Fermt2 PE=1 SV=1 |
| 1061 | 1 | UniProt | n | Q32ME9 | 39 | 109498 | 1 | 1 | 1 | 1 | 0.04 | 1 | 0.727 | 1.079 | 1.058 | 1.577 | 1.037 | 0.922 | 1.201 | Desmocollin 1, isoform CRA_a OS=Mus musculus GN=Dsc1 PE=1 SV=1 |
| 1015 | 1 | UniProt | n | D3YX62 | 44 | 33716 | 1 | 1 | 1 | 1 | 0.13 | 1 | 0.912 | 0.963 | 1.022 | 0.929 | 1.813 | 0.958 | 1.198 | Heme oxygenase 2 (Fragment) OS=Mus musculus GN=Hmox2 PE=1 SV=1 |
| 3 | 3 | UniProt | n | Q3U962 | 6574 | 164313 | 168 | 168 | 40 | 40 | 3.48 | 1 | 1.04 | 1.039 | 1.205 | 1.189 | 1.187 | 1.026 | 1.194 | Collagen alpha-2(V) chain OS=Mus musculus GN=Col5a2 PE=1 SV=1 |
| 19 | 11 | UniProt | n | Q3UV17 | 493 | 74171 | 49 | 49 | 6 | 6 | 0.41 | 1 | 0.81 | 1.054 | 1.066 | 1.437 | 1.11 | 0.949 | 1.194 | Keratin, type II cytoskeletal 2 oral OS=Mus musculus GN=Krt76 PE=1 SV=1 |
| 98 | 6 | UniProt | n | P04223 | 534 | 45896 | 18 | 18 | 6 | 6 | 0.74 | 1 | 0.956 | 0.957 | 1.238 | 1.187 | 1.147 | 0.971 | 1.190 | H-2 class I histocompatibility antigen, K-K alpha chain OS=Mus musculus GN=H2-K1 PE=1 SV=1 |
| 907 | 1 | UniProt | n | Q8JZK9 | 54 | 67469 | 2 | 2 | 2 | 2 | 0.13 | 1 | 0.878 | 0.859 | 2.304 | 0.796 | 0.916 | 0.910 | 1.189 | Hydroxymethylglutaryl-CoA synthase, cytoplasmic OS=Mus musculus GN=Hmgcs1 PE=1 SV=1 |
| 867 | 1 | UniProt | n | Q6PB93 | 58 | 75110 | 2 | 2 | 2 | 2 | 0.12 | 1 | 0.928 | 1.076 | 1.265 | 1.233 | 1.072 | 1.000 | 1.187 | Polypeptide N-acetylgalactosaminyltransferase 2 OS=Mus musculus GN=Galnt2 PE=1 SV=1 |
| 19 | 7 | UniProt | n | Q6IFZ6 | 928 | 70190 | 54 | 54 | 7 | 7 | 0.72 | 1 | 0.751 | 0.994 | 1.131 | 1.437 | 1.023 | 0.907 | 1.185 | Keratin, type II cytoskeletal 1b OS=Mus musculus GN=Krt77 PE=1 SV=1 |
| 951 | 1 | UniProt | n | P97350 | 49 | 94162 | 1 | 1 | 1 | 1 | 0.05 | 1 | 0.84 | 1.246 | 1.201 | 1.332 | 1.035 | 1.015 | 1.183 | Plakophilin-1 OS=Mus musculus GN=Pkp1 PE=1 SV=1 |
| 191 | 1 | UniProt | n | Q9Z0J0 | 472 | 20966 | 19 | 19 | 5 | 5 | 1.71 | 1 | 1.081 | 1.05 | 1.175 | 1.227 | 1.148 | 1.043 | 1.183 | Epididymal secretory protein E1 OS=Mus musculus GN=Npc2 PE=1 SV=1 |
| 1148 | 1 | UniProt | n | A0A1D5RI | 32 | 91397 | 2 | 2 | 2 | 2 | 0.1 | 1 | 1.096 | 1.15 | 1.22 | 1.097 | 1.235 | 1.080 | 1.182 | Nuclear pore complex protein Nup93 OS=Mus musculus GN=Nup93 PE=1 SV=1 |
| 977 | 1 | UniProt | n | A0A0J9YV | 47 | 62355 | 1 | 1 | 1 | 1 | 0.07 | 1 | 0.868 | 0.908 | 1.327 | 1.071 | 1.152 | 0.924 | 1.179 | Platelet-derived growth factor receptor alpha OS=Mus musculus GN=Pdgfra PE=1 SV=1 |
| 166 | 1 | UniProt | n | O88968 | 586 | 54008 | 30 | 30 | 13 | 13 | 1.99 | 1 | 1.003 | 0.99 | 1.198 | 1.149 | 1.188 | 0.998 | 1.178 | Transcobalamin-2 OS=Mus musculus GN=Tcn2 PE=1 SV=1 |
| 19 | 9 | UniProt | n | E9Q1Z0 | 899 | 68853 | 42 | 42 | 10 | 10 | 1.09 | 1 | 0.845 | 1.047 | 1.021 | 1.459 | 1.095 | 0.960 | 1.177 | Keratin 90 OS=Mus musculus GN=Krt90 PE=1 SV=1 |
| 98 | 8 | UniProt | n | P01902 | 460 | 45648 | 16 | 16 | 6 | 6 | 0.91 | 1 | 1.066 | 1.044 | 1.152 | 1.165 | 1.207 | 1.036 | 1.174 | H-2 class I histocompatibility antigen, K-D alpha chain OS=Mus musculus GN=H2-K1 PE=1 SV=1 |
| 107 | 1 | UniProt | n | J3QPG5 | 974 | 74704 | 51 | 51 | 18 | 18 | 2.11 | 1 | 1.183 | 1.065 | 1.147 | 1.213 | 1.153 | 1.080 | 1.171 | Prosaposin OS=Mus musculus GN=Psap PE=1 SV=1 |
| 884 | 1 | UniProt | n | Q9WV91 | 56 | 109784 | 5 | 5 | 4 | 4 | 0.17 | 1 | 0.832 | 0.943 | 1.167 | 1.172 | 1.165 | 0.922 | 1.168 | Prostaglandin F2 receptor negative regulator OS=Mus musculus GN=Ptgfrn PE=1 SV=2 |
| 115 | 1 | UniProt | n | P82198 | 882 | 86920 | 32 | 32 | 16 | 16 | 1.29 | 1 | 1.038 | 1.002 | 1.135 | 1.144 | 1.227 | 1.013 | 1.168 | Transforming growth factor-beta-induced protein ig-h3 OS=Mus musculus GN=Tgfbi PE=1 SV=1 |
| 95 | 1 | UniProt | n | P12023 | 1070 | 100576 | 38 | 38 | 13 | 13 | 0.88 | 1 | 0.938 | 0.915 | 1.24 | 1.093 | 1.174 | 0.950 | 1.167 | Amyloid beta A4 protein OS=Mus musculus GN=App PE=1 SV=3 |
| 573 | 1 | UniProt | n | Q9D554 | 115 | 71858 | 3 | 3 | 1 | 1 | 0.13 | 1 | 0.915 | 1.052 | 0.671 | 3.467 | 0.678 | 0.987 | 1.164 | Splicing factor 3A subunit 3 OS=Mus musculus GN=Sf3a3 PE=1 SV=2 |
| 413 | 1 | UniProt | n | Q9CZT5 | 200 | 76786 | 9 | 9 | 4 | 4 | 0.32 | 1 | 0.899 | 0.954 | 1.326 | 1.083 | 1.094 | 0.950 | 1.163 | Vasorin OS=Mus musculus GN=Vasn PE=2 SV=2 |
| 367 | 1 | UniProt | n | Q8VCF1 | 231 | 51450 | 10 | 10 | 6 | 6 | 0.64 | 1 | 0.938 | 0.948 | 1.166 | 1.121 | 1.196 | 0.962 | 1.161 | Soluble calcium-activated nucleotidase 1 OS=Mus musculus GN=Cant1 PE=2 SV=1 |
| 366 | 1 | UniProt | n | Q9R0Q7 | 231 | 22894 | 4 | 4 | 2 | 2 | 0.73 | 1 | 1.144 | 1.224 | 1.141 | 1.186 | 1.147 | 1.119 | 1.158 | Prostaglandin E synthase 3 OS=Mus musculus GN=Ptges3 PE=1 SV=1 |
| 98 | 5 | UniProt | n | P14426 | 596 | 44216 | 24 | 24 | 7 | 7 | 1.15 | 1 | 0.93 | 0.935 | 1.227 | 1.147 | 1.1 | 0.954 | 1.157 | H-2 class I histocompatibility antigen, D-K alpha chain OS=Mus musculus GN=H2-D1 PE=1 SV=1 |
| 883 | 1 | UniProt | n | P42703-2 | 56 | 93273 | 1 | 1 | 1 | 1 | 0.05 | 1 | 0.655 | 0.825 | 1.342 | 1.388 | 0.829 | 0.815 | 1.156 | Isoform 2 of Leukemia inhibitory factor receptor OS=Mus musculus GN=Lifr |
| 867 | 2 | UniProt | n | O08832 | 39 | 76328 | 2 | 2 | 2 | 2 | 0.12 | 1 | 0.997 | 1.249 | 1.139 | 1.24 | 1.093 | 1.076 | 1.156 | Polypeptide N-acetylgalactosaminyltransferase 4 OS=Mus musculus GN=Galnt4 PE=2 SV=1 |
| 525 | 1 | UniProt | n | P10148 | 131 | 20150 | 21 | 21 | 4 | 4 | 1.28 | 1 | 1.117 | 1.019 | 1.096 | 1.173 | 1.196 | 1.044 | 1.154 | C-C motif chemokine 2 OS=Mus musculus GN=Ccl2 PE=1 SV=1 |
| 308 | 1 | UniProt | n | Q08761 | 286 | 90535 | 5 | 5 | 3 | 3 | 0.15 | 1 | 0.9 | 0.951 | 1.168 | 1.07 | 1.22 | 0.949 | 1.151 | Vitamin K-dependent protein S OS=Mus musculus GN=Pros1 PE=2 SV=1 |
| 625 | 1 | UniProt | n | O35664-3 | 98 | 33627 | 4 | 4 | 3 | 3 | 0.46 | 1 | 1.054 | 0.945 | 1.125 | 1.163 | 1.165 | 0.999 | 1.151 | Isoform 3 of Interferon alpha/beta receptor 2 OS=Mus musculus GN=Ifnar2 |
| 1229 | 1 | UniProt | n | A0A0R4J2 | 26 | 98689 | 2 | 2 | 2 | 2 | 0.09 | 1 | 0.845 | 0.886 | 1.195 | 1.15 | 1.106 | 0.908 | 1.150 | Protein-glutamine gamma-glutamyltransferase K OS=Mus musculus GN=Tgm1 PE=1 SV=1 |
| 1198 | 1 | UniProt | n | P26516 | 28 | 48123 | 2 | 2 | 2 | 2 | 0.19 | 1 | 1.025 | 1 | 1.057 | 1.265 | 1.129 | 1.008 | 1.147 | 26S proteasome non-ATPase regulatory subunit 7 OS=Mus musculus GN=Psmd7 PE=1 SV=2 |
| 126 | 1 | UniProt | n | E9Q0F0 | 812 | 121575 | 31 | 31 | 2 | 2 | 0.15 | 1 | 0.837 | 1.006 | 0.957 | 1.369 | 1.151 | 0.944 | 1.147 | Keratin 78 OS=Mus musculus GN=Krt78 PE=1 SV=1 |
| 445 | 1 | UniProt | n | Q60648 | 174 | 24829 | 8 | 8 | 4 | 4 | 0.96 | 1 | 1.08 | 1.349 | 0.952 | 1.35 | 1.172 | 1.134 | 1.146 | Ganglioside GM2 activator OS=Mus musculus GN=Gm2a PE=1 SV=2 |
| 502 | 1 | UniProt | n | P70428 | 144 | 94778 | 8 | 8 | 7 | 7 | 0.37 | 1 | 0.966 | 0.996 | 1.196 | 1.078 | 1.141 | 0.987 | 1.137 | Exostosin-2 OS=Mus musculus GN=Ext2 PE=1 SV=2 |
| 1146 | 1 | UniProt | n | Q3UKJ7 | 32 | 68052 | 1 | 1 | 1 | 1 | 0.06 | 1 | 0.639 | 0.654 | 2.773 | 0.805 | 0.656 | 0.748 | 1.136 | WD40 repeat-containing protein SMU1 OS=Mus musculus GN=Smu1 PE=2 SV=2 |
| 608 | 1 | UniProt | n | Z4YN97 | 102 | 13231 | 3 | 3 | 2 | 2 | 0.87 | 1 | 1.065 | 1.023 | 1.151 | 1.078 | 1.172 | 1.029 | 1.133 | Adenylate kinase isoenzyme 1 (Fragment) OS=Mus musculus GN=Ak1 PE=1 SV=1 |
| 761 | 1 | UniProt | n | A0A140LH | 71 | 57575 | 2 | 2 | 2 | 2 | 0.16 | 1 | 0.862 | 0.844 | 1.143 | 0.988 | 1.286 | 0.899 | 1.132 | Lysosomal Pro-X carboxypeptidase OS=Mus musculus GN=Prcp PE=1 SV=1 |
| 810 | 1 | UniProt | n | E9Q557 | 64 | 409822 | 4 | 4 | 4 | 4 | 0.04 | 1 | 0.944 | 1.087 | 1.055 | 1.229 | 1.113 | 1.009 | 1.130 | Desmoplakin OS=Mus musculus GN=Dsp PE=1 SV=1 |
| 386 | 1 | UniProt | n | Q02257 | 219 | 92385 | 11 | 11 | 5 | 5 | 0.38 | 1 | 0.903 | 1.008 | 1.105 | 1.172 | 1.107 | 0.969 | 1.128 | Junction plakoglobin OS=Mus musculus GN=Jup PE=1 SV=3 |
| 719 | 1 | UniProt | n | F8VQ06 | 78 | 151048 | 2 | 2 | 2 | 2 | 0.06 | 1 | 0.952 | 0.992 | 1.112 | 1.135 | 1.134 | 0.981 | 1.127 | Latent-transforming growth factor beta-binding protein 3 OS=Mus musculus GN=Ltbp3 PE=1 SV=1 |
| 1177 | 1 | UniProt | n | O35598 | 30 | 101996 | 2 | 2 | 2 | 2 | 0.09 | 1 | 0.831 | 0.853 | 1.211 | 1.195 | 0.989 | 0.892 | 1.127 | Disintegrin and metalloproteinase domain-containing protein 10 OS=Mus musculus GN=Adam10 PE=1 SV=2 |
| 1052 | 1 | UniProt | n | A0A1W2P | 40 | 184855 | 2 | 2 | 2 | 2 | 0.05 | 1 | 0.913 | 0.997 | 1.244 | 1.069 | 1.076 | 0.969 | 1.127 | Receptor-type tyrosine-protein phosphatase kappa OS=Mus musculus GN=Ptprk PE=1 SV=1 |
| 338 | 1 | UniProt | n | Q6IR41 | 250 | 32897 | 8 | 8 | 2 | 2 | 1.16 | 1 | 0.997 | 1.295 | 1.072 | 1.127 | 1.172 | 1.089 | 1.123 | Complement C1q tumor necrosis factor-related protein 6 OS=Mus musculus GN=C1qtnf6 PE=2 SV=1 |
| 410 | 1 | UniProt | n | Q3UF75 | 201 | 46080 | 7 | 7 | 4 | 4 | 0.44 | 1 | 1.1 | 1.237 | 1.11 | 1.168 | 1.091 | 1.108 | 1.123 | Alpha-parvin OS=Mus musculus GN=Parva PE=1 SV=1 |
| 882 | 1 | UniProt | n | P25118 | 56 | 57865 | 1 | 1 | 1 | 1 | 0.08 | 1 | 0.999 | 0.944 | 0.872 | 0.964 | 1.682 | 0.981 | 1.122 | Tumor necrosis factor receptor superfamily member 1A OS=Mus musculus GN=Tnfrsf1a PE=1 SV=1 |
| 29 | 3 | UniProt | n | Q61781 | 1553 | 60715 | 66 | 66 | 15 | 15 | 5.56 | 1 | 0.845 | 0.985 | 0.937 | 1.329 | 1.128 | 0.941 | 1.120 | Keratin, type I cytoskeletal 14 OS=Mus musculus GN=Krt14 PE=1 SV=2 |
| 98 | 3 | UniProt | n | P01901 | 598 | 46161 | 20 | 20 | 6 | 6 | 0.9 | 1 | 0.962 | 0.942 | 1.159 | 1.096 | 1.098 | 0.968 | 1.117 | H-2 class I histocompatibility antigen, K-B alpha chain OS=Mus musculus GN=H2-K1 PE=1 SV=1 |
| 255 | 1 | UniProt | n | P11152 | 347 | 64183 | 9 | 9 | 5 | 5 | 0.39 | 1 | 0.963 | 1.084 | 1.07 | 1.15 | 1.133 | 1.014 | 1.117 | Lipoprotein lipase OS=Mus musculus GN=Lpl PE=1 SV=3 |
| 1124 | 1 | UniProt | n | Q61735 | 34 | 38965 | 1 | 1 | 1 | 1 | 0.11 | 1 | 0.869 | 0.934 | 1.17 | 1.062 | 1.115 | 0.933 | 1.115 | Leukocyte surface antigen CD47 OS=Mus musculus GN=Cd47 PE=1 SV=2 |
| 19 | 10 | UniProt | n | Q6NXH9 | 811 | 70028 | 39 | 39 | 6 | 6 | 0.62 | 1 | 0.728 | 0.92 | 0.984 | 1.386 | 1.013 | 0.875 | 1.114 | Keratin, type II cytoskeletal 73 OS=Mus musculus GN=Krt73 PE=1 SV=1 |
| 111 | 1 | UniProt | n | Q3UQ28 | 923 | 181411 | 41 | 41 | 24 | 24 | 0.8 | 1 | 0.982 | 0.996 | 1.092 | 1.083 | 1.163 | 0.993 | 1.112 | Peroxidasin homolog OS=Mus musculus GN=Pxdn PE=1 SV=2 |
| 536 | 1 | UniProt | n | Q9D1A2 | 127 | 64661 | 5 | 5 | 3 | 3 | 0.3 | 1 | 1.083 | 1.374 | 1.474 | 1.088 | 0.85 | 1.142 | 1.109 | Cytosolic non-specific dipeptidase OS=Mus musculus GN=Cndp2 PE=1 SV=1 |
| 991 | 1 | UniProt | n | Q9Z2L6 | 46 | 62614 | 2 | 2 | 2 | 2 | 0.14 | 1 | 0.949 | 0.921 | 1.101 | 1.155 | 1.06 | 0.956 | 1.105 | Multiple inositol polyphosphate phosphatase 1 OS=Mus musculus GN=Minpp1 PE=1 SV=3 |
| 432 | 1 | UniProt | n | P37804 | 182 | 28083 | 14 | 14 | 7 | 7 | 2.3 | 1 | 1.044 | 1.022 | 1.119 | 1.222 | 0.973 | 1.022 | 1.100 | Transgelin OS=Mus musculus GN=Tagln PE=1 SV=3 |
| 1100 | 1 | UniProt | n | P32233 | 36 | 52277 | 1 | 1 | 1 | 1 | 0.08 | 1 | 0.943 | 1.094 | 1.182 | 1.121 | 1 | 1.010 | 1.098 | Developmentally-regulated GTP-binding protein 1 OS=Mus musculus GN=Drg1 PE=1 SV=1 |
| 1094 | 1 | UniProt | n | A0A1L1SR | 36 | 102255 | 1 | 1 | 1 | 1 | 0.04 | 1 | 0.821 | 1.059 | 0.925 | 1.053 | 1.357 | 0.954 | 1.097 | Low-density lipoprotein receptor OS=Mus musculus GN=Ldlr PE=1 SV=1 |
| 4 | 2 | UniProt | n | Q05793 | 9788 | 428156 | 385 | 385 | 91 | 91 | 2.26 | 1 | 0.958 | 0.981 | 1.122 | 1.059 | 1.109 | 0.980 | 1.096 | Basement membrane-specific heparan sulfate proteoglycan core protein OS=Mus musculus GN=Hspg2 PE=1 SV=1 |
| 4 | 1 | UniProt | n | E9PZ16 | 11264 | 502469 | 426 | 426 | 101 | 101 | 1.96 | 1 | 0.962 | 0.98 | 1.12 | 1.056 | 1.111 | 0.981 | 1.095 | Basement membrane-specific heparan sulfate proteoglycan core protein OS=Mus musculus GN=Hspg2 PE=1 SV=1 |
| 1200 | 1 | UniProt | n | Q80U93 | 28 | 241008 | 1 | 1 | 1 | 1 | 0.02 | 1 | 0.905 | 1.002 | 1.131 | 1.07 | 1.08 | 0.968 | 1.093 | Nuclear pore complex protein Nup214 OS=Mus musculus GN=Nup214 PE=1 SV=2 |
| 1215 | 1 | UniProt | n | Q8C9X6 | 26 | 106329 | 1 | 1 | 1 | 1 | 0.04 | 1 | 0.918 | 0.943 | 1.368 | 1.101 | 0.86 | 0.953 | 1.090 | Enhancer of polycomb homolog 1 OS=Mus musculus GN=Epc1 PE=1 SV=1 |
| 127 | 1 | UniProt | n | Q3V1G4 | 794 | 93175 | 23 | 23 | 9 | 9 | 0.51 | 1 | 1.01 | 1.054 | 1.16 | 1.058 | 1.05 | 1.021 | 1.088 | Olfactomedin-like protein 2B OS=Mus musculus GN=Olfml2b PE=1 SV=2 |
| 3 | 1 | UniProt | n | P11087 | 22815 | 155811 | 782 | 782 | 73 | 73 | 19.58 | 1 | 1.033 | 1.029 | 1.072 | 1.06 | 1.126 | 1.021 | 1.086 | Collagen alpha-1(I) chain OS=Mus musculus GN=Col1a1 PE=1 SV=4 |
| 121 | 1 | UniProt | n | P98063 | 831 | 131033 | 29 | 29 | 19 | 19 | 0.91 | 1 | 1.06 | 1.078 | 1.06 | 1.066 | 1.128 | 1.045 | 1.084 | Bone morphogenetic protein 1 OS=Mus musculus GN=Bmp1 PE=1 SV=2 |
| 556 | 1 | UniProt | n | P54923 | 120 | 44835 | 6 | 6 | 4 | 4 | 0.46 | 1 | 1.589 | 1.429 | 1.154 | 1.089 | 1.013 | 1.314 | 1.084 | [Protein ADP-ribosylarginine] hydrolase OS=Mus musculus GN=Adprh PE=1 SV=1 |
| 29 | 4 | UniProt | n | Q9Z2K1 | 818 | 57980 | 41 | 41 | 8 | 8 | 1.07 | 1 | 0.777 | 0.928 | 0.886 | 1.302 | 1.102 | 0.897 | 1.083 | Keratin, type I cytoskeletal 16 OS=Mus musculus GN=Krt16 PE=1 SV=3 |
| 29 | 6 | UniProt | n | B1AQ77 | 495 | 55685 | 36 | 36 | 9 | 9 | 1.48 | 1 | 0.781 | 0.916 | 0.912 | 1.286 | 1.079 | 0.894 | 1.082 | Keratin 15, isoform CRA_a OS=Mus musculus GN=Krt15 PE=1 SV=1 |
| 616 | 1 | UniProt | n | Q91XA2 | 101 | 51434 | 6 | 6 | 2 | 2 | 0.18 | 1 | 1.033 | 1.111 | 1.026 | 1.126 | 1.091 | 1.047 | 1.080 | Golgi membrane protein 1 OS=Mus musculus GN=Golm1 PE=1 SV=2 |
| 1239 | 1 | UniProt | n | A0A0G2JC | 24 | 16435 | 1 | 1 | 1 | 1 | 0.29 | 1 | 0.955 | 0.925 | 1.058 | 1.168 | 1.017 | 0.960 | 1.079 | Cold shock domain-containing protein E1 (Fragment) OS=Mus musculus GN=Csde1 PE=1 SV=1 |
| 755 | 1 | UniProt | n | G3X9I1 | 72 | 27483 | 5 | 5 | 3 | 3 | 0.84 | 1 | 1.03 | 0.879 | 1.066 | 1.016 | 1.158 | 0.967 | 1.078 | DNA segment, Chr 6, Wayne State University 176, expressed, isoform CRA_a OS=Mus musculus GN=Fam3c PE=1 SV=1 |
| 131 | 1 | UniProt | n | P06797 | 770 | 45284 | 24 | 24 | 7 | 7 | 0.92 | 1 | 1.037 | 1.082 | 1.07 | 1.092 | 1.073 | 1.039 | 1.078 | Cathepsin L1 OS=Mus musculus GN=Ctsl PE=1 SV=2 |
| 113 | 1 | UniProt | n | A0A0A0M | 893 | 127866 | 45 | 45 | 18 | 18 | 0.94 | 1 | 0.961 | 1.008 | 1.114 | 1.032 | 1.088 | 0.989 | 1.077 | Matrilin-2 OS=Mus musculus GN=Matn2 PE=1 SV=1 |
| 326 | 3 | UniProt | n | Q99L20 | 57 | 32833 | 2 | 2 | 2 | 2 | 0.29 | 1 | 0.885 | 0.893 | 1.076 | 1.03 | 1.128 | 0.925 | 1.077 | Glutathione S-transferase theta-3 OS=Mus musculus GN=Gstt3 PE=1 SV=1 |
| 98 | 2 | UniProt | n | P01900 | 610 | 45619 | 22 | 22 | 6 | 6 | 1.3 | 1 | 0.966 | 0.948 | 1.134 | 1.042 | 1.057 | 0.971 | 1.077 | H-2 class I histocompatibility antigen, D-D alpha chain OS=Mus musculus GN=H2-D1 PE=1 SV=1 |
| 44 | 1 | UniProt | n | Q07797 | 2186 | 69749 | 67 | 67 | 16 | 16 | 3.29 | 1 | 0.998 | 0.984 | 1.102 | 1.022 | 1.095 | 0.994 | 1.072 | Galectin-3-binding protein OS=Mus musculus GN=Lgals3bp PE=1 SV=1 |
| 676 | 1 | UniProt | n | A0A1W2P | 87 | 21918 | 2 | 2 | 1 | 1 | 0.21 | 1 | 0.996 | 1.086 | 1.029 | 1.085 | 1.103 | 1.027 | 1.072 | Tetraspanin (Fragment) OS=Mus musculus GN=Cd63 PE=1 SV=1 |
| 1077 | 1 | UniProt | n | A0A1D5RI | 38 | 23899 | 2 | 2 | 1 | 1 | 0.19 | 1 | 1.022 | 0.882 | 1.188 | 1.011 | 1.02 | 0.966 | 1.070 | Translin-associated protein X OS=Mus musculus GN=Tsnax PE=1 SV=1 |
| 19 | 15 | UniProt | n | P11679 | 222 | 64874 | 36 | 36 | 8 | 8 | 0.69 | 1 | 0.781 | 1.01 | 1.061 | 1.255 | 0.918 | 0.924 | 1.069 | Keratin, type II cytoskeletal 8 OS=Mus musculus GN=Krt8 PE=1 SV=4 |
| 135 | 1 | UniProt | n | A0A0R4J0 | 720 | 70049 | 28 | 28 | 11 | 11 | 1.33 | 1 | 0.999 | 0.966 | 1.04 | 1.06 | 1.107 | 0.988 | 1.069 | Lysyl oxidase homolog 1 OS=Mus musculus GN=Loxl1 PE=1 SV=1 |
| 161 | 1 | UniProt | n | Q8R422 | 602 | 186842 | 24 | 24 | 14 | 14 | 0.41 | 1 | 0.937 | 0.96 | 1.082 | 1.075 | 1.048 | 0.965 | 1.068 | CD109 antigen OS=Mus musculus GN=Cd109 PE=1 SV=1 |
| 141 | 1 | UniProt | n | P15116 | 707 | 111358 | 31 | 31 | 15 | 15 | 0.99 | 1 | 0.96 | 0.974 | 1.101 | 1.07 | 1.034 | 0.978 | 1.068 | Cadherin-2 OS=Mus musculus GN=Cdh2 PE=1 SV=2 |
| 1040 | 1 | UniProt | n | P29758 | 41 | 56859 | 2 | 2 | 2 | 2 | 0.16 | 1 | 0.992 | 1.02 | 0.932 | 1.352 | 0.966 | 1.004 | 1.068 | Ornithine aminotransferase, mitochondrial OS=Mus musculus GN=Oat PE=1 SV=1 |
| 873 | 1 | UniProt | n | P01029 | 57 | 220593 | 2 | 2 | 2 | 2 | 0.04 | 1 | 1.052 | 0.988 | 1.019 | 1.042 | 1.146 | 1.013 | 1.068 | Complement C4-B OS=Mus musculus GN=C4b PE=1 SV=3 |
| 622 | 1 | UniProt | n | Q9ET22 | 99 | 61240 | 5 | 5 | 3 | 3 | 0.23 | 1 | 0.957 | 0.963 | 1.074 | 1.115 | 1.013 | 0.973 | 1.067 | Dipeptidyl peptidase 2 OS=Mus musculus GN=Dpp7 PE=1 SV=2 |
| 116 | 1 | UniProt | n | P18760 | 856 | 26641 | 32 | 32 | 8 | 8 | 2.52 | 1 | 1.071 | 1.041 | 1.105 | 1.094 | 1.003 | 1.037 | 1.066 | Cofilin-1 OS=Mus musculus GN=Cfl1 PE=1 SV=3 |
| 675 | 1 | UniProt | n | Q9WUM3 | 87 | 61381 | 1 | 1 | 1 | 1 | 0.07 | 1 | 1.146 | 1.189 | 1.1 | 1.086 | 1.008 | 1.109 | 1.064 | Coronin-1B OS=Mus musculus GN=Coro1b PE=1 SV=1 |
| 1038 | 1 | UniProt | n | J3QNB3 | 41 | 29399 | 1 | 1 | 1 | 1 | 0.15 | 1 | 0.897 | 0.84 | 1.114 | 1.225 | 0.882 | 0.910 | 1.064 | Disintegrin and metalloproteinase domain-containing protein 17 OS=Mus musculus GN=Adam17 PE=1 SV=1 |
| 240 | 1 | UniProt | n | Q9R0E1 | 364 | 93542 | 15 | 15 | 9 | 9 | 0.65 | 1 | 1.105 | 0.994 | 1.075 | 1.062 | 1.054 | 1.032 | 1.064 | Procollagen-lysine,2-oxoglutarate 5-dioxygenase 3 OS=Mus musculus GN=Plod3 PE=1 SV=1 |
| 1158 | 1 | UniProt | n | O88632-2 | 31 | 94922 | 2 | 2 | 2 | 2 | 0.09 | 1 | 0.977 | 1.311 | 1.123 | 1.096 | 0.977 | 1.086 | 1.063 | Isoform A of Semaphorin-3F OS=Mus musculus GN=Sema3f |
| 976 | 1 | UniProt | n | P56480 | 47 | 63566 | 2 | 2 | 2 | 2 | 0.14 | 1 | 0.866 | 1.061 | 1.25 | 1.116 | 0.86 | 0.972 | 1.063 | ATP synthase subunit beta, mitochondrial OS=Mus musculus GN=Atp5b PE=1 SV=2 |
| 664 | 1 | UniProt | n | P04925 | 89 | 31748 | 3 | 3 | 1 | 1 | 0.3 | 1 | 1.156 | 1.084 | 0.942 | 1.207 | 1.054 | 1.078 | 1.062 | Major prion protein OS=Mus musculus GN=Prnp PE=1 SV=2 |
| 1062 | 1 | UniProt | n | Q3TSR0 | 39 | 41656 | 2 | 2 | 2 | 2 | 0.22 | 1 | 0.968 | 0.976 | 1.173 | 1.099 | 0.929 | 0.981 | 1.062 | Exostosin-like 2 OS=Mus musculus GN=Extl2 PE=1 SV=1 |
| 1016 | 1 | UniProt | n | A0A0R4J0 | 43 | 36128 | 1 | 1 | 1 | 1 | 0.12 | 1 | 1.212 | 1.386 | 0.807 | 1.514 | 0.979 | 1.189 | 1.062 | Hydroxyacylglutathione hydrolase, mitochondrial OS=Mus musculus GN=Hagh PE=1 SV=1 |
| 953 | 1 | UniProt | n | A2AQV2 | 49 | 43315 | 3 | 3 | 3 | 3 | 0.34 | 1 | 0.906 | 0.952 | 1.093 | 1.153 | 0.942 | 0.952 | 1.059 | Carbohydrate sulfotransferase OS=Mus musculus GN=Chst14 PE=1 SV=1 |
| 805 | 1 | UniProt | n | E0CYT6 | 64 | 23638 | 2 | 2 | 1 | 1 | 0.19 | 1 | 0.899 | 1.038 | 1.087 | 1.046 | 1.04 | 0.977 | 1.057 | Syndecan-1 (Fragment) OS=Mus musculus GN=Sdc1 PE=1 SV=1 |
| 505 | 1 | UniProt | n | Q00493 | 141 | 60541 | 8 | 8 | 6 | 6 | 0.63 | 1 | 1.008 | 1.021 | 1.126 | 1.044 | 1.002 | 1.010 | 1.056 | Carboxypeptidase E OS=Mus musculus GN=Cpe PE=1 SV=2 |
| 1169 | 1 | UniProt | n | Q64514 | 30 | 167194 | 1 | 1 | 1 | 1 | 0.03 | 1 | 0.961 | 1.152 | 1.43 | 0.899 | 0.915 | 1.034 | 1.056 | Tripeptidyl-peptidase 2 OS=Mus musculus GN=Tpp2 PE=1 SV=3 |
| 78 | 1 | UniProt | n | Q64299 | 1242 | 46825 | 59 | 59 | 11 | 11 | 2.23 | 1 | 1.001 | 0.895 | 1.065 | 1.024 | 1.078 | 0.964 | 1.055 | Protein NOV homolog OS=Mus musculus GN=Nov PE=1 SV=1 |
| 297 | 1 | UniProt | n | O35375 | 293 | 119358 | 16 | 16 | 11 | 11 | 0.53 | 1 | 0.94 | 0.974 | 1.04 | 1.107 | 1.015 | 0.971 | 1.053 | Neuropilin-2 OS=Mus musculus GN=Nrp2 PE=1 SV=2 |
| 1261 | 1 | UniProt | n | Q8BZX4-2 | 21 | 91166 | 1 | 1 | 1 | 1 | 0.05 | 1 | 1.191 | 1.058 | 1.03 | 1.114 | 1.016 | 1.080 | 1.052 | Isoform 2 of Splicing regulatory glutamine/lysine-rich protein 1 OS=Mus musculus GN=Srek1 |
| 116 | 3 | UniProt | n | Q9R0P5 | 238 | 25478 | 11 | 11 | 5 | 5 | 1.27 | 1 | 1.051 | 1.076 | 1.101 | 1.071 | 0.982 | 1.042 | 1.050 | Destrin OS=Mus musculus GN=Dstn PE=1 SV=3 |
| 29 | 1 | UniProt | n | A2A513 | 3280 | 64141 | 149 | 149 | 9 | 9 | 1.52 | 1 | 0.551 | 0.715 | 0.898 | 1.266 | 1.012 | 0.733 | 1.048 | Keratin, type I cytoskeletal 10 OS=Mus musculus GN=Krt10 PE=1 SV=1 |
| 969 | 1 | UniProt | n | Q3TMX0 | 47 | 38194 | 2 | 2 | 2 | 2 | 0.25 | 1 | 0.856 | 0.881 | 1.074 | 1.002 | 1.058 | 0.910 | 1.044 | MCG4375, isoform CRA_b OS=Mus musculus GN=Sdcbp PE=1 SV=1 |
| 933 | 1 | UniProt | n | Q3UZI5 | 51 | 25164 | 3 | 3 | 1 | 1 | 0.18 | 1 | 1.096 | 1.091 | 1.083 | 1.048 | 1.003 | 1.061 | 1.044 | Pleiotrophin OS=Mus musculus GN=Ptn PE=1 SV=1 |
| 260 | 1 | UniProt | n | Q8CG19 | 337 | 216519 | 14 | 14 | 10 | 10 | 0.22 | 1 | 0.967 | 0.921 | 1.012 | 1.052 | 1.064 | 0.962 | 1.042 | Latent-transforming growth factor beta-binding protein 1 OS=Mus musculus GN=Ltbp1 PE=1 SV=2 |
| 1210 | 1 | UniProt | n | G3X8R2 | 27 | 29293 | 1 | 1 | 1 | 1 | 0.15 | 1 | 0.997 | 0.948 | 0.941 | 0.888 | 1.335 | 0.981 | 1.037 | Steroid receptor RNA activator 1 (Fragment) OS=Mus musculus GN=Sra1 PE=1 SV=1 |
| 910 | 1 | UniProt | n | A0A1W2P | 54 | 96048 | 1 | 1 | 1 | 1 | 0.05 | 1 | 0.865 | 1.006 | 1.043 | 0.972 | 1.095 | 0.955 | 1.035 | alpha-1,2-Mannosidase (Fragment) OS=Mus musculus GN=Man1a PE=1 SV=1 |
| 98 | 4 | UniProt | n | P01897 | 597 | 44916 | 25 | 25 | 9 | 9 | 1.81 | 1 | 0.884 | 0.908 | 1.096 | 1.037 | 0.976 | 0.929 | 1.035 | H-2 class I histocompatibility antigen, L-D alpha chain OS=Mus musculus GN=H2-L PE=1 SV=2 |
| 449 | 1 | UniProt | n | A0A140LH | 174 | 107212 | 5 | 5 | 2 | 2 | 0.13 | 1 | 1.054 | 1.003 | 1.035 | 0.976 | 1.093 | 1.019 | 1.034 | Disintegrin and metalloproteinase domain-containing protein 9 OS=Mus musculus GN=Adam9 PE=1 SV=1 |
| 881 | 1 | UniProt | n | O09159 | 57 | 127812 | 5 | 5 | 3 | 3 | 0.1 | 1 | 0.975 | 1.062 | 1.019 | 0.999 | 1.083 | 1.012 | 1.033 | Lysosomal alpha-mannosidase OS=Mus musculus GN=Man2b1 PE=1 SV=4 |
| 343 | 1 | UniProt | n | A0A0R4J0 | 247 | 43232 | 6 | 6 | 4 | 4 | 0.48 | 1 | 1.066 | 1.029 | 1.007 | 1.033 | 1.058 | 1.031 | 1.032 | Growth arrest-specific protein 1 OS=Mus musculus GN=Gas1 PE=1 SV=1 |
| 603 | 1 | UniProt | n | Q8R5L1 | 103 | 36757 | 2 | 2 | 1 | 1 | 0.12 | 1 | 0.936 | 0.986 | 1.043 | 0.853 | 1.235 | 0.974 | 1.032 | Complement component 1 Q subcomponent-binding protein, mitochondrial OS=Mus musculus GN=C1qbp PE=1 SV=1 |
| 765 | 1 | UniProt | n | Q64674 | 71 | 39300 | 4 | 4 | 2 | 2 | 0.24 | 1 | 1.041 | 1.176 | 0.936 | 1.055 | 1.11 | 1.070 | 1.031 | Spermidine synthase OS=Mus musculus GN=Srm PE=1 SV=1 |
| 736 | 1 | UniProt | n | Q99LL3 | 76 | 56594 | 3 | 3 | 2 | 2 | 0.16 | 1 | 0.875 | 0.952 | 1.116 | 1.027 | 0.953 | 0.941 | 1.030 | Carbohydrate sulfotransferase 12 OS=Mus musculus GN=Chst12 PE=2 SV=2 |
| 196 | 1 | UniProt | n | Q9JJH1 | 455 | 19207 | 9 | 9 | 4 | 4 | 1.38 | 1 | 0.984 | 0.947 | 1.032 | 1.026 | 1.029 | 0.977 | 1.029 | Ribonuclease 4 OS=Mus musculus GN=Rnase4 PE=1 SV=1 |
| 725 | 1 | UniProt | n | E0CX39 | 77 | 15030 | 1 | 1 | 1 | 1 | 0.32 | 1 | 0.911 | 1.12 | 0.985 | 1.125 | 0.982 | 1.007 | 1.029 | Gamma-glutamylaminecyclotransferase (Fragment) OS=Mus musculus GN=Ggact PE=1 SV=8 |
| 1192 | 1 | UniProt | n | E9Q2S9 | 29 | 32671 | 1 | 1 | 1 | 1 | 0.14 | 1 | 0.865 | 0.907 | 1.079 | 1.033 | 0.976 | 0.922 | 1.028 | Coiled-coil domain-containing protein 43 OS=Mus musculus GN=Ccdc43 PE=1 SV=1 |
| 1190 | 1 | UniProt | n | Q9Z0M5 | 29 | 52827 | 1 | 1 | 1 | 1 | 0.08 | 1 | 1.038 | 0.94 | 1.044 | 1.044 | 0.997 | 0.992 | 1.028 | Lysosomal acid lipase/cholesteryl ester hydrolase OS=Mus musculus GN=Lipa PE=1 SV=2 |
| 149 | 2 | UniProt | n | E9PWE8 | 255 | 86374 | 11 | 11 | 7 | 7 | 0.41 | 1 | 0.917 | 0.953 | 1.02 | 0.981 | 1.084 | 0.956 | 1.027 | Dihydropyrimidinase-related protein 3 OS=Mus musculus GN=Dpysl3 PE=1 SV=1 |
| 369 | 1 | UniProt | n | P27046 | 229 | 152527 | 12 | 12 | 8 | 8 | 0.25 | 1 | 0.936 | 0.996 | 1.025 | 1.004 | 1.048 | 0.977 | 1.026 | Alpha-mannosidase 2 OS=Mus musculus GN=Man2a1 PE=1 SV=2 |
| 1113 | 1 | UniProt | n | Q8CCH2 | 35 | 43010 | 1 | 1 | 1 | 1 | 0.1 | 1 | 0.971 | 0.917 | 1.009 | 1.059 | 1.005 | 0.962 | 1.024 | NHL repeat-containing protein 3 OS=Mus musculus GN=Nhlrc3 PE=1 SV=1 |
| 1048 | 1 | UniProt | n | D3Z4J1 | 41 | 23794 | 1 | 1 | 1 | 1 | 0.19 | 1 | 1.188 | 1.086 | 1.006 | 0.869 | 1.219 | 1.089 | 1.021 | DAZ-associated protein 1 (Fragment) OS=Mus musculus GN=Dazap1 PE=1 SV=1 |
| 1223 | 1 | UniProt | n | Q3UNS8 | 26 | 58059 | 1 | 1 | 1 | 1 | 0.08 | 1 | 1.101 | 0.978 | 0.973 | 1.039 | 1.054 | 1.025 | 1.021 | Kynurenine/alpha-aminoadipate aminotransferase, mitochondrial OS=Mus musculus GN=Aadat PE=1 SV=1 |
| 96 | 1 | UniProt | n | P05213 | 1068 | 56756 | 36 | 36 | 13 | 13 | 2.29 | 1 | 1.021 | 1.097 | 0.965 | 1.037 | 1.055 | 1.039 | 1.018 | Tubulin alpha-1B chain OS=Mus musculus GN=Tuba1b PE=1 SV=2 |
| 55 | 2 | UniProt | n | Q08879-2 | 1467 | 84020 | 65 | 65 | 16 | 16 | 1.36 | 1 | 1.053 | 1.049 | 1.05 | 0.973 | 1.03 | 1.034 | 1.017 | Isoform C of Fibulin-1 OS=Mus musculus GN=Fbln1 |
| 344 | 1 | UniProt | n | Q04999 | 247 | 50389 | 7 | 7 | 5 | 5 | 0.52 | 1 | 0.896 | 0.978 | 1.105 | 0.952 | 0.999 | 0.957 | 1.017 | Inhibin beta B chain OS=Mus musculus GN=Inhbb PE=2 SV=4 |
| 824 | 1 | UniProt | n | D3YXK2 | 62 | 123706 | 1 | 1 | 1 | 1 | 0.03 | 1 | 0.945 | 0.871 | 1.078 | 1.094 | 0.889 | 0.937 | 1.016 | Scaffold attachment factor B1 OS=Mus musculus GN=Safb PE=1 SV=2 |
| 278 | 1 | UniProt | n | A0A0R4J0 | 320 | 572640 | 18 | 18 | 12 | 12 | 0.1 | 1 | 0.972 | 1.03 | 1.029 | 1.001 | 1.015 | 1.000 | 1.015 | Low density lipoprotein receptor-related protein 1 OS=Mus musculus GN=Lrp1 PE=1 SV=1 |
| 1011 | 1 | UniProt | n | P35235 | 44 | 82567 | 4 | 4 | 4 | 4 | 0.23 | 1 | 0.966 | 0.964 | 1.044 | 0.914 | 1.094 | 0.977 | 1.014 | Tyrosine-protein phosphatase non-receptor type 11 OS=Mus musculus GN=Ptpn11 PE=1 SV=2 |
| 454 | 1 | UniProt | n | Q920A5 | 170 | 61505 | 6 | 6 | 5 | 5 | 0.41 | 1 | 1.091 | 1.136 | 0.949 | 1.072 | 1.026 | 1.074 | 1.014 | Retinoid-inducible serine carboxypeptidase OS=Mus musculus GN=Scpep1 PE=1 SV=2 |
| 134 | 1 | UniProt | n | P30412 | 740 | 27435 | 28 | 28 | 6 | 6 | 2.4 | 1 | 0.966 | 0.91 | 1.038 | 0.98 | 1.026 | 0.958 | 1.014 | Peptidyl-prolyl cis-trans isomerase C OS=Mus musculus GN=Ppic PE=1 SV=1 |
| 526 | 1 | UniProt | n | P97314 | 131 | 27732 | 4 | 4 | 3 | 3 | 0.57 | 1 | 0.879 | 0.935 | 1.057 | 1.015 | 0.971 | 0.937 | 1.014 | Cysteine and glycine-rich protein 2 OS=Mus musculus GN=Csrp2 PE=1 SV=3 |
| 727 | 1 | UniProt | n | Q61599 | 77 | 29879 | 1 | 1 | 1 | 1 | 0.15 | 1 | 0.907 | 1.033 | 0.991 | 0.988 | 1.063 | 0.979 | 1.013 | Rho GDP-dissociation inhibitor 2 OS=Mus musculus GN=Arhgdib PE=1 SV=3 |
| 620 | 2 | UniProt | n | A0A0G2JC | 93 | 38510 | 5 | 5 | 4 | 4 | 0.55 | 1 | 1.081 | 1.189 | 1.037 | 1.004 | 0.996 | 1.087 | 1.012 | Calponin OS=Mus musculus GN=Cnn3 PE=1 SV=1 |
| 828 | 1 | UniProt | n | Q9D0B6 | 62 | 26561 | 1 | 1 | 1 | 1 | 0.17 | 1 | 0.866 | 1.051 | 0.991 | 1.062 | 0.982 | 0.969 | 1.011 | Protein PBDC1 OS=Mus musculus GN=Pbdc1 PE=1 SV=1 |
| 1170 | 1 | UniProt | n | M0QWF0 | 30 | 38248 | 1 | 1 | 1 | 1 | 0.12 | 1 | 0.78 | 0.948 | 1.047 | 0.943 | 1.046 | 0.904 | 1.011 | Zinc finger protein 207 (Fragment) OS=Mus musculus GN=Zfp207 PE=1 SV=1 |
| 362 | 1 | UniProt | n | G5E924 | 233 | 76873 | 7 | 7 | 4 | 4 | 0.32 | 1 | 0.904 | 1.094 | 1.027 | 0.949 | 1.056 | 0.996 | 1.010 | Heterogeneous nuclear ribonucleoprotein L (Fragment) OS=Mus musculus GN=Hnrnpl PE=1 SV=1 |
| 116 | 2 | UniProt | n | P45591 | 408 | 26091 | 19 | 19 | 5 | 5 | 1.23 | 1 | 1.091 | 1.037 | 1.042 | 1.065 | 0.927 | 1.042 | 1.009 | Cofilin-2 OS=Mus musculus GN=Cfl2 PE=1 SV=1 |
| 358 | 1 | UniProt | n | E9Q668 | 237 | 90577 | 9 | 9 | 5 | 5 | 0.26 | 1 | 0.908 | 0.986 | 1.035 | 1.024 | 0.968 | 0.964 | 1.009 | Semaphorin-3A (Fragment) OS=Mus musculus GN=Sema3a PE=1 SV=1 |
| 208 | 3 | UniProt | n | O35639 | 106 | 44713 | 4 | 4 | 3 | 3 | 0.33 | 1 | 0.962 | 1.019 | 1.062 | 1.007 | 0.958 | 0.993 | 1.008 | Annexin A3 OS=Mus musculus GN=Anxa3 PE=1 SV=4 |
| 878 | 1 | UniProt | n | E0CXA9 | 57 | 27866 | 2 | 2 | 1 | 1 | 0.16 | 1 | 0.693 | 0.868 | 1.068 | 1.175 | 0.816 | 0.844 | 1.008 | MOB-like protein phocein OS=Mus musculus GN=Mob4 PE=1 SV=1 |
| 27 | 2 | UniProt | n | A0A1B0GI | 1762 | 38093 | 84 | 84 | 19 | 19 | 9.21 | 1 | 0.96 | 0.955 | 1.052 | 1.001 | 0.971 | 0.971 | 1.007 | Nucleobindin-1 (Fragment) OS=Mus musculus GN=Nucb1 PE=1 SV=1 |
| 572 | 1 | UniProt | n | Q8K3C3 | 115 | 27653 | 2 | 2 | 2 | 2 | 0.35 | 1 | 1.007 | 0.955 | 1.015 | 1.114 | 0.901 | 0.987 | 1.006 | Protein LZIC OS=Mus musculus GN=Lzic PE=1 SV=1 |
| 1222 | 1 | UniProt | n | Q8K1K2 | 26 | 46166 | 1 | 1 | 1 | 1 | 0.1 | 1 | 0.642 | 0.788 | 0.999 | 1.512 | 0.673 | 0.797 | 1.005 | 26S proteasome regulatory subunit 8 OS=Mus musculus GN=Psmc5 PE=1 SV=1 |
| 25 | 1 | UniProt | n | P01027 | 3863 | 224112 | 167 | 167 | 56 | 56 | 2.24 | 1 | 1.002 | 1.031 | 1.002 | 0.979 | 1.032 | 1.011 | 1.004 | Complement C3 OS=Mus musculus GN=C3 PE=1 SV=3 |
| 880 | 1 | UniProt | n | Q8BM72 | 57 | 60241 | 2 | 2 | 1 | 1 | 0.07 | 1 | 1.127 | 1.227 | 1.078 | 0.994 | 0.942 | 1.114 | 1.003 | Heat shock 70 kDa protein 13 OS=Mus musculus GN=Hspa13 PE=1 SV=1 |
| 1257 | 1 | UniProt | n | P60122 | 21 | 63235 | 1 | 1 | 1 | 1 | 0.07 | 1 | 0.897 | 1.164 | 1.042 | 0.98 | 0.988 | 1.014 | 1.003 | RuvB-like 1 OS=Mus musculus GN=Ruvbl1 PE=1 SV=1 |
| 27 | 1 | UniProt | n | Q02819 | 3421 | 62502 | 145 | 145 | 27 | 27 | 7.15 | 1 | 0.943 | 0.931 | 1.041 | 0.993 | 0.975 | 0.958 | 1.003 | Nucleobindin-1 OS=Mus musculus GN=Nucb1 PE=1 SV=2 |
| 850 | 1 | UniProt | n | H3BJF6 | 60 | 11387 | 8 | 8 | 1 | 1 | 1.05 | 1 | 1.108 | 1.141 | 0.949 | 0.948 | 1.117 | 1.081 | 1.002 | Protein zwilch homolog (Fragment) OS=Mus musculus GN=Zwilch PE=1 SV=1 |
| 55 | 1 | UniProt | n | Q08879 | 1632 | 86767 | 69 | 69 | 19 | 19 | 1.65 | 1 | 1.04 | 1.047 | 1.03 | 0.961 | 1.014 | 1.029 | 1.001 | Fibulin-1 OS=Mus musculus GN=Fbln1 PE=1 SV=2 |
| 1022 | 1 | UniProt | n | O35188 | 43 | 47611 | 2 | 2 | 2 | 2 | 0.19 | 1 | 0.927 | 0.912 | 1.036 | 1.021 | 0.948 | 0.946 | 1.001 | Fractalkine OS=Mus musculus GN=Cx3cl1 PE=1 SV=3 |
| 964 | 1 | UniProt | n | Q9Z0L8 | 48 | 42932 | 1 | 1 | 1 | 1 | 0.1 | 1 | 0.865 | 1.076 | 1.172 | 0.956 | 0.894 | 0.976 | 1.001 | Gamma-glutamyl hydrolase OS=Mus musculus GN=Ggh PE=1 SV=2 |
| 324 | 1 | UniProt | n | Q8C9W3 | 266 | 153348 | 5 | 5 | 4 | 4 | 0.12 | 1 | 0.905 | 0.919 | 0.991 | 1.024 | 0.985 | 0.940 | 1.000 | A disintegrin and metalloproteinase with thrombospondin motifs 2 OS=Mus musculus GN=Adamts2 PE=1 SV=2 |
| 1179 | 1 | UniProt | n | Q8K1L5 | 30 | 17807 | 1 | 1 | 1 | 1 | 0.26 | 1 | 0.765 | 0.756 | 0.98 | 0.984 | 1.036 | 0.833 | 1.000 | Protein phosphatase 1 regulatory subunit 11 OS=Mus musculus GN=Ppp1r11 PE=1 SV=1 |
| 156 | 1 | UniProt | n | M0QWP1 | 639 | 239161 | 27 | 27 | 17 | 17 | 0.38 | 1 | 0.918 | 0.944 | 1.102 | 0.957 | 0.947 | 0.953 | 1.000 | Agrin OS=Mus musculus GN=Agrn PE=1 SV=1 |
| 1092 | 1 | UniProt | n | P09671 | 36 | 29943 | 1 | 1 | 1 | 1 | 0.15 | 1 | 1.022 | 1.014 | 0.943 | 1.056 | 0.999 | 1.012 | 0.998 | Superoxide dismutase [Mn], mitochondrial OS=Mus musculus GN=Sod2 PE=1 SV=3 |
| 863 | 1 | UniProt | n | Q9D0R8 | 58 | 27393 | 1 | 1 | 1 | 1 | 0.17 | 1 | 0.975 | 1.075 | 0.975 | 1.062 | 0.959 | 1.016 | 0.998 | Protein LSM12 homolog OS=Mus musculus GN=Lsm12 PE=1 SV=1 |
| 1114 | 1 | UniProt | n | F6TVP2 | 35 | 89625 | 1 | 1 | 1 | 1 | 0.05 | 1 | 1.014 | 1.193 | 1.067 | 0.978 | 0.951 | 1.066 | 0.997 | Deoxynucleoside triphosphate triphosphohydrolase SAMHD1 (Fragment) OS=Mus musculus GN=Samhd1 PE=1 SV=1 |
| 870 | 1 | UniProt | n | E9Q7G0 | 57 | 281688 | 2 | 2 | 2 | 2 | 0.03 | 1 | 0.917 | 1.06 | 1.037 | 1.07 | 0.893 | 0.991 | 0.997 | Nuclear mitotic apparatus protein 1 OS=Mus musculus GN=Numa1 PE=1 SV=1 |
| 900 | 1 | UniProt | n | Z4YJD9 | 55 | 59803 | 2 | 2 | 2 | 2 | 0.15 | 1 | 0.587 | 0.627 | 0.924 | 1.135 | 0.939 | 0.717 | 0.995 | Keratin, type I cytoskeletal 12 OS=Mus musculus GN=Krt12 PE=1 SV=1 |
| 110 | 1 | UniProt | n | Q61553 | 944 | 63590 | 31 | 31 | 15 | 15 | 1.71 | 1 | 1.002 | 1.02 | 1.037 | 0.943 | 1.005 | 1.007 | 0.994 | Fascin OS=Mus musculus GN=Fscn1 PE=1 SV=4 |
| 662 | 1 | UniProt | n | D3Z598 | 89 | 176298 | 3 | 3 | 3 | 3 | 0.07 | 1 | 0.952 | 1.037 | 0.887 | 1.157 | 0.955 | 0.996 | 0.993 | Latent-transforming growth factor beta-binding protein 4 OS=Mus musculus GN=Ltbp4 PE=1 SV=1 |
| 69 | 1 | UniProt | n | A0A0A6Y\ | 1458 | 112026 | 52 | 52 | 19 | 19 | 1.3 | 1 | 1.064 | 1.095 | 1.023 | 0.987 | 0.969 | 1.052 | 0.993 | Neural cell adhesion molecule 1 OS=Mus musculus GN=Ncam1 PE=1 SV=1 |
| 71 | 2 | UniProt | n | Q9EPL2 | 1386 | 125053 | 61 | 61 | 18 | 18 | 1.11 | 1 | 0.996 | 1.004 | 1.026 | 0.978 | 0.974 | 1.000 | 0.992 | Calsyntenin-1 OS=Mus musculus GN=Clstn1 PE=1 SV=1 |

| | | | | | | | | | | | | | | | | | | | |
|---|---|---|---|---|---|---|---|---|---|---|---|---|---|---|---|---|---|---|---|
| 1153 | 1 | UniProt n | Q9DBZ5 | 31 | 29255 | 2 | 2 | 2 | 2 | 0.33 | 1 | 0.982 | 1.035 | 1.071 | 1.042 | 0.872 | 1.005 | 0.991 | Eukaryotic translation initiation factor 3 subunit K OS=Mus musculus GN=Eif3k PE=1 SV=1 |
| 2 | 1 | UniProt n | Q01149 | 30114 | 144798 | 712 | 712 | 65 | 65 | 15.71 | 1 | 1.046 | 1.021 | 0.964 | 0.972 | 1.037 | 1.022 | 0.990 | Collagen alpha-2(I) chain OS=Mus musculus GN=Col1a2 PE=1 SV=2 |
| 1107 | 1 | UniProt n | Q9CRB6 | 35 | 27609 | 1 | 1 | 1 | 1 | 0.16 | 1 | 0.931 | 1.111 | 0.962 | 1.019 | 0.991 | 1.011 | 0.990 | Tubulin polymerization-promoting protein family member 3 OS=Mus musculus GN=Tppp3 PE=1 SV=1 |
| 902 | 1 | UniProt n | Q3UPI1 | 55 | 66769 | 2 | 2 | 1 | 1 | 0.14 | 1 | 1.04 | 0.996 | 0.956 | 1.005 | 1.011 | 1.012 | 0.990 | Protein FAM198B OS=Mus musculus GN=Fam198b PE=2 SV=1 |
| 501 | 1 | UniProt n | B1AWE0 | 145 | 27161 | 3 | 3 | 2 | 2 | 0.36 | 1 | 0.976 | 1.06 | 1.119 | 0.963 | 0.901 | 1.011 | 0.990 | Clathrin light chain OS=Mus musculus GN=Clta PE=1 SV=1 |
| 779 | 1 | UniProt n | Q8BVU5 | 69 | 46065 | 1 | 1 | 1 | 1 | 0.1 | 1 | 1.077 | 1.098 | 1.109 | 0.922 | 0.949 | 1.057 | 0.990 | ADP-ribose pyrophosphatase, mitochondrial OS=Mus musculus GN=Nudt9 PE=1 SV=1 |
| 528 | 1 | UniProt n | Q08024 | 130 | 24634 | 1 | 1 | 1 | 1 | 0.19 | 1 | 1.027 | 1.071 | 0.977 | 0.999 | 0.99 | 1.032 | 0.989 | Core-binding factor subunit beta OS=Mus musculus GN=Cbfb PE=1 SV=1 |
| 159 | 1 | UniProt n | Q64449 | 614 | 183795 | 19 | 19 | 10 | 10 | 0.26 | 1 | 0.964 | 0.99 | 1.036 | 0.968 | 0.963 | 0.985 | 0.988 | C-type mannose receptor 2 OS=Mus musculus GN=Mrc2 PE=1 SV=3 |
| 634 | 1 | UniProt n | Q8CC47 | 95 | 53806 | 2 | 2 | 1 | 1 | 0.08 | 1 | 1.085 | 1.057 | 0.93 | 0.959 | 1.081 | 1.047 | 0.988 | N-acetylgalactosamine-6-sulfatase OS=Mus musculus GN=Galns PE=1 SV=1 |
| 997 | 1 | UniProt n | Q99LJ1 | 45 | 60995 | 1 | 1 | 1 | 1 | 0.07 | 1 | 0.973 | 1.138 | 0.978 | 1.004 | 0.981 | 1.035 | 0.988 | Tissue alpha-L-fucosidase OS=Mus musculus GN=Fuca1 PE=1 SV=1 |
| 1191 | 1 | UniProt n | Q9WTZ2 | 29 | 133799 | 2 | 2 | 1 | 1 | 0.03 | 1 | 1.05 | 1.064 | 0.926 | 1.035 | 1.005 | 1.038 | 0.988 | Membrane-bound transcription factor site-1 protease OS=Mus musculus GN=Mbtps1 PE=1 SV=1 |
| 844 | 1 | UniProt n | Q9WUK2 | 60 | 31021 | 1 | 1 | 1 | 1 | 0.15 | 1 | 1.064 | 1.003 | 1.066 | 0.87 | 1.036 | 1.022 | 0.987 | Eukaryotic translation initiation factor 4H OS=Mus musculus GN=Eif4h PE=1 SV=3 |
| 1271 | 1 | UniProt n | Q99MQ4 | 19 | 53514 | 1 | 1 | 1 | 1 | 0.08 | 1 | 0.942 | 1.095 | 0.912 | 1.135 | 0.928 | 1.010 | 0.987 | Asporin OS=Mus musculus GN=Aspn PE=1 SV=1 |
| 1051 | 1 | UniProt n | Q9D8B3 | 40 | 33134 | 1 | 1 | 1 | 1 | 0.14 | 1 | 1.137 | 1.327 | 1.01 | 1.047 | 0.901 | 1.147 | 0.984 | Charged multivesicular body protein 4b OS=Mus musculus GN=Chmp4b PE=1 SV=2 |
| 567 | 1 | UniProt n | Q06335 | 117 | 92832 | 5 | 5 | 4 | 4 | 0.26 | 1 | 0.88 | 0.863 | 0.978 | 1.058 | 0.92 | 0.912 | 0.984 | Amyloid-like protein 2 OS=Mus musculus GN=Aplp2 PE=1 SV=4 |
| 287 | 1 | UniProt n | P61161 | 309 | 52872 | 10 | 10 | 8 | 8 | 0.9 | 1 | 0.912 | 0.956 | 0.938 | 0.959 | 1.058 | 0.955 | 0.984 | Actin-related protein 2 OS=Mus musculus GN=Actr2 PE=1 SV=1 |
| 1006 | 1 | UniProt n | Q9D819 | 44 | 42140 | 1 | 1 | 1 | 1 | 0.11 | 1 | 1.003 | 1.127 | 0.936 | 1.078 | 0.943 | 1.042 | 0.984 | Inorganic pyrophosphatase OS=Mus musculus GN=Ppa1 PE=1 SV=1 |
| 475 | 1 | UniProt n | Q3UW53 | 157 | 120006 | 4 | 4 | 3 | 3 | 0.11 | 1 | 0.928 | 1.077 | 1.214 | 1.162 | 0.674 | 1.000 | 0.983 | Protein Niban OS=Mus musculus GN=Fam129a PE=1 SV=2 |
| 446 | 1 | UniProt n | O55135 | 174 | 28733 | 3 | 3 | 2 | 2 | 0.55 | 1 | 0.934 | 1.08 | 0.978 | 0.968 | 1.004 | 1.003 | 0.983 | Eukaryotic translation initiation factor 6 OS=Mus musculus GN=Eif6 PE=1 SV=2 |
| 375 | 1 | UniProt n | Q8K482 | 226 | 135284 | 10 | 10 | 6 | 6 | 0.21 | 1 | 0.884 | 0.915 | 1.013 | 0.97 | 0.965 | 0.932 | 0.982 | EMILIN-2 OS=Mus musculus GN=Emilin2 PE=1 SV=1 |
| 87 | 1 | UniProt n | Q62165 | 1144 | 112331 | 47 | 47 | 15 | 15 | 1.3 | 1 | 1.027 | 1.001 | 0.991 | 0.954 | 0.995 | 1.009 | 0.980 | Dystroglycan OS=Mus musculus GN=Dag1 PE=1 SV=4 |
| 238 | 1 | UniProt n | G3X8T3 | 368 | 63193 | 13 | 13 | 7 | 7 | 0.71 | 1 | 0.973 | 1.008 | 0.977 | 0.97 | 0.992 | 0.994 | 0.980 | Carboxypeptidase OS=Mus musculus GN=Ctsa PE=1 SV=1 |
| 591 | 1 | UniProt n | Q8VDM4 | 107 | 116601 | 4 | 4 | 3 | 3 | 0.12 | 1 | 1.007 | 1.159 | 0.914 | 1.008 | 1.02 | 1.053 | 0.979 | 26S proteasome non-ATPase regulatory subunit 2 OS=Mus musculus GN=Psmd2 PE=1 SV=1 |
| 394 | 1 | UniProt n | J3QPW1 | 210 | 41615 | 7 | 7 | 5 | 5 | 0.66 | 1 | 1.087 | 1.034 | 0.959 | 1.002 | 0.976 | 1.040 | 0.979 | Phosphatidylinositol transfer protein alpha isoform OS=Mus musculus GN=Pitpna PE=1 SV=1 |
| 704 | 1 | UniProt n | E9Q440 | 81 | 125543 | 2 | 2 | 1 | 1 | 0.03 | 1 | 0.846 | 0.805 | 0.996 | 0.961 | 0.979 | 0.880 | 0.979 | Protein HEG homolog 1 OS=Mus musculus GN=Heg1 PE=1 SV=1 |
| 146 | 1 | UniProt n | P99024 | 689 | 54874 | 33 | 33 | 14 | 14 | 2.18 | 1 | 1.008 | 1.073 | 0.932 | 0.995 | 1.01 | 1.026 | 0.978 | Tubulin beta-5 chain OS=Mus musculus GN=Tubb5 PE=1 SV=1 |
| 512 | 1 | UniProt n | P47941 | 137 | 38464 | 4 | 4 | 2 | 2 | 0.39 | 1 | 0.921 | 1.012 | 1.04 | 0.995 | 0.905 | 0.977 | 0.978 | Crk-like protein OS=Mus musculus GN=Crkl PE=1 SV=2 |
| 71 | 1 | UniProt n | Q9EPL2-2 | 1396 | 123600 | 59 | 59 | 18 | 18 | 1.13 | 1 | 0.999 | 1 | 1.007 | 0.965 | 0.962 | 1.000 | 0.978 | Isoform 2 of Calsyntenin-1 OS=Mus musculus GN=Clstn1 |
| 1079 | 1 | UniProt n | E0CYQ2 | 38 | 16942 | 1 | 1 | 1 | 1 | 0.28 | 1 | 0.949 | 1.177 | 1.025 | 0.965 | 0.943 | 1.038 | 0.977 | NudC domain-containing protein 2 OS=Mus musculus GN=Nudcd2 PE=1 SV=1 |
| 370 | 1 | UniProt n | B0V2N1 | 229 | 236769 | 6 | 6 | 5 | 5 | 0.09 | 1 | 0.891 | 1.005 | 0.958 | 1.006 | 0.966 | 0.964 | 0.976 | Receptor-type tyrosine-protein phosphatase S OS=Mus musculus GN=Ptprs PE=1 SV=1 |
| 1027 | 1 | UniProt n | A2AH25 | 43 | 63971 | 1 | 1 | 1 | 1 | 0.07 | 1 | 1.156 | 1.195 | 0.914 | 1.047 | 0.972 | 1.114 | 0.976 | Rho GTPase-activating protein 1 OS=Mus musculus GN=Arhgap1 PE=1 SV=1 |
| 462 | 1 | UniProt n | Q9D1Q6 | 167 | 53838 | 6 | 6 | 5 | 5 | 0.48 | 1 | 0.855 | 0.914 | 1.003 | 1.02 | 0.908 | 0.921 | 0.976 | Endoplasmic reticulum resident protein 44 OS=Mus musculus GN=Erp44 PE=1 SV=1 |
| 1130 | 1 | UniProt n | O89017 | 33 | 57268 | 1 | 1 | 1 | 1 | 0.08 | 1 | 1.078 | 1.106 | 0.867 | 1.014 | 1.053 | 1.060 | 0.975 | Legumain OS=Mus musculus GN=Lgmn PE=1 SV=1 |
| 1091 | 1 | UniProt n | A0A0B4J1 | 37 | 77897 | 1 | 1 | 1 | 1 | 0.06 | 1 | 0.776 | 0.924 | 1.01 | 0.897 | 1.021 | 0.895 | 0.974 | SNW domain-containing protein 1 OS=Mus musculus GN=Snw1 PE=1 SV=1 |
| 518 | 1 | UniProt n | Q8QZR4 | 134 | 36269 | 7 | 7 | 6 | 6 | 1.01 | 1 | 1.072 | 0.923 | 1.005 | 0.994 | 0.925 | 0.996 | 0.974 | Out at first protein homolog OS=Mus musculus GN=Oaf PE=2 SV=1 |
| 72 | 1 | UniProt n | Q99K41 | 1387 | 113443 | 30 | 30 | 15 | 15 | 0.82 | 1 | 0.925 | 0.99 | 1.029 | 0.95 | 0.945 | 0.971 | 0.974 | EMILIN-1 OS=Mus musculus GN=Emilin1 PE=1 SV=1 |
| 180 | 1 | UniProt n | Q6P5H2 | 516 | 230321 | 65 | 65 | 4 | 4 | 0.08 | 1 | 0.723 | 1.023 | 0.955 | 1.182 | 0.818 | 0.904 | 0.974 | Nestin OS=Mus musculus GN=Nes PE=1 SV=1 |
| 637 | 1 | UniProt n | P10922 | 94 | 37884 | 3 | 3 | 3 | 3 | 0.4 | 1 | 1.061 | 1.08 | 0.919 | 0.978 | 1.023 | 1.046 | 0.972 | Histone H1.0 OS=Mus musculus GN=H1f0 PE=2 SV=4 |
| 42 | 2 | UniProt n | G3UXY9 | 1995 | 124033 | 76 | 76 | 26 | 26 | 1.99 | 1 | 1.033 | 0.981 | 0.976 | 0.958 | 0.982 | 1.004 | 0.972 | Ectonucleotide pyrophosphatase/phosphodiesterase family member 2 OS=Mus musculus GN=Enpp2 PE=1 SV=1 |
| 627 | 2 | UniProt n | B8JJN0 | 90 | 168411 | 10 | 10 | 5 | 5 | 0.19 | 1 | 1.029 | 1.015 | 0.999 | 0.992 | 0.925 | 1.015 | 0.971 | Predicted gene 20547 OS=Mus musculus GN=Gm20547 PE=3 SV=1 |
| 468 | 1 | UniProt n | Q9Z1Z2 | 161 | 46861 | 7 | 7 | 5 | 5 | 0.57 | 1 | 1.047 | 1.084 | 0.976 | 0.924 | 1.015 | 1.043 | 0.971 | Serine-threonine kinase receptor-associated protein OS=Mus musculus GN=Strap PE=1 SV=2 |
| 167 | 1 | UniProt n | Q8BKG3 | 580 | 132647 | 27 | 27 | 12 | 12 | 0.47 | 1 | 1.011 | 0.979 | 0.979 | 0.958 | 0.975 | 0.997 | 0.971 | Inactive tyrosine-protein kinase 7 OS=Mus musculus GN=Ptk7 PE=1 SV=1 |
| 1058 | 1 | UniProt n | Q6PGB6 | 40 | 24757 | 1 | 1 | 1 | 1 | 0.18 | 1 | 1.283 | 1.223 | 1.104 | 0.8 | 1.035 | 1.162 | 0.971 | N-alpha-acetyltransferase 50 OS=Mus musculus GN=Naa50 PE=1 SV=1 |
| 43 | 1 | UniProt n | Q8BND5-3 | 2191 | 70587 | 74 | 74 | 20 | 20 | 3.48 | 1 | 0.966 | 0.964 | 0.984 | 0.969 | 0.955 | 0.977 | 0.969 | Isoform 3 of Sulfhydryl oxidase 1 OS=Mus musculus GN=Qsox1 |
| 1231 | 1 | UniProt n | A0A0G2JF | 25 | 80164 | 1 | 1 | 1 | 1 | 0.05 | 1 | 0.829 | 1.049 | 0.947 | 1.101 | 0.872 | 0.955 | 0.969 | MCG8104 OS=Mus musculus GN=Tsc22d2 PE=1 SV=1 |
| 762 | 1 | UniProt n | P10287 | 71 | 102391 | 2 | 2 | 2 | 2 | 0.09 | 1 | 0.928 | 0.925 | 0.98 | 0.926 | 1.001 | 0.950 | 0.968 | Cadherin-3 OS=Mus musculus GN=Cdh3 PE=1 SV=2 |
| 860 | 1 | UniProt n | Q5SX75 | 59 | 70379 | 2 | 2 | 2 | 2 | 0.13 | 1 | 0.948 | 1.032 | 0.98 | 1.002 | 0.925 | 0.993 | 0.968 | Procollagen-proline, 2-oxoglutarate 4-dioxygenase (Proline 4-hydroxylase), alpha II polypeptide, isoform CRA_f OS=Mus musculus GN=P4ha2 PE=1 SV=1 |
| 724 | 1 | UniProt n | A0A1B0G | 77 | 119060 | 2 | 2 | 2 | 2 | 0.07 | 1 | 0.973 | 1.052 | 1.023 | 0.914 | 0.971 | 1.008 | 0.968 | Large proline-rich protein BAG6 (Fragment) OS=Mus musculus GN=Bag6 PE=1 SV=1 |
| 43 | 2 | UniProt n | Q8BND5 | 2189 | 91590 | 75 | 75 | 21 | 21 | 2.33 | 1 | 0.967 | 0.964 | 0.982 | 0.966 | 0.955 | 0.977 | 0.968 | Sulfhydryl oxidase 1 OS=Mus musculus GN=Qsox1 PE=1 SV=1 |
| 452 | 1 | UniProt n | Q3UPN1 | 171 | 97395 | 8 | 8 | 6 | 6 | 0.3 | 1 | 0.95 | 1.028 | 0.982 | 1.013 | 0.908 | 0.992 | 0.967 | Vascular cell adhesion protein 1 OS=Mus musculus GN=Vcam1 PE=1 SV=1 |
| 723 | 1 | UniProt n | P17047 | 77 | 53367 | 5 | 5 | 2 | 2 | 0.17 | 1 | 1.182 | 1.157 | 0.899 | 1.055 | 0.946 | 1.110 | 0.964 | Lysosome-associated membrane glycoprotein 2 OS=Mus musculus GN=Lamp2 PE=1 SV=2 |
| 65 | 1 | UniProt n | Q9R0E2 | 1484 | 95562 | 50 | 50 | 22 | 22 | 2.46 | 1 | 1.144 | 0.971 | 0.972 | 0.911 | 1.013 | 1.036 | 0.964 | Procollagen-lysine,2-oxoglutarate 5-dioxygenase 1 OS=Mus musculus GN=Plod1 PE=1 SV=1 |
| 119 | 2 | UniProt n | Q63ZW6 | 578 | 187707 | 16 | 16 | 5 | 5 | 0.12 | 1 | 0.976 | 0.932 | 0.987 | 0.926 | 0.981 | 0.969 | 0.964 | Col4a5 protein OS=Mus musculus GN=Col4a5 PE=1 SV=1 |
| 168 | 1 | UniProt n | Q8VHY0 | 577 | 264615 | 23 | 23 | 19 | 19 | 0.38 | 1 | 0.89 | 0.928 | 0.976 | 0.925 | 0.989 | 0.938 | 0.963 | Chondroitin sulfate proteoglycan 4 OS=Mus musculus GN=Cspg4 PE=1 SV=3 |
| 627 | 1 | UniProt n | B8JJM9 | 97 | 63001 | 10 | 10 | 5 | 5 | 0.6 | 1 | 1.047 | 1.028 | 0.982 | 0.97 | 0.936 | 1.025 | 0.962 | Complement C2 OS=Mus musculus GN=C2 PE=1 SV=1 |
| 923 | 1 | UniProt n | A0A1L1SR | 53 | 50783 | 3 | 3 | 2 | 2 | 0.18 | 1 | 0.92 | 0.95 | 0.981 | 1.009 | 0.899 | 0.956 | 0.962 | Neogenin (Fragment) OS=Mus musculus GN=Neo1 PE=1 SV=1 |
| 1098 | 1 | UniProt n | O88428 | 36 | 83284 | 2 | 2 | 1 | 1 | 0.05 | 1 | 1.025 | 1.05 | 1.042 | 0.94 | 0.907 | 1.025 | 0.961 | Bifunctional 3'-phosphoadenosine 5'-phosphosulfate synthase 2 OS=Mus musculus GN=Papss2 PE=1 SV=2 |
| 687 | 1 | UniProt n | Q91Z50 | 85 | 54432 | 1 | 1 | 1 | 1 | 0.08 | 1 | 1.11 | 1.084 | 0.93 | 1.021 | 0.931 | 1.064 | 0.960 | Flap endonuclease 1 OS=Mus musculus GN=Fen1 PE=1 SV=1 |
| 807 | 1 | UniProt n | Q9JKR6 | 64 | 135328 | 2 | 2 | 2 | 2 | 0.06 | 1 | 1.032 | 1.112 | 0.988 | 0.995 | 0.898 | 1.047 | 0.959 | Hypoxia up-regulated protein 1 OS=Mus musculus GN=Hyou1 PE=1 SV=1 |
| 309 | 1 | UniProt n | Q60675 | 286 | 403975 | 17 | 17 | 14 | 14 | 0.17 | 1 | 0.907 | 0.995 | 0.985 | 0.962 | 0.931 | 0.966 | 0.959 | Laminin subunit alpha-2 OS=Mus musculus GN=Lama2 PE=1 SV=2 |
| 774 | 1 | UniProt n | A2A9Q2 | 69 | 165604 | 5 | 5 | 5 | 5 | 0.14 | 1 | 0.96 | 1.106 | 1.09 | 0.957 | 0.844 | 1.020 | 0.958 | Nardilysin, N-arginine dibasic convertase, NRD convertase 1 OS=Mus musculus GN=Nrd1 PE=1 SV=1 |
| 382 | 1 | UniProt n | P70699 | 222 | 114079 | 7 | 7 | 4 | 4 | 0.2 | 1 | 0.865 | 0.893 | 1.029 | 1.01 | 0.846 | 0.918 | 0.958 | Lysosomal alpha-glucosidase OS=Mus musculus GN=Gaa PE=1 SV=2 |
| 128 | 1 | UniProt n | Q6PI17 | 792 | 30672 | 31 | 31 | 6 | 6 | 2.44 | 1 | 1.307 | 0.993 | 1.008 | 0.911 | 0.955 | 1.091 | 0.957 | Metalloproteinase inhibitor 2 OS=Mus musculus GN=Timp2 PE=1 SV=1 |
| 146 | 2 | UniProt n | P68372 | 628 | 55034 | 32 | 32 | 14 | 14 | 2.16 | 1 | 0.984 | 1.054 | 0.901 | 0.974 | 0.997 | 1.012 | 0.956 | Tubulin beta-4B chain OS=Mus musculus GN=Tubb4b PE=1 SV=1 |
| 1217 | 1 | UniProt n | A0A0N4S | 26 | 14091 | 1 | 1 | 1 | 1 | 0.34 | 1 | 1.036 | 1.01 | 0.813 | 0.988 | 1.089 | 1.015 | 0.956 | RAB7, member RAS oncogene family (Fragment) OS=Mus musculus GN=Rab7 PE=1 SV=3 |
| 29 | 8 | UniProt n | A6BLY7 | 246 | 58897 | 21 | 21 | 3 | 3 | 0.54 | 1 | 0.632 | 0.783 | 0.822 | 1.123 | 0.947 | 0.791 | 0.956 | Keratin, type I cytoskeletal 28 OS=Mus musculus GN=Krt28 PE=1 SV=1 |
| 1090 | 1 | UniProt n | Q62384 | 37 | 58094 | 1 | 1 | 1 | 1 | 0.08 | 1 | 0.892 | 1.063 | 0.865 | 1.099 | 0.918 | 0.982 | 0.956 | Zinc finger protein ZPR1 OS=Mus musculus GN=Zpr1 PE=1 SV=1 |
| 7 | 1 | UniProt n | Q04857 | 7177 | 126681 | 199 | 199 | 32 | 32 | 2.95 | 1 | 1.062 | 1.067 | 0.94 | 0.962 | 0.964 | 1.043 | 0.955 | Collagen alpha-1(VI) chain OS=Mus musculus GN=Col6a1 PE=1 SV=1 |
| 829 | 1 | UniProt n | B1AT92 | 62 | 28836 | 3 | 3 | 2 | 2 | 0.34 | 1 | 1.088 | 1.119 | 0.916 | 1.024 | 0.927 | 1.068 | 0.954 | Growth factor receptor-bound protein 2 OS=Mus musculus GN=Grb2 PE=1 SV=1 |
| 1145 | 1 | UniProt n | G3UYY1 | 32 | 57471 | 1 | 1 | 1 | 1 | 0.08 | 1 | 0.88 | 1.08 | 0.931 | 1.059 | 0.88 | 0.983 | 0.954 | Serine hydroxymethyltransferase (Fragment) OS=Mus musculus GN=Shmt1 PE=1 SV=1 |
| 986 | 1 | UniProt n | I7HIQ2 | 47 | 28459 | 2 | 2 | 2 | 2 | 0.34 | 1 | 0.857 | 0.837 | 0.92 | 1.003 | 0.94 | 0.895 | 0.954 | C-X-C motif chemokine 16 (Fragment) OS=Mus musculus GN=Cxcl16 PE=1 SV=1 |
| 901 | 1 | UniProt n | Q9EP71 | 55 | 140366 | 1 | 1 | 1 | 1 | 0.03 | 1 | 0.926 | 0.904 | 0.989 | 0.887 | 0.987 | 0.942 | 0.953 | Ankycorbin OS=Mus musculus GN=Rai14 PE=1 SV=1 |
| 529 | 1 | UniProt n | Q9R1P1 | 129 | 27134 | 4 | 4 | 3 | 3 | 0.86 | 1 | 1.091 | 1.034 | 1.004 | 1.013 | 0.849 | 1.041 | 0.952 | Proteasome subunit beta type-3 OS=Mus musculus GN=Psmb3 PE=1 SV=1 |
| 396 | 1 | UniProt n | Q9QUR6 | 209 | 96084 | 10 | 10 | 8 | 8 | 0.42 | 1 | 0.912 | 0.95 | 0.964 | 0.943 | 0.947 | 0.953 | 0.951 | Prolyl endopeptidase OS=Mus musculus GN=Prep PE=1 SV=1 |
| 42 | 1 | UniProt n | Q9R1E6 | 2209 | 116158 | 86 | 86 | 26 | 26 | 2.34 | 1 | 1.027 | 0.987 | 0.962 | 0.935 | 0.956 | 1.005 | 0.951 | Ectonucleotide pyrophosphatase/phosphodiesterase family member 2 OS=Mus musculus GN=Enpp2 PE=1 SV=3 |
| 511 | 1 | UniProt n | Q9WUU7 | 137 | 37872 | 12 | 12 | 5 | 5 | 0.95 | 1 | 0.966 | 0.942 | 0.979 | 0.946 | 0.928 | 0.969 | 0.951 | Cathepsin Z OS=Mus musculus GN=Ctsz PE=1 SV=1 |
| 1189 | 1 | UniProt n | Q9QUH0 | 29 | 14527 | 1 | 1 | 1 | 1 | 0.33 | 1 | 0.856 | 1.058 | 0.89 | 1.026 | 0.941 | 0.968 | 0.951 | Glutaredoxin-1 OS=Mus musculus GN=Glrx PE=1 SV=3 |
| 972 | 1 | UniProt n | J3QP71 | 47 | 25580 | 1 | 1 | 1 | 1 | 0.18 | 1 | 1.001 | 0.995 | 0.973 | 0.973 | 0.906 | 0.999 | 0.950 | Basigin (Fragment) OS=Mus musculus GN=Bsg PE=1 SV=1 |
| 851 | 1 | UniProt n | P61967 | 60 | 23468 | 1 | 1 | 1 | 1 | 0.2 | 1 | 1.011 | 1.105 | 1.006 | 1.095 | 0.778 | 1.038 | 0.950 | AP-1 complex subunit sigma-1A OS=Mus musculus GN=Ap1s1 PE=1 SV=1 |
| 484 | 1 | UniProt n | Q3TGU7 | 153 | 57028 | 10 | 10 | 6 | 6 | 0.68 | 1 | 1 | 1.089 | 1.002 | 0.932 | 0.917 | 1.029 | 0.950 | Proliferation-associated 2G4 OS=Mus musculus GN=Pa2g4 PE=1 SV=1 |
| 954 | 1 | UniProt n | D3YVF4 | 49 | 16187 | 1 | 1 | 1 | 1 | 0.29 | 1 | 0.849 | 1.053 | 0.859 | 1.042 | 0.953 | 0.963 | 0.948 | 40S ribosomal protein S14 (Fragment) OS=Mus musculus GN=Rps14 PE=1 SV=1 |
| 73 | 1 | UniProt n | P10605 | 1348 | 42859 | 47 | 47 | 11 | 11 | 2.59 | 1 | 1.056 | 1.028 | 0.942 | 0.98 | 0.923 | 1.028 | 0.948 | Cathepsin B OS=Mus musculus GN=Ctsb PE=1 SV=2 |
| 491 | 1 | UniProt n | P62774 | 149 | 16945 | 3 | 3 | 1 | 1 | 0.28 | 1 | 0.874 | 0.925 | 1.061 | 1.097 | 0.731 | 0.932 | 0.948 | Myotrophin OS=Mus musculus GN=Mtpn PE=1 SV=2 |
| 864 | 1 | UniProt n | Q99PM0 | 58 | 23236 | 2 | 2 | 1 | 1 | 0.2 | 1 | 0.924 | 0.979 | 1.058 | 0.86 | 0.934 | 0.967 | 0.947 | Microfibril-associated glycoprotein 1 OS=Mus musculus GN=Mfap2 PE=1 SV=1 |
| 431 | 1 | UniProt n | Q80UE4 | 184 | 110024 | 4 | 4 | 2 | 2 | 0.08 | 1 | 0.928 | 0.993 | 0.988 | 0.934 | 0.915 | 0.973 | 0.945 | Band 4.1-like protein 2 OS=Mus musculus GN=Epb41l2 PE=1 SV=1 |
| 1224 | 1 | UniProt n | A2RSX9 | 26 | 47869 | 1 | 1 | 1 | 1 | 0.09 | 1 | 0.986 | 1.074 | 0.995 | 0.91 | 0.932 | 1.019 | 0.945 | ADP-ribosylation factor-interacting protein 1 OS=Mus musculus GN=Arfip1 PE=1 SV=1 |
| 46 | 1 | UniProt n | P09411 | 2161 | 57621 | 63 | 63 | 17 | 17 | 4.02 | 1 | 1.068 | 1.073 | 1.005 | 0.897 | 0.931 | 1.046 | 0.943 | Phosphoglycerate kinase 1 OS=Mus musculus GN=Pgk1 PE=1 SV=4 |
| 783 | 1 | UniProt n | D3Z6F5 | 68 | 63779 | 2 | 2 | 2 | 2 | 0.14 | 1 | 0.711 | 0.842 | 0.957 | 1.075 | 0.815 | 0.843 | 0.943 | ATP synthase subunit alpha OS=Mus musculus GN=Atp5a1 PE=1 SV=1 |
| 589 | 1 | UniProt n | Q9CPN9 | 107 | 30607 | 3 | 3 | 1 | 1 | 0.32 | 1 | 0.558 | 0.794 | 1.248 | 1.378 | 0.483 | 0.762 | 0.940 | RIKEN cDNA 2210010C04 gene OS=Mus musculus GN=2210010C04Rik PE=1 SV=1 |
| 944 | 1 | UniProt n | O09164 | 50 | 30388 | 1 | 1 | 1 | 1 | 0.15 | 1 | 1.214 | 0.862 | 1.212 | 0.6 | 1.14 | 1.015 | 0.939 | Extracellular superoxide dismutase [Cu-Zn] OS=Mus musculus GN=Sod3 PE=1 SV=1 |
| 368 | 2 | UniProt n | Q61656 | 103 | 80308 | 6 | 6 | 5 | 5 | 0.3 | 1 | 1.03 | 0.978 | 0.924 | 0.925 | 0.968 | 1.002 | 0.939 | Probable ATP-dependent RNA helicase DDX5 OS=Mus musculus GN=Ddx5 PE=1 SV=2 |
| 890 | 1 | UniProt n | P38647 | 55 | 90073 | 3 | 3 | 3 | 3 | 0.15 | 1 | 1.014 | 1.074 | 0.977 | 0.97 | 0.873 | 1.029 | 0.939 | Stress-70 protein, mitochondrial OS=Mus musculus GN=Hspa9 PE=1 SV=3 |
| 721 | 1 | UniProt n | E9PYI8 | 78 | 65430 | 5 | 5 | 3 | 3 | 0.21 | 1 | 0.964 | 1.063 | 0.98 | 0.958 | 0.877 | 1.008 | 0.937 | Ubiquitin carboxyl-terminal hydrolase 14 OS=Mus musculus GN=Usp14 PE=1 SV=1 |
| 823 | 1 | UniProt n | P83917 | 62 | 30318 | 2 | 2 | 2 | 2 | 0.32 | 1 | 0.888 | 1.014 | 1.008 | 0.96 | 0.85 | 0.966 | 0.937 | Chromobox protein homolog 1 OS=Mus musculus GN=Cbx1 PE=1 SV=1 |
| 984 | 1 | UniProt n | F2Z405 | 47 | 33331 | 1 | 1 | 1 | 1 | 0.13 | 1 | 0.882 | 1.055 | 0.869 | 0.981 | 0.964 | 0.976 | 0.937 | Serine protease inhibitor A3G OS=Mus musculus GN=Serpina3g PE=1 SV=1 |
| 893 | 1 | UniProt n | A0A0A6Y | 55 | 102226 | 1 | 1 | 1 | 1 | 0.04 | 1 | 0.912 | 0.945 | 0.864 | 1.157 | 0.821 | 0.952 | 0.936 | Protocadherin 7 (Fragment) OS=Mus musculus GN=Pcdh7 PE=1 SV=1 |
| 1252 | 1 | UniProt n | A0A0G2J | 23 | 72640 | 1 | 1 | 1 | 1 | 0.06 | 1 | 1.065 | 0.989 | 0.771 | 1.058 | 1.003 | 1.017 | 0.935 | Calcium/calmodulin-dependent protein kinase type II subunit delta OS=Mus musculus GN=Camk2d PE=1 SV=1 |
| 943 | 1 | UniProt n | P54823 | 50 | 65126 | 3 | 3 | 3 | 3 | 0.22 | 1 | 0.904 | 0.976 | 1.271 | 0.848 | 0.759 | 0.959 | 0.935 | Probable ATP-dependent RNA helicase DDX6 OS=Mus musculus GN=Ddx6 PE=1 SV=1 |
| 373 | 1 | UniProt n | Q8C483 | 229 | 77361 | 9 | 9 | 7 | 7 | 0.47 | 1 | 0.926 | 0.948 | 0.945 | 0.876 | 0.982 | 0.958 | 0.933 | Serine--tRNA ligase, cytoplasmic OS=Mus musculus GN=Sars PE=1 SV=1 |
| 472 | 1 | UniProt n | B2RXS4 | 159 | 236386 | 9 | 9 | 6 | 6 | 0.13 | 1 | 0.812 | 0.867 | 1.026 | 0.923 | 0.858 | 0.890 | 0.933 | Plexin-B2 OS=Mus musculus GN=Plxnb2 PE=1 SV=1 |
| 330 | 1 | UniProt n | Q9WU78 | 262 | 116197 | 12 | 12 | 8 | 8 | 0.34 | 1 | 0.995 | 1.071 | 0.931 | 0.938 | 0.93 | 1.021 | 0.933 | Programmed cell death 6-interacting protein OS=Mus musculus GN=Pdcd6ip PE=1 SV=3 |
| 859 | 1 | UniProt n | P27612 | 59 | 101700 | 2 | 2 | 2 | 2 | 0.09 | 1 | 0.963 | 0.983 | 1 | 0.853 | 0.952 | 0.982 | 0.933 | Phospholipase A-2-activating protein OS=Mus musculus GN=Plaa PE=1 SV=4 |
| 941 | 1 | UniProt n | P97346 | 50 | 55024 | 1 | 1 | 1 | 1 | 0.08 | 1 | 0.8 | 0.921 | 0.791 | 0.927 | 1.106 | 0.903 | 0.933 | Nucleoredoxin OS=Mus musculus GN=Nxn PE=1 SV=1 |
| 869 | 1 | UniProt n | Q99KV1 | 58 | 48973 | 2 | 2 | 2 | 2 | 0.19 | 1 | 0.835 | 0.828 | 1.021 | 0.904 | 0.877 | 0.884 | 0.932 | DnaJ homolog subfamily B member 11 OS=Mus musculus GN=Dnajb11 PE=1 SV=1 |
| 1154 | 1 | UniProt n | Q61543 | 31 | 163543 | 1 | 1 | 1 | 1 | 0.03 | 1 | 0.836 | 0.822 | 0.995 | 0.907 | 0.896 | 0.882 | 0.932 | Golgi apparatus protein 1 OS=Mus musculus GN=Glg1 PE=1 SV=1 |
| 41 | 1 | UniProt n | P33434 | 2221 | 88968 | 77 | 77 | 23 | 23 | 2.45 | 1 | 0.951 | 0.927 | 0.956 | 0.91 | 0.929 | 0.959 | 0.931 | 72 kDa type IV collagenase OS=Mus musculus GN=Mmp2 PE=1 SV=1 |
| 1266 | 1 | UniProt n | D3YVL0 | 20 | 139548 | 1 | 1 | 1 | 1 | 0.03 | 1 | 1.003 | 0.976 | 1.125 | 0.988 | 0.727 | 0.993 | 0.931 | Putative helicase MOV-10 OS=Mus musculus GN=Mov10 PE=1 SV=2 |
| 657 | 1 | UniProt n | P62257 | 89 | 24947 | 6 | 6 | 3 | 3 | 0.65 | 1 | 0.992 | 0.986 | 0.972 | 0.921 | 0.902 | 0.993 | 0.931 | Ubiquitin-conjugating enzyme E2 H OS=Mus musculus GN=Ube2h PE=1 SV=1 |
| 476 | 1 | UniProt n | Q3UZP7 | 157 | 23677 | 2 | 2 | 1 | 1 | 0.19 | 1 | 1.019 | 1.122 | 0.905 | 0.928 | 0.958 | 1.046 | 0.930 | Paralemmin-1 OS=Mus musculus GN=Palm PE=1 SV=1 |
| 158 | 1 | UniProt n | P10126 | 615 | 64960 | 37 | 37 | 12 | 12 | 1.33 | 1 | 1.057 | 1.121 | 0.929 | 0.914 | 0.946 | 1.058 | 0.930 | Elongation factor 1-alpha 1 OS=Mus musculus GN=Eef1a1 PE=1 SV=3 |
| 498 | 1 | UniProt n | A2AD97 | 146 | 36154 | 7 | 7 | 3 | 3 | 0.59 | 1 | 0.951 | 0.885 | 0.969 | 0.903 | 0.913 | 0.944 | 0.928 | Matrix-remodeling-associated protein 8 (Fragment) OS=Mus musculus GN=Mxra8 PE=1 SV=8 |
| 660 | 1 | UniProt n | Q60817 | 89 | 28541 | 3 | 3 | 2 | 2 | 0.34 | 1 | 0.977 | 1.125 | 0.958 | 1.022 | 0.814 | 1.032 | 0.927 | Nascent polypeptide-associated complex subunit alpha OS=Mus musculus GN=Naca PE=1 SV=1 |
| 682 | 1 | UniProt n | P50428 | 86 | 58008 | 4 | 4 | 1 | 1 | 0.24 | 1 | 1.105 | 1.352 | 0.96 | 0.907 | 0.914 | 1.143 | 0.927 | Arylsulfatase A OS=Mus musculus GN=Arsa PE=1 SV=2 |
| 842 | 1 | UniProt n | D3Z111 | 61 | 24714 | 2 | 2 | 1 | 1 | 0.18 | 1 | 1.102 | 1.153 | 0.892 | 0.933 | 0.956 | 1.083 | 0.927 | Acyl-protein thioesterase 1 OS=Mus musculus GN=Lypla1 PE=1 SV=1 |
| 1167 | 1 | UniProt n | E9Q0U1 | 30 | 30663 | 1 | 1 | 1 | 1 | 0.15 | 1 | 0.947 | 1.056 | 0.993 | 0.851 | 0.941 | 1.000 | 0.926 | 26S proteasome non-ATPase regulatory subunit 13 OS=Mus musculus GN=Psmd13 PE=1 SV=1 |
| 428 | 1 | UniProt n | Q3TGS7 | 184 | 22718 | 8 | 8 | 3 | 3 | 0.74 | 1 | 1.096 | 1.122 | 0.93 | 0.923 | 0.926 | 1.071 | 0.926 | Sorting nexin 12, isoform CRA_b OS=Mus musculus GN=Snx12 PE=1 SV=1 |
| 1151 | 1 | UniProt n | Q9WUL7 | 31 | 25175 | 1 | 1 | 1 | 1 | 0.18 | 1 | 0.952 | 1.067 | 0.871 | 1.145 | 0.797 | 1.005 | 0.926 | ADP-ribosylation factor-like protein 3 OS=Mus musculus GN=Arl3 PE=1 SV=1 |
| 404 | 1 | UniProt n | A0A087W | 205 | 17198 | 6 | 6 | 3 | 3 | 1.63 | 1 | 0.989 | 1.008 | 0.917 | 0.991 | 0.874 | 0.999 | 0.926 | Elongin-C OS=Mus musculus GN=Eloc PE=1 SV=1 |
| 1258 | 1 | UniProt n | Q497S0 | 21 | 29662 | 1 | 1 | 1 | 1 | 0.15 | 1 | 0.959 | 1.1 | 0.863 | 1.02 | 0.894 | 1.018 | 0.923 | EG546282 protein OS=Mus musculus GN=Gm5935 PE=2 SV=1 |
| 750 | 1 | UniProt n | P28271 | 73 | 114084 | 2 | 2 | 2 | 2 | 0.08 | 1 | 0.898 | 1.039 | 0.935 | 0.966 | 0.871 | 0.977 | 0.923 | Cytoplasmic aconitate hydratase OS=Mus musculus GN=Aco1 PE=1 SV=3 |
| 493 | 1 | UniProt n | Q05186 | 148 | 46303 | 7 | 7 | 4 | 4 | 0.44 | 1 | 1.001 | 1.045 | 0.947 | 0.898 | 0.925 | 1.015 | 0.923 | Reticulocalbin-1 OS=Mus musculus GN=Rcn1 PE=1 SV=1 |
| 649 | 1 | UniProt n | Q99JF5 | 90 | 49796 | 3 | 3 | 2 | 2 | 0.18 | 1 | 0.966 | 1.086 | 0.902 | 0.854 | 1.02 | 1.016 | 0.923 | Diphosphomevalonate decarboxylase OS=Mus musculus GN=Mvd PE=1 SV=2 |
| 1138 | 1 | UniProt n | P62242 | 33 | 34155 | 1 | 1 | 1 | 1 | 0.13 | 1 | 1.035 | 1.102 | 0.816 | 1.095 | 0.877 | 1.045 | 0.922 | 40S ribosomal protein S8 OS=Mus musculus GN=Rps8 PE=1 SV=2 |
| 483 | 1 | UniProt n | A0A0R4J1 | 154 | 66014 | 6 | 6 | 5 | 5 | 0.38 | 1 | 0.967 | 0.938 | 0.893 | 0.921 | 0.95 | 0.968 | 0.921 | Biotinidase OS=Mus musculus GN=Btd PE=1 SV=1 |
| 368 | 1 | UniProt n | Q3U741 | 230 | 83084 | 9 | 9 | 7 | 7 | 0.43 | 1 | 1.011 | 0.995 | 0.918 | 0.909 | 0.936 | 1.002 | 0.921 | DEAD (Asp-Glu-Ala-Asp) box polypeptide 17, isoform CRA_a OS=Mus musculus GN=Ddx17 PE=1 SV=1 |
| 1009 | 1 | UniProt n | A0A1L1SV | 44 | 187059 | 1 | 1 | 1 | 1 | 0.02 | 1 | 1.005 | 1.157 | 0.981 | 1.044 | 0.762 | 1.052 | 0.921 | Ubiquitin carboxyl-terminal hydrolase 47 OS=Mus musculus GN=Usp47 PE=1 SV=1 |
| 1123 | 1 | UniProt n | E9Q5E1 | 34 | 136088 | 1 | 1 | 1 | 1 | 0.03 | 1 | 1.55 | 0.796 | 0.669 | 0.827 | 1.409 | 1.073 | 0.920 | Protocadherin-19 OS=Mus musculus GN=Pcdh19 PE=1 SV=1 |
| 818 | 1 | UniProt n | Q9EP52 | 63 | 29279 | 2 | 2 | 2 | 2 | 0.33 | 1 | 0.896 | 0.907 | 0.843 | 1.06 | 0.872 | 0.933 | 0.920 | Twisted gastrulation protein homolog 1 OS=Mus musculus GN=Twsg1 PE=1 SV=1 |
| 988 | 1 | UniProt n | Q922Q8 | 46 | 46783 | 1 | 1 | 1 | 1 | 0.09 | 1 | 1.014 | 0.91 | 0.949 | 0.849 | 0.966 | 0.974 | 0.920 | Leucine-rich repeat-containing protein 59 OS=Mus musculus GN=Lrrc59 PE=1 SV=1 |
| 760 | 1 | UniProt n | A0A0R4J0 | 72 | 41998 | 1 | 1 | 1 | 1 | 0.11 | 1 | 0.872 | 1.046 | 1.021 | 0.912 | 0.835 | 0.970 | 0.920 | Chitinase domain-containing protein 1 OS=Mus musculus GN=Chid1 PE=1 SV=1 |
| 735 | 1 | UniProt n | A0A0G2JD | 76 | 123092 | 2 | 2 | 2 | 2 | 0.07 | 1 | 0.924 | 0.963 | 0.963 | 0.958 | 0.837 | 0.962 | 0.917 | Ubiquitin associated protein 2-like, isoform CRA_g OS=Mus musculus GN=Ubap2l PE=1 SV=1 |
| 631 | 1 | UniProt n | P97825 | 96 | 18809 | 5 | 5 | 3 | 3 | 2.04 | 1 | 0.984 | 1.004 | 0.871 | 0.958 | 0.925 | 0.996 | 0.917 | Jupiter microtubule associated homolog 1 OS=Mus musculus GN=JPT1 PE=1 SV=3 |
| 917 | 1 | UniProt n | A2A4J1 | 53 | 29263 | 2 | 2 | 2 | 2 | 0.33 | 1 | 0.89 | 0.959 | 1.051 | 0.916 | 0.801 | 0.949 | 0.917 | Proteasome activator complex subunit 3 (Fragment) OS=Mus musculus GN=Psme3 PE=1 SV=1 |
| 596 | 1 | UniProt n | H3BJ30 | 105 | 64582 | 2 | 2 | 2 | 2 | 0.14 | 1 | 0.992 | 0.938 | 0.915 | 0.943 | 0.893 | 0.976 | 0.917 | Cleavage and polyadenylation-specificity factor subunit 6 OS=Mus musculus GN=Cpsf6 PE=1 SV=1 |
| 329 | 1 | UniProt n | Q02053 | 263 | 135431 | 12 | 12 | 10 | 10 | 0.37 | 1 | 0.943 | 1.047 | 0.885 | 0.969 | 0.898 | 0.996 | 0.917 | Ubiquitin-like modifier-activating enzyme 1 OS=Mus musculus GN=Uba1 PE=1 SV=1 |
| 1233 | 1 | UniProt n | Q8VCR4 | 25 | 13722 | 1 | 1 | 1 | 1 | 0.35 | 1 | 0.904 | 0.922 | 0.889 | 0.919 | 0.94 | 0.941 | 0.916 | 0610038D11Rik protein OS=Mus musculus GN=Trmt112 PE=1 SV=1 |
| 295 | 1 | UniProt n | P29416 | 301 | 68031 | 11 | 11 | 6 | 6 | 0.45 | 1 | 0.919 | 0.961 | 0.87 | 0.933 | 0.942 | 0.959 | 0.914 | Beta-hexosaminidase subunit alpha OS=Mus musculus GN=Hexa PE=1 SV=2 |
| 1119 | 1 | UniProt n | A0A1Y7VJ | 34 | 69396 | 1 | 1 | 1 | 1 | 0.06 | 1 | 0.574 | 0.953 | 0.859 | 0.971 | 0.914 | 0.818 | 0.914 | Zinc finger protein 934 OS=Mus musculus GN=Zfp934 PE=4 SV=1 |
| 155 | 1 | UniProt n | Q3TKX1 | 643 | 49341 | 18 | 18 | 4 | 4 | 0.53 | 1 | 1.004 | 0.965 | 0.902 | 0.867 | 0.974 | 0.990 | 0.913 | ATPase, H+ transporting, lysosomal accessory protein OS=Mus musculus GN=Atp6ap1 PE=1 SV=1 |
| 651 | 1 | UniProt n | Q9Z1T1 | 90 | 149331 | 2 | 2 | 2 | 2 | 0.06 | 1 | 0.823 | 1.036 | 0.888 | 0.949 | 0.903 | 0.948 | 0.913 | AP-3 complex subunit beta-1 OS=Mus musculus GN=Ap3b1 PE=1 SV=2 |
| 938 | 1 | UniProt n | P54751 | 50 | 46016 | 2 | 2 | 2 | 2 | 0.2 | 1 | 0.774 | 0.892 | 0.958 | 0.905 | 0.877 | 0.884 | 0.913 | CMP-N-acetylneuraminate-beta-galactosamide-alpha-2,3-sialyltransferase 1 OS=Mus musculus GN=St3gal1 PE=2 SV=1 |
| 541 | 1 | UniProt n | Q3TGS9 | 125 | 51905 | 4 | 4 | 2 | 2 | 0.28 | 1 | 1.018 | 1.051 | 0.951 | 0.956 | 0.836 | 1.023 | 0.913 | ADP-ribosylation factor GTPase-activating protein 1 OS=Mus musculus GN=Arfgap1 PE=1 SV=1 |
| 808 | 1 | UniProt n | A0A1W2P | 64 | 9553 | 1 | 1 | 1 | 1 | 0.53 | 1 | 1.063 | 1.069 | 0.878 | 0.988 | 0.874 | 1.044 | 0.912 | Enhancer of rudimentary homolog OS=Mus musculus GN=Erh PE=1 SV=1 |
| 1075 | 1 | UniProt n | O35674 | 38 | 115917 | 1 | 1 | 1 | 1 | 0.04 | 1 | 1.013 | 0.946 | 1.117 | 0.853 | 0.79 | 0.986 | 0.910 | Disintegrin and metalloproteinase domain-containing protein 19 OS=Mus musculus GN=Adam19 PE=2 SV=2 |
| 407 | 1 | UniProt n | Q9WVJ3 | 204 | 59432 | 15 | 15 | 9 | 9 | 0.9 | 1 | 0.976 | 0.987 | 0.921 | 0.891 | 0.914 | 0.988 | 0.909 | Carboxypeptidase Q OS=Mus musculus GN=Cpq PE=1 SV=1 |
| 679 | 1 | UniProt n | Q9D1M0 | 86 | 41433 | 4 | 4 | 3 | 3 | 0.36 | 1 | 0.84 | 0.934 | 1.145 | 0.829 | 0.789 | 0.922 | 0.908 | Protein SEC13 homolog OS=Mus musculus GN=Sec13 PE=1 SV=3 |
| 482 | 1 | UniProt n | A0A1B0G | 154 | 34561 | 4 | 4 | 3 | 3 | 0.44 | 1 | 1.025 | 1.043 | 0.952 | 0.952 | 0.826 | 1.023 | 0.908 | Reticulocalbin-3 (Fragment) OS=Mus musculus GN=Rcn3 PE=1 SV=1 |
| 435 | 1 | UniProt n | D3YW23 | 180 | 12906 | 13 | 13 | 3 | 3 | 1.61 | 1 | 1.124 | 0.957 | 0.877 | 0.947 | 0.901 | 1.025 | 0.908 | C-X-C motif chemokine OS=Mus musculus GN=Cxcl10 PE=3 SV=1 |
| 224 | 1 | UniProt n | F8WIR1 | 388 | 53681 | 18 | 18 | 7 | 7 | 1.03 | 1 | 0.965 | 1.052 | 0.929 | 0.901 | 0.893 | 1.005 | 0.908 | Cathepsin D OS=Mus musculus GN=Ctsd PE=1 SV=2 |
| 477 | 1 | UniProt n | Q8BWY3 | 156 | 60744 | 8 | 8 | 6 | 6 | 0.52 | 1 | 0.988 | 1.056 | 0.966 | 0.914 | 0.846 | 1.014 | 0.907 | Eukaryotic peptide chain release factor subunit 1 OS=Mus musculus GN=Etf1 PE=1 SV=4 |
| 1059 | 1 | UniProt n | Q05D44 | 40 | 191136 | 1 | 1 | 1 | 1 | 0.02 | 1 | 0.804 | 1.07 | 0.932 | 0.902 | 0.888 | 0.951 | 0.907 | Eukaryotic translation initiation factor 5B OS=Mus musculus GN=Eif5b PE=1 SV=2 |
| 517 | 1 | UniProt n | P50247 | 134 | 57502 | 8 | 8 | 7 | 7 | 0.67 | 1 | 1.033 | 1.112 | 0.929 | 0.857 | 0.937 | 1.047 | 0.907 | Adenosylhomocysteinase OS=Mus musculus GN=Ahcy PE=1 SV=3 |
| 1109 | 1 | UniProt n | B1ATE0 | 35 | 49175 | 1 | 1 | 1 | 1 | 0.09 | 1 | 0.85 | 0.892 | 0.907 | 0.886 | 0.928 | 0.912 | 0.907 | N-sulfoglucosamine sulfohydrolase (sulfamidase) OS=Mus musculus GN=Sgsh PE=1 SV=1 |
| 558 | 1 | UniProt n | A2AUD5 | 120 | 29965 | 2 | 2 | 2 | 2 | 0.32 | 1 | 1.022 | 1.064 | 0.878 | 0.907 | 0.936 | 1.028 | 0.907 | Tumor protein D54 OS=Mus musculus GN=Tpd52l2 PE=1 SV=1 |
| 433 | 1 | UniProt n | A0A0R4J0 | 181 | 38410 | 7 | 7 | 5 | 5 | 0.73 | 1 | 1.023 | 0.955 | 0.783 | 0.863 | 1.103 | 0.992 | 0.907 | N-acetylglucosamine-1-phosphotransferase subunit gamma OS=Mus musculus GN=Gnptg PE=1 SV=1 |
| 741 | 1 | UniProt n | P21956 | 75 | 57907 | 3 | 3 | 3 | 3 | 0.24 | 1 | 0.841 | 0.87 | 0.933 | 0.878 | 0.909 | 0.901 | 0.906 | Lactadherin OS=Mus musculus GN=Mfge8 PE=1 SV=3 |
| 1157 | 1 | UniProt n | Q8JZX4 | 31 | 56716 | 1 | 1 | 1 | 1 | 0.08 | 1 | 0.858 | 0.879 | 1.427 | 0.757 | 0.688 | 0.910 | 0.906 | Splicing factor 45 OS=Mus musculus GN=Rbm17 PE=1 SV=1 |
| 957 | 1 | UniProt n | E9PZF5 | 49 | 22053 | 1 | 1 | 1 | 1 | 0.21 | 1 | 1.049 | 0.925 | 0.863 | 0.942 | 0.913 | 0.990 | 0.905 | Acidic leucine-rich nuclear phosphoprotein 32 family member E (Fragment) OS=Mus musculus GN=Anp32e PE=1 SV=1 |
| 143 | 1 | UniProt n | Q8BHN3 | 692 | 116338 | 27 | 27 | 15 | 15 | 0.79 | 1 | 0.918 | 0.956 | 0.914 | 0.964 | 0.841 | 0.957 | 0.905 | Neutral alpha-glucosidase AB OS=Mus musculus GN=Ganab PE=1 SV=1 |
| 1074 | 1 | UniProt n | O88531 | 38 | 41924 | 1 | 1 | 1 | 1 | 0.11 | 1 | 1.005 | 0.999 | 0.964 | 0.816 | 0.941 | 1.001 | 0.905 | Palmitoyl-protein thioesterase 1 OS=Mus musculus GN=Ppt1 PE=1 SV=2 |
| 1072 | 1 | UniProt n | Q99M71 | 37 | 28529 | 3 | 3 | 3 | 3 | 0.55 | 1 | 1.013 | 0.967 | 0.93 | 0.921 | 0.864 | 0.993 | 0.905 | Mammalian ependymin-related protein 1 OS=Mus musculus GN=Epdr1 PE=1 SV=1 |
| 1065 | 1 | UniProt n | G3UVV4 | 39 | 121422 | 1 | 1 | 1 | 1 | 0.04 | 1 | 1.087 | 1.069 | 0.892 | 0.863 | 0.961 | 1.051 | 0.904 | Hexokinase 1, isoform CRA_f OS=Mus musculus GN=Hk1 PE=1 SV=1 |
| 1184 | 1 | UniProt n | Q9D115 | 29 | 12843 | 1 | 1 | 1 | 1 | 0.38 | 1 | 0.93 | 1.186 | 0.989 | 1.012 | 0.737 | 1.033 | 0.904 | Zinc finger protein 706 OS=Mus musculus GN=Znf706 PE=1 SV=1 |
| 1211 | 1 | UniProt n | B1AT36 | 27 | 63636 | 1 | 1 | 1 | 1 | 0.07 | 1 | 1.067 | 1.093 | 0.708 | 1.076 | 0.968 | 1.053 | 0.903 | 26S proteasome non-ATPase regulatory subunit 12 OS=Mus musculus GN=Psmd12 PE=1 SV=1 |
| 999 | 1 | UniProt n | P61082 | 45 | 26896 | 2 | 2 | 1 | 1 | 0.17 | 1 | 0.908 | 1.032 | 0.973 | 0.895 | 0.844 | 0.979 | 0.902 | NEDD8-conjugating enzyme Ubc12 OS=Mus musculus GN=Ube2m PE=1 SV=1 |
| 1188 | 1 | UniProt n | Q6P1B9 | 29 | 65646 | 1 | 1 | 1 | 1 | 0.07 | 1 | 1.22 | 1.263 | 0.973 | 0.95 | 0.792 | 1.155 | 0.901 | Bin1 protein OS=Mus musculus GN=Bin1 PE=1 SV=1 |
| 336 | 1 | UniProt n | Q3TW96 | 254 | 62955 | 10 | 10 | 5 | 5 | 0.4 | 1 | 0.962 | 1.044 | 0.98 | 0.883 | 0.846 | 1.001 | 0.901 | UDP-N-acetylhexosamine pyrophosphorylase-like protein 1 OS=Mus musculus GN=Uap1l1 PE=1 SV=1 |

| | | | | | | | | | | | | | | | | | | | | |
|---|---|---|---|---|---|---|---|---|---|---|---|---|---|---|---|---|---|---|---|---|
| 184 | 1 | UniProt | n | D3YWF6 | 499 | 33328 | 19 | 19 | 6 | 6 | 1.13 | 1 | 0.971 | 1.008 | 0.878 | 0.956 | 0.867 | 0.993 | 0.899 | Ubiquitin thioesterase OTUB1 OS=Mus musculus GN=Otub1 PE=1 SV=1 |
| 85 | 1 | UniProt | n | A0A1Y7VJ | 1166 | 54837 | 40 | 40 | 11 | 11 | 2.18 | 1 | 0.996 | 0.951 | 0.862 | 0.86 | 0.981 | 0.982 | 0.899 | Fibulin-5 OS=Mus musculus GN=Fbln5 PE=4 SV=1 |
| 1164 | 1 | UniProt | n | A0A1B0GI | 31 | 20147 | 2 | 2 | 2 | 2 | 0.51 | 1 | 0.904 | 0.889 | 0.948 | 0.894 | 0.854 | 0.930 | 0.898 | 40S ribosomal protein S11 OS=Mus musculus GN=Rps11 PE=1 SV=1 |
| 630 | 1 | UniProt | n | P48758 | 96 | 37240 | 3 | 3 | 3 | 3 | 0.4 | 1 | 0.912 | 0.929 | 0.908 | 0.946 | 0.84 | 0.946 | 0.897 | Carbonyl reductase [NADPH] 1 OS=Mus musculus GN=Cbr1 PE=1 SV=3 |
| 1102 | 1 | UniProt | n | Q9QWR8 | 35 | 53397 | 2 | 2 | 2 | 2 | 0.17 | 1 | 1.019 | 1.22 | 0.954 | 0.828 | 0.912 | 1.075 | 0.896 | Alpha-N-acetylgalactosaminidase OS=Mus musculus GN=Naga PE=1 SV=2 |
| 551 | 1 | UniProt | n | H3BJC6 | 122 | 17978 | 3 | 3 | 2 | 2 | 0.59 | 1 | 0.903 | 0.876 | 0.886 | 0.845 | 0.962 | 0.925 | 0.896 | S-formylglutathione hydrolase (Fragment) OS=Mus musculus GN=Esd PE=1 SV=1 |
| 428 | 2 | UniProt | n | O70492 | 97 | 22138 | 4 | 4 | 3 | 3 | 0.76 | 1 | 1.138 | 1.188 | 0.874 | 0.936 | 0.879 | 1.106 | 0.896 | Sorting nexin-3 OS=Mus musculus GN=Snx3 PE=1 SV=3 |
| 14 | 1 | UniProt | n | E9PWQ3 | 5043 | 402761 | 190 | 190 | 76 | 76 | 1.35 | 1 | 0.99 | 1.052 | 0.904 | 0.841 | 0.945 | 1.014 | 0.896 | Collagen, type VI, alpha 3 OS=Mus musculus GN=Col6a3 PE=1 SV=2 |
| 694 | 2 | UniProt | n | Q9WV55 | 77 | 35322 | 4 | 4 | 3 | 3 | 0.43 | 1 | 1.124 | 1.256 | 0.863 | 0.918 | 0.906 | 1.122 | 0.895 | Vesicle-associated membrane protein-associated protein A OS=Mus musculus GN=Vapa PE=1 SV=2 |
| 966 | 1 | UniProt | n | A0A087W | 48 | 170660 | 2 | 2 | 2 | 2 | 0.05 | 1 | 0.852 | 0.858 | 0.882 | 0.822 | 0.99 | 0.901 | 0.895 | ATP-dependent RNA helicase A OS=Mus musculus GN=Dhx9 PE=1 SV=1 |
| 14 | 2 | UniProt | n | A0A087W | 4962 | 332908 | 187 | 187 | 73 | 73 | 1.71 | 1 | 0.989 | 1.054 | 0.903 | 0.841 | 0.945 | 1.014 | 0.895 | Collagen, type VI, alpha 3 OS=Mus musculus GN=Col6a3 PE=1 SV=1 |
| 84 | 1 | UniProt | n | P19157 | 1171 | 27687 | 39 | 39 | 7 | 7 | 5.18 | 1 | 1.002 | 1.005 | 0.888 | 0.892 | 0.906 | 1.002 | 0.895 | Glutathione S-transferase P 1 OS=Mus musculus GN=Gstp1 PE=1 SV=2 |
| 1234 | 1 | UniProt | n | A0A0G2JD | 25 | 7416 | 1 | 1 | 1 | 1 | 0.72 | 1 | 0.898 | 0.976 | 0.992 | 0.797 | 0.907 | 0.957 | 0.895 | Guanylate-binding protein 3 (Fragment) OS=Mus musculus GN=Gbp3 PE=1 SV=1 |
| 145 | 1 | UniProt | n | P58022 | 689 | 100162 | 32 | 32 | 13 | 13 | 1.05 | 1 | 0.97 | 0.988 | 0.925 | 0.877 | 0.88 | 0.986 | 0.894 | Lysyl oxidase homolog 2 OS=Mus musculus GN=Loxl2 PE=1 SV=2 |
| 1247 | 1 | UniProt | n | Q5PSV9 | 23 | 209039 | 1 | 1 | 1 | 1 | 0.02 | 1 | 0.814 | 0.983 | 0.885 | 1.041 | 0.773 | 0.928 | 0.893 | Mediator of DNA damage checkpoint protein 1 OS=Mus musculus GN=Mdc1 PE=1 SV=1 |
| 692 | 1 | UniProt | n | D3Z7E6 | 84 | 21750 | 4 | 4 | 2 | 2 | 0.47 | 1 | 0.855 | 0.86 | 0.936 | 0.898 | 0.847 | 0.903 | 0.893 | Platelet-activating factor acetylhydrolase IB subunit gamma OS=Mus musculus GN=Pafah1b3 PE=1 SV=1 |
| 806 | 1 | UniProt | n | A0A0J9YV | 64 | 57967 | 2 | 2 | 1 | 1 | 0.08 | 1 | 0.851 | 0.946 | 0.916 | 0.899 | 0.864 | 0.930 | 0.893 | Protein phosphatase 1G OS=Mus musculus GN=Ppm1g PE=1 SV=1 |
| 1209 | 1 | UniProt | n | Q64695 | 27 | 30277 | 1 | 1 | 1 | 1 | 0.15 | 1 | 0.808 | 0.896 | 0.856 | 0.908 | 0.915 | 0.898 | 0.893 | Endothelial protein C receptor OS=Mus musculus GN=Procr PE=1 SV=3 |
| 764 | 1 | UniProt | n | P00375 | 71 | 26810 | 2 | 2 | 1 | 1 | 0.17 | 1 | 0.991 | 1.042 | 0.908 | 0.892 | 0.878 | 1.011 | 0.893 | Dihydrofolate reductase OS=Mus musculus GN=Dhfr PE=1 SV=3 |
| 300 | 1 | UniProt | n | Q61753 | 292 | 65102 | 9 | 9 | 6 | 6 | 0.48 | 1 | 0.961 | 1.062 | 0.849 | 0.909 | 0.917 | 1.007 | 0.891 | D-3-phosphoglycerate dehydrogenase OS=Mus musculus GN=Phgdh PE=1 SV=3 |
| 748 | 1 | UniProt | n | E9Q0X4 | 74 | 125111 | 4 | 4 | 3 | 3 | 0.11 | 1 | 0.842 | 0.977 | 0.922 | 0.883 | 0.869 | 0.937 | 0.891 | Collagen alpha-1(XVI) chain OS=Mus musculus GN=Col16a1 PE=1 SV=1 |
| 648 | 1 | UniProt | n | Q9WV54 | 91 | 53176 | 6 | 6 | 4 | 4 | 0.37 | 1 | 0.964 | 0.98 | 0.92 | 0.892 | 0.862 | 0.981 | 0.891 | Acid ceramidase OS=Mus musculus GN=Asah1 PE=1 SV=1 |
| 722 | 1 | UniProt | n | Q9QUR7 | 75 | 22102 | 2 | 2 | 2 | 2 | 0.46 | 1 | 0.967 | 0.967 | 0.849 | 0.888 | 0.938 | 0.978 | 0.891 | Peptidyl-prolyl cis-trans isomerase NIMA-interacting 1 OS=Mus musculus GN=Pin1 PE=1 SV=1 |
| 650 | 1 | UniProt | n | A2AFI3 | 90 | 37857 | 2 | 2 | 1 | 1 | 0.12 | 1 | 1.06 | 1.065 | 0.919 | 0.851 | 0.904 | 1.041 | 0.891 | RNA binding motif protein, X chromosome, isoform CRA b OS=Mus musculus GN=Rbmx PE=1 SV=1 |
| 658 | 1 | UniProt | n | Q8BG32 | 89 | 58588 | 4 | 4 | 4 | 4 | 0.33 | 1 | 0.914 | 0.919 | 0.942 | 0.883 | 0.848 | 0.944 | 0.890 | 26S proteasome non-ATPase regulatory subunit 11 OS=Mus musculus GN=Psmd11 PE=1 SV=3 |
| 1245 | 1 | UniProt | n | D3YYI8 | 24 | 68249 | 1 | 1 | 1 | 1 | 0.06 | 1 | 0.922 | 0.904 | 1.005 | 0.825 | 0.85 | 0.941 | 0.890 | Histone deacetylase OS=Mus musculus GN=Gm10093 PE=3 SV=1 |
| 796 | 1 | UniProt | n | O88325 | 66 | 88479 | 2 | 2 | 2 | 2 | 0.1 | 1 | 0.907 | 0.954 | 0.869 | 0.903 | 0.897 | 0.953 | 0.890 | Alpha-N-acetylglucosaminidase OS=Mus musculus GN=Naglu PE=1 SV=1 |
| 1168 | 1 | UniProt | n | Q9DB05 | 30 | 41750 | 1 | 1 | 1 | 1 | 0.11 | 1 | 0.835 | 0.87 | 0.929 | 0.903 | 0.835 | 0.899 | 0.888 | Alpha-soluble NSF attachment protein OS=Mus musculus GN=Napa PE=1 SV=1 |
| 296 | 1 | UniProt | n | Q99KI0 | 301 | 102739 | 14 | 14 | 8 | 8 | 0.39 | 1 | 1.025 | 1.066 | 0.901 | 0.853 | 0.909 | 1.030 | 0.887 | Aconitate hydratase, mitochondrial OS=Mus musculus GN=Aco2 PE=1 SV=1 |
| 1162 | 1 | UniProt | n | G3X977 | 31 | 127303 | 1 | 1 | 1 | 1 | 0.03 | 1 | 0.961 | 0.937 | 0.848 | 0.874 | 0.942 | 0.966 | 0.887 | Inter-alpha trypsin inhibitor, heavy chain 2 OS=Mus musculus GN=Itih2 PE=1 SV=1 |
| 672 | 1 | UniProt | n | A2AAW9 | 88 | 45752 | 4 | 4 | 3 | 3 | 0.32 | 1 | 1.033 | 1.094 | 0.865 | 0.919 | 0.877 | 1.042 | 0.887 | Eukaryotic translation initiation factor 2 subunit 3, X-linked OS=Mus musculus GN=Eif2s3x PE=1 SV=1 |
| 703 | 1 | UniProt | n | Q9CWJ9 | 81 | 75238 | 5 | 5 | 5 | 5 | 0.32 | 1 | 0.894 | 0.991 | 0.867 | 0.841 | 0.954 | 0.960 | 0.886 | Bifunctional purine biosynthesis protein PURH OS=Mus musculus GN=Atic PE=1 SV=2 |
| 1041 | 1 | UniProt | n | A0A087W | 41 | 28259 | 1 | 1 | 1 | 1 | 0.16 | 1 | 0.876 | 1.031 | 0.95 | 0.892 | 0.819 | 0.967 | 0.885 | Aminoacylase-1 (Fragment) OS=Mus musculus GN=Acy1 PE=1 SV=1 |
| 400 | 1 | UniProt | n | P46061 | 208 | 76073 | 9 | 9 | 7 | 7 | 0.48 | 1 | 1.014 | 1.108 | 0.916 | 0.889 | 0.852 | 1.040 | 0.885 | Ran GTPase-activating protein 1 OS=Mus musculus GN=Rangap1 PE=1 SV=2 |
| 63 | 1 | UniProt | n | P55065 | 1523 | 60382 | 57 | 57 | 15 | 15 | 4.37 | 1 | 1.034 | 1.067 | 0.904 | 0.928 | 0.823 | 1.033 | 0.884 | Phospholipid transfer protein OS=Mus musculus GN=Pltp PE=1 SV=1 |
| 1047 | 1 | UniProt | n | Q9DBF1 | 41 | 67148 | 1 | 1 | 1 | 1 | 0.07 | 1 | 1.007 | 1.014 | 0.843 | 0.847 | 0.966 | 1.007 | 0.884 | Alpha-aminoadipic semialdehyde dehydrogenase OS=Mus musculus GN=Aldh7a1 PE=1 SV=4 |
| 793 | 1 | UniProt | n | D3Z3X5 | 67 | 29858 | 3 | 3 | 2 | 2 | 0.32 | 1 | 0.948 | 0.946 | 0.821 | 0.953 | 0.881 | 0.964 | 0.883 | Glutathione S-transferase theta-1 OS=Mus musculus GN=Gstt1 PE=1 SV=1 |
| 294 | 1 | UniProt | n | P20060 | 301 | 70221 | 17 | 17 | 11 | 11 | 0.94 | 1 | 1.052 | 1.014 | 0.858 | 0.895 | 0.896 | 1.022 | 0.883 | Beta-hexosaminidase subunit beta OS=Mus musculus GN=Hexb PE=1 SV=2 |
| 629 | 1 | UniProt | n | A0A0R4J0 | 96 | 46009 | 4 | 4 | 3 | 3 | 0.32 | 1 | 1.036 | 0.976 | 0.913 | 0.872 | 0.864 | 1.004 | 0.883 | Glutaminyl-peptide cyclotransferase OS=Mus musculus GN=Qpct PE=1 SV=1 |
| 1101 | 1 | UniProt | n | Q9D8E6 | 36 | 64694 | 1 | 1 | 1 | 1 | 0.07 | 1 | 1.168 | 1.007 | 0.647 | 1.049 | 1.013 | 1.056 | 0.883 | 60S ribosomal protein L4 OS=Mus musculus GN=Rpl4 PE=1 SV=3 |
| 911 | 1 | UniProt | n | Q8BP47 | 54 | 75621 | 2 | 2 | 2 | 2 | 0.12 | 1 | 0.953 | 1.122 | 0.918 | 0.871 | 0.859 | 1.023 | 0.882 | Asparagine--tRNA ligase, cytoplasmic OS=Mus musculus GN=Nars PE=1 SV=2 |
| 1165 | 1 | UniProt | n | Q60749 | 31 | 56388 | 3 | 3 | 3 | 3 | 0.25 | 1 | 0.928 | 0.954 | 0.887 | 0.922 | 0.839 | 0.960 | 0.882 | KH domain-containing, RNA-binding, signal transduction-associated protein 1 OS=Mus musculus GN=Khdrbs1 PE=1 SV=2 |
| 45 | 2 | UniProt | n | P10493 | 1643 | 147876 | 65 | 65 | 25 | 25 | 1.11 | 1 | 0.944 | 0.928 | 0.897 | 0.866 | 0.882 | 0.957 | 0.882 | Nidogen-1 OS=Mus musculus GN=Nid1 PE=1 SV=2 |
| 1031 | 1 | UniProt | n | F8WIK0 | 42 | 41777 | 5 | 5 | 3 | 3 | 0.35 | 1 | 1.121 | 1.142 | 0.92 | 0.85 | 0.875 | 1.086 | 0.881 | Anamorsin OS=Mus musculus GN=Ciapin1 PE=1 SV=2 |
| 776 | 1 | UniProt | n | A0A0R4J1 | 69 | 89627 | 2 | 2 | 2 | 2 | 0.1 | 1 | 1.012 | 1.059 | 0.899 | 0.849 | 0.896 | 1.023 | 0.881 | Acylamino-acid-releasing enzyme (Fragment) OS=Mus musculus GN=Apeh PE=1 SV=1 |
| 1081 | 1 | UniProt | n | A0A0R4J2 | 38 | 132307 | 1 | 1 | 1 | 1 | 0.03 | 1 | 0.841 | 1.069 | 0.827 | 1.059 | 0.779 | 0.965 | 0.880 | Oncostatin-M-specific receptor subunit beta OS=Mus musculus GN=Osmr PE=1 SV=1 |
| 1201 | 1 | UniProt | n | Q91VH6 | 28 | 38251 | 1 | 1 | 1 | 1 | 0.12 | 1 | 1.004 | 1.196 | 0.817 | 0.796 | 1.049 | 1.063 | 0.880 | Protein MEMO1 OS=Mus musculus GN=Memo1 PE=1 SV=1 |
| 341 | 1 | UniProt | n | Q91ZJ5 | 248 | 69858 | 11 | 11 | 7 | 7 | 0.53 | 1 | 0.975 | 1.049 | 0.885 | 0.855 | 0.899 | 1.008 | 0.879 | UTP--glucose-1-phosphate uridylyltransferase OS=Mus musculus GN=Ugp2 PE=1 SV=3 |
| 716 | 1 | UniProt | n | A2ATU9 | 79 | 58944 | 2 | 2 | 2 | 2 | 0.15 | 1 | 1.021 | 1.114 | 0.878 | 0.853 | 0.906 | 1.044 | 0.879 | Histone acetyltransferase type B catalytic subunit OS=Mus musculus GN=Hat1 PE=1 SV=1 |
| 586 | 1 | UniProt | n | Q9D8S4 | 109 | 33598 | 9 | 9 | 4 | 4 | 0.65 | 1 | 0.992 | 1.005 | 0.815 | 0.914 | 0.91 | 0.999 | 0.878 | Oligoribonuclease, mitochondrial OS=Mus musculus GN=Rexo2 PE=1 SV=2 |
| 674 | 1 | UniProt | n | Q9JIF7 | 87 | 129212 | 4 | 4 | 4 | 4 | 0.14 | 1 | 0.987 | 1.085 | 0.89 | 0.9 | 0.845 | 1.023 | 0.878 | Coatomer subunit beta OS=Mus musculus GN=Copb1 PE=1 SV=1 |
| 26 | 2 | UniProt | n | P05063 | 131 | 46080 | 5 | 5 | 3 | 3 | 0.32 | 1 | 1.157 | 1.2 | 0.861 | 0.943 | 0.831 | 1.116 | 0.877 | Fructose-bisphosphate aldolase C OS=Mus musculus GN=Aldoc PE=1 SV=4 |
| 568 | 1 | UniProt | n | Q3TWV4 | 117 | 61498 | 6 | 6 | 6 | 6 | 0.51 | 1 | 1.019 | 1.098 | 0.883 | 0.908 | 0.839 | 1.038 | 0.876 | AP-2 complex subunit mu OS=Mus musculus GN=Ap2m1 PE=1 SV=1 |
| 1159 | 1 | UniProt | n | A0A087W | 31 | 108457 | 1 | 1 | 1 | 1 | 0.04 | 1 | 1.003 | 1.145 | 0.798 | 0.971 | 0.868 | 1.047 | 0.876 | Bcl-2-associated transcription factor 1 OS=Mus musculus GN=Bclaf1 PE=1 SV=1 |
| 936 | 1 | UniProt | n | Q9DBG5 | 51 | 53409 | 1 | 1 | 1 | 1 | 0.08 | 1 | 1.035 | 1.099 | 0.909 | 0.834 | 0.887 | 1.044 | 0.876 | Perilipin-3 OS=Mus musculus GN=Plin3 PE=1 SV=1 |
| 108 | 1 | UniProt | n | Q8CG16 | 971 | 92169 | 42 | 42 | 15 | 15 | 1.09 | 1 | 0.949 | 0.952 | 0.918 | 0.836 | 0.876 | 0.967 | 0.876 | Complement C1r-A subcomponent OS=Mus musculus GN=C1ra PE=1 SV=1 |
| 1000 | 1 | UniProt | n | Q8CG72 | 45 | 43316 | 2 | 2 | 2 | 2 | 0.22 | 1 | 0.955 | 1.077 | 1.024 | 0.852 | 0.769 | 1.009 | 0.875 | Poly(ADP-ribose) glycohydrolase ARH3 OS=Mus musculus GN=Adprhl2 PE=1 SV=1 |
| 752 | 1 | UniProt | n | Q8R016 | 73 | 63934 | 4 | 4 | 3 | 3 | 0.22 | 1 | 1.056 | 1.017 | 0.883 | 0.992 | 0.765 | 1.024 | 0.875 | Bleomycin hydrolase OS=Mus musculus GN=Blmh PE=1 SV=1 |
| 571 | 1 | UniProt | n | Q8R1B4 | 116 | 129440 | 2 | 2 | 1 | 1 | 0.03 | 1 | 0.856 | 0.967 | 0.935 | 0.884 | 0.809 | 0.939 | 0.874 | Eukaryotic translation initiation factor 3 subunit C OS=Mus musculus GN=Eif3c PE=1 SV=1 |
| 872 | 1 | UniProt | n | P17897 | 57 | 19585 | 2 | 2 | 1 | 1 | 0.24 | 1 | 0.68 | 0.674 | 0.638 | 0.726 | 1.442 | 0.771 | 0.874 | Lysozyme C-1 OS=Mus musculus GN=Lyz1 PE=1 SV=1 |
| 345 | 1 | UniProt | n | A0A0R4J0 | 247 | 107282 | 10 | 10 | 6 | 6 | 0.32 | 1 | 1.016 | 1.043 | 0.902 | 0.876 | 0.844 | 1.020 | 0.874 | Transforming growth factor beta receptor type 3 OS=Mus musculus GN=Tgfbr3 PE=1 SV=1 |
| 659 | 1 | UniProt | n | P61164 | 89 | 49980 | 2 | 2 | 1 | 1 | 0.09 | 1 | 0.888 | 1.077 | 0.824 | 0.879 | 0.918 | 0.985 | 0.873 | Alpha-centractin OS=Mus musculus GN=Actr1a PE=1 SV=1 |
| 538 | 1 | UniProt | n | Q9WVK4 | 125 | 74300 | 4 | 4 | 3 | 3 | 0.19 | 1 | 0.954 | 1.008 | 0.907 | 0.83 | 0.882 | 0.987 | 0.872 | EH domain-containing protein 1 OS=Mus musculus GN=Ehd1 PE=1 SV=1 |
| 262 | 1 | UniProt | n | Q64191 | 336 | 43192 | 11 | 11 | 5 | 5 | 0.63 | 1 | 0.964 | 0.972 | 0.889 | 0.888 | 0.84 | 0.979 | 0.872 | N(4)-(beta-N-acetylglucosaminyl)-L-asparaginase OS=Mus musculus GN=Aga PE=1 SV=1 |
| 857 | 1 | UniProt | n | P99025 | 59 | 11191 | 2 | 2 | 1 | 1 | 1.08 | 1 | 0.911 | 0.894 | 0.807 | 0.93 | 0.883 | 0.934 | 0.872 | GTP cyclohydrolase 1 feedback regulatory protein OS=Mus musculus GN=Gchfr PE=1 SV=3 |
| 742 | 1 | UniProt | n | P97494 | 74 | 84424 | 4 | 4 | 3 | 3 | 0.16 | 1 | 1.14 | 1.174 | 0.847 | 0.916 | 0.853 | 1.102 | 0.871 | Glutamate--cysteine ligase catalytic subunit OS=Mus musculus GN=Gclc PE=1 SV=4 |
| 792 | 1 | UniProt | n | Q9Z1Z0 | 67 | 125738 | 4 | 4 | 3 | 3 | 0.14 | 1 | 0.878 | 0.926 | 0.895 | 0.922 | 0.801 | 0.933 | 0.871 | General vesicular transport factor p115 OS=Mus musculus GN=Uso1 PE=1 SV=2 |
| 323 | 1 | UniProt | n | Q14AA6 | 267 | 30518 | 16 | 16 | 5 | 5 | 0.99 | 1 | 1.063 | 1.144 | 0.866 | 0.862 | 0.885 | 1.067 | 0.871 | GTP-binding nuclear protein Ran OS=Mus musculus GN=1700009N14Rik PE=2 SV=1 |
| 1045 | 1 | UniProt | n | D3YUB9 | 41 | 21698 | 3 | 3 | 1 | 1 | 0.21 | 1 | 1.274 | 1.38 | 0.8 | 0.961 | 0.857 | 1.207 | 0.870 | Polyadenylate-binding protein 2 (Fragment) OS=Mus musculus GN=Pabpn1 PE=4 SV=2 |
| 926 | 1 | UniProt | n | Q9CPT4 | 52 | 21409 | 4 | 4 | 3 | 3 | 0.79 | 1 | 1.031 | 1.066 | 0.882 | 0.942 | 0.793 | 1.032 | 0.870 | Myeloid-derived growth factor OS=Mus musculus GN=Mydgf PE=1 SV=1 |
| 948 | 1 | UniProt | n | P51655 | 50 | 75284 | 2 | 2 | 2 | 2 | 0.12 | 1 | 0.817 | 1.053 | 0.938 | 0.904 | 0.777 | 0.951 | 0.870 | Glypican-4 OS=Mus musculus GN=Gpc4 PE=1 SV=2 |
| 304 | 1 | UniProt | n | P34022 | 290 | 31325 | 11 | 11 | 4 | 4 | 0.71 | 1 | 1.064 | 1.031 | 0.922 | 0.848 | 0.842 | 1.031 | 0.870 | Ran-specific GTPase-activating protein OS=Mus musculus GN=Ranbp1 PE=1 SV=2 |
| 459 | 1 | UniProt | n | Q64010 | 168 | 37490 | 6 | 6 | 4 | 4 | 0.57 | 1 | 0.989 | 1.057 | 0.838 | 0.909 | 0.864 | 1.015 | 0.870 | Adapter molecule crk OS=Mus musculus GN=Crk PE=1 SV=1 |
| 1120 | 1 | UniProt | n | Q6ZPJ3 | 34 | 166840 | 1 | 1 | 1 | 1 | 0.03 | 1 | 1.059 | 0.905 | 0.88 | 0.817 | 0.915 | 0.986 | 0.870 | (E3-independent) E2 ubiquitin-conjugating enzyme UBE2O OS=Mus musculus GN=Ube2o PE=1 SV=3 |
| 3 | 2 | UniProt | n | P08121 | 8791 | 158730 | 292 | 292 | 46 | 46 | 5.68 | 1 | 1.01 | 1.002 | 0.889 | 0.86 | 0.859 | 1.004 | 0.869 | Collagen alpha-1(III) chain OS=Mus musculus GN=Col3a1 PE=1 SV=4 |
| 713 | 1 | UniProt | n | O70194 | 79 | 76318 | 4 | 4 | 3 | 3 | 0.18 | 1 | 0.934 | 1.015 | 0.893 | 0.857 | 0.858 | 0.982 | 0.869 | Eukaryotic translation initiation factor 3 subunit D OS=Mus musculus GN=Eif3d PE=1 SV=2 |
| 814 | 1 | UniProt | n | C0HKG5 | 64 | 35525 | 6 | 6 | 3 | 3 | 0.43 | 1 | 1.043 | 0.953 | 0.879 | 0.936 | 0.798 | 0.998 | 0.869 | Ribonuclease T2-A OS=Mus musculus GN=Rnaset2a PE=1 SV=1 |
| 578 | 1 | UniProt | n | F8WHL2 | 113 | 166254 | 5 | 5 | 4 | 4 | 0.11 | 1 | 0.997 | 1.031 | 0.847 | 0.836 | 0.926 | 1.009 | 0.869 | Coatomer subunit alpha OS=Mus musculus GN=Copa PE=1 SV=1 |
| 527 | 1 | UniProt | n | Q9CZ30 | 130 | 58574 | 8 | 8 | 3 | 3 | 0.33 | 1 | 1.033 | 1.032 | 0.82 | 0.881 | 0.903 | 1.022 | 0.867 | Obg-like ATPase 1 OS=Mus musculus GN=Ola1 PE=1 SV=1 |
| 952 | 1 | UniProt | n | Q9ER00 | 49 | 36744 | 2 | 2 | 1 | 1 | 0.12 | 1 | 1.01 | 1.043 | 0.991 | 0.861 | 0.764 | 1.018 | 0.867 | Syntaxin-12 OS=Mus musculus GN=Stx12 PE=1 SV=1 |
| 494 | 1 | UniProt | n | P61961 | 148 | 11287 | 3 | 3 | 2 | 2 | 1.07 | 1 | 1.055 | 0.954 | 0.8 | 0.935 | 0.868 | 1.002 | 0.866 | Ubiquitin-fold modifier 1 OS=Mus musculus GN=Ufm1 PE=1 SV=1 |
| 335 | 1 | UniProt | n | P70349 | 256 | 16902 | 10 | 10 | 4 | 4 | 2.43 | 1 | 1.017 | 1.044 | 0.888 | 0.872 | 0.837 | 1.020 | 0.865 | Histidine triad nucleotide-binding protein 1 OS=Mus musculus GN=Hint1 PE=1 SV=3 |
| 1203 | 1 | UniProt | n | B1AU25 | 27 | 78166 | 1 | 1 | 1 | 1 | 0.06 | 1 | 1.048 | 1.821 | 0.915 | 0.834 | 0.849 | 1.240 | 0.865 | Apoptosis-inducing factor 1, mitochondrial OS=Mus musculus GN=Aifm1 PE=1 SV=1 |
| 641 | 1 | UniProt | n | Q5SWN2 | 93 | 86172 | 3 | 3 | 3 | 3 | 0.16 | 1 | 0.958 | 1.079 | 0.799 | 0.906 | 0.892 | 1.011 | 0.864 | Replication protein A 70 kDa DNA-binding subunit OS=Mus musculus GN=Rpa1 PE=1 SV=2 |
| 204 | 1 | UniProt | n | Q3U1J4 | 440 | 144515 | 27 | 27 | 16 | 16 | 0.6 | 1 | 0.952 | 1.011 | 0.896 | 0.876 | 0.821 | 0.987 | 0.864 | DNA damage-binding protein 1 OS=Mus musculus GN=Ddb1 PE=1 SV=2 |
| 865 | 1 | UniProt | n | Q91ZW3 | 58 | 151048 | 1 | 1 | 1 | 1 | 0.03 | 1 | 1.032 | 1.162 | 0.752 | 0.766 | 1.116 | 1.062 | 0.863 | SWI/SNF-related matrix-associated actin-dependent regulator of chromatin subfamily A member 5 OS=Mus musculus GN=Smarca5 PE=1 SV=1 |
| 109 | 1 | UniProt | n | Q8CG14 | 951 | 89305 | 35 | 35 | 17 | 17 | 1.58 | 1 | 1.016 | 0.981 | 0.858 | 0.851 | 0.879 | 0.999 | 0.863 | Complement C1s-A subcomponent OS=Mus musculus GN=C1sa PE=2 SV=2 |
| 891 | 1 | UniProt | n | Q9JMG7 | 55 | 31285 | 2 | 2 | 1 | 1 | 0.14 | 1 | 0.977 | 1.124 | 0.755 | 0.973 | 0.873 | 1.032 | 0.862 | Hepatoma-derived growth factor-related protein 3 OS=Mus musculus GN=Hdgfl3 PE=1 SV=2 |
| 488 | 1 | UniProt | n | Q9DAK9 | 150 | 17168 | 10 | 10 | 4 | 4 | 2.35 | 1 | 0.913 | 0.961 | 0.925 | 0.854 | 0.809 | 0.957 | 0.861 | 14 kDa phosphohistidine phosphatase OS=Mus musculus GN=Phpt1 PE=1 SV=1 |
| 946 | 1 | UniProt | n | A0A0G2JD | 50 | 20711 | 1 | 1 | 1 | 1 | 0.22 | 1 | 0.9 | 1.018 | 0.877 | 0.808 | 0.898 | 0.971 | 0.860 | DCN1-like protein (Fragment) OS=Mus musculus GN=Dcun1d1 PE=1 SV=1 |
| 614 | 1 | UniProt | n | Q6PDQ2 | 101 | 272654 | 1 | 1 | 1 | 1 | 0.02 | 1 | 1.016 | 1.25 | 0.876 | 0.864 | 0.84 | 1.083 | 0.860 | Chromodomain-helicase-DNA-binding protein 4 OS=Mus musculus GN=Chd4 PE=1 SV=1 |
| 848 | 1 | UniProt | n | Q11136 | 60 | 61555 | 3 | 3 | 3 | 3 | 0.23 | 1 | 0.83 | 0.953 | 0.915 | 0.809 | 0.857 | 0.925 | 0.859 | Xaa-Pro dipeptidase OS=Mus musculus GN=Pepd PE=1 SV=3 |
| 929 | 1 | UniProt | n | Q9JHJ0 | 52 | 48392 | 1 | 1 | 1 | 1 | 0.09 | 1 | 0.972 | 1.123 | 0.811 | 0.917 | 0.853 | 1.030 | 0.859 | Tropomodulin-3 OS=Mus musculus GN=Tmod3 PE=1 SV=1 |
| 427 | 1 | UniProt | n | Q8BMS2 | 185 | 39656 | 10 | 10 | 6 | 6 | 0.89 | 1 | 0.965 | 0.926 | 0.864 | 0.832 | 0.882 | 0.963 | 0.859 | Spondin-2 OS=Mus musculus GN=Spon2 PE=2 SV=2 |
| 137 | 1 | UniProt | n | O88342 | 714 | 79998 | 36 | 36 | 17 | 17 | 1.88 | 1 | 1.008 | 1.045 | 0.921 | 0.845 | 0.814 | 1.017 | 0.859 | WD repeat-containing protein 1 OS=Mus musculus GN=Wdr1 PE=1 SV=3 |
| 1173 | 1 | UniProt | n | O35103 | 30 | 56866 | 2 | 2 | 1 | 1 | 0.08 | 1 | 1.493 | 1.615 | 1.069 | 0.745 | 0.795 | 1.341 | 0.859 | Osteomodulin OS=Mus musculus GN=Omd PE=2 SV=1 |
| 198 | 2 | UniProt | n | Q9DBG3 | 364 | 124113 | 21 | 21 | 11 | 11 | 0.56 | 1 | 1.011 | 1.088 | 0.972 | 0.803 | 0.811 | 1.032 | 0.859 | AP-2 complex subunit beta OS=Mus musculus GN=Ap2b1 PE=1 SV=1 |
| 105 | 2 | UniProt | n | Q61292 | 705 | 212847 | 27 | 27 | 18 | 18 | 0.43 | 1 | 0.929 | 1.028 | 0.893 | 0.848 | 0.834 | 0.985 | 0.858 | Laminin subunit beta-2 OS=Mus musculus GN=Lamb2 PE=1 SV=2 |
| 221 | 2 | UniProt | n | E9Q0M9 | 359 | 81131 | 24 | 24 | 9 | 9 | 0.6 | 1 | 0.924 | 0.95 | 0.921 | 0.871 | 0.786 | 0.957 | 0.857 | Caldesmon 1 OS=Mus musculus GN=Cald1 PE=1 SV=1 |
| 272 | 1 | UniProt | n | Q60864 | 325 | 82213 | 16 | 16 | 11 | 11 | 0.86 | 1 | 0.934 | 0.938 | 0.861 | 0.875 | 0.835 | 0.957 | 0.857 | Stress-induced-phosphoprotein 1 OS=Mus musculus GN=Stip1 PE=1 SV=1 |
| 395 | 3 | UniProt | n | B1AQY9 | 86 | 62645 | 5 | 5 | 4 | 4 | 0.31 | 1 | 0.824 | 0.927 | 0.907 | 0.96 | 0.722 | 0.914 | 0.857 | Septin-8 OS=Mus musculus GN=Sept8 PE=1 SV=1 |
| 1017 | 1 | UniProt | n | Q9D7S9 | 43 | 32211 | 1 | 1 | 1 | 1 | 0.14 | 1 | 0.911 | 0.878 | 0.86 | 0.771 | 0.948 | 0.928 | 0.857 | Charged multivesicular body protein 5 OS=Mus musculus GN=Chmp5 PE=1 SV=1 |
| 1216 | 1 | UniProt | n | Q5F226 | 26 | 528745 | 1 | 1 | 1 | 1 | 0.01 | 1 | 1.081 | 1.064 | 0.99 | 0.82 | 0.774 | 1.048 | 0.857 | Protocadherin Fat 2 OS=Mus musculus GN=Fat2 PE=1 SV=1 |
| 1067 | 1 | UniProt | n | A2ALF0 | 39 | 35858 | 1 | 1 | 1 | 1 | 0.12 | 1 | 1.048 | 0.904 | 0.972 | 0.862 | 0.747 | 0.982 | 0.855 | DnaJ homolog subfamily C member 8 OS=Mus musculus GN=Dnajc8 PE=1 SV=1 |
| 620 | 1 | UniProt | n | D3Z7R6 | 100 | 34433 | 3 | 3 | 2 | 2 | 0.28 | 1 | 1.091 | 1.048 | 0.907 | 0.803 | 0.858 | 1.046 | 0.855 | Calponin OS=Mus musculus GN=Cnn2 PE=1 SV=1 |
| 361 | 1 | UniProt | n | Q9CWS0 | 233 | 37463 | 9 | 9 | 6 | 6 | 0.96 | 1 | 0.961 | 0.974 | 0.836 | 0.86 | 0.868 | 0.978 | 0.855 | N(G),N(G)-dimethylarginine dimethylaminohydrolase 1 OS=Mus musculus GN=Ddah1 PE=1 SV=3 |
| 1110 | 1 | UniProt | n | A0A087W | 35 | 29657 | 2 | 2 | 2 | 2 | 0.33 | 1 | 0.909 | 0.95 | 0.904 | 0.92 | 0.748 | 0.952 | 0.854 | Protein CDV3 OS=Mus musculus GN=Cdv3 PE=1 SV=1 |
| 931 | 1 | UniProt | n | Q3TVI8 | 51 | 93301 | 1 | 1 | 1 | 1 | 0.05 | 1 | 1.15 | 1.187 | 0.946 | 0.947 | 0.694 | 1.109 | 0.853 | Pre-B-cell leukemia transcription factor-interacting protein 1 OS=Mus musculus GN=Pbxip1 PE=1 SV=2 |
| 1218 | 1 | UniProt | n | P62702 | 26 | 38281 | 1 | 1 | 1 | 1 | 0.12 | 1 | 0.918 | 1.077 | 0.769 | 0.883 | 0.913 | 0.996 | 0.853 | 40S ribosomal protein S4, X isoform OS=Mus musculus GN=Rps4x PE=1 SV=2 |
| 359 | 1 | UniProt | n | Q9ERK4 | 234 | 128976 | 8 | 8 | 5 | 5 | 0.18 | 1 | 0.947 | 0.991 | 0.832 | 0.839 | 0.887 | 0.979 | 0.852 | Exportin-2 OS=Mus musculus GN=Cse1l PE=1 SV=1 |
| 363 | 1 | UniProt | n | P11983 | 232 | 73251 | 11 | 11 | 8 | 8 | 0.68 | 1 | 0.979 | 1.024 | 0.837 | 0.846 | 0.873 | 1.001 | 0.852 | T-complex protein 1 subunit alpha OS=Mus musculus GN=Tcp1 PE=1 SV=3 |
| 896 | 1 | UniProt | n | Q00519 | 55 | 173114 | 2 | 2 | 2 | 2 | 0.05 | 1 | 0.973 | 1.01 | 0.838 | 0.914 | 0.807 | 0.994 | 0.852 | Xanthine dehydrogenase/oxidase OS=Mus musculus GN=Xdh PE=1 SV=5 |
| 332 | 1 | UniProt | n | Q9CR16 | 261 | 53511 | 19 | 19 | 12 | 12 | 1.79 | 1 | 1.049 | 1.048 | 0.882 | 0.84 | 0.834 | 1.032 | 0.852 | Peptidyl-prolyl cis-trans isomerase D OS=Mus musculus GN=Ppid PE=1 SV=3 |
| 868 | 1 | UniProt | n | A0A1W2P | 58 | 21462 | 5 | 5 | 3 | 3 | 0.79 | 1 | 1.053 | 0.986 | 0.92 | 0.843 | 0.795 | 1.013 | 0.851 | 40S ribosomal protein S12 OS=Mus musculus GN=Rps12 PE=1 SV=1 |
| 1089 | 1 | UniProt | n | Q91VH2 | 37 | 81123 | 2 | 2 | 1 | 1 | 0.11 | 1 | 0.817 | 0.891 | 0.968 | 0.938 | 0.679 | 0.900 | 0.851 | Sorting nexin-9 OS=Mus musculus GN=Snx9 PE=1 SV=1 |
| 119 | 1 | UniProt | n | P02463 | 840 | 188877 | 24 | 24 | 9 | 9 | 0.25 | 1 | 0.971 | 0.944 | 0.873 | 0.832 | 0.848 | 0.971 | 0.851 | Collagen alpha-1(IV) chain OS=Mus musculus GN=Col4a1 PE=1 SV=4 |
| 668 | 1 | UniProt | n | Q9Z204 | 89 | 43536 | 2 | 2 | 2 | 2 | 0.21 | 1 | 0.844 | 0.907 | 0.874 | 0.878 | 0.799 | 0.915 | 0.850 | Heterogeneous nuclear ribonucleoproteins C1/C2 OS=Mus musculus GN=Hnrnpc PE=1 SV=1 |
| 1193 | 1 | UniProt | n | Q7TNV0 | 29 | 64306 | 1 | 1 | 1 | 1 | 0.07 | 1 | 1.058 | 1 | 0.824 | 0.85 | 0.875 | 1.019 | 0.849 | Protein DEK OS=Mus musculus GN=Dek PE=1 SV=1 |
| 1246 | 1 | UniProt | n | Q8R1T1-2 | 23 | 48520 | 1 | 1 | 1 | 1 | 0.09 | 1 | 1.067 | 1.17 | 0.922 | 0.848 | 0.783 | 1.077 | 0.849 | Isoform 2 of Charged multivesicular body protein 7 OS=Mus musculus GN=Chmp7 |
| 524 | 1 | UniProt | n | P46935 | 131 | 118266 | 5 | 5 | 2 | 2 | 0.07 | 1 | 1.017 | 1.089 | 0.859 | 0.922 | 0.771 | 1.035 | 0.848 | E3 ubiquitin-protein ligase NEDD4 OS=Mus musculus GN=Nedd4 PE=1 SV=3 |
| 665 | 1 | UniProt | n | Q9D832 | 89 | 47935 | 4 | 4 | 2 | 2 | 0.19 | 1 | 1.029 | 1.089 | 0.839 | 0.929 | 0.783 | 1.039 | 0.848 | DnaJ homolog subfamily B member 4 OS=Mus musculus GN=Dnajb4 PE=1 SV=1 |
| 211 | 1 | UniProt | n | P27546 | 414 | 145804 | 16 | 16 | 10 | 10 | 0.34 | 1 | 0.997 | 1.022 | 0.888 | 0.837 | 0.821 | 1.006 | 0.848 | Microtubule-associated protein 4 OS=Mus musculus GN=Map4 PE=1 SV=3 |
| 242 | 1 | UniProt | n | A0A0R4J0 | 359 | 93287 | 17 | 17 | 10 | 10 | 0.65 | 1 | 0.994 | 1.041 | 0.871 | 0.848 | 0.826 | 1.011 | 0.848 | Protein disulfide-isomerase A4 OS=Mus musculus GN=Pdia4 PE=1 SV=1 |
| 1180 | 1 | UniProt | n | Q8BU30 | 30 | 170532 | 1 | 1 | 1 | 1 | 0.03 | 1 | 0.969 | 1.111 | 0.92 | 0.76 | 0.872 | 1.025 | 0.848 | Isoleucine--tRNA ligase, cytoplasmic OS=Mus musculus GN=Iars PE=1 SV=2 |
| 1082 | 1 | UniProt | n | A0A0J9YT | 38 | 17627 | 1 | 1 | 1 | 1 | 0.27 | 1 | 0.971 | 0.91 | 1.062 | 0.807 | 0.709 | 0.960 | 0.847 | Ras GTPase-activating protein-binding protein 2 (Fragment) OS=Mus musculus GN=G3bp2 PE=1 SV=1 |
| 693 | 1 | UniProt | n | Q9WV32 | 84 | 49895 | 2 | 2 | 2 | 2 | 0.18 | 1 | 1.028 | 1.1 | 0.892 | 0.86 | 0.791 | 1.042 | 0.847 | Actin-related protein 2/3 complex subunit 1B OS=Mus musculus GN=Arpc1b PE=1 SV=4 |
| 221 | 1 | UniProt | n | E9QA15 | 393 | 116341 | 24 | 24 | 9 | 9 | 0.39 | 1 | 0.91 | 0.937 | 0.91 | 0.859 | 0.776 | 0.948 | 0.847 | Caldesmon 1 OS=Mus musculus GN=Cald1 PE=1 SV=1 |
| 1046 | 1 | UniProt | n | P05533 | 41 | 16744 | 1 | 1 | 1 | 1 | 0.28 | 1 | 0.812 | 0.956 | 0.776 | 0.891 | 0.877 | 0.919 | 0.846 | Lymphocyte antigen 6A-2/6E-1 OS=Mus musculus GN=Ly6a PE=1 SV=1 |
| 694 | 1 | UniProt | n | Q8BH80 | 84 | 33777 | 3 | 3 | 2 | 2 | 0.28 | 1 | 1.128 | 1.23 | 0.805 | 0.881 | 0.855 | 1.115 | 0.846 | Vesicle-associated membrane protein, associated protein B and C OS=Mus musculus GN=Vapb PE=1 SV=1 |
| 912 | 1 | UniProt | n | H7BXB9 | 54 | 18943 | 1 | 1 | 1 | 1 | 0.25 | 1 | 1.177 | 1.189 | 0.705 | 0.942 | 0.912 | 1.119 | 0.846 | RIKEN cDNA 1700037H04 gene (Fragment) OS=Mus musculus GN=1700037H04Rik PE=1 SV=1 |
| 500 | 1 | UniProt | n | Q99KQ4 | 146 | 69940 | 5 | 5 | 4 | 4 | 0.27 | 1 | 0.978 | 1.065 | 0.882 | 0.863 | 0.795 | 1.014 | 0.846 | Nicotinamide phosphoribosyltransferase OS=Mus musculus GN=Nampt PE=1 SV=1 |
| 811 | 1 | UniProt | n | Q9R1T2 | 64 | 47785 | 3 | 3 | 3 | 3 | 0.3 | 1 | 0.998 | 0.886 | 0.886 | 0.842 | 0.811 | 0.960 | 0.846 | SUMO-activating enzyme subunit 1 OS=Mus musculus GN=Sae1 PE=1 SV=1 |
| 945 | 1 | UniProt | n | Q8VDN2 | 50 | 131307 | 2 | 2 | 2 | 2 | 0.07 | 1 | 1.03 | 1.146 | 0.847 | 0.83 | 0.859 | 1.057 | 0.845 | Sodium/potassium-transporting ATPase subunit alpha-1 OS=Mus musculus GN=Atp1a1 PE=1 SV=1 |
| 833 | 1 | UniProt | n | A2BFF5 | 61 | 84046 | 2 | 2 | 1 | 1 | 0.05 | 1 | 0.807 | 0.861 | 0.926 | 0.807 | 0.808 | 0.886 | 0.845 | Cytoplasmic dynein 1 intermediate chain 2 OS=Mus musculus GN=Dync1i2 PE=1 SV=1 |
| 555 | 1 | UniProt | n | E9Q600 | 120 | 94980 | 3 | 3 | 3 | 3 | 0.14 | 1 | 0.831 | 0.922 | 0.859 | 0.807 | 0.871 | 0.915 | 0.845 | Lysyl oxidase homolog 4 OS=Mus musculus GN=Loxl4 PE=1 SV=1 |
| 1115 | 1 | UniProt | n | Q3TIR6 | 34 | 19953 | 1 | 1 | 1 | 1 | 0.23 | 1 | 0.903 | 0.977 | 0.726 | 1.056 | 0.787 | 0.959 | 0.845 | Prefoldin subunit 3 OS=Mus musculus GN=Vbp1 PE=1 SV=1 |
| 545 | 1 | UniProt | n | A0A0A6Y) | 124 | 23491 | 2 | 2 | 2 | 2 | 0.43 | 1 | 0.839 | 0.929 | 0.898 | 0.831 | 0.808 | 0.920 | 0.845 | Transforming protein RhoA (Fragment) OS=Mus musculus GN=Rhoa PE=1 SV=1 |
| 928 | 1 | UniProt | n | A2ATI6 | 52 | 40826 | 1 | 1 | 1 | 1 | 0.11 | 1 | 1.125 | 0.929 | 0.643 | 0.99 | 0.947 | 1.015 | 0.845 | Golgi reassembly stacking protein 2, isoform CRA c OS=Mus musculus GN=Gorasp2 PE=1 SV=1 |
| 1272 | 1 | UniProt | n | Q921F4 | 19 | 75421 | 1 | 1 | 1 | 1 | 0.06 | 1 | 0.832 | 0.764 | 0.989 | 0.768 | 0.793 | 0.860 | 0.845 | Heterogeneous nuclear ribonucleoprotein L-like OS=Mus musculus GN=Hnrnpll PE=1 SV=3 |
| 1084 | 1 | UniProt | n | M0QWS4 | 37 | 13519 | 2 | 2 | 1 | 1 | 0.36 | 1 | 0.986 | 1.05 | 0.868 | 0.964 | 0.719 | 1.012 | 0.844 | Ubiquitin-fold modifier conjugating enzyme 1, isoform CRA c OS=Mus musculus GN=Ufc1 PE=1 SV=1 |
| 200 | 1 | UniProt | n | A0A0A0M | 450 | 20127 | 21 | 21 | 6 | 6 | 5.48 | 1 | 1.058 | 1.021 | 0.844 | 0.838 | 0.849 | 1.026 | 0.844 | Eukaryotic translation initiation factor 5A (Fragment) OS=Mus musculus GN=Eif5a PE=1 SV=1 |
| 885 | 1 | UniProt | n | Q80ZP8 | 56 | 26318 | 2 | 2 | 2 | 2 | 0.38 | 1 | 0.98 | 0.954 | 0.902 | 0.901 | 0.738 | 0.978 | 0.843 | Armet protein OS=Mus musculus GN=Manf PE=1 SV=1 |
| 717 | 1 | UniProt | n | O88839 | 78 | 99678 | 3 | 3 | 3 | 3 | 0.14 | 1 | 0.901 | 0.836 | 0.833 | 0.848 | 0.848 | 0.910 | 0.843 | Disintegrin and metalloproteinase domain-containing protein 15 OS=Mus musculus GN=Adam15 PE=1 SV=2 |
| 356 | 1 | UniProt | n | P80314 | 238 | 69277 | 10 | 10 | 7 | 7 | 0.63 | 1 | 0.823 | 0.921 | 0.869 | 0.844 | 0.816 | 0.912 | 0.843 | T-complex protein 1 subunit beta OS=Mus musculus GN=Cct2 PE=1 SV=4 |
| 1263 | 1 | UniProt | n | Q91XC8 | 20 | 14798 | 1 | 1 | 1 | 1 | 0.32 | 1 | 1.042 | 0.973 | 0.905 | 0.865 | 0.764 | 1.005 | 0.843 | Death-associated protein 1 OS=Mus musculus GN=Dap PE=1 SV=3 |
| 577 | 1 | UniProt | n | P70677 | 113 | 39123 | 3 | 3 | 3 | 3 | 0.38 | 1 | 1.006 | 1.041 | 0.887 | 0.818 | 0.824 | 1.016 | 0.842 | Caspase-3 OS=Mus musculus GN=Casp3 PE=1 SV=1 |
| 667 | 1 | UniProt | n | Q3UJQ9 | 89 | 63777 | 3 | 3 | 2 | 2 | 0.14 | 1 | 1.069 | 1.147 | 0.83 | 0.817 | 0.881 | 1.070 | 0.842 | Succinyl-CoA:3-ketoacid-coenzyme A transferase OS=Mus musculus GN=Oxct1 PE=1 SV=1 |
| 103 | 1 | UniProt | n | A0A140T8 | 1006 | 34214 | 35 | 35 | 11 | 11 | 6.16 | 1 | 1.035 | 1.037 | 0.894 | 0.834 | 0.799 | 1.024 | 0.841 | Proliferating cell nuclear antigen OS=Mus musculus GN=Pcna-ps2 PE=3 SV=1 |
| 731 | 1 | UniProt | n | Q9D662 | 77 | 96548 | 4 | 4 | 4 | 4 | 0.19 | 1 | 0.929 | 0.988 | 0.833 | 0.873 | 0.819 | 0.972 | 0.841 | Protein transport protein Sec23B OS=Mus musculus GN=Sec23b PE=1 SV=1 |
| 889 | 1 | UniProt | n | P97351 | 55 | 41914 | 1 | 1 | 1 | 1 | 0.11 | 1 | 1.129 | 0.875 | 0.761 | 0.864 | 0.902 | 0.996 | 0.840 | 40S ribosomal protein S3a OS=Mus musculus GN=Rps3a PE=1 SV=3 |
| 587 | 1 | UniProt | n | P11438 | 108 | 50593 | 3 | 3 | 1 | 1 | 0.09 | 1 | 1.175 | 1.203 | 0.849 | 0.885 | 0.789 | 1.122 | 0.840 | Lysosome-associated membrane glycoprotein 1 OS=Mus musculus GN=Lamp1 PE=1 SV=2 |
| 395 | 2 | UniProt | n | A0A0J9YT | 111 | 64388 | 7 | 7 | 6 | 6 | 0.48 | 1 | 0.893 | 0.993 | 0.934 | 0.909 | 0.698 | 0.961 | 0.840 | Septin-11 OS=Mus musculus GN=Sept11 PE=1 SV=1 |
| 927 | 1 | UniProt | n | D3Z3T5 | 52 | 33973 | 1 | 1 | 1 | 1 | 0.13 | 1 | 1.105 | 1.259 | 1.064 | 0.652 | 0.854 | 1.116 | 0.840 | Choline-phosphate cytidylyltransferase A OS=Mus musculus GN=Pcyt1a PE=1 SV=1 |
| 560 | 1 | UniProt | n | Q76MZ3 | 119 | 75964 | 6 | 6 | 4 | 4 | 0.25 | 1 | 0.95 | 1.064 | 0.897 | 0.822 | 0.802 | 1.004 | 0.839 | Serine/threonine-protein phosphatase 2A 65 kDa regulatory subunit A alpha isoform OS=Mus musculus GN=Ppp2r1a PE=1 SV=3 |
| 376 | 1 | UniProt | n | P97310 | 226 | 116254 | 8 | 8 | 6 | 6 | 0.24 | 1 | 1 | 1.053 | 0.832 | 0.83 | 0.855 | 1.017 | 0.839 | DNA replication licensing factor MCM2 OS=Mus musculus GN=Mcm2 PE=1 SV=3 |
| 322 | 1 | UniProt | n | P06801 | 269 | 75979 | 12 | 12 | 9 | 9 | 0.75 | 1 | 1.018 | 1.018 | 0.854 | 0.866 | 0.798 | 1.012 | 0.839 | NADP-dependent malic enzyme OS=Mus musculus GN=Me1 PE=1 SV=2 |
| 197 | 1 | UniProt | n | Q9QYB1 | 453 | 36500 | 18 | 18 | 9 | 9 | 2.17 | 1 | 1.074 | 1.078 | 0.866 | 0.832 | 0.819 | 1.050 | 0.839 | Chloride intracellular channel protein 4 OS=Mus musculus GN=Clic4 PE=1 SV=3 |
| 22 | 1 | UniProt | n | P17182 | 4012 | 58034 | 124 | 124 | 20 | 20 | 7.26 | 1 | 1.107 | 1.08 | 0.86 | 0.793 | 0.864 | 1.061 | 0.838 | Alpha-enolase OS=Mus musculus GN=Eno1 PE=1 SV=3 |
| 1187 | 1 | UniProt | n | Q8R086 | 29 | 67216 | 1 | 1 | 1 | 1 | 0.06 | 1 | 0.929 | 1.13 | 1.091 | 0.581 | 0.929 | 1.016 | 0.838 | Sulfite oxidase, mitochondrial OS=Mus musculus GN=Suox PE=1 SV=2 |
| 355 | 1 | UniProt | n | Q9WTP6 | 238 | 33070 | 11 | 11 | 5 | 5 | 0.89 | 1 | 0.894 | 0.892 | 0.892 | 0.833 | 0.791 | 0.927 | 0.838 | Adenylate kinase 2, mitochondrial OS=Mus musculus GN=Ak2 PE=1 SV=5 |
| 734 | 1 | UniProt | n | O70310 | 76 | 70209 | 6 | 6 | 3 | 3 | 0.2 | 1 | 0.93 | 0.886 | 0.835 | 0.913 | 0.77 | 0.938 | 0.837 | Glycylpeptide N-tetradecanoyltransferase 1 OS=Mus musculus GN=Nmt1 PE=1 SV=1 |
| 580 | 1 | UniProt | n | Q9D1J1 | 112 | 33540 | 2 | 2 | 2 | 2 | 0.29 | 1 | 1.035 | 1.059 | 0.84 | 0.856 | 0.815 | 1.031 | 0.837 | Adaptin ear-binding coat-associated protein 2 OS=Mus musculus GN=Necap2 PE=1 SV=1 |
| 539 | 1 | UniProt | n | Q9EQH3 | 125 | 106292 | 6 | 6 | 4 | 4 | 0.17 | 1 | 0.963 | 1.128 | 0.901 | 0.846 | 0.768 | 1.028 | 0.837 | Vacuolar protein sorting-associated protein 35 OS=Mus musculus GN=Vps35 PE=1 SV=1 |
| 235 | 1 | UniProt | n | Q8VHM5 | 371 | 84763 | 14 | 14 | 10 | 10 | 0.65 | 1 | 0.969 | 0.978 | 0.844 | 0.852 | 0.814 | 0.982 | 0.837 | Heterogeneous nuclear ribonucleoprotein R OS=Mus musculus GN=Hnrnpr PE=1 SV=1 |
| 909 | 1 | UniProt | n | Q8BTS4 | 54 | 64916 | 1 | 1 | 1 | 1 | 0.07 | 1 | 0.877 | 1.15 | 0.849 | 0.819 | 0.841 | 1.003 | 0.836 | Nuclear pore complex protein Nup54 OS=Mus musculus GN=Nup54 PE=1 SV=1 |
| 581 | 1 | UniProt | n | Q3UM45 | 112 | 49268 | 5 | 5 | 4 | 4 | 0.41 | 1 | 0.984 | 1.02 | 0.774 | 0.942 | 0.802 | 1.001 | 0.836 | Protein phosphatase 1 regulatory subunit 7 OS=Mus musculus GN=Ppp1r7 PE=1 SV=2 |
| 1096 | 1 | UniProt | n | A2CES4 | 36 | 22388 | 1 | 1 | 1 | 1 | 0.2 | 1 | 1.144 | 1.066 | 0.765 | 0.792 | 0.961 | 1.068 | 0.835 | U2 small nuclear ribonucleoprotein B'' (Fragment) OS=Mus musculus GN=Snrpb2 PE=1 SV=1 |

| | | | | | | | | | | | | | | | | | | |
|---|---|---|---|---|---|---|---|---|---|---|---|---|---|---|---|---|---|---|
| 1135 | 1 | UniProt n | Q9CQ45 | 33 | 22239 | 1 | 1 | 1 | 1 | 0.21 | 1 | 0.936 | 0.905 | 0.932 | 0.882 | 0.708 | 0.946 | 0.835 | Neudesin OS=Mus musculus GN=Nenf PE=1 SV=1 |
| 439 | 1 | UniProt n | G5E839 | 177 | 66801 | 11 | 11 | 6 | 6 | 0.46 | 1 | 1.048 | 1.071 | 0.822 | 0.875 | 0.809 | 1.039 | 0.835 | T-complex protein 1 subunit delta OS=Mus musculus GN=Cct4 PE=1 SV=1 |
| 612 | 1 | UniProt n | Q9JMH6 | 101 | 80176 | 2 | 2 | 2 | 2 | 0.11 | 1 | 0.816 | 0.859 | 0.803 | 0.888 | 0.816 | 0.888 | 0.835 | Thioredoxin reductase 1, cytoplasmic OS=Mus musculus GN=Txnrd1 PE=1 SV=3 |
| 201 | 1 | UniProt n | P99029 | 449 | 27027 | 15 | 15 | 8 | 8 | 2.47 | 1 | 1.05 | 0.949 | 0.809 | 0.849 | 0.847 | 0.999 | 0.835 | Peroxiredoxin-5, mitochondrial OS=Mus musculus GN=Prdx5 PE=1 SV=2 |
| 316 | 1 | UniProt n | Q99JY9 | 274 | 54996 | 12 | 12 | 6 | 6 | 0.85 | 1 | 0.952 | 1.009 | 0.847 | 0.813 | 0.844 | 0.987 | 0.835 | Actin-related protein 3 OS=Mus musculus GN=Actr3 PE=1 SV=3 |
| 174 | 1 | UniProt n | Q9JKF1 | 542 | 229423 | 17 | 17 | 11 | 11 | 0.25 | 1 | 0.931 | 1.014 | 0.849 | 0.848 | 0.806 | 0.981 | 0.834 | Ras GTPase-activating-like protein IQGAP1 OS=Mus musculus GN=Iqgap1 PE=1 SV=2 |
| 1044 | 1 | UniProt n | P14131 | 41 | 21699 | 1 | 1 | 1 | 1 | 0.21 | 1 | 0.943 | 1.081 | 0.713 | 0.808 | 1.006 | 1.006 | 0.834 | 40S ribosomal protein S16 OS=Mus musculus GN=Rps16 PE=1 SV=4 |
| 653 | 1 | UniProt n | A0A140T8 | 90 | 30803 | 4 | 4 | 3 | 3 | 0.5 | 1 | 1.103 | 1.059 | 0.767 | 0.822 | 0.918 | 1.053 | 0.833 | Ribosomal protein S2, pseudogene 6 OS=Mus musculus GN=Rps2-ps6 PE=3 SV=1 |
| 690 | 1 | UniProt n | Q8VDJ3 | 84 | 175882 | 4 | 4 | 3 | 3 | 0.08 | 1 | 1.026 | 1.105 | 0.907 | 0.833 | 0.766 | 1.043 | 0.833 | Vigilin OS=Mus musculus GN=Hdlbp PE=1 SV=1 |
| 469 | 1 | UniProt n | A0A171K | 161 | 44913 | 6 | 6 | 4 | 4 | 0.46 | 1 | 1.035 | 1.13 | 0.842 | 0.802 | 0.857 | 1.054 | 0.833 | Protein arginine N-methyltransferase 1 OS=Mus musculus GN=Prmt1 PE=1 SV=1 |
| 922 | 1 | UniProt n | Q9D358-2 | 53 | 22252 | 4 | 4 | 2 | 2 | 0.46 | 1 | 0.875 | 0.848 | 0.787 | 0.851 | 0.863 | 0.905 | 0.833 | Isoform 2 of Low molecular weight phosphotyrosine protein phosphatase OS=Mus musculus GN=Acp1 |
| 904 | 1 | UniProt n | Q99KR3 | 54 | 38439 | 2 | 2 | 1 | 1 | 0.12 | 1 | 0.99 | 0.99 | 0.848 | 0.871 | 0.782 | 0.993 | 0.833 | Endoribonuclease LACTB2 OS=Mus musculus GN=Lactb2 PE=1 SV=1 |
| 314 | 1 | UniProt n | B1AU75 | 275 | 103965 | 12 | 12 | 8 | 8 | 0.44 | 1 | 1.002 | 1.054 | 0.846 | 0.856 | 0.796 | 1.018 | 0.832 | Nuclear autoantigenic sperm protein OS=Mus musculus GN=Nasp PE=1 SV=1 |
| 497 | 1 | UniProt n | Q9D1E6 | 146 | 32466 | 4 | 4 | 3 | 3 | 0.48 | 1 | 0.945 | 0.943 | 0.9 | 0.842 | 0.76 | 0.962 | 0.832 | Tubulin-folding cofactor B OS=Mus musculus GN=Tbcb PE=1 SV=2 |
| 414 | 1 | UniProt n | Q922W6 | 200 | 65556 | 11 | 11 | 8 | 8 | 0.68 | 1 | 0.927 | 0.957 | 0.819 | 0.881 | 0.798 | 0.961 | 0.832 | Matrix metalloproteinase OS=Mus musculus GN=Mmp3 PE=1 SV=1 |
| 147 | 1 | UniProt n | P10639 | 680 | 15898 | 30 | 30 | 6 | 6 | 5.23 | 1 | 0.992 | 0.898 | 0.799 | 0.913 | 0.788 | 0.962 | 0.831 | Thioredoxin OS=Mus musculus GN=Txn PE=1 SV=3 |
| 1021 | 1 | UniProt n | Q8BMJ3 | 43 | 21714 | 1 | 1 | 1 | 1 | 0.21 | 1 | 0.917 | 0.922 | 0.854 | 0.938 | 0.716 | 0.946 | 0.831 | Eukaryotic translation initiation factor 1A, X-chromosomal OS=Mus musculus GN=Eif1ax PE=2 SV=3 |
| 52 | 1 | UniProt n | F8VQJ3 | 1821 | 206617 | 62 | 62 | 32 | 32 | 1.1 | 1 | 0.988 | 1.034 | 0.848 | 0.785 | 0.861 | 1.007 | 0.831 | Laminin subunit gamma-1 OS=Mus musculus GN=Lamc1 PE=1 SV=1 |
| 328 | 1 | UniProt n | Q99KP6 | 264 | 65003 | 9 | 9 | 5 | 5 | 0.38 | 1 | 1.017 | 1.05 | 0.855 | 0.808 | 0.829 | 1.022 | 0.830 | Pre-mRNA-processing factor 19 OS=Mus musculus GN=Prpf19 PE=1 SV=1 |
| 726 | 1 | UniProt n | P63276 | 77 | 19819 | 1 | 1 | 1 | 1 | 0.23 | 1 | 1.035 | 1.119 | 0.827 | 0.835 | 0.829 | 1.050 | 0.830 | 40S ribosomal protein S17 OS=Mus musculus GN=Rps17 PE=1 SV=2 |
| 888 | 1 | UniProt n | Q922B2 | 56 | 66909 | 2 | 2 | 2 | 2 | 0.13 | 1 | 0.97 | 1.073 | 0.828 | 0.806 | 0.857 | 1.013 | 0.830 | Aspartate--tRNA ligase, cytoplasmic OS=Mus musculus GN=Dars PE=1 SV=2 |
| 846 | 1 | UniProt n | E9Q390 | 60 | 278879 | 4 | 4 | 3 | 3 | 0.05 | 1 | 1.012 | 0.994 | 0.85 | 0.844 | 0.797 | 1.002 | 0.830 | Myoferlin OS=Mus musculus GN=Myof PE=1 SV=2 |
| 932 | 1 | UniProt n | P60487 | 51 | 32727 | 1 | 1 | 1 | 1 | 0.14 | 1 | 0.924 | 0.813 | 0.928 | 0.805 | 0.765 | 0.909 | 0.830 | Pyridoxal phosphate phosphatase OS=Mus musculus GN=Pdxp PE=1 SV=1 |
| 601 | 1 | UniProt n | Q8BYK6 | 103 | 75269 | 2 | 2 | 1 | 1 | 0.06 | 1 | 0.848 | 0.984 | 0.85 | 0.9 | 0.747 | 0.941 | 0.830 | YTH domain-containing family protein 3 OS=Mus musculus GN=Ythdf3 PE=1 SV=2 |
| 79 | 1 | UniProt n | Q9R118 | 1221 | 61348 | 42 | 42 | 17 | 17 | 3.25 | 1 | 1.039 | 1.007 | 0.856 | 0.81 | 0.824 | 1.015 | 0.830 | Serine protease HTRA1 OS=Mus musculus GN=Htra1 PE=1 SV=2 |
| 853 | 1 | UniProt n | Q3THK3 | 60 | 76113 | 1 | 1 | 1 | 1 | 0.06 | 1 | 0.931 | 0.964 | 0.741 | 0.804 | 0.958 | 0.965 | 0.829 | General transcription factor IIF subunit 1 OS=Mus musculus GN=Gtf2f1 PE=1 SV=2 |
| 490 | 1 | UniProt n | Q9D0R2 | 150 | 103508 | 10 | 10 | 7 | 7 | 0.33 | 1 | 0.929 | 0.945 | 0.837 | 0.817 | 0.834 | 0.958 | 0.829 | Threonine--tRNA ligase, cytoplasmic OS=Mus musculus GN=Tars PE=1 SV=2 |
| 392 | 1 | UniProt n | Q64433 | 211 | 14606 | 11 | 11 | 4 | 4 | 2.1 | 1 | 1.075 | 1.062 | 0.844 | 0.846 | 0.798 | 1.045 | 0.829 | 10 kDa heat shock protein, mitochondrial OS=Mus musculus GN=Hspe1 PE=1 SV=2 |
| 136 | 1 | UniProt n | Q9WVA4 | 720 | 26778 | 29 | 29 | 10 | 10 | 6.67 | 1 | 1.052 | 0.978 | 0.858 | 0.825 | 0.804 | 1.010 | 0.829 | Transgelin-2 OS=Mus musculus GN=Tagln2 PE=1 SV=4 |
| 236 | 1 | UniProt n | P42208 | 370 | 49897 | 15 | 15 | 8 | 8 | 0.97 | 1 | 0.92 | 0.965 | 0.783 | 0.889 | 0.817 | 0.961 | 0.829 | Septin-2 OS=Mus musculus GN=Sept2 PE=1 SV=2 |
| 276 | 1 | UniProt n | P29341 | 323 | 83586 | 18 | 18 | 12 | 12 | 1.03 | 1 | 1.002 | 0.935 | 0.84 | 0.8 | 0.846 | 0.978 | 0.828 | Polyadenylate-binding protein 1 OS=Mus musculus GN=Pabpc1 PE=1 SV=2 |
| 1080 | 1 | UniProt n | A0A0R4J1 | 38 | 46008 | 1 | 1 | 1 | 1 | 0.1 | 1 | 0.941 | 1.016 | 0.803 | 0.817 | 0.866 | 0.985 | 0.828 | Regulation of nuclear pre-mRNA domain-containing protein 1B OS=Mus musculus GN=Rprd1b PE=1 SV=1 |
| 916 | 1 | UniProt n | Q8QZY1 | 53 | 78314 | 5 | 5 | 4 | 4 | 0.24 | 1 | 0.808 | 0.847 | 0.84 | 0.857 | 0.788 | 0.881 | 0.828 | Eukaryotic translation initiation factor 3 subunit L OS=Mus musculus GN=Eif3l PE=1 SV=1 |
| 803 | 1 | UniProt n | Q6NZM8 | 65 | 80304 | 2 | 2 | 2 | 2 | 0.11 | 1 | 0.925 | 0.961 | 0.8 | 0.871 | 0.814 | 0.962 | 0.828 | 3'-phosphoadenosine 5'-phosphosulfate synthase 1, isoform CRA a OS=Mus musculus GN=Papss1 PE=1 SV=1 |
| 934 | 1 | UniProt n | P63147 | 51 | 19477 | 2 | 2 | 2 | 2 | 0.54 | 1 | 0.905 | 0.937 | 0.866 | 0.844 | 0.776 | 0.947 | 0.828 | Ubiquitin-conjugating enzyme E2 B OS=Mus musculus GN=Ube2b PE=1 SV=1 |
| 744 | 1 | UniProt n | Q61166 | 74 | 37436 | 4 | 4 | 3 | 3 | 0.4 | 1 | 0.991 | 1.021 | 0.808 | 0.873 | 0.804 | 1.004 | 0.828 | Microtubule-associated protein RP/EB family member 1 OS=Mus musculus GN=Mapre1 PE=1 SV=3 |
| 298 | 1 | UniProt n | Q9CPW4 | 293 | 20887 | 8 | 8 | 3 | 3 | 1.72 | 1 | 1.032 | 1.015 | 0.834 | 0.817 | 0.832 | 1.016 | 0.828 | Actin-related protein 2/3 complex subunit 5 OS=Mus musculus GN=Arpc5 PE=1 SV=3 |
| 575 | 1 | UniProt n | Q91YR9 | 114 | 44331 | 5 | 5 | 4 | 4 | 0.46 | 1 | 0.861 | 0.942 | 0.823 | 0.899 | 0.766 | 0.933 | 0.828 | Prostaglandin reductase 1 OS=Mus musculus GN=Ptgr1 PE=1 SV=2 |
| 626 | 1 | UniProt n | G3UXT7 | 97 | 17834 | 6 | 6 | 4 | 4 | 1.54 | 1 | 0.961 | 0.983 | 0.852 | 0.808 | 0.823 | 0.981 | 0.827 | RNA-binding protein FUS (Fragment) OS=Mus musculus GN=Fus PE=1 SV=1 |
| 780 | 1 | UniProt n | Q8K1J6 | 68 | 61562 | 3 | 3 | 2 | 2 | 0.15 | 1 | 1.093 | 1.183 | 0.863 | 0.788 | 0.833 | 1.089 | 0.827 | CCA tRNA nucleotidyltransferase 1, mitochondrial OS=Mus musculus GN=Trnt1 PE=1 SV=1 |
| 105 | 1 | UniProt n | E9QN70 | 981 | 237095 | 38 | 38 | 23 | 23 | 0.57 | 1 | 0.974 | 1.037 | 0.879 | 0.81 | 0.795 | 1.003 | 0.827 | Laminin subunit beta-1 OS=Mus musculus GN=Lamb1 PE=1 SV=1 |
| 283 | 1 | UniProt n | Q3UE92 | 313 | 86320 | 11 | 11 | 8 | 8 | 0.48 | 1 | 1.022 | 1.042 | 0.795 | 0.798 | 0.891 | 1.021 | 0.827 | X-prolyl aminopeptidase (Aminopeptidase P) 1, soluble, isoform CRA b OS=Mus musculus GN=Xpnpep1 PE=1 SV=1 |
| 711 | 1 | UniProt n | B2KFA8 | 80 | 29785 | 4 | 4 | 2 | 2 | 0.33 | 1 | 0.963 | 0.995 | 0.765 | 0.83 | 0.89 | 0.986 | 0.827 | Branched-chain-amino-acid aminotransferase, cytosolic (Fragment) OS=Mus musculus GN=Bcat1 PE=1 SV=8 |
| 798 | 1 | UniProt n | Q64669 | 65 | 37983 | 3 | 3 | 2 | 2 | 0.25 | 1 | 0.986 | 0.987 | 0.806 | 0.84 | 0.834 | 0.991 | 0.827 | NAD(P)H dehydrogenase [quinone] 1 OS=Mus musculus GN=Nqo1 PE=1 SV=3 |
| 397 | 1 | UniProt n | P80317 | 209 | 72937 | 9 | 9 | 7 | 7 | 0.5 | 1 | 0.991 | 0.955 | 0.83 | 0.832 | 0.817 | 0.982 | 0.826 | T-complex protein 1 subunit zeta OS=Mus musculus GN=Cct6a PE=1 SV=3 |
| 605 | 1 | UniProt n | Q8BH58 | 102 | 37151 | 7 | 7 | 4 | 4 | 0.57 | 1 | 0.945 | 0.972 | 0.87 | 0.824 | 0.787 | 0.972 | 0.826 | TIP41-like protein OS=Mus musculus GN=Tiprl PE=1 SV=1 |
| 503 | 1 | UniProt n | Q9D1J3 | 144 | 31778 | 5 | 5 | 3 | 3 | 0.49 | 1 | 0.94 | 0.977 | 0.834 | 0.836 | 0.809 | 0.972 | 0.826 | SAP domain-containing ribonucleoprotein OS=Mus musculus GN=Sarnp PE=1 SV=3 |
| 685 | 1 | UniProt n | P28474 | 85 | 49596 | 6 | 6 | 4 | 4 | 0.41 | 1 | 0.87 | 0.914 | 0.928 | 0.837 | 0.726 | 0.926 | 0.826 | Alcohol dehydrogenase class-3 OS=Mus musculus GN=Adh5 PE=1 SV=3 |
| 496 | 1 | UniProt n | Q3U4W8 | 146 | 109804 | 6 | 6 | 5 | 5 | 0.21 | 1 | 0.976 | 1.041 | 0.893 | 0.73 | 0.865 | 1.005 | 0.826 | Ubiquitinyl hydrolase 1 OS=Mus musculus GN=Usp5 PE=1 SV=1 |
| 1064 | 1 | UniProt n | Q8VD62 | 39 | 30093 | 2 | 2 | 2 | 2 | 0.32 | 1 | 0.882 | 0.917 | 1.026 | 0.82 | 0.67 | 0.932 | 0.826 | UPF0696 protein C11orf68 homolog OS=Mus musculus GN=Bles03 PE=1 SV=1 |
| 621 | 1 | UniProt n | Q9CXW3 | 100 | 36624 | 5 | 5 | 4 | 4 | 0.58 | 1 | 1.022 | 1.042 | 0.89 | 0.813 | 0.779 | 1.021 | 0.826 | Calcyclin-binding protein OS=Mus musculus GN=Cacybp PE=1 SV=1 |
| 550 | 1 | UniProt n | F7ANV6 | 122 | 29825 | 5 | 5 | 4 | 4 | 0.76 | 1 | 0.978 | 1.042 | 0.861 | 0.811 | 0.807 | 1.006 | 0.826 | Annexin (Fragment) OS=Mus musculus GN=Anxa4 PE=1 SV=1 |
| 812 | 1 | UniProt n | F6ZDS4 | 64 | 327990 | 1 | 1 | 1 | 1 | 0.01 | 1 | 0.891 | 0.89 | 0.797 | 0.774 | 0.912 | 0.926 | 0.826 | Nucleoprotein TPR OS=Mus musculus GN=Tpr PE=1 SV=1 |
| 1063 | 1 | UniProt n | A0A0J9YU | 36 | 15544 | 2 | 2 | 2 | 2 | 0.71 | 1 | 1.073 | 1.076 | 0.873 | 0.841 | 0.766 | 1.049 | 0.825 | Ubiquitin-conjugating enzyme E2 A OS=Mus musculus GN=Ube2a PE=1 SV=1 |
| 1007 | 1 | UniProt n | P97370 | 44 | 39636 | 1 | 1 | 1 | 1 | 0.11 | 1 | 1.084 | 0.977 | 0.757 | 0.942 | 0.788 | 1.019 | 0.825 | Sodium/potassium-transporting ATPase subunit beta-3 OS=Mus musculus GN=Atp1b3 PE=1 SV=1 |
| 1255 | 1 | UniProt n | Q8BGC0 | 22 | 107241 | 1 | 1 | 1 | 1 | 0.04 | 1 | 1.023 | 0.954 | 0.783 | 0.924 | 0.776 | 0.992 | 0.825 | HIV Tat-specific factor 1 homolog OS=Mus musculus GN=Htatsf1 PE=1 SV=1 |
| 655 | 1 | UniProt n | A0A0G2JE | 90 | 16669 | 6 | 6 | 3 | 3 | 1.11 | 1 | 1.119 | 1.169 | 0.803 | 0.891 | 0.782 | 1.094 | 0.824 | RNA-binding protein 8A OS=Mus musculus GN=Rbm8a PE=1 SV=1 |
| 1176 | 1 | UniProt n | D3YYB0 | 30 | 15139 | 1 | 1 | 1 | 1 | 0.32 | 1 | 0.896 | 0.898 | 0.78 | 0.893 | 0.803 | 0.930 | 0.824 | 40S ribosomal protein S27 OS=Mus musculus GN=Rps27l PE=1 SV=1 |
| 466 | 1 | UniProt n | Q8CI51 | 162 | 74918 | 8 | 8 | 6 | 6 | 0.49 | 1 | 0.971 | 0.948 | 0.802 | 0.833 | 0.837 | 0.973 | 0.824 | PDZ and LIM domain protein 5 OS=Mus musculus GN=Pdlim5 PE=1 SV=4 |
| 185 | 3 | UniProt n | E9Q5B6 | 143 | 17653 | 6 | 6 | 2 | 2 | 0.6 | 1 | 1.093 | 1.033 | 0.854 | 0.816 | 0.802 | 1.041 | 0.824 | Heterogeneous nuclear ribonucleoprotein D0 (Fragment) OS=Mus musculus GN=Hnrnpd PE=1 SV=1 |
| 258 | 1 | UniProt n | E0CYV0 | 341 | 36294 | 9 | 9 | 4 | 4 | 0.59 | 1 | 1.011 | 1.05 | 0.87 | 0.837 | 0.767 | 1.020 | 0.824 | Protein-L-isoaspartate O-methyltransferase OS=Mus musculus GN=Pcmt1 PE=1 SV=1 |
| 1185 | 1 | UniProt n | D3YVV9 | 29 | 150787 | 2 | 2 | 1 | 1 | 0.03 | 1 | 1.235 | 0.821 | 0.559 | 0.864 | 1.156 | 1.005 | 0.823 | Synaptopodin-2 OS=Mus musculus GN=Synpo2 PE=1 SV=1 |
| 786 | 1 | UniProt n | B2RSH2 | 68 | 50225 | 2 | 2 | 2 | 2 | 0.18 | 1 | 0.558 | 0.802 | 0.778 | 0.935 | 0.767 | 0.765 | 0.823 | Guanine nucleotide-binding protein G(i) subunit alpha-1 OS=Mus musculus GN=Gnai1 PE=1 SV=1 |
| 903 | 1 | UniProt n | A0A1D5RI | 55 | 79078 | 2 | 2 | 2 | 2 | 0.11 | 1 | 0.973 | 0.872 | 0.817 | 0.806 | 0.847 | 0.947 | 0.823 | Epidermal growth factor receptor substrate 15-like 1 OS=Mus musculus GN=Eps15l1 PE=1 SV=1 |
| 831 | 1 | UniProt n | H3BIX0 | 62 | 136559 | 2 | 2 | 2 | 2 | 0.06 | 1 | 0.896 | 0.918 | 0.798 | 0.898 | 0.778 | 0.937 | 0.823 | Treacle protein (Fragment) OS=Mus musculus GN=Tcof1 PE=1 SV=1 |
| 182 | 1 | UniProt n | B2M1R6 | 507 | 55409 | 22 | 22 | 10 | 10 | 1.14 | 1 | 0.977 | 0.989 | 0.846 | 0.832 | 0.792 | 0.989 | 0.823 | Heterogeneous nuclear ribonucleoprotein K OS=Mus musculus GN=Hnrnpk PE=1 SV=1 |
| 513 | 1 | UniProt n | P17427 | 136 | 121114 | 8 | 8 | 7 | 7 | 0.28 | 1 | 1.016 | 1.058 | 0.843 | 0.843 | 0.784 | 1.024 | 0.823 | AP-2 complex subunit alpha-2 OS=Mus musculus GN=Ap2a2 PE=1 SV=2 |
| 460 | 1 | UniProt n | Q9CZD3 | 167 | 97984 | 3 | 3 | 2 | 2 | 0.09 | 1 | 0.884 | 0.882 | 0.797 | 0.774 | 0.903 | 0.920 | 0.823 | Glycine--tRNA ligase OS=Mus musculus GN=Gars PE=1 SV=1 |
| 640 | 1 | UniProt n | Q8C281 | 93 | 101940 | 2 | 2 | 2 | 2 | 0.09 | 1 | 0.95 | 1.012 | 0.882 | 0.756 | 0.834 | 0.987 | 0.822 | Calpastatin OS=Mus musculus GN=Cast PE=2 SV=1 |
| 8 | 3 | UniProt n | P68134 | 2780 | 48384 | 156 | 156 | 13 | 13 | 4.25 | 1 | 1.112 | 1.178 | 0.795 | 0.777 | 0.9 | 1.094 | 0.822 | Actin, alpha skeletal muscle OS=Mus musculus GN=Acta1 PE=1 SV=1 |
| 172 | 1 | UniProt n | P28352 | 543 | 45220 | 23 | 23 | 6 | 6 | 0.75 | 1 | 0.989 | 1.007 | 0.846 | 0.836 | 0.786 | 0.999 | 0.822 | DNA-(apurinic or apyrimidinic site) lyase OS=Mus musculus GN=Apex1 PE=1 SV=2 |
| 133 | 1 | UniProt n | P10107 | 749 | 48066 | 30 | 30 | 9 | 9 | 1.41 | 1 | 1.028 | 1.034 | 0.784 | 0.829 | 0.855 | 1.021 | 0.822 | Annexin A1 OS=Mus musculus GN=Anxa1 PE=1 SV=2 |
| 520 | 1 | UniProt n | P62908 | 133 | 33184 | 5 | 5 | 4 | 4 | 0.66 | 1 | 0.95 | 1.054 | 0.808 | 0.79 | 0.87 | 1.000 | 0.822 | 40S ribosomal protein S3 OS=Mus musculus GN=Rps3 PE=1 SV=1 |
| 8 | 2 | UniProt n | P68033 | 2892 | 48352 | 156 | 156 | 12 | 12 | 4.25 | 1 | 1.111 | 1.177 | 0.795 | 0.777 | 0.899 | 1.094 | 0.822 | Actin, alpha cardiac muscle 1 OS=Mus musculus GN=Actc1 PE=1 SV=1 |
| 399 | 1 | UniProt n | E9Q718 | 208 | 102562 | 9 | 9 | 6 | 6 | 0.28 | 1 | 1.198 | 0.966 | 0.841 | 0.849 | 0.777 | 1.050 | 0.822 | Procollagen-lysine,2-oxoglutarate 5-dioxygenase 2 OS=Mus musculus GN=Plod2 PE=1 SV=1 |
| 402 | 2 | UniProt n | Q91V41 | 87 | 28021 | 3 | 3 | 2 | 2 | 0.35 | 1 | 0.96 | 1.069 | 0.846 | 0.746 | 0.879 | 1.009 | 0.822 | Ras-related protein Rab-14 OS=Mus musculus GN=Rab14 PE=1 SV=3 |
| 470 | 1 | UniProt n | A0A0G2J( | 160 | 27079 | 3 | 3 | 3 | 3 | 0.59 | 1 | 0.942 | 1.001 | 0.768 | 1.001 | 0.721 | 0.981 | 0.821 | Dihydropteridine reductase OS=Mus musculus GN=Qdpr PE=1 SV=1 |
| 815 | 1 | UniProt n | A0A0A0M | 63 | 14768 | 3 | 3 | 2 | 2 | 0.75 | 1 | 0.939 | 0.89 | 0.877 | 0.798 | 0.792 | 0.942 | 0.821 | Neuroplastin OS=Mus musculus GN=Nptn PE=1 SV=1 |
| 855 | 1 | UniProt n | A0A0U1R( | 59 | 15867 | 2 | 2 | 1 | 1 | 0.3 | 1 | 0.91 | 0.97 | 0.85 | 0.82 | 0.795 | 0.959 | 0.821 | Ubiquitin domain-containing protein UBFD1 (Fragment) OS=Mus musculus GN=Ubfd1 PE=1 SV=1 |
| 1213 | 1 | UniProt n | A0A140LJ | 27 | 64586 | 1 | 1 | 1 | 1 | 0.07 | 1 | 0.951 | 0.917 | 0.808 | 0.829 | 0.827 | 0.955 | 0.821 | COP9 signalosome complex subunit 1 OS=Mus musculus GN=Gps1 PE=1 SV=1 |
| 185 | 2 | UniProt n | D3YTQ3 | 159 | 55202 | 5 | 5 | 2 | 2 | 0.17 | 1 | 1.092 | 1.024 | 0.852 | 0.809 | 0.803 | 1.038 | 0.821 | Heterogeneous nuclear ribonucleoprotein D-like OS=Mus musculus GN=Hnrnpdl PE=1 SV=1 |
| 866 | 1 | UniProt n | P0DP26 | 58 | 19565 | 3 | 3 | 2 | 2 | 0.53 | 1 | 1.052 | 0.796 | 0.653 | 0.947 | 0.895 | 0.943 | 0.821 | Calmodulin-1 OS=Mus musculus GN=Calm1 PE=1 SV=1 |
| 993 | 1 | UniProt n | E9PX03 | 46 | 19898 | 2 | 2 | 2 | 2 | 0.52 | 1 | 0.815 | 0.973 | 0.924 | 0.794 | 0.754 | 0.926 | 0.821 | Coronin OS=Mus musculus GN=Coro1c PE=1 SV=1 |
| 781 | 1 | UniProt n | F6TN03 | 68 | 27383 | 3 | 3 | 2 | 2 | 0.36 | 1 | 0.839 | 0.872 | 0.865 | 0.818 | 0.781 | 0.901 | 0.821 | Ribosome maturation protein SBDS (Fragment) OS=Mus musculus GN=Sbds PE=1 SV=1 |
| 371 | 1 | UniProt n | A0A087W | 229 | 26604 | 8 | 8 | 4 | 4 | 0.88 | 1 | 1.134 | 1.047 | 0.779 | 0.84 | 0.844 | 1.059 | 0.820 | Eukaryotic translation elongation factor 1 beta 2 OS=Mus musculus GN=Eef1b2 PE=1 SV=1 |
| 302 | 1 | UniProt n | P63028 | 291 | 24105 | 13 | 13 | 4 | 4 | 1 | 1 | 1.123 | 1.087 | 0.788 | 0.872 | 0.803 | 1.069 | 0.820 | Translationally-controlled tumor protein OS=Mus musculus GN=Tpt1 PE=1 SV=1 |
| 479 | 1 | UniProt n | O08997 | 155 | 9905 | 8 | 8 | 5 | 5 | 6.91 | 1 | 1.074 | 1.015 | 0.849 | 0.863 | 0.753 | 1.029 | 0.820 | Copper transport protein ATOX1 OS=Mus musculus GN=Atox1 PE=1 SV=1 |
| 955 | 1 | UniProt n | Q9EPL8 | 49 | 136895 | 1 | 1 | 1 | 1 | 0.03 | 1 | 0.952 | 1.094 | 0.929 | 0.755 | 0.786 | 1.014 | 0.820 | Importin-7 OS=Mus musculus GN=Ipo7 PE=1 SV=2 |
| 856 | 1 | UniProt n | Q9CQM5 | 59 | 18236 | 5 | 5 | 2 | 2 | 0.58 | 1 | 0.936 | 0.831 | 0.849 | 0.86 | 0.754 | 0.920 | 0.820 | Thioredoxin domain-containing protein 17 OS=Mus musculus GN=Txndc17 PE=1 SV=1 |
| 228 | 1 | UniProt n | Q9CPV4 | 382 | 40826 | 16 | 16 | 8 | 8 | 1.53 | 1 | 0.968 | 0.974 | 0.788 | 0.873 | 0.8 | 0.981 | 0.819 | Glyoxalase domain-containing protein 4 OS=Mus musculus GN=Glod4 PE=1 SV=1 |
| 1001 | 1 | UniProt n | B1AUX2 | 44 | 238642 | 1 | 1 | 1 | 1 | 0.02 | 1 | 1.191 | 1.215 | 0.76 | 0.797 | 0.907 | 1.131 | 0.819 | Host cell factor 1 OS=Mus musculus GN=Hcfc1 PE=1 SV=1 |
| 695 | 1 | UniProt n | Q3ULG5 | 84 | 104199 | 5 | 5 | 4 | 4 | 0.18 | 1 | 0.918 | 1.03 | 0.827 | 0.801 | 0.828 | 0.982 | 0.819 | DNA helicase OS=Mus musculus GN=Mcm6 PE=1 SV=1 |
| 773 | 1 | UniProt n | A0A1B0G | 70 | 50913 | 4 | 4 | 3 | 3 | 0.28 | 1 | 1.033 | 1.09 | 0.868 | 0.842 | 0.75 | 1.040 | 0.818 | Glutaredoxin-3 OS=Mus musculus GN=Glrx3 PE=1 SV=1 |
| 61 | 1 | UniProt n | P07356 | 1525 | 49225 | 50 | 50 | 14 | 14 | 2.63 | 1 | 1.054 | 1.037 | 0.781 | 0.821 | 0.854 | 1.030 | 0.818 | Annexin A2 OS=Mus musculus GN=Anxa2 PE=1 SV=2 |
| 487 | 1 | UniProt n | G3UYP0 | 152 | 15172 | 8 | 8 | 4 | 4 | 1.98 | 1 | 1.072 | 1.082 | 0.794 | 0.854 | 0.807 | 1.051 | 0.818 | SUMO-conjugating enzyme (Fragment) OS=Mus musculus GN=Ube2i PE=1 SV=1 |
| 1242 | 1 | UniProt n | P35550 | 24 | 40462 | 1 | 1 | 1 | 1 | 0.11 | 1 | 0.708 | 0.819 | 0.799 | 0.859 | 0.796 | 0.834 | 0.817 | rRNA 2'-O-methyltransferase fibrillarin OS=Mus musculus GN=Fbl PE=1 SV=2 |
| 652 | 1 | UniProt n | A0A023T7 | 90 | 21604 | 3 | 3 | 2 | 2 | 0.47 | 1 | 0.969 | 0.88 | 0.784 | 0.882 | 0.79 | 0.948 | 0.817 | Mago nashi protein OS=Mus musculus GN=Magohb PE=1 SV=1 |
| 251 | 2 | UniProt n | B7ZBY7 | 316 | 22553 | 11 | 11 | 6 | 6 | 2.67 | 1 | 1.042 | 1.122 | 0.834 | 0.818 | 0.798 | 1.053 | 0.817 | Ubiquitin-conjugating enzyme E2 variant 1 OS=Mus musculus GN=Ube2v1 PE=3 SV=1 |
| 448 | 1 | UniProt n | P14602 | 174 | 25479 | 4 | 4 | 2 | 2 | 0.39 | 1 | 0.914 | 0.906 | 0.893 | 0.75 | 0.812 | 0.939 | 0.816 | Heat shock protein beta-1 OS=Mus musculus GN=Hspb1 PE=1 SV=3 |
| 249 | 1 | UniProt n | D3Z794 | 352 | 10890 | 12 | 12 | 2 | 2 | 2.1 | 1 | 1.065 | 1.069 | 0.855 | 0.808 | 0.786 | 1.044 | 0.816 | Small ubiquitin-related modifier 2 OS=Mus musculus GN=Sumo2 PE=1 SV=1 |
| 354 | 1 | UniProt n | Q8BJY1 | 239 | 62647 | 5 | 5 | 3 | 3 | 0.22 | 1 | 0.98 | 0.982 | 0.836 | 0.763 | 0.851 | 0.987 | 0.816 | 26S proteasome non-ATPase regulatory subunit 5 OS=Mus musculus GN=Psmd5 PE=1 SV=4 |
| 978 | 1 | UniProt n | A0A1L1SR | 47 | 6936 | 1 | 1 | 1 | 1 | 0.79 | 1 | 1.124 | 1.095 | 0.921 | 0.705 | 0.836 | 1.072 | 0.816 | Porphobilinogen deaminase (Fragment) OS=Mus musculus GN=Hmbs PE=1 SV=1 |
| 549 | 1 | UniProt n | Q99L45 | 122 | 52946 | 4 | 4 | 3 | 3 | 0.27 | 1 | 1.048 | 1.057 | 0.827 | 0.834 | 0.787 | 1.035 | 0.816 | Eukaryotic translation initiation factor 2 subunit 2 OS=Mus musculus GN=Eif2s2 PE=1 SV=1 |
| 1140 | 1 | UniProt n | Q8VEJ4 | 32 | 60816 | 1 | 1 | 1 | 1 | 0.07 | 1 | 1.153 | 1.111 | 0.836 | 0.779 | 0.832 | 1.086 | 0.815 | Notchless protein homolog 1 OS=Mus musculus GN=Nle1 PE=1 SV=4 |
| 532 | 1 | UniProt n | Q3UPL0 | 127 | 153478 | 4 | 4 | 2 | 2 | 0.09 | 1 | 1.018 | 0.992 | 0.806 | 0.853 | 0.788 | 1.003 | 0.815 | Protein transport protein Sec31A OS=Mus musculus GN=Sec31a PE=1 SV=2 |
| 19 | 2 | UniProt n | P20152 | 3243 | 60698 | 153 | 153 | 25 | 25 | 7.66 | 1 | 0.884 | 0.724 | 0.848 | 0.87 | 0.733 | 0.862 | 0.815 | Vimentin OS=Mus musculus GN=Vim PE=1 SV=3 |
| 75 | 1 | UniProt n | P09103 | 1295 | 72859 | 55 | 55 | 17 | 17 | 2.02 | 1 | 1.075 | 1.087 | 0.854 | 0.787 | 0.804 | 1.053 | 0.815 | Protein disulfide-isomerase OS=Mus musculus GN=P4hb PE=1 SV=2 |
| 21 | 1 | UniProt n | B7FAU9 | 4466 | 331602 | 164 | 164 | 74 | 74 | 1.75 | 1 | 1.006 | 1.035 | 0.84 | 0.8 | 0.804 | 1.014 | 0.814 | Filamin, alpha OS=Mus musculus GN=Flna PE=1 SV=1 |
| 215 | 1 | UniProt n | Q9CZ44 | 405 | 45857 | 12 | 12 | 6 | 6 | 0.74 | 1 | 0.971 | 0.988 | 0.809 | 0.813 | 0.821 | 0.986 | 0.814 | NSFL1 cofactor p47 OS=Mus musculus GN=Nsfl1c PE=1 SV=1 |
| 840 | 1 | UniProt n | A2A5F5 | 61 | 10864 | 2 | 2 | 1 | 1 | 0.46 | 1 | 0.975 | 1.016 | 0.768 | 0.847 | 0.83 | 0.997 | 0.814 | Ras-related protein Rab-5C (Fragment) OS=Mus musculus GN=Rab5c PE=1 SV=1 |
| 252 | 1 | UniProt n | P14206 | 350 | 36560 | 13 | 13 | 7 | 7 | 1.52 | 1 | 0.994 | 1.068 | 0.779 | 0.809 | 0.856 | 1.020 | 0.814 | 40S ribosomal protein SA OS=Mus musculus GN=Rpsa PE=1 SV=4 |
| 1178 | 1 | UniProt n | Q9QYJ3 | 30 | 46799 | 1 | 1 | 1 | 1 | 0.09 | 1 | 0.95 | 0.915 | 0.861 | 0.845 | 0.74 | 0.954 | 0.814 | DnaJ homolog subfamily B member 1 OS=Mus musculus GN=Dnajb1 PE=1 SV=3 |
| 632 | 1 | UniProt n | Q3TXS7 | 95 | 128089 | 3 | 3 | 3 | 3 | 0.1 | 1 | 1.04 | 1.093 | 0.816 | 0.726 | 0.907 | 1.044 | 0.813 | 26S proteasome non-ATPase regulatory subunit 1 OS=Mus musculus GN=Psmd1 PE=1 SV=1 |
| 342 | 1 | UniProt n | P67984 | 247 | 20880 | 7 | 7 | 3 | 3 | 0.82 | 1 | 0.998 | 0.996 | 0.861 | 0.79 | 0.789 | 0.998 | 0.813 | 60S ribosomal protein L22 OS=Mus musculus GN=Rpl22 PE=1 SV=2 |
| 235 | 2 | UniProt n | A0A0R4J2 | 156 | 75770 | 8 | 8 | 7 | 7 | 0.48 | 1 | 0.939 | 0.958 | 0.866 | 0.854 | 0.725 | 0.965 | 0.812 | Heterogeneous nuclear ribonucleoprotein Q OS=Mus musculus GN=Syncrip PE=1 SV=1 |
| 1230 | 1 | UniProt n | A2A701 | 26 | 20859 | 1 | 1 | 1 | 1 | 0.22 | 1 | 0.785 | 1 | 0.809 | 0.849 | 0.78 | 0.922 | 0.812 | Eukaryotic translation initiation factor 3 subunit M (Fragment) OS=Mus musculus GN=Eif3m PE=1 SV=1 |
| 394 | 2 | UniProt n | P53811 | 186 | 40215 | 8 | 8 | 5 | 5 | 0.69 | 1 | 1.002 | 1.002 | 0.847 | 0.816 | 0.772 | 1.001 | 0.811 | Phosphatidylinositol transfer protein beta isoform OS=Mus musculus GN=Pitpnb PE=1 SV=2 |
| 1033 | 1 | UniProt n | G3X956 | 42 | 153077 | 2 | 2 | 2 | 2 | 0.06 | 1 | 1.023 | 1.114 | 0.845 | 0.833 | 0.758 | 1.045 | 0.811 | Suppressor of Ty 16 OS=Mus musculus GN=Supt16 PE=1 SV=1 |
| 707 | 1 | UniProt n | E9Q3X0 | 80 | 111325 | 3 | 3 | 3 | 3 | 0.12 | 1 | 0.69 | 0.903 | 0.859 | 0.795 | 0.78 | 0.854 | 0.811 | Major vault protein OS=Mus musculus GN=Mvp PE=1 SV=1 |
| 47 | 1 | UniProt n | P17742 | 2080 | 22357 | 123 | 123 | 12 | 12 | 50.15 | 1 | 1.07 | 1.036 | 0.81 | 0.755 | 0.87 | 1.035 | 0.810 | Peptidyl-prolyl cis-trans isomerase A OS=Mus musculus GN=Ppia PE=1 SV=2 |
| 546 | 1 | UniProt n | Q9WTQ5 | 123 | 219680 | 5 | 5 | 2 | 2 | 0.06 | 1 | 0.782 | 0.845 | 0.884 | 0.824 | 0.729 | 0.871 | 0.810 | A-kinase anchor protein 12 OS=Mus musculus GN=Akap12 PE=1 SV=1 |
| 162 | 1 | UniProt n | Q8CDN6 | 595 | 38319 | 16 | 16 | 8 | 8 | 1.69 | 1 | 0.917 | 1.001 | 0.828 | 0.743 | 0.863 | 0.972 | 0.810 | Thioredoxin-like protein 1 OS=Mus musculus GN=Txnl1 PE=1 SV=3 |
| 599 | 1 | UniProt n | P35505 | 103 | 51593 | 4 | 4 | 4 | 4 | 0.39 | 1 | 0.935 | 1.032 | 0.808 | 0.902 | 0.728 | 0.988 | 0.810 | Fumarylacetoacetase OS=Mus musculus GN=Fah PE=1 SV=2 |
| 784 | 1 | UniProt n | I1E4X7 | 68 | 20739 | 3 | 3 | 2 | 2 | 0.5 | 1 | 0.96 | 0.931 | 0.859 | 0.776 | 0.795 | 0.963 | 0.809 | Diphosphoinositol polyphosphate phosphohydrolase 1 OS=Mus musculus GN=Nudt3 PE=1 SV=1 |
| 271 | 2 | UniProt n | P60766 | 147 | 26997 | 5 | 5 | 3 | 3 | 0.59 | 1 | 0.992 | 1.037 | 0.808 | 0.812 | 0.807 | 1.009 | 0.809 | Cell division control protein 42 homolog OS=Mus musculus GN=Cdc42 PE=1 SV=2 |
| 743 | 1 | UniProt n | Q9D8U8 | 74 | 60383 | 2 | 2 | 1 | 1 | 0.07 | 1 | 0.815 | 0.687 | 0.705 | 0.909 | 0.826 | 0.824 | 0.809 | Sorting nexin-5 OS=Mus musculus GN=Snx5 PE=1 SV=1 |
| 772 | 1 | UniProt n | E9PYX7 | 70 | 222074 | 2 | 2 | 2 | 2 | 0.04 | 1 | 0.952 | 0.991 | 0.8 | 0.921 | 0.718 | 0.981 | 0.809 | Afadin OS=Mus musculus GN=Afdn PE=1 SV=1 |
| 1253 | 1 | UniProt n | G5DDB7 | 23 | 103823 | 1 | 1 | 1 | 1 | 0.04 | 1 | 1.035 | 1 | 0.821 | 0.677 | 0.95 | 1.012 | 0.808 | Phosphoinositide phospholipase C OS=Mus musculus GN=Plcd1 PE=1 SV=1 |
| 617 | 1 | UniProt n | D3YYD5 | 101 | 16590 | 6 | 6 | 3 | 3 | 1.12 | 1 | 0.943 | 1.051 | 0.851 | 0.825 | 0.752 | 0.997 | 0.808 | Vacuolar protein sorting-associated protein 29 (Fragment) OS=Mus musculus GN=Vps29 PE=1 SV=1 |
| 680 | 1 | UniProt n | Q5SUH6 | 86 | 82537 | 3 | 3 | 2 | 2 | 0.17 | 1 | 1.154 | 1.124 | 0.834 | 0.835 | 0.758 | 1.091 | 0.808 | Clathrin interactor 1 OS=Mus musculus GN=Clint1 PE=1 SV=1 |
| 467 | 1 | UniProt n | Q5SUR0 | 161 | 159302 | 5 | 5 | 5 | 5 | 0.14 | 1 | 1 | 1.111 | 0.81 | 0.781 | 0.833 | 1.036 | 0.808 | Phosphoribosylformylglycinamidine synthase OS=Mus musculus GN=Pfas PE=1 SV=1 |
| 333 | 1 | UniProt n | P70168 | 260 | 111519 | 17 | 17 | 8 | 8 | 0.41 | 1 | 0.986 | 1.016 | 0.815 | 0.819 | 0.789 | 1.001 | 0.808 | Importin subunit beta-1 OS=Mus musculus GN=Kpnb1 PE=1 SV=2 |
| 474 | 1 | UniProt n | Q9R0P4 | 157 | 25252 | 2 | 2 | 2 | 2 | 0.39 | 1 | 0.962 | 0.975 | 0.784 | 0.849 | 0.791 | 0.979 | 0.807 | Small acidic protein OS=Mus musculus GN=Smap PE=1 SV=1 |
| 733 | 1 | UniProt n | O35350 | 76 | 95383 | 4 | 4 | 3 | 3 | 0.14 | 1 | 0.859 | 0.875 | 0.74 | 0.884 | 0.804 | 0.909 | 0.807 | Calpain-1 catalytic subunit OS=Mus musculus GN=Capn1 PE=1 SV=1 |
| 15 | 2 | UniProt n | Q3UH59 | 796 | 294366 | 30 | 30 | 14 | 14 | 0.22 | 1 | 1.069 | 1.122 | 0.799 | 0.8 | 0.822 | 1.062 | 0.807 | Myosin-10 OS=Mus musculus GN=Myh10 PE=1 SV=1 |
| 66 | 1 | UniProt n | Q5SQB0 | 1477 | 38466 | 46 | 46 | 7 | 7 | 2.33 | 1 | 1.016 | 0.997 | 0.82 | 0.814 | 0.787 | 1.004 | 0.807 | Nucleophosmin OS=Mus musculus GN=Npm1 PE=1 SV=1 |
| 789 | 1 | UniProt n | P58389 | 67 | 44218 | 3 | 3 | 2 | 2 | 0.21 | 1 | 1.017 | 1.025 | 0.782 | 0.847 | 0.793 | 1.014 | 0.807 | Serine/threonine-protein phosphatase 2A activator OS=Mus musculus GN=Ptpa PE=1 SV=1 |
| 74 | 1 | UniProt n | Q9DCD0 | 1299 | 64274 | 41 | 41 | 14 | 14 | 1.87 | 1 | 1.081 | 1.09 | 0.811 | 0.78 | 0.829 | 1.056 | 0.806 | 6-phosphogluconate dehydrogenase, decarboxylating OS=Mus musculus GN=Pgd PE=1 SV=3 |
| 1236 | 1 | UniProt n | Q9WTM5 | 24 | 59736 | 1 | 1 | 1 | 1 | 0.07 | 1 | 0.765 | 0.923 | 0.831 | 0.865 | 0.729 | 0.890 | 0.806 | RuvB-like 2 OS=Mus musculus GN=Ruvbl2 PE=1 SV=3 |
| 635 | 1 | UniProt n | P60229 | 94 | 60723 | 4 | 4 | 3 | 3 | 0.23 | 1 | 0.818 | 0.802 | 0.813 | 0.858 | 0.751 | 0.869 | 0.806 | Eukaryotic translation initiation factor 3 subunit E OS=Mus musculus GN=Eif3e PE=1 SV=1 |
| 245 | 1 | UniProt n | P24527 | 356 | 81377 | 21 | 21 | 11 | 11 | 0.87 | 1 | 1.027 | 1.026 | 0.816 | 0.812 | 0.789 | 1.018 | 0.806 | Leukotriene A-4 hydrolase OS=Mus musculus GN=Lta4h PE=1 SV=4 |
| 80 | 1 | UniProt n | P09405 | 1214 | 104405 | 64 | 64 | 23 | 23 | 1.87 | 1 | 1.05 | 1.02 | 0.81 | 0.791 | 0.814 | 1.023 | 0.805 | Nucleolin OS=Mus musculus GN=Ncl PE=1 SV=2 |
| 777 | 1 | UniProt n | A0A1L1SS | 69 | 17883 | 1 | 1 | 1 | 1 | 0.26 | 1 | 1.221 | 1.269 | 0.822 | 0.75 | 0.845 | 1.157 | 0.805 | Cysteine and histidine-rich domain-containing protein 1 OS=Mus musculus GN=Chordc1 PE=1 SV=1 |
| 383 | 1 | UniProt n | Q8K4Z5 | 222 | 106483 | 8 | 8 | 3 | 3 | 0.22 | 1 | 0.978 | 0.975 | 0.834 | 0.909 | 0.687 | 0.984 | 0.805 | Splicing factor 3A subunit 1 OS=Mus musculus GN=Sf3a1 PE=1 SV=1 |
| 251 | 1 | UniProt n | Q9D2M8 | 350 | 19444 | 11 | 11 | 5 | 5 | 3.49 | 1 | 1.037 | 1.125 | 0.814 | 0.806 | 0.793 | 1.053 | 0.804 | Ubiquitin-conjugating enzyme E2 variant 2 OS=Mus musculus GN=Ube2v2 PE=1 SV=4 |
| 1259 | 1 | UniProt n | A0A1W2P | 21 | 294240 | 1 | 1 | 1 | 1 | 0.01 | 1 | 0.948 | 0.996 | 0.719 | 0.796 | 0.909 | 0.981 | 0.804 | Tyrosine-protein phosphatase OS=Mus musculus GN=Ptprb PE=1 SV=1 |
| 257 | 1 | UniProt n | Q8BGQ7 | 342 | 127000 | 13 | 13 | 9 | 9 | 0.35 | 1 | 0.969 | 1.044 | 0.839 | 0.786 | 0.788 | 1.004 | 0.804 | Alanine--tRNA ligase, cytoplasmic OS=Mus musculus GN=Aars PE=1 SV=1 |
| 365 | 1 | UniProt n | Q3KQM4 | 231 | 39327 | 10 | 10 | 6 | 6 | 1.12 | 1 | 1.06 | 1.044 | 0.753 | 0.834 | 0.827 | 1.034 | 0.804 | Splicing factor U2AF 65 kDa subunit OS=Mus musculus GN=U2af2 PE=1 SV=1 |
| 234 | 2 | UniProt n | P80318 | 336 | 72003 | 20 | 20 | 11 | 11 | 1.28 | 1 | 0.986 | 1.026 | 0.803 | 0.78 | 0.828 | 1.004 | 0.803 | T-complex protein 1 subunit gamma OS=Mus musculus GN=Cct3 PE=1 SV=1 |
| 519 | 1 | UniProt n | F6QTS1 | 133 | 33850 | 5 | 5 | 3 | 3 | 0.86 | 1 | 0.969 | 0.928 | 0.754 | 0.854 | 0.805 | 0.965 | 0.803 | COP9 signalosome complex subunit 4 (Fragment) OS=Mus musculus GN=Cops4 PE=1 SV=1 |
| 299 | 1 | UniProt n | Q99LS3 | 293 | 29523 | 14 | 14 | 5 | 5 | 1.71 | 1 | 0.93 | 0.886 | 0.796 | 0.855 | 0.761 | 0.938 | 0.803 | Phosphoserine phosphatase OS=Mus musculus GN=Psph PE=1 SV=1 |
| 405 | 1 | UniProt n | Q99KC8 | 205 | 105025 | 11 | 11 | 7 | 7 | 0.38 | 1 | 0.942 | 0.917 | 0.802 | 0.848 | 0.761 | 0.952 | 0.803 | von Willebrand factor A domain-containing protein 5A OS=Mus musculus GN=Vwa5a PE=1 SV=2 |
| 787 | 1 | UniProt n | A0A0U1RI | 68 | 47335 | 2 | 2 | 1 | 1 | 0.09 | 1 | 0.888 | 1.012 | 0.798 | 0.737 | 0.88 | 0.965 | 0.803 | S-adenosylmethionine synthase OS=Mus musculus GN=Mat2a PE=1 SV=1 |
| 8 | 1 | UniProt n | P60710 | 5890 | 48070 | 228 | 228 | 19 | 19 | 20.63 | 1 | 1.076 | 1.115 | 0.778 | 0.758 | 0.876 | 1.063 | 0.802 | Actin, cytoplasmic 1 OS=Mus musculus GN=Actb PE=1 SV=1 |
| 439 | 2 | UniProt n | P80316 | 162 | 73643 | 8 | 8 | 5 | 5 | 0.33 | 1 | 0.949 | 1.088 | 0.807 | 0.827 | 0.774 | 1.011 | 0.802 | T-complex protein 1 subunit epsilon OS=Mus musculus GN=Cct5 PE=1 SV=1 |
| 266 | 1 | UniProt n | O70435 | 328 | 34655 | 19 | 19 | 8 | 8 | 1.98 | 1 | 1.055 | 1.078 | 0.78 | 0.837 | 0.791 | 1.044 | 0.802 | Proteasome subunit alpha type-3 OS=Mus musculus GN=Psma3 PE=1 SV=3 |
| 990 | 1 | UniProt n | Q9D820 | 46 | 23200 | 1 | 1 | 1 | 1 | 0.2 | 1 | 0.774 | 0.956 | 0.776 | 0.913 | 0.727 | 0.904 | 0.802 | Prolyl-tRNA synthetase associated domain-containing protein 1 OS=Mus musculus GN=Prorsd1 PE=1 SV=1 |
| 175 | 1 | UniProt n | P47753 | 532 | 39141 | 17 | 17 | 7 | 7 | 1.12 | 1 | 0.979 | 0.978 | 0.795 | 0.794 | 0.815 | 0.986 | 0.801 | F-actin-capping protein subunit alpha-1 OS=Mus musculus GN=Capza1 PE=1 SV=4 |
| 82 | 1 | UniProt n | P19096 | 1199 | 305797 | 53 | 53 | 30 | 30 | 0.54 | 1 | 0.973 | 1.055 | 0.807 | 0.783 | 0.814 | 1.009 | 0.801 | Fatty acid synthase OS=Mus musculus GN=Fasn PE=1 SV=2 |
| 17 | 1 | UniProt n | P63017 | 4729 | 87742 | 179 | 179 | 29 | 29 | 6.99 | 1 | 1.008 | 1.036 | 0.821 | 0.811 | 0.77 | 1.015 | 0.800 | Heat shock cognate 71 kDa protein OS=Mus musculus GN=Hspa8 PE=1 SV=1 |
| 402 | 1 | UniProt n | Q5SW88 | 207 | 28018 | 7 | 7 | 3 | 3 | 0.57 | 1 | 0.871 | 0.979 | 0.797 | 0.803 | 0.801 | 0.948 | 0.800 | RAB1A, member RAS oncogene family OS=Mus musculus GN=Rab1a PE=1 SV=1 |
| 843 | 1 | UniProt n | F6Q3W0 | 60 | 40129 | 2 | 2 | 1 | 1 | 0.11 | 1 | 0.889 | 1.006 | 0.724 | 0.878 | 0.806 | 0.963 | 0.800 | WD repeat-containing protein 5 (Fragment) OS=Mus musculus GN=Wdr5 PE=1 SV=1 |
| 1060 | 1 | UniProt n | P62814 | 40 | 63788 | 1 | 1 | 1 | 1 | 0.07 | 1 | 0.896 | 1.018 | 0.772 | 0.782 | 0.847 | 0.970 | 0.800 | V-type proton ATPase subunit B, brain isoform OS=Mus musculus GN=Atp6v1b2 PE=1 SV=1 |
| 285 | 1 | UniProt n | Q61081 | 310 | 57755 | 13 | 13 | 6 | 6 | 0.55 | 1 | 1.077 | 1.072 | 0.822 | 0.8 | 0.777 | 1.049 | 0.799 | Hsp90 co-chaperone Cdc37 OS=Mus musculus GN=Cdc37 PE=1 SV=1 |
| 1056 | 1 | UniProt n | Q8R0Y6 | 40 | 117284 | 1 | 1 | 1 | 1 | 0.04 | 1 | 1.106 | 1.178 | 0.721 | 0.787 | 0.9 | 1.092 | 0.799 | Cytosolic 10-formyltetrahydrofolate dehydrogenase OS=Mus musculus GN=Aldh1l1 PE=1 SV=1 |
| 1106 | 1 | UniProt n | P35585 | 35 | 58689 | 2 | 2 | 2 | 2 | 0.15 | 1 | 0.98 | 1.01 | 0.788 | 0.773 | 0.837 | 0.997 | 0.799 | AP-1 complex subunit mu-1 OS=Mus musculus GN=Ap1m1 PE=1 SV=3 |
| 715 | 1 | UniProt n | Q8K157 | 79 | 43435 | 3 | 3 | 2 | 2 | 0.21 | 1 | 0.811 | 0.834 | 0.794 | 0.818 | 0.784 | 0.878 | 0.799 | Aldose 1-epimerase OS=Mus musculus GN=Galm PE=1 SV=1 |

| | | | | | | | | | | | | | | | | | | | | |
|---|---|---|---|---|---|---|---|---|---|---|---|---|---|---|---|---|---|---|---|---|
| 250 | 2 | UniProt | n | A0A0G2JF | 41 | 77791 | 4 | 4 | 4 | 4 | 0.24 | 1 | 1.002 | 1.012 | 0.888 | 0.765 | 0.748 | 1.005 | 0.798 | Far upstream element-binding protein 1 OS=Mus musculus GN=Fubp1 PE=1 SV=1 |
| 795 | 1 | UniProt | n | P27048 | 66 | 27124 | 1 | 1 | 1 | 1 | 0.17 | 1 | 1.064 | 1.046 | 0.767 | 0.8 | 0.827 | 1.036 | 0.798 | Small nuclear ribonucleoprotein-associated protein B OS=Mus musculus GN=Snrpb PE=1 SV=1 |
| 94 | 1 | UniProt | n | Q3U2G2 | 1078 | 120042 | 50 | 50 | 25 | 25 | 1.6 | 1 | 0.967 | 0.985 | 0.815 | 0.789 | 0.789 | 0.984 | 0.798 | Heat shock 70 kDa protein 4 OS=Mus musculus GN=Hspa4 PE=1 SV=1 |
| 114 | 1 | UniProt | n | Q99LX0 | 885 | 25060 | 34 | 34 | 8 | 8 | 6.42 | 1 | 1.089 | 1.105 | 0.823 | 0.788 | 0.782 | 1.064 | 0.797 | Protein DJ-1 OS=Mus musculus GN=Park7 PE=1 SV=1 |
| 507 | 1 | UniProt | n | D6RCG1 | 140 | 38640 | 2 | 2 | 2 | 2 | 0.24 | 1 | 0.776 | 0.921 | 0.707 | 0.853 | 0.84 | 0.894 | 0.797 | Trifunctional purine biosynthetic protein adenosine-3 OS=Mus musculus GN=Gart PE=1 SV=1 |
| 699 | 1 | UniProt | n | F6RPJ9 | 82 | 136644 | 5 | 5 | 4 | 4 | 0.13 | 1 | 0.998 | 1.069 | 0.787 | 0.806 | 0.798 | 1.022 | 0.797 | Insulin-degrading enzyme (Fragment) OS=Mus musculus GN=Ide PE=1 SV=1 |
| 1111 | 1 | UniProt | n | A0A0R4J1 | 35 | 24534 | 2 | 2 | 1 | 1 | 0.41 | 1 | 0.955 | 0.921 | 0.762 | 0.76 | 0.874 | 0.958 | 0.797 | Methionine sulfoxide reductase B3 OS=Mus musculus GN=Msrb3 PE=1 SV=1 |
| 809 | 1 | UniProt | n | O35864 | 64 | 45922 | 4 | 4 | 4 | 4 | 0.44 | 1 | 0.998 | 1.202 | 0.861 | 0.757 | 0.776 | 1.063 | 0.797 | COP9 signalosome complex subunit 5 OS=Mus musculus GN=Cops5 PE=1 SV=3 |
| 757 | 1 | UniProt | n | Q91YP2 | 72 | 94969 | 3 | 3 | 3 | 3 | 0.14 | 1 | 0.925 | 0.994 | 0.851 | 0.788 | 0.754 | 0.972 | 0.797 | Neurolysin, mitochondrial OS=Mus musculus GN=Nln PE=1 SV=1 |
| 149 | 1 | UniProt | n | O08553 | 670 | 72599 | 19 | 19 | 12 | 12 | 1.4 | 1 | 0.989 | 1.009 | 0.8 | 0.794 | 0.796 | 0.999 | 0.797 | Dihydropyrimidinase-related protein 2 OS=Mus musculus GN=Dpysl2 PE=1 SV=2 |
| 426 | 2 | UniProt | n | O35737 | 143 | 54570 | 4 | 4 | 3 | 3 | 0.26 | 1 | 1.002 | 0.926 | 0.761 | 0.854 | 0.778 | 0.975 | 0.797 | Heterogeneous nuclear ribonucleoprotein H OS=Mus musculus GN=Hnrnph1 PE=1 SV=3 |
| 714 | 1 | UniProt | n | Q9JLV1 | 79 | 71436 | 3 | 3 | 2 | 2 | 0.13 | 1 | 0.928 | 1.02 | 0.819 | 0.77 | 0.801 | 0.982 | 0.796 | BAG family molecular chaperone regulator 3 OS=Mus musculus GN=Bag3 PE=1 SV=2 |
| 165 | 1 | UniProt | n | Q9D8N0 | 587 | 60648 | 33 | 33 | 11 | 11 | 1.31 | 1 | 1.018 | 1.037 | 0.795 | 0.805 | 0.789 | 1.018 | 0.796 | Elongation factor 1-gamma OS=Mus musculus GN=Eef1g PE=1 SV=3 |
| 613 | 1 | UniProt | n | Q06138 | 101 | 49994 | 4 | 4 | 3 | 3 | 0.29 | 1 | 0.841 | 0.822 | 0.729 | 1.012 | 0.684 | 0.884 | 0.796 | Calcium-binding protein 39 OS=Mus musculus GN=Cab39 PE=1 SV=2 |
| 584 | 1 | UniProt | n | Q9CQE8 | 111 | 34615 | 4 | 4 | 4 | 4 | 0.63 | 1 | 0.933 | 0.8 | 0.85 | 0.815 | 0.728 | 0.907 | 0.796 | UPF0568 protein C14orf166 homolog OS=Mus musculus PE=1 SV=1 |
| 177 | 2 | UniProt | n | E9Q7D8 | 338 | 29153 | 10 | 10 | 4 | 4 | 1.37 | 1 | 0.964 | 0.977 | 0.804 | 0.744 | 0.843 | 0.980 | 0.796 | Poly(rC)-binding protein 3 OS=Mus musculus GN=Pcbp3 PE=1 SV=1 |
| 887 | 1 | UniProt | n | Q9JLI8 | 56 | 130013 | 1 | 1 | 1 | 1 | 0.03 | 1 | 0.911 | 1.069 | 0.818 | 0.881 | 0.699 | 0.991 | 0.796 | Squamous cell carcinoma antigen recognized by T-cells 3 OS=Mus musculus GN=Sart3 PE=1 SV=1 |
| 918 | 1 | UniProt | n | Q9D5V6 | 53 | 51105 | 1 | 1 | 1 | 1 | 0.09 | 1 | 0.855 | 0.886 | 0.933 | 0.78 | 0.691 | 0.912 | 0.795 | Synapse-associated protein 1 OS=Mus musculus GN=Syap1 PE=1 SV=1 |
| 253 | 1 | UniProt | n | P68037 | 349 | 23768 | 25 | 25 | 5 | 5 | 1.41 | 1 | 1.063 | 1.069 | 0.769 | 0.822 | 0.794 | 1.044 | 0.795 | Ubiquitin-conjugating enzyme E2 L3 OS=Mus musculus GN=Ube2l3 PE=1 SV=1 |
| 839 | 1 | UniProt | n | E9Q1S3 | 61 | 92765 | 4 | 4 | 3 | 3 | 0.15 | 1 | 0.974 | 1.088 | 0.746 | 0.853 | 0.788 | 1.020 | 0.794 | Protein transport protein Sec23A OS=Mus musculus GN=Sec23a PE=1 SV=1 |
| 510 | 2 | UniProt | n | Q9QZM0 | 96 | 70959 | 2 | 2 | 2 | 2 | 0.13 | 1 | 0.739 | 0.856 | 0.875 | 0.754 | 0.76 | 0.858 | 0.794 | Ubiquilin-2 OS=Mus musculus GN=Ubqln2 PE=1 SV=2 |
| 937 | 1 | UniProt | n | E9Q5C9 | 50 | 107117 | 1 | 1 | 1 | 1 | 0.04 | 1 | 1.077 | 0.931 | 0.742 | 0.803 | 0.841 | 1.001 | 0.794 | Nucleolar and coiled-body phosphoprotein 1 OS=Mus musculus GN=Nolc1 PE=1 SV=1 |
| 492 | 1 | UniProt | n | Q8CGC7 | 148 | 215203 | 9 | 9 | 8 | 8 | 0.17 | 1 | 0.959 | 1.015 | 0.805 | 0.789 | 0.788 | 0.991 | 0.794 | Bifunctional glutamate/proline--tRNA ligase OS=Mus musculus GN=Eprs PE=1 SV=4 |
| 112 | 1 | UniProt | n | Q68FD5 | 894 | 222672 | 32 | 32 | 21 | 21 | 0.49 | 1 | 0.963 | 1.072 | 0.779 | 0.772 | 0.832 | 1.011 | 0.794 | Clathrin heavy chain 1 OS=Mus musculus GN=Cltc PE=1 SV=3 |
| 595 | 1 | UniProt | n | Q9CQI6 | 105 | 19676 | 7 | 7 | 4 | 4 | 1.34 | 1 | 0.943 | 0.855 | 0.779 | 0.864 | 0.743 | 0.931 | 0.794 | Coactosin-like protein OS=Mus musculus GN=Cotl1 PE=1 SV=3 |
| 440 | 1 | UniProt | n | P30681 | 176 | 36757 | 9 | 9 | 2 | 2 | 0.26 | 1 | 1.11 | 1.168 | 0.824 | 0.794 | 0.764 | 1.090 | 0.794 | High mobility group protein B2 OS=Mus musculus GN=Hmgb2 PE=1 SV=3 |
| 76 | 1 | UniProt | n | P17751 | 1290 | 39582 | 44 | 44 | 11 | 11 | 3.93 | 1 | 1.037 | 0.966 | 0.796 | 0.782 | 0.803 | 1.001 | 0.794 | Triosephosphate isomerase OS=Mus musculus GN=Tpi1 PE=1 SV=4 |
| 160 | 1 | UniProt | n | P26883 | 604 | 14699 | 19 | 19 | 3 | 3 | 3.11 | 1 | 0.973 | 0.98 | 0.773 | 0.824 | 0.784 | 0.984 | 0.793 | Peptidyl-prolyl cis-trans isomerase FKBP1A OS=Mus musculus GN=Fkbp1a PE=1 SV=2 |
| 1029 | 1 | UniProt | n | Q9WTL7 | 42 | 28704 | 2 | 2 | 2 | 2 | 0.34 | 1 | 0.888 | 0.918 | 0.879 | 0.789 | 0.72 | 0.934 | 0.793 | Acyl-protein thioesterase 2 OS=Mus musculus GN=Lypla2 PE=1 SV=1 |
| 877 | 1 | UniProt | n | Q8QZY9 | 57 | 48632 | 1 | 1 | 1 | 1 | 0.09 | 1 | 0.997 | 1.03 | 0.803 | 0.85 | 0.729 | 1.009 | 0.792 | Splicing factor 3B subunit 4 OS=Mus musculus GN=Sf3b4 PE=1 SV=1 |
| 633 | 1 | UniProt | n | P12382 | 95 | 97205 | 3 | 3 | 1 | 1 | 0.09 | 1 | 0.905 | 1.32 | 0.74 | 0.679 | 0.989 | 1.061 | 0.792 | ATP-dependent 6-phosphofructokinase, liver type OS=Mus musculus GN=Pfkl PE=1 SV=4 |
| 318 | 2 | UniProt | n | P53996 | 225 | 24241 | 13 | 13 | 6 | 6 | 1.82 | 1 | 0.943 | 0.854 | 0.798 | 0.771 | 0.806 | 0.930 | 0.792 | Cellular nucleic acid-binding protein OS=Mus musculus GN=Cnbp PE=1 SV=2 |
| 788 | 1 | UniProt | n | Q9WU84 | 68 | 32700 | 1 | 1 | 1 | 1 | 0.14 | 1 | 1.05 | 1.225 | 0.857 | 0.751 | 0.768 | 1.088 | 0.791 | Copper chaperone for superoxide dismutase OS=Mus musculus GN=Ccs PE=1 SV=1 |
| 177 | 1 | UniProt | n | P60335 | 528 | 42451 | 18 | 18 | 9 | 9 | 1.99 | 1 | 0.941 | 0.933 | 0.782 | 0.766 | 0.825 | 0.958 | 0.791 | Poly(rC)-binding protein 1 OS=Mus musculus GN=Pcbp1 PE=1 SV=1 |
| 874 | 1 | UniProt | n | A2AV19 | 57 | 32342 | 1 | 1 | 1 | 1 | 0.14 | 1 | 1.061 | 0.867 | 0.728 | 0.74 | 0.915 | 0.973 | 0.790 | Exosome complex component RRP4 (Fragment) OS=Mus musculus GN=Exosc2 PE=1 SV=1 |
| 827 | 1 | UniProt | n | P00493 | 62 | 30214 | 3 | 3 | 2 | 2 | 0.32 | 1 | 0.965 | 0.959 | 0.776 | 0.77 | 0.824 | 0.975 | 0.790 | Hypoxanthine-guanine phosphoribosyltransferase OS=Mus musculus GN=Hprt1 PE=1 SV=3 |
| 804 | 1 | UniProt | n | A2AJK8 | 64 | 36813 | 2 | 2 | 2 | 2 | 0.26 | 1 | 0.998 | 1.024 | 0.747 | 0.847 | 0.778 | 1.007 | 0.790 | Tetratricopeptide repeat protein 1 OS=Mus musculus GN=Ttc1 PE=1 SV=1 |
| 705 | 2 | UniProt | n | Q99K48 | 60 | 63116 | 4 | 4 | 3 | 3 | 0.31 | 1 | 1.012 | 0.982 | 0.824 | 0.802 | 0.743 | 0.998 | 0.789 | Non-POU domain-containing octamer-binding protein OS=Mus musculus GN=Nono PE=1 SV=3 |
| 12 | 3 | UniProt | n | P08113 | 1498 | 116072 | 85 | 85 | 29 | 29 | 2.22 | 1 | 1.044 | 1.076 | 0.804 | 0.783 | 0.779 | 1.040 | 0.789 | Endoplasmin OS=Mus musculus GN=Hsp90b1 PE=1 SV=2 |
| 247 | 1 | UniProt | n | Q05816 | 354 | 19966 | 18 | 18 | 5 | 5 | 2.5 | 1 | 0.973 | 0.94 | 0.819 | 0.777 | 0.77 | 0.971 | 0.788 | Fatty acid-binding protein, epidermal OS=Mus musculus GN=Fabp5 PE=1 SV=3 |
| 40 | 2 | UniProt | n | Q7TPR4 | 2182 | 119937 | 92 | 92 | 34 | 34 | 3.11 | 1 | 1.034 | 1.073 | 0.798 | 0.773 | 0.794 | 1.035 | 0.788 | Alpha-actinin-1 OS=Mus musculus GN=Actn1 PE=1 SV=1 |
| 185 | 1 | UniProt | n | Q20BD0 | 498 | 45407 | 20 | 20 | 6 | 6 | 0.92 | 1 | 1.041 | 0.989 | 0.809 | 0.778 | 0.778 | 1.010 | 0.788 | Heterogeneous nuclear ribonucleoprotein A/B OS=Mus musculus GN=Hnrnpab PE=1 SV=1 |
| 292 | 1 | UniProt | n | Q8BJW6 | 303 | 81204 | 9 | 9 | 5 | 5 | 0.37 | 1 | 0.884 | 0.864 | 0.78 | 0.762 | 0.823 | 0.914 | 0.788 | Eukaryotic translation initiation factor 2A OS=Mus musculus GN=Eif2a PE=1 SV=2 |
| 554 | 1 | UniProt | n | Q6P5F9 | 121 | 143402 | 4 | 4 | 3 | 3 | 0.09 | 1 | 1.031 | 1.04 | 0.748 | 0.737 | 0.886 | 1.024 | 0.788 | Exportin-1 OS=Mus musculus GN=Xpo1 PE=1 SV=1 |
| 436 | 1 | UniProt | n | Q9CZU6 | 179 | 60451 | 4 | 4 | 3 | 3 | 0.23 | 1 | 0.948 | 0.98 | 0.729 | 0.815 | 0.822 | 0.976 | 0.788 | Citrate synthase, mitochondrial OS=Mus musculus GN=Cs PE=1 SV=1 |
| 424 | 1 | UniProt | n | A0A0R4J1 | 189 | 43822 | 5 | 5 | 4 | 4 | 0.47 | 1 | 0.99 | 0.996 | 0.76 | 0.802 | 0.801 | 0.995 | 0.787 | Thioredoxin domain-containing protein 5 OS=Mus musculus GN=Txndc5 PE=1 SV=1 |
| 905 | 1 | UniProt | n | Q921K2 | 54 | 151670 | 2 | 2 | 2 | 2 | 0.06 | 1 | 0.855 | 0.836 | 0.824 | 0.743 | 0.797 | 0.894 | 0.787 | Poly [ADP-ribose] polymerase OS=Mus musculus GN=Parp1 PE=1 SV=1 |
| 609 | 1 | UniProt | n | A0A1L1ST | 102 | 117487 | 6 | 6 | 4 | 4 | 0.16 | 1 | 1.076 | 1.046 | 0.829 | 0.775 | 0.759 | 1.040 | 0.787 | Interleukin enhancer-binding factor 3 OS=Mus musculus GN=Ilf3 PE=1 SV=1 |
| 225 | 1 | UniProt | n | A2AVJ7 | 386 | 221639 | 17 | 17 | 10 | 10 | 0.23 | 1 | 0.944 | 0.966 | 0.916 | 0.758 | 0.702 | 0.970 | 0.787 | Ribosome-binding protein 1 OS=Mus musculus GN=Rrbp1 PE=1 SV=1 |
| 318 | 1 | UniProt | n | A0A0N4S\ | 272 | 23448 | 16 | 16 | 6 | 6 | 1.92 | 1 | 0.943 | 0.849 | 0.807 | 0.759 | 0.795 | 0.929 | 0.787 | Cellular nucleic acid-binding protein OS=Mus musculus GN=Cnbp PE=1 SV=1 |
| 444 | 1 | UniProt | n | P62897 | 175 | 17470 | 7 | 7 | 3 | 3 | 1.05 | 1 | 0.98 | 0.789 | 0.739 | 0.816 | 0.807 | 0.918 | 0.787 | Cytochrome c, somatic OS=Mus musculus GN=Cycs PE=1 SV=2 |
| 686 | 1 | UniProt | n | Q3UHX2 | 85 | 29110 | 2 | 2 | 1 | 1 | 0.33 | 1 | 1.09 | 1.141 | 0.786 | 0.815 | 0.758 | 1.075 | 0.786 | 28 kDa heat- and acid-stable phosphoprotein OS=Mus musculus GN=Pdap1 PE=1 SV=1 |
| 576 | 1 | UniProt | n | Q62418 | 114 | 54984 | 4 | 4 | 4 | 4 | 0.36 | 1 | 0.834 | 0.856 | 0.772 | 0.846 | 0.743 | 0.894 | 0.786 | Drebrin-like protein OS=Mus musculus GN=Dbnl PE=1 SV=2 |
| 471 | 1 | UniProt | n | Q91V98 | 159 | 86412 | 6 | 6 | 3 | 3 | 0.16 | 1 | 0.904 | 0.954 | 0.856 | 0.804 | 0.705 | 0.952 | 0.786 | Endosialin OS=Mus musculus GN=Cd248 PE=1 SV=1 |
| 325 | 1 | UniProt | n | P68040 | 265 | 40898 | 14 | 14 | 7 | 7 | 1.06 | 1 | 0.999 | 1.078 | 0.788 | 0.786 | 0.783 | 1.025 | 0.786 | Receptor of activated protein C kinase 1 OS=Mus musculus GN=Rack1 PE=1 SV=3 |
| 1160 | 1 | UniProt | n | Q9CR95 | 31 | 35143 | 1 | 1 | 1 | 1 | 0.13 | 1 | 0.749 | 1.102 | 0.767 | 0.697 | 0.906 | 0.938 | 0.785 | Adaptin ear-binding coat-associated protein 1 OS=Mus musculus GN=Necap1 PE=1 SV=2 |
| 790 | 1 | UniProt | n | O54724 | 67 | 56095 | 2 | 2 | 2 | 2 | 0.16 | 1 | 0.97 | 0.97 | 0.733 | 0.825 | 0.797 | 0.980 | 0.784 | Caveolae-associated protein 1 OS=Mus musculus GN=Cavin1 PE=1 SV=1 |
| 1105 | 1 | UniProt | n | Q6PB44 | 35 | 209214 | 1 | 1 | 1 | 1 | 0.02 | 1 | 0.881 | 1.187 | 0.908 | 0.763 | 0.695 | 1.015 | 0.784 | Tyrosine-protein phosphatase non-receptor type 23 OS=Mus musculus GN=Ptpn23 PE=1 SV=2 |
| 420 | 1 | UniProt | n | P25206 | 193 | 110339 | 7 | 7 | 7 | 7 | 0.31 | 1 | 1.025 | 1.09 | 0.786 | 0.836 | 0.732 | 1.038 | 0.784 | DNA replication licensing factor MCM3 OS=Mus musculus GN=Mcm3 PE=1 SV=2 |
| 212 | 1 | UniProt | n | Q8VCG1 | 410 | 24418 | 10 | 10 | 4 | 4 | 0.99 | 1 | 0.936 | 0.924 | 0.835 | 0.802 | 0.718 | 0.953 | 0.783 | Deoxyuridine triphosphatase OS=Mus musculus GN=Dut PE=1 SV=1 |
| 153 | 1 | UniProt | n | P26039 | 655 | 321290 | 25 | 25 | 15 | 15 | 0.27 | 1 | 0.931 | 0.954 | 0.79 | 0.782 | 0.778 | 0.961 | 0.783 | Talin-1 OS=Mus musculus GN=Tln1 PE=1 SV=2 |
| 152 | 1 | UniProt | n | Q543K9 | 665 | 35818 | 20 | 20 | 7 | 7 | 2.23 | 1 | 0.955 | 0.96 | 0.77 | 0.792 | 0.787 | 0.971 | 0.783 | Purine nucleoside phosphorylase OS=Mus musculus GN=Pnp PE=1 SV=1 |
| 280 | 1 | UniProt | n | P05202 | 318 | 57133 | 17 | 17 | 10 | 10 | 1.09 | 1 | 0.931 | 0.894 | 0.785 | 0.771 | 0.792 | 0.941 | 0.783 | Aspartate aminotransferase, mitochondrial OS=Mus musculus GN=Got2 PE=1 SV=1 |
| 426 | 1 | UniProt | n | Q9Z2X1 | 185 | 51148 | 6 | 6 | 4 | 4 | 0.39 | 1 | 0.921 | 0.915 | 0.806 | 0.811 | 0.733 | 0.945 | 0.783 | Heterogeneous nuclear ribonucleoprotein F OS=Mus musculus GN=Hnrnpf PE=1 SV=3 |
| 644 | 1 | UniProt | n | Q9D892 | 91 | 26418 | 5 | 5 | 3 | 3 | 0.61 | 1 | 1.022 | 1.025 | 0.795 | 0.797 | 0.756 | 1.016 | 0.782 | Inosine triphosphate pyrophosphatase OS=Mus musculus GN=Itpa PE=1 SV=2 |
| 348 | 1 | UniProt | n | Q9ERG0 | 246 | 104896 | 10 | 10 | 6 | 6 | 0.38 | 1 | 0.897 | 0.931 | 0.812 | 0.78 | 0.756 | 0.942 | 0.782 | LIM domain and actin-binding protein 1 OS=Mus musculus GN=Lima1 PE=1 SV=3 |
| 70 | 1 | UniProt | n | Q9DBJ1 | 1458 | 34686 | 65 | 65 | 12 | 12 | 6.89 | 1 | 1.06 | 0.998 | 0.823 | 0.755 | 0.77 | 1.019 | 0.782 | Phosphoglycerate mutase 1 OS=Mus musculus GN=Pgam1 PE=1 SV=3 |
| 1172 | 1 | UniProt | n | Q8VEA4 | 30 | 18575 | 3 | 3 | 2 | 2 | 0.96 | 1 | 1.047 | 1.003 | 0.802 | 0.754 | 0.79 | 1.016 | 0.782 | Mitochondrial intermembrane space import and assembly protein 40 OS=Mus musculus GN=Chchd4 PE=1 SV=1 |
| 97 | 1 | UniProt | n | A0A0A6Y\ | 1064 | 11156 | 59 | 59 | 7 | 7 | 17.73 | 1 | 1.024 | 0.944 | 0.838 | 0.888 | 0.639 | 0.989 | 0.781 | MCG23377, isoform CRA_b OS=Mus musculus GN=Gm8797 PE=4 SV=1 |
| 169 | 1 | UniProt | n | D3Z1Z8 | 567 | 16589 | 41 | 41 | 5 | 5 | 3.51 | 1 | 1.048 | 0.977 | 0.793 | 0.798 | 0.751 | 1.008 | 0.780 | Stathmin (Fragment) OS=Mus musculus GN=Stmn1 PE=1 SV=1 |
| 623 | 1 | UniProt | n | P45878 | 99 | 19427 | 4 | 4 | 1 | 1 | 0.24 | 1 | 1.008 | 0.954 | 0.793 | 0.799 | 0.75 | 0.987 | 0.780 | Peptidyl-prolyl cis-trans isomerase FKBP2 OS=Mus musculus GN=Fkbp2 PE=1 SV=1 |
| 17 | 2 | UniProt | n | P20029 | 2463 | 91330 | 75 | 75 | 21 | 21 | 2.19 | 1 | 1.005 | 1.023 | 0.798 | 0.766 | 0.777 | 1.009 | 0.780 | 78 kDa glucose-regulated protein OS=Mus musculus GN=Hspa5 PE=1 SV=3 |
| 590 | 1 | UniProt | n | A0A0N4S\ | 107 | 16888 | 2 | 2 | 1 | 1 | 0.28 | 1 | 0.843 | 0.851 | 0.747 | 0.747 | 0.85 | 0.895 | 0.780 | Gamma-glutamylcyclotransferase (Fragment) OS=Mus musculus GN=Ggct PE=1 SV=1 |
| 689 | 1 | UniProt | n | Q3TCN2 | 84 | 73262 | 3 | 3 | 2 | 2 | 0.12 | 1 | 0.941 | 1.017 | 0.872 | 0.731 | 0.744 | 0.985 | 0.780 | Putative phospholipase B-like 2 OS=Mus musculus GN=Plbd2 PE=1 SV=2 |
| 219 | 1 | UniProt | n | A2AMW0 | 397 | 36808 | 22 | 22 | 9 | 9 | 1.8 | 1 | 1.045 | 1.057 | 0.773 | 0.794 | 0.772 | 1.034 | 0.780 | Capping protein (Actin filament) muscle Z-line, beta, isoform CRA_a OS=Mus musculus GN=Capzb PE=1 SV=1 |
| 310 | 1 | UniProt | n | Q62426 | 283 | 14173 | 11 | 11 | 4 | 4 | 3.29 | 1 | 0.994 | 0.893 | 0.705 | 0.882 | 0.762 | 0.961 | 0.780 | Cystatin-B OS=Mus musculus GN=Cstb PE=1 SV=1 |
| 1010 | 1 | UniProt | n | D3Z3J6 | 44 | 47786 | 3 | 3 | 2 | 2 | 0.19 | 1 | 0.836 | 0.896 | 0.838 | 0.794 | 0.712 | 0.908 | 0.780 | Polyadenylate-binding protein-interacting protein 1 OS=Mus musculus GN=Paip1 PE=1 SV=1 |
| 1053 | 1 | UniProt | n | P51660 | 40 | 95056 | 3 | 3 | 2 | 2 | 0.09 | 1 | 0.866 | 0.866 | 0.768 | 0.767 | 0.804 | 0.909 | 0.779 | Peroxisomal multifunctional enzyme type 2 OS=Mus musculus GN=Hsd17b4 PE=1 SV=3 |
| 421 | 1 | UniProt | n | Q8VEK3 | 192 | 109193 | 6 | 6 | 3 | 3 | 0.12 | 1 | 0.953 | 0.924 | 0.705 | 0.781 | 0.86 | 0.958 | 0.779 | Heterogeneous nuclear ribonucleoprotein U OS=Mus musculus GN=Hnrnpu PE=1 SV=1 |
| 422 | 1 | UniProt | n | Q91VW3 | 191 | 12037 | 3 | 3 | 1 | 1 | 0.41 | 1 | 0.937 | 0.851 | 0.749 | 0.898 | 0.704 | 0.927 | 0.779 | SH3 domain-binding glutamic acid-rich-like protein 3 OS=Mus musculus GN=Sh3bgrl3 PE=1 SV=1 |
| 35 | 1 | UniProt | n | E9Q8I0 | 2790 | 171416 | 101 | 101 | 33 | 33 | 1.76 | 1 | 1.044 | 1.044 | 0.772 | 0.755 | 0.812 | 1.029 | 0.779 | Complement factor H OS=Mus musculus GN=Cfh PE=1 SV=1 |
| 458 | 1 | UniProt | n | Q61792 | 168 | 36990 | 5 | 5 | 2 | 2 | 0.41 | 1 | 0.979 | 0.972 | 0.759 | 0.751 | 0.83 | 0.984 | 0.779 | LIM and SH3 domain protein 1 OS=Mus musculus GN=Lasp1 PE=1 SV=1 |
| 320 | 1 | UniProt | n | P51859 | 270 | 34862 | 15 | 15 | 7 | 7 | 1.33 | 1 | 1.028 | 1.043 | 0.774 | 0.781 | 0.782 | 1.024 | 0.779 | Hepatoma-derived growth factor OS=Mus musculus GN=Hdgf PE=1 SV=2 |
| 62 | 2 | UniProt | n | P28654 | 1383 | 47969 | 71 | 71 | 13 | 13 | 2.43 | 1 | 1.133 | 1.102 | 0.794 | 0.761 | 0.782 | 1.077 | 0.779 | Decorin OS=Mus musculus GN=Dcn PE=1 SV=1 |
| 139 | 1 | UniProt | n | Q61171 | 708 | 26162 | 29 | 29 | 7 | 7 | 2.6 | 1 | 1.033 | 1.002 | 0.791 | 0.79 | 0.756 | 1.012 | 0.779 | Peroxiredoxin-2 OS=Mus musculus GN=Prdx2 PE=1 SV=3 |
| 1225 | 1 | UniProt | n | Q3UMU9 | 26 | 96896 | 1 | 1 | 1 | 1 | 0.04 | 1 | 1.046 | 1.49 | 0.767 | 0.709 | 0.868 | 1.159 | 0.779 | Hepatoma-derived growth factor-related protein 2 OS=Mus musculus GN=Hdgfl2 PE=1 SV=1 |
| 1248 | 1 | UniProt | n | Q6IRU5 | 23 | 30724 | 1 | 1 | 1 | 1 | 0.15 | 1 | 0.969 | 0.959 | 0.794 | 0.688 | 0.864 | 0.976 | 0.779 | Clathrin light chain B OS=Mus musculus GN=Cltb PE=1 SV=1 |
| 563 | 1 | UniProt | n | A0A0G2JE | 118 | 17035 | 5 | 5 | 2 | 2 | 0.63 | 1 | 1.053 | 1.045 | 0.788 | 0.759 | 0.786 | 1.032 | 0.778 | Ubiquitin-conjugating enzyme E2 D3 (Fragment) OS=Mus musculus GN=Ube2d3 PE=1 SV=1 |
| 303 | 1 | UniProt | n | Q8BP92 | 291 | 42419 | 7 | 7 | 3 | 3 | 0.49 | 1 | 0.914 | 1.067 | 0.783 | 0.811 | 0.74 | 0.992 | 0.777 | Reticulocalbin-2 OS=Mus musculus GN=Rcn2 PE=1 SV=1 |
| 275 | 1 | UniProt | n | Q8BG13 | 324 | 17968 | 10 | 10 | 3 | 3 | 2.18 | 1 | 0.981 | 0.896 | 0.758 | 0.781 | 0.793 | 0.958 | 0.777 | RNA-binding protein 3 OS=Mus musculus GN=Rbm3 PE=1 SV=1 |
| 710 | 1 | UniProt | n | H3BJA3 | 80 | 26277 | 2 | 2 | 2 | 2 | 0.38 | 1 | 0.91 | 0.845 | 0.712 | 0.799 | 0.825 | 0.916 | 0.777 | Glutamate--cysteine ligase regulatory subunit (Fragment) OS=Mus musculus GN=Gclm PE=1 SV=1 |
| 1181 | 1 | UniProt | n | P97807 | 30 | 64545 | 1 | 1 | 1 | 1 | 0.07 | 1 | 1.232 | 1.203 | 0.63 | 0.93 | 0.801 | 1.140 | 0.777 | Fumarate hydratase, mitochondrial OS=Mus musculus GN=Fh PE=1 SV=3 |
| 218 | 1 | UniProt | n | P49722 | 401 | 30261 | 13 | 13 | 7 | 7 | 1.64 | 1 | 1.081 | 1.026 | 0.737 | 0.784 | 0.812 | 1.035 | 0.777 | Proteasome subunit alpha type-2 OS=Mus musculus GN=Psma2 PE=1 SV=3 |
| 960 | 1 | UniProt | n | Q9D0S9 | 48 | 19789 | 1 | 1 | 1 | 1 | 0.24 | 1 | 0.676 | 0.574 | 0.633 | 1.068 | 0.694 | 0.729 | 0.777 | Histidine triad nucleotide-binding protein 2, mitochondrial OS=Mus musculus GN=Hint2 PE=1 SV=1 |
| 195 | 1 | UniProt | n | Q61001 | 458 | 439324 | 10 | 10 | 7 | 7 | 0.07 | 1 | 0.987 | 1.004 | 0.748 | 0.781 | 0.803 | 0.997 | 0.777 | Laminin subunit alpha-5 OS=Mus musculus GN=Lama5 PE=1 SV=4 |
| 241 | 1 | UniProt | n | A0A1C7C\ | 360 | 86526 | 19 | 19 | 13 | 13 | 0.98 | 1 | 1 | 1.043 | 0.809 | 0.751 | 0.772 | 1.014 | 0.777 | Plastin-3 (Fragment) OS=Mus musculus GN=Pls3 PE=1 SV=1 |
| 1083 | 1 | UniProt | n | Q9JHU4 | 38 | 624570 | 1 | 1 | 1 | 1 | 0.01 | 1 | 0.764 | 0.898 | 0.956 | 0.621 | 0.79 | 0.882 | 0.777 | Cytoplasmic dynein 1 heavy chain 1 OS=Mus musculus GN=Dync1h1 PE=1 SV=2 |
| 817 | 1 | UniProt | n | Q9Z1F9 | 63 | 85678 | 2 | 2 | 2 | 2 | 0.1 | 1 | 0.928 | 1.09 | 0.776 | 0.79 | 0.765 | 1.004 | 0.777 | SUMO-activating enzyme subunit 2 OS=Mus musculus GN=Uba2 PE=1 SV=1 |
| 513 | 2 | UniProt | n | P17426 | 76 | 125505 | 3 | 3 | 3 | 3 | 0.11 | 1 | 0.949 | 0.951 | 0.816 | 0.784 | 0.732 | 0.966 | 0.777 | AP-2 complex subunit alpha-1 OS=Mus musculus GN=Ap2a1 PE=1 SV=1 |
| 32 | 1 | UniProt | n | Q02788 | 2963 | 127308 | 120 | 120 | 30 | 30 | 2.43 | 1 | 1.005 | 0.972 | 0.79 | 0.76 | 0.779 | 0.992 | 0.776 | Collagen alpha-2(VI) chain OS=Mus musculus GN=Col6a2 PE=1 SV=3 |
| 1207 | 1 | UniProt | n | A2AA71 | 27 | 128912 | 1 | 1 | 1 | 1 | 0.03 | 1 | 0.944 | 0.967 | 0.789 | 0.753 | 0.785 | 0.970 | 0.775 | Protein transport protein Sec24A OS=Mus musculus GN=Sec24a PE=1 SV=1 |
| 234 | 1 | UniProt | n | P42932 | 372 | 73362 | 19 | 19 | 15 | 15 | 1.38 | 1 | 0.939 | 0.984 | 0.816 | 0.762 | 0.75 | 0.974 | 0.775 | T-complex protein 1 subunit theta OS=Mus musculus GN=Cct8 PE=1 SV=3 |
| 250 | 1 | UniProt | n | Q3U0V1 | 352 | 86830 | 14 | 14 | 9 | 9 | 0.63 | 1 | 0.999 | 0.982 | 0.797 | 0.777 | 0.753 | 0.994 | 0.775 | Far upstream element-binding protein 2 OS=Mus musculus GN=Khsrp PE=1 SV=2 |
| 86 | 2 | UniProt | n | Q6IRU2 | 1133 | 36452 | 51 | 51 | 13 | 13 | 4.05 | 1 | 1.086 | 1.072 | 0.821 | 0.799 | 0.71 | 1.052 | 0.775 | Tropomyosin alpha-4 chain OS=Mus musculus GN=Tpm4 PE=1 SV=3 |
| 876 | 1 | UniProt | n | P98083 | 57 | 70283 | 2 | 2 | 1 | 1 | 0.06 | 1 | 0.998 | 0.994 | 0.85 | 0.735 | 0.745 | 0.997 | 0.775 | SHC-transforming protein 1 OS=Mus musculus GN=Shc1 PE=1 SV=3 |
| 15 | 1 | UniProt | n | Q8VDD5 | 4946 | 289864 | 215 | 215 | 70 | 70 | 2.53 | 1 | 1.036 | 1.071 | 0.789 | 0.765 | 0.771 | 1.035 | 0.775 | Myosin-9 OS=Mus musculus GN=Myh9 PE=1 SV=4 |
| 106 | 2 | UniProt | n | O08807 | 321 | 36389 | 21 | 21 | 6 | 6 | 1 | 1 | 1.007 | 0.936 | 0.75 | 0.795 | 0.78 | 0.980 | 0.775 | Peroxiredoxin-4 OS=Mus musculus GN=Prdx4 PE=1 SV=1 |
| 973 | 1 | UniProt | n | Q9D8S9 | 47 | 16029 | 1 | 1 | 1 | 1 | 0.3 | 1 | 0.847 | 0.915 | 0.816 | 0.822 | 0.693 | 0.919 | 0.775 | BolA-like protein 1 OS=Mus musculus GN=Bola1 PE=1 SV=1 |
| 1128 | 1 | UniProt | n | Q3U128 | 33 | 64370 | 1 | 1 | 1 | 1 | 0.07 | 1 | 0.968 | 1.108 | 0.758 | 0.724 | 0.847 | 1.024 | 0.775 | Protein APCDD1 OS=Mus musculus GN=Apcdd1 PE=2 SV=1 |
| 120 | 1 | UniProt | n | F8WIV2 | 832 | 55152 | 37 | 37 | 14 | 14 | 3.29 | 1 | 1.023 | 0.977 | 0.796 | 0.75 | 0.777 | 1.000 | 0.774 | Serine (or cysteine) peptidase inhibitor, clade B, member 6a OS=Mus musculus GN=Serpinb6a PE=1 SV=1 |
| 243 | 1 | UniProt | n | P34884 | 358 | 13851 | 15 | 15 | 3 | 3 | 2.29 | 1 | 0.999 | 0.897 | 0.855 | 0.771 | 0.703 | 0.964 | 0.774 | Macrophage migration inhibitory factor OS=Mus musculus GN=Mif PE=1 SV=2 |
| 175 | 2 | UniProt | n | Q5RKN9 | 452 | 39155 | 14 | 14 | 7 | 7 | 1.12 | 1 | 0.973 | 0.942 | 0.76 | 0.765 | 0.794 | 0.971 | 0.773 | Capping protein (Actin filament) muscle Z-line, alpha 1 OS=Mus musculus GN=Capza1 PE=1 SV=1 |
| 40 | 1 | UniProt | n | A0A1L1SV | 2230 | 124749 | 91 | 91 | 36 | 36 | 3.03 | 1 | 1.039 | 1.091 | 0.779 | 0.747 | 0.793 | 1.043 | 0.773 | Alpha-actinin-4 OS=Mus musculus GN=Actn4 PE=1 SV=1 |
| 198 | 1 | UniProt | n | Q5SVG4 | 452 | 121688 | 25 | 25 | 13 | 13 | 0.75 | 1 | 1.02 | 1.088 | 0.756 | 0.79 | 0.77 | 1.035 | 0.772 | AP complex subunit beta OS=Mus musculus GN=Ap1b1 PE=1 SV=1 |
| 521 | 1 | UniProt | n | P62141 | 132 | 43891 | 4 | 4 | 3 | 3 | 0.33 | 1 | 0.973 | 0.903 | 0.772 | 0.804 | 0.74 | 0.958 | 0.772 | Serine/threonine-protein phosphatase PP1-beta catalytic subunit OS=Mus musculus GN=Ppp1cb PE=1 SV=3 |
| 237 | 1 | UniProt | n | Q9JJU8 | 368 | 15329 | 11 | 11 | 6 | 6 | 4.07 | 1 | 1.017 | 1.007 | 0.756 | 0.739 | 0.822 | 1.008 | 0.772 | SH3 domain-binding glutamic acid-rich-like protein OS=Mus musculus GN=Sh3bgrl PE=1 SV=1 |
| 738 | 1 | UniProt | n | Q9JHW2 | 75 | 36843 | 2 | 2 | 2 | 2 | 0.26 | 1 | 0.99 | 1.128 | 0.834 | 0.775 | 0.71 | 1.037 | 0.771 | Omega-amidase NIT2 OS=Mus musculus GN=Nit2 PE=1 SV=1 |
| 317 | 1 | UniProt | n | P63005 | 273 | 57747 | 14 | 14 | 8 | 8 | 0.8 | 1 | 0.93 | 1.026 | 0.793 | 0.801 | 0.722 | 0.984 | 0.771 | Platelet-activating factor acetylhydrolase IB subunit alpha OS=Mus musculus GN=Pafah1b1 PE=1 SV=2 |
| 1026 | 1 | UniProt | n | A0A0G2JG | 43 | 22022 | 1 | 1 | 1 | 1 | 0.21 | 1 | 0.863 | 0.944 | 0.796 | 0.839 | 0.686 | 0.934 | 0.771 | 26S proteasome non-ATPase regulatory subunit 9 (Fragment) OS=Mus musculus GN=Psmd9 PE=1 SV=1 |
| 1037 | 1 | UniProt | n | O89051 | 41 | 36738 | 1 | 1 | 1 | 1 | 0.12 | 1 | 0.925 | 1.13 | 0.846 | 0.807 | 0.671 | 1.015 | 0.771 | Integral membrane protein 2B OS=Mus musculus GN=Itm2b PE=1 SV=1 |
| 720 | 1 | UniProt | n | A0A0G2JD | 78 | 18468 | 2 | 2 | 2 | 2 | 0.57 | 1 | 0.611 | 0.777 | 0.717 | 0.838 | 0.762 | 0.780 | 0.771 | MCG10748, isoform CRA_b OS=Mus musculus GN=Rap1a PE=1 SV=1 |
| 104 | 1 | UniProt | n | Q3V117 | 991 | 144269 | 33 | 33 | 22 | 22 | 0.97 | 1 | 0.983 | 1.114 | 0.762 | 0.755 | 0.794 | 1.031 | 0.770 | ATP-citrate synthase OS=Mus musculus GN=Acly PE=1 SV=1 |
| 12 | 2 | UniProt | n | P07901 | 4496 | 109697 | 167 | 167 | 28 | 28 | 2.58 | 1 | 1.084 | 1.118 | 0.786 | 0.76 | 0.764 | 1.066 | 0.770 | Heat shock protein HSP 90-alpha OS=Mus musculus GN=Hsp90aa1 PE=1 SV=4 |
| 183 | 1 | UniProt | n | Q921M3 | 506 | 152461 | 20 | 20 | 12 | 12 | 0.4 | 1 | 0.934 | 0.969 | 0.791 | 0.756 | 0.763 | 0.967 | 0.770 | Splicing factor 3B subunit 3 OS=Mus musculus GN=Sf3b3 PE=1 SV=1 |
| 597 | 1 | UniProt | n | P15864 | 104 | 39202 | 6 | 6 | 2 | 2 | 0.24 | 1 | 1.054 | 1.026 | 0.762 | 0.741 | 0.808 | 1.026 | 0.770 | Histone H1.2 OS=Mus musculus GN=Hist1h1c PE=1 SV=2 |
| 423 | 1 | UniProt | n | P70195 | 191 | 36232 | 8 | 8 | 4 | 4 | 0.79 | 1 | 0.984 | 0.999 | 0.722 | 0.932 | 0.678 | 0.994 | 0.770 | Proteasome subunit beta type-7 OS=Mus musculus GN=Psmb7 PE=1 SV=1 |
| 389 | 1 | UniProt | n | Q8BFR4 | 214 | 70860 | 15 | 15 | 8 | 8 | 0.61 | 1 | 0.973 | 1.008 | 0.786 | 0.764 | 0.759 | 0.994 | 0.770 | N-acetylglucosamine-6-sulfatase OS=Mus musculus GN=Gns PE=1 SV=1 |
| 1030 | 1 | UniProt | n | A0A0R4J1 | 42 | 36889 | 2 | 2 | 2 | 2 | 0.26 | 1 | 0.804 | 0.846 | 0.801 | 0.789 | 0.721 | 0.879 | 0.770 | Putative deoxyribonuclease TATDN1 OS=Mus musculus GN=Tatdn1 PE=1 SV=1 |
| 315 | 2 | UniProt | n | A0A0U1RI | 69 | 43188 | 3 | 3 | 3 | 3 | 0.34 | 1 | 1.06 | 1.122 | 0.842 | 0.726 | 0.745 | 1.059 | 0.769 | Mitogen-activated protein kinase (Fragment) OS=Mus musculus GN=Mapk3 PE=1 SV=1 |
| 974 | 1 | UniProt | n | Q9CX34 | 47 | 48404 | 1 | 1 | 1 | 1 | 0.09 | 1 | 1.168 | 1.044 | 0.586 | 0.89 | 0.873 | 1.068 | 0.769 | Protein SGT1 homolog OS=Mus musculus GN=Sugt1 PE=1 SV=3 |
| 178 | 1 | UniProt | n | P40124 | 526 | 64281 | 16 | 16 | 8 | 8 | 0.69 | 1 | 0.994 | 1.019 | 0.798 | 0.727 | 0.784 | 1.004 | 0.769 | Adenylyl cyclase-associated protein 1 OS=Mus musculus GN=Cap1 PE=1 SV=4 |
| 542 | 1 | UniProt | n | E9Q3T0 | 125 | 13951 | 3 | 3 | 2 | 2 | 0.81 | 1 | 0.988 | 0.975 | 0.764 | 0.733 | 0.812 | 0.988 | 0.769 | Predicted pseudogene 10073 OS=Mus musculus GN=Gm10073 PE=1 SV=1 |
| 530 | 1 | UniProt | n | Q8BV57 | 129 | 156562 | 5 | 5 | 5 | 5 | 0.15 | 1 | 0.885 | 0.87 | 0.798 | 0.793 | 0.718 | 0.917 | 0.769 | Soluble scavenger receptor cysteine-rich domain-containing protein SSC5D OS=Mus musculus GN=Ssc5d PE=1 SV=1 |
| 1274 | 1 | UniProt | n | Q9D5T7-2 | 19 | 55115 | 1 | 1 | 1 | 1 | 0.08 | 1 | 0.632 | 0.492 | --- | 0.601 | 0.983 | 0.677 | 0.769 | Isoform 2 of HORMA domain-containing protein 1 OS=Mus musculus GN=Hormad1 |
| 1032 | 1 | UniProt | n | P50516 | 42 | 79510 | 2 | 2 | 2 | 2 | 0.11 | 1 | 0.885 | 0.826 | 0.74 | 0.741 | 0.828 | 0.901 | 0.769 | V-type proton ATPase catalytic subunit A OS=Mus musculus GN=Atp6v1a PE=1 SV=2 |
| 747 | 1 | UniProt | n | Q8BTI8 | 74 | 337336 | 3 | 3 | 3 | 3 | 0.04 | 1 | 0.766 | 0.832 | 0.79 | 0.799 | 0.719 | 0.861 | 0.768 | Serine/arginine repetitive matrix protein 2 OS=Mus musculus GN=Srrm2 PE=1 SV=3 |
| 264 | 1 | UniProt | n | P06745 | 331 | 75688 | 13 | 13 | 6 | 6 | 0.56 | 1 | 1.033 | 1.061 | 0.8 | 0.747 | 0.757 | 1.031 | 0.768 | Glucose-6-phosphate isomerase OS=Mus musculus GN=Gpi PE=1 SV=4 |
| 791 | 1 | UniProt | n | Q9QZQ8 | 67 | 54450 | 1 | 1 | 1 | 1 | 0.08 | 1 | 1.094 | 1.013 | 0.835 | 0.755 | 0.717 | 1.035 | 0.767 | Core histone macro-H2A.1 OS=Mus musculus GN=H2afy PE=1 SV=3 |
| 908 | 1 | UniProt | n | Q99JI4 | 54 | 53951 | 2 | 2 | 2 | 2 | 0.17 | 1 | 0.855 | 0.834 | 0.934 | 0.741 | 0.653 | 0.893 | 0.767 | 26S proteasome non-ATPase regulatory subunit 6 OS=Mus musculus GN=Psmd6 PE=1 SV=1 |
| 1132 | 1 | UniProt | n | A0A0G2JG | 33 | 71215 | 1 | 1 | 1 | 1 | 0.06 | 1 | 0.855 | 1.007 | 0.824 | 0.788 | 0.696 | 0.951 | 0.767 | RAP1, GTP-GDP dissociation stimulator 1 OS=Mus musculus GN=Rap1gds1 PE=1 SV=1 |
| 288 | 1 | UniProt | n | Q8BGJ5 | 305 | 67337 | 10 | 10 | 5 | 5 | 0.55 | 1 | 1.114 | 1.103 | 0.766 | 0.758 | 0.778 | 1.071 | 0.767 | MCG13402, isoform CRA_a OS=Mus musculus GN=Ptbp1 PE=1 SV=1 |
| 12 | 1 | UniProt | n | P11499 | 5284 | 106625 | 200 | 200 | 33 | 33 | 4.31 | 1 | 1.082 | 1.112 | 0.776 | 0.755 | 0.771 | 1.064 | 0.767 | Heat shock protein HSP 90-beta OS=Mus musculus GN=Hsp90ab1 PE=1 SV=3 |
| 698 | 1 | UniProt | n | P97377 | 83 | 46622 | 3 | 3 | 3 | 3 | 0.31 | 1 | 1.002 | 0.981 | 0.749 | 0.781 | 0.772 | 0.994 | 0.767 | Cyclin-dependent kinase 2 OS=Mus musculus GN=Cdk2 PE=1 SV=2 |
| 447 | 1 | UniProt | n | Q8BPW9 | 174 | 56652 | 8 | 8 | 6 | 6 | 0.57 | 1 | 0.982 | 1.023 | 0.828 | 0.767 | 0.71 | 1.002 | 0.767 | Aspartyl aminopeptidase OS=Mus musculus GN=Dnpep PE=1 SV=1 |
| 1112 | 1 | UniProt | n | Q07113 | 35 | 321580 | 1 | 1 | 1 | 1 | 0.01 | 1 | 0.812 | 0.994 | 0.819 | 0.691 | 0.796 | 0.931 | 0.767 | Cation-independent mannose-6-phosphate receptor OS=Mus musculus GN=Igf2r PE=1 SV=1 |
| 17 | 3 | UniProt | n | P17156 | 1689 | 85952 | 64 | 64 | 9 | 9 | 0.64 | 1 | 0.979 | 0.997 | 0.794 | 0.739 | 0.767 | 0.992 | 0.766 | Heat shock-related 70 kDa protein 2 OS=Mus musculus GN=Hspa2 PE=1 SV=2 |
| 559 | 1 | UniProt | n | Q9QZF2 | 119 | 69495 | 5 | 5 | 3 | 3 | 0.2 | 1 | 0.681 | 0.7 | 0.788 | 0.751 | 0.758 | 0.781 | 0.765 | Glypican-1 OS=Mus musculus GN=Gpc1 PE=1 SV=1 |
| 1005 | 1 | UniProt | n | Q8K003 | 44 | 13450 | 3 | 3 | 1 | 1 | 0.36 | 1 | 0.986 | 0.99 | 0.755 | 0.795 | 0.747 | 0.992 | 0.765 | Translation machinery-associated protein 7 OS=Mus musculus GN=Tma7 PE=3 SV=1 |
| 32 | 2 | UniProt | n | D3Z7D5 | 2492 | 115923 | 95 | 95 | 22 | 22 | 1.69 | 1 | 1.006 | 0.959 | 0.771 | 0.754 | 0.77 | 0.988 | 0.765 | Collagen alpha-2(VI) chain OS=Mus musculus GN=Col6a2 PE=1 SV=1 |
| 387 | 1 | UniProt | n | Q99KK7 | 218 | 93815 | 4 | 4 | 3 | 3 | 0.15 | 1 | 0.816 | 0.882 | 0.81 | 0.744 | 0.742 | 0.896 | 0.765 | Dipeptidyl peptidase 3 OS=Mus musculus GN=Dpp3 PE=1 SV=2 |
| 1250 | 1 | UniProt | n | Q8CGF7 | 23 | 154010 | 1 | 1 | 1 | 1 | 0.03 | 1 | 1.096 | 0.937 | 0.843 | 0.714 | 0.742 | 1.009 | 0.764 | Transcription elongation regulator 1 OS=Mus musculus GN=Tcerg1 PE=1 SV=2 |
| 1117 | 1 | UniProt | n | Q9D8T2 | 34 | 62016 | 1 | 1 | 1 | 1 | 0.07 | 1 | 1.227 | 0.985 | 0.67 | 0.868 | 0.767 | 1.065 | 0.764 | Gasdermin-D OS=Mus musculus GN=Gsdmdc1 PE=1 SV=1 |
| 1108 | 1 | UniProt | n | E9Q2T3 | 35 | 373334 | 3 | 3 | 3 | 3 | 0.03 | 1 | 0.95 | 1.025 | 0.808 | 0.749 | 0.737 | 0.991 | 0.764 | Tenascin XB OS=Mus musculus GN=Tnxb PE=1 SV=1 |
| 821 | 1 | UniProt | n | D3YVI6 | 62 | 22000 | 1 | 1 | 1 | 1 | 0.21 | 1 | 0.996 | 1.051 | 0.782 | 0.792 | 0.718 | 1.015 | 0.763 | COP9 signalosome complex subunit 7a (Fragment) OS=Mus musculus GN=Cops7a PE=1 SV=1 |
| 274 | 1 | UniProt | n | Q8CFZ6 | 324 | 28039 | 14 | 14 | 4 | 4 | 1.11 | 1 | 1.027 | 1.05 | 0.774 | 0.808 | 0.711 | 1.025 | 0.763 | C-type lectin domain family 3, member b OS=Mus musculus GN=Clec3b PE=1 SV=1 |
| 1035 | 1 | UniProt | n | A2API8 | 42 | 119290 | 1 | 1 | 1 | 1 | 0.04 | 1 | 0.866 | 0.966 | 0.742 | 0.77 | 0.778 | 0.942 | 0.763 | A-kinase anchor protein 2 OS=Mus musculus GN=Akap2 PE=1 SV=1 |
| 767 | 1 | UniProt | n | P10711 | 71 | 44266 | 3 | 3 | 3 | 3 | 0.33 | 1 | 0.985 | 0.948 | 0.78 | 0.764 | 0.745 | 0.977 | 0.763 | Transcription elongation factor A protein 1 OS=Mus musculus GN=Tcea1 PE=1 SV=2 |
| 357 | 1 | UniProt | n | O09131 | 237 | 34661 | 16 | 16 | 8 | 8 | 1.64 | 1 | 1.008 | 1.074 | 0.757 | 0.779 | 0.752 | 1.027 | 0.763 | Glutathione S-transferase omega-1 OS=Mus musculus GN=Gsto1 PE=1 SV=2 |
| 940 | 1 | UniProt | n | P35564 | 50 | 85506 | 1 | 1 | 1 | 1 | 0.05 | 1 | 1.048 | 1.078 | 0.67 | 0.701 | 0.944 | 1.042 | 0.763 | Calnexin OS=Mus musculus GN=Canx PE=1 SV=1 |
| 434 | 1 | UniProt | n | E9QKZ2 | 180 | 129481 | 6 | 6 | 5 | 5 | 0.18 | 1 | 0.949 | 1.122 | 0.801 | 0.773 | 0.716 | 1.021 | 0.763 | Importin-9 OS=Mus musculus GN=Ipo9 PE=1 SV=1 |
| 898 | 1 | UniProt | n | A0A0G2JG | 55 | 98637 | 1 | 1 | 1 | 1 | 0.04 | 1 | 1.036 | 0.873 | 0.832 | 0.721 | 0.738 | 0.967 | 0.762 | Serine/threonine-protein kinase DCLK1 OS=Mus musculus GN=Dclk1 PE=1 SV=1 |
| 495 | 1 | UniProt | n | A0A1W2P | 148 | 27642 | 7 | 7 | 6 | 6 | 1.49 | 1 | 1 | 1.015 | 0.822 | 0.715 | 0.753 | 1.005 | 0.762 | Small glutamine-rich tetratricopeptide repeat-containing protein alpha (Fragment) OS=Mus musculus GN=Sgta PE=1 SV=1 |
| 797 | 1 | UniProt | n | Q3U7R1 | 66 | 138089 | 3 | 3 | 3 | 3 | 0.1 | 1 | 0.944 | 0.993 | 0.766 | 0.773 | 0.747 | 0.979 | 0.762 | Extended synaptotagmin-1 OS=Mus musculus GN=Esyt1 PE=1 SV=2 |
| 199 | 1 | UniProt | n | A0A1B0G! | 452 | 62009 | 13 | 13 | 8 | 8 | 0.73 | 1 | 0.994 | 1.035 | 0.811 | 0.755 | 0.722 | 1.010 | 0.762 | Ribonuclease inhibitor OS=Mus musculus GN=Rnh1 PE=1 SV=1 |
| 19 | 1 | UniProt | n | P48678 | 4634 | 86591 | 223 | 223 | 38 | 38 | 7.61 | 1 | 0.983 | 0.893 | 0.796 | 0.766 | 0.725 | 0.957 | 0.762 | Prelamin-A/C OS=Mus musculus GN=Lmna PE=1 SV=2 |
| 583 | 1 | UniProt | n | Q9ER72 | 111 | 115688 | 3 | 3 | 3 | 3 | 0.12 | 1 | 0.883 | 0.945 | 0.792 | 0.736 | 0.758 | 0.941 | 0.762 | Cysteine--tRNA ligase, cytoplasmic OS=Mus musculus GN=Cars PE=1 SV=2 |
| 412 | 1 | UniProt | n | G3X9V0 | 200 | 32669 | 7 | 7 | 5 | 5 | 0.9 | 1 | 1.004 | 1.087 | 0.764 | 0.766 | 0.755 | 1.030 | 0.762 | MCG22048, isoform CRA_a OS=Mus musculus GN=Psme2 PE=1 SV=1 |

| | | | | | | | | | | | | | | | | | | |
|---|---|---|---|---|---|---|---|---|---|---|---|---|---|---|---|---|---|---|
| 942 | 1 | UniProt n | D6RHA2 | 50 | 55022 | 1 | 1 | 1 | 1 | 0.08 | 1 | 0.787 | 0.878 | 0.824 | 0.762 | 0.703 | 0.884 | 0.761 | N-acylglucosamine 2-epimerase OS=Mus musculus GN=Renbp PE=1 SV=2 |
| 282 | 1 | UniProt n | Q9QUN9 | 315 | 42066 | 11 | 11 | 6 | 6 | 0.82 | 1 | 0.902 | 0.87 | 0.818 | 0.735 | 0.734 | 0.922 | 0.761 | Dickkopf-related protein 3 OS=Mus musculus GN=Dkk3 PE=2 SV=1 |
| 187 | 2 | UniProt n | P10630 | 315 | 53553 | 11 | 11 | 6 | 6 | 0.74 | 1 | 1.04 | 1.041 | 0.755 | 0.717 | 0.815 | 1.027 | 0.761 | Eukaryotic initiation factor 4A-II OS=Mus musculus GN=Eif4a2 PE=1 SV=2 |
| 187 | 1 | UniProt n | P60843 | 491 | 52697 | 17 | 17 | 9 | 9 | 1.23 | 1 | 1.028 | 1.029 | 0.752 | 0.717 | 0.817 | 1.019 | 0.761 | Eukaryotic initiation factor 4A-I OS=Mus musculus GN=Eif4a1 PE=1 SV=1 |
| 522 | 1 | UniProt n | G3UY93 | 131 | 161908 | 3 | 3 | 3 | 3 | 0.08 | 1 | 0.986 | 1.065 | 0.759 | 0.744 | 0.78 | 1.016 | 0.761 | Valine--tRNA ligase (Fragment) OS=Mus musculus GN=Vars PE=1 SV=1 |
| 1186 | 1 | UniProt n | Q6Q899 | 29 | 132003 | 1 | 1 | 1 | 1 | 0.03 | 1 | 0.74 | 0.883 | 0.737 | 0.822 | 0.725 | 0.868 | 0.760 | Probable ATP-dependent RNA helicase DDX58 OS=Mus musculus GN=Ddx58 PE=1 SV=2 |
| 782 | 1 | UniProt n | Q9Z247 | 68 | 70624 | 4 | 4 | 2 | 2 | 0.13 | 1 | 0.994 | 1.012 | 0.77 | 0.779 | 0.732 | 1.002 | 0.760 | Peptidyl-prolyl cis-trans isomerase FKBP9 OS=Mus musculus GN=Fkbp9 PE=1 SV=1 |
| 192 | 1 | UniProt n | Q9Z1N5 | 470 | 58498 | 20 | 20 | 10 | 10 | 1.06 | 1 | 1.026 | 1.058 | 0.774 | 0.737 | 0.769 | 1.028 | 0.760 | Spliceosome RNA helicase Ddx39b OS=Mus musculus GN=Ddx39b PE=1 SV=1 |
| 515 | 1 | UniProt n | Q8CI94 | 136 | 113647 | 7 | 7 | 5 | 5 | 0.21 | 1 | 0.971 | 0.993 | 0.75 | 0.772 | 0.757 | 0.988 | 0.760 | Glycogen phosphorylase, brain form OS=Mus musculus GN=Pygb PE=1 SV=3 |
| 190 | 1 | UniProt n | P35979 | 473 | 22799 | 18 | 18 | 6 | 6 | 3.35 | 1 | 0.999 | 1.009 | 0.713 | 0.812 | 0.757 | 1.003 | 0.760 | 60S ribosomal protein L12 OS=Mus musculus GN=Rpl12 PE=1 SV=2 |
| 307 | 1 | UniProt n | A0A0J9YU | 286 | 19801 | 10 | 10 | 5 | 5 | 1.87 | 1 | 0.986 | 0.973 | 0.777 | 0.781 | 0.722 | 0.986 | 0.760 | Ubiquitin-conjugating enzyme E2 K OS=Mus musculus GN=Ube2k PE=1 SV=1 |
| 173 | 1 | UniProt n | Q3U9N4 | 543 | 78145 | 22 | 22 | 9 | 9 | 0.82 | 1 | 0.95 | 0.84 | 0.771 | 0.72 | 0.789 | 0.928 | 0.759 | Granulins OS=Mus musculus GN=Grn PE=1 SV=1 |
| 506 | 1 | UniProt n | P97765 | 140 | 32973 | 5 | 5 | 3 | 3 | 0.47 | 1 | 0.98 | 1.025 | 0.82 | 0.769 | 0.694 | 1.001 | 0.759 | WW domain-binding protein 2 OS=Mus musculus GN=Wbp2 PE=1 SV=1 |
| 1023 | 1 | UniProt n | B2KF35 | 43 | 34974 | 1 | 1 | 1 | 1 | 0.13 | 1 | 0.839 | 0.8 | 0.817 | 0.708 | 0.756 | 0.876 | 0.759 | Mitogen-activated protein kinase OS=Mus musculus GN=Mapk14 PE=1 SV=1 |
| 401 | 1 | UniProt n | Q9DCC4 | 208 | 32113 | 9 | 9 | 4 | 4 | 0.69 | 1 | 0.911 | 0.869 | 0.735 | 0.762 | 0.779 | 0.925 | 0.758 | Pyrroline-5-carboxylate reductase 3 OS=Mus musculus GN=Pycr3 PE=1 SV=2 |
| 543 | 1 | UniProt n | Q9CPX4 | 125 | 23848 | 5 | 5 | 4 | 4 | 1.4 | 1 | 0.86 | 0.841 | 0.757 | 0.809 | 0.712 | 0.898 | 0.758 | Ferritin OS=Mus musculus GN=Ftl1 PE=1 SV=1 |
| 284 | 1 | UniProt n | P97855 | 312 | 57927 | 8 | 8 | 6 | 6 | 0.55 | 1 | 0.921 | 0.971 | 0.702 | 0.812 | 0.764 | 0.963 | 0.758 | Ras GTPase-activating protein-binding protein 1 OS=Mus musculus GN=G3bp1 PE=1 SV=1 |
| 232 | 1 | UniProt n | Q8R2P8 | 376 | 85607 | 14 | 14 | 7 | 7 | 0.41 | 1 | 0.991 | 1.111 | 0.807 | 0.737 | 0.732 | 1.033 | 0.758 | Lysine--tRNA ligase OS=Mus musculus GN=Kars PE=1 SV=1 |
| 606 | 1 | UniProt n | P23116 | 102 | 188930 | 5 | 5 | 5 | 5 | 0.12 | 1 | 0.913 | 0.923 | 0.804 | 0.741 | 0.73 | 0.945 | 0.758 | Eukaryotic translation initiation factor 3 subunit A OS=Mus musculus GN=Eif3a PE=1 SV=5 |
| 565 | 1 | UniProt n | Q6PGH2 | 117 | 25225 | 5 | 5 | 4 | 4 | 0.94 | 1 | 0.992 | 1.014 | 0.736 | 0.774 | 0.762 | 1.002 | 0.757 | Jupiter microtubule associated homolog 2 OS=Mus musculus GN=Jpt2 PE=1 SV=1 |
| 1013 | 1 | UniProt n | Q9CR51 | 44 | 17458 | 1 | 1 | 1 | 1 | 0.27 | 1 | 0.768 | 0.788 | 0.733 | 0.872 | 0.679 | 0.846 | 0.757 | V-type proton ATPase subunit G 1 OS=Mus musculus GN=Atp6v1g1 PE=1 SV=3 |
| 416 | 1 | UniProt n | Q5XJF6 | 198 | 34688 | 13 | 13 | 4 | 4 | 0.83 | 1 | 1.066 | 1.029 | 0.697 | 0.758 | 0.821 | 1.031 | 0.757 | Ribosomal protein OS=Mus musculus GN=Rpl10a PE=1 SV=1 |
| 510 | 1 | UniProt n | Q8R317 | 137 | 65588 | 3 | 3 | 2 | 2 | 0.14 | 1 | 0.886 | 0.929 | 0.773 | 0.75 | 0.748 | 0.937 | 0.757 | Ubiquilin-1 OS=Mus musculus GN=Ubqln1 PE=1 SV=1 |
| 372 | 1 | UniProt n | Q9CXY6 | 229 | 48695 | 6 | 6 | 6 | 6 | 0.68 | 1 | 0.968 | 0.955 | 0.769 | 0.768 | 0.734 | 0.974 | 0.757 | Interleukin enhancer-binding factor 2 OS=Mus musculus GN=Ilf2 PE=1 SV=1 |
| 746 | 1 | UniProt n | P05132 | 74 | 51588 | 2 | 2 | 2 | 2 | 0.18 | 1 | 0.841 | 0.857 | 0.793 | 0.748 | 0.729 | 0.897 | 0.756 | cAMP-dependent protein kinase catalytic subunit alpha OS=Mus musculus GN=Prkaca PE=1 SV=3 |
| 508 | 1 | UniProt n | Q61035 | 139 | 70587 | 8 | 8 | 8 | 8 | 0.62 | 1 | 1.016 | 1.107 | 0.767 | 0.762 | 0.738 | 1.040 | 0.756 | Histidine--tRNA ligase, cytoplasmic OS=Mus musculus GN=Hars PE=1 SV=2 |
| 834 | 1 | UniProt n | A0A0G2JG | 61 | 79734 | 5 | 5 | 4 | 4 | 0.24 | 1 | 0.87 | 0.981 | 0.794 | 0.77 | 0.704 | 0.949 | 0.755 | Pre-mRNA-splicing factor ATP-dependent RNA helicase DHX15 OS=Mus musculus GN=Dhx15 PE=1 SV=1 |
| 749 | 1 | UniProt n | P54728 | 73 | 49616 | 7 | 7 | 4 | 4 | 0.41 | 1 | 0.971 | 1.032 | 0.796 | 0.752 | 0.719 | 1.001 | 0.755 | UV excision repair protein RAD23 homolog B OS=Mus musculus GN=Rad23b PE=1 SV=2 |
| 678 | 1 | UniProt n | A2A8L5 | 86 | 236870 | 3 | 3 | 3 | 3 | 0.06 | 1 | 0.978 | 1.094 | 0.735 | 0.795 | 0.736 | 1.023 | 0.755 | Receptor-type tyrosine-protein phosphatase F OS=Mus musculus GN=Ptprf PE=1 SV=1 |
| 485 | 2 | UniProt n | Q9EST5 | 56 | 35761 | 5 | 5 | 3 | 3 | 0.42 | 1 | 1.045 | 0.954 | 0.778 | 0.79 | 0.699 | 0.999 | 0.755 | Acidic leucine-rich nuclear phosphoprotein 32 family member B OS=Mus musculus GN=Anp32b PE=1 SV=1 |
| 83 | 2 | UniProt n | P49312 | 747 | 40047 | 38 | 38 | 7 | 7 | 1.86 | 1 | 1.036 | 1.04 | 0.76 | 0.767 | 0.737 | 1.025 | 0.755 | Heterogeneous nuclear ribonucleoprotein A1 OS=Mus musculus GN=Hnrnpa1 PE=1 SV=2 |
| 18 | 1 | UniProt n | P37889 | 4699 | 143968 | 163 | 163 | 31 | 31 | 1.98 | 1 | 0.981 | 0.963 | 0.778 | 0.718 | 0.769 | 0.981 | 0.755 | Fibulin-2 OS=Mus musculus GN=Fbln2 PE=1 SV=2 |
| 594 | 1 | UniProt n | P10852 | 105 | 67214 | 4 | 4 | 3 | 3 | 0.21 | 1 | 0.76 | 0.857 | 0.71 | 0.758 | 0.798 | 0.867 | 0.754 | 4F2 cell-surface antigen heavy chain OS=Mus musculus GN=Slc3a2 PE=1 SV=1 |
| 1099 | 1 | UniProt n | Q8R1F1 | 36 | 98352 | 1 | 1 | 1 | 1 | 0.04 | 1 | 1.068 | 1.136 | 0.771 | 0.69 | 0.807 | 1.067 | 0.754 | Niban-like protein 1 OS=Mus musculus GN=Fam129b PE=1 SV=2 |
| 99 | 1 | UniProt n | A0A0A0M | 1011 | 47681 | 39 | 39 | 9 | 9 | 2.16 | 1 | 1.072 | 1.1 | 0.767 | 0.742 | 0.753 | 1.056 | 0.754 | Glyceraldehyde-3-phosphate dehydrogenase OS=Mus musculus GN=Gapdh PE=1 SV=1 |
| 544 | 1 | UniProt n | Q60865 | 124 | 87081 | 4 | 4 | 2 | 2 | 0.1 | 1 | 0.943 | 0.944 | 0.809 | 0.711 | 0.745 | 0.962 | 0.754 | Caprin-1 OS=Mus musculus GN=Caprin1 PE=1 SV=2 |
| 60 | 1 | UniProt n | P40142 | 1530 | 81829 | 58 | 58 | 17 | 17 | 2.12 | 1 | 1.012 | 1.043 | 0.776 | 0.738 | 0.747 | 1.018 | 0.753 | Transketolase OS=Mus musculus GN=Tkt PE=1 SV=1 |
| 729 | 1 | UniProt n | A0A0G2JF | 77 | 21388 | 3 | 3 | 2 | 2 | 0.48 | 1 | 1.026 | 1.028 | 0.752 | 0.838 | 0.676 | 1.018 | 0.752 | Actin-related protein 2/3 complex subunit 3 (Fragment) OS=Mus musculus GN=Arpc3 PE=1 SV=1 |
| 553 | 1 | UniProt n | A0A0R4J0 | 121 | 32667 | 5 | 5 | 4 | 4 | 0.67 | 1 | 0.973 | 0.917 | 0.755 | 0.734 | 0.767 | 0.963 | 0.752 | UMP-CMP kinase OS=Mus musculus GN=Cmpk1 PE=1 SV=1 |
| 430 | 1 | UniProt n | A0A0R4IZ' | 184 | 91702 | 10 | 10 | 8 | 8 | 0.52 | 1 | 0.977 | 1.024 | 0.742 | 0.754 | 0.759 | 1.000 | 0.752 | Thimet oligopeptidase OS=Mus musculus GN=Thop1 PE=1 SV=1 |
| 1093 | 1 | UniProt n | H3BIV6 | 36 | 50224 | 1 | 1 | 1 | 1 | 0.09 | 1 | 0.834 | 0.987 | 0.697 | 0.686 | 0.888 | 0.937 | 0.752 | Cysteine and histidine-rich protein 1 OS=Mus musculus GN=Cyhr1 PE=1 SV=1 |
| 213 | 1 | UniProt n | Q9D0F9 | 409 | 72865 | 18 | 18 | 11 | 11 | 0.9 | 1 | 0.96 | 0.977 | 0.745 | 0.74 | 0.77 | 0.979 | 0.752 | Phosphoglucomutase-1 OS=Mus musculus GN=Pgm1 PE=1 SV=4 |
| 246 | 1 | UniProt n | Q9R1P4 | 354 | 33712 | 24 | 24 | 9 | 9 | 2.07 | 1 | 0.994 | 0.987 | 0.786 | 0.72 | 0.75 | 0.994 | 0.752 | Proteasome subunit alpha type-1 OS=Mus musculus GN=Psma1 PE=1 SV=1 |
| 643 | 1 | UniProt n | Q3UID0 | 92 | 154125 | 4 | 4 | 3 | 3 | 0.09 | 1 | 0.981 | 1 | 0.78 | 0.686 | 0.793 | 0.994 | 0.751 | SWI/SNF complex subunit SMARCC2 OS=Mus musculus GN=Smarcc2 PE=1 SV=1 |
| 1020 | 1 | UniProt n | Q62383 | 43 | 236128 | 1 | 1 | 1 | 1 | 0.02 | 1 | 0.926 | 0.826 | 0.835 | 0.678 | 0.749 | 0.915 | 0.751 | Transcription elongation factor SPT6 OS=Mus musculus GN=Supt6h PE=1 SV=2 |
| 12 | 4 | UniProt n | Q9CQN1 | 336 | 95341 | 7 | 7 | 2 | 2 | 0.09 | 1 | 1.113 | 1.16 | 0.784 | 0.703 | 0.769 | 1.089 | 0.751 | Heat shock protein 75 kDa, mitochondrial OS=Mus musculus GN=Trap1 PE=1 SV=1 |
| 208 | 2 | UniProt n | F8WIT2 | 383 | 92340 | 16 | 16 | 9 | 9 | 0.66 | 1 | 1.029 | 1.117 | 0.738 | 0.759 | 0.755 | 1.048 | 0.751 | Annexin OS=Mus musculus GN=Anxa6 PE=1 SV=1 |
| 58 | 1 | UniProt n | P26041 | 1565 | 87590 | 77 | 77 | 26 | 26 | 4.44 | 1 | 0.991 | 1.016 | 0.771 | 0.753 | 0.728 | 1.002 | 0.750 | Moesin OS=Mus musculus GN=Msn PE=1 SV=3 |
| 1002 | 1 | UniProt n | P70372 | 44 | 42064 | 1 | 1 | 1 | 1 | 0.11 | 1 | 0.972 | 0.923 | 0.771 | 0.741 | 0.739 | 0.964 | 0.750 | ELAV-like protein 1 OS=Mus musculus GN=Elavl1 PE=1 SV=2 |
| 39 | 1 | UniProt n | P58252 | 2385 | 115808 | 104 | 104 | 35 | 35 | 3.33 | 1 | 1.061 | 1.071 | 0.759 | 0.714 | 0.779 | 1.044 | 0.750 | Elongation factor 2 OS=Mus musculus GN=Eef2 PE=1 SV=2 |
| 464 | 1 | UniProt n | Q9WTX5 | 163 | 24274 | 8 | 8 | 4 | 4 | 0.99 | 1 | 1.261 | 1.084 | 0.788 | 0.774 | 0.691 | 1.110 | 0.750 | S-phase kinase-associated protein 1 OS=Mus musculus GN=Skp1 PE=1 SV=3 |
| 987 | 1 | UniProt n | A2AQE4 | 46 | 58416 | 3 | 3 | 2 | 2 | 0.16 | 1 | 0.92 | 0.832 | 0.698 | 0.745 | 0.809 | 0.915 | 0.749 | COP9 signalosome complex subunit 2 OS=Mus musculus GN=Cops2 PE=1 SV=1 |
| 208 | 1 | UniProt n | P14824 | 428 | 92937 | 16 | 16 | 9 | 9 | 0.65 | 1 | 1.026 | 1.12 | 0.737 | 0.756 | 0.755 | 1.047 | 0.749 | Annexin A6 OS=Mus musculus GN=Anxa6 PE=1 SV=3 |
| 150 | 1 | UniProt n | Q920E5 | 668 | 49045 | 27 | 27 | 13 | 13 | 2.06 | 1 | 1.068 | 1.08 | 0.742 | 0.766 | 0.74 | 1.049 | 0.749 | Farnesyl pyrophosphate synthase OS=Mus musculus GN=Fdps PE=1 SV=1 |
| 533 | 1 | UniProt n | Q9CY64 | 127 | 40334 | 6 | 6 | 4 | 4 | 0.52 | 1 | 0.964 | 0.957 | 0.754 | 0.728 | 0.766 | 0.973 | 0.749 | Biliverdin reductase A OS=Mus musculus GN=Blvra PE=1 SV=1 |
| 939 | 1 | UniProt n | Q9CQR2 | 50 | 11053 | 4 | 4 | 2 | 2 | 1.1 | 1 | 0.736 | 0.72 | 0.614 | 0.889 | 0.77 | 0.809 | 0.749 | 40S ribosomal protein S21 OS=Mus musculus GN=Rps21 PE=1 SV=1 |
| 1039 | 1 | UniProt n | P59999 | 41 | 24005 | 2 | 2 | 2 | 2 | 0.42 | 1 | 0.919 | 0.885 | 0.687 | 0.802 | 0.762 | 0.933 | 0.749 | Actin-related protein 2/3 complex subunit 4 OS=Mus musculus GN=Arpc4 PE=1 SV=3 |
| 23 | 1 | UniProt n | P52480 | 3894 | 69828 | 142 | 142 | 30 | 30 | 11.01 | 1 | 1.072 | 1.075 | 0.778 | 0.723 | 0.746 | 1.048 | 0.749 | Pyruvate kinase PKM OS=Mus musculus GN=Pkm PE=1 SV=4 |
| 521 | 2 | UniProt n | P62137 | 127 | 44806 | 4 | 4 | 3 | 3 | 0.33 | 1 | 1.017 | 0.911 | 0.76 | 0.77 | 0.717 | 0.975 | 0.749 | Serine/threonine-protein phosphatase PP1-alpha catalytic subunit OS=Mus musculus GN=Ppp1ca PE=1 SV=1 |
| 411 | 1 | UniProt n | Q9CVB6 | 200 | 42337 | 13 | 13 | 8 | 8 | 1.45 | 1 | 0.997 | 1.007 | 0.745 | 0.745 | 0.754 | 1.001 | 0.748 | Actin-related protein 2/3 complex subunit 2 OS=Mus musculus GN=Arpc2 PE=1 SV=3 |
| 34 | 1 | UniProt n | Q64727 | 2793 | 140224 | 125 | 125 | 45 | 45 | 4.13 | 1 | 1.003 | 1.063 | 0.761 | 0.762 | 0.721 | 1.022 | 0.748 | Vinculin OS=Mus musculus GN=Vcl PE=1 SV=4 |
| 49 | 1 | UniProt n | P62962 | 2041 | 18128 | 70 | 70 | 8 | 8 | 30.53 | 1 | 1.004 | 0.978 | 0.772 | 0.742 | 0.729 | 0.994 | 0.747 | Profilin-1 OS=Mus musculus GN=Pfn1 PE=1 SV=2 |
| 220 | 1 | UniProt n | O35685 | 393 | 48981 | 20 | 20 | 10 | 10 | 1.37 | 1 | 0.99 | 1.037 | 0.798 | 0.756 | 0.692 | 1.009 | 0.747 | Nuclear migration protein nudC OS=Mus musculus GN=Nudc PE=1 SV=1 |
| 349 | 1 | UniProt n | P46664 | 243 | 59133 | 9 | 9 | 6 | 6 | 0.65 | 1 | 0.97 | 1.033 | 0.778 | 0.726 | 0.739 | 1.001 | 0.747 | Adenylosuccinate synthetase isozyme 2 OS=Mus musculus GN=Adss PE=1 SV=2 |
| 254 | 1 | UniProt n | Q9QUM9 | 348 | 33807 | 18 | 18 | 10 | 10 | 2.92 | 1 | 1.006 | 0.986 | 0.717 | 0.781 | 0.744 | 0.997 | 0.747 | Proteasome subunit alpha type-6 OS=Mus musculus GN=Psma6 PE=1 SV=1 |
| 775 | 1 | UniProt n | Q9DCH4 | 69 | 41399 | 3 | 3 | 2 | 2 | 0.23 | 1 | 1.114 | 0.998 | 0.672 | 0.951 | 0.65 | 1.036 | 0.746 | Eukaryotic translation initiation factor 3 subunit F OS=Mus musculus GN=Eif3f PE=1 SV=2 |
| 666 | 1 | UniProt n | A0A0J9YU | 89 | 174203 | 5 | 5 | 3 | 3 | 0.08 | 1 | 1.002 | 1.096 | 0.768 | 0.746 | 0.724 | 1.032 | 0.746 | Eukaryotic translation initiation factor 4 gamma 1 OS=Mus musculus GN=Eif4g1 PE=1 SV=1 |
| 841 | 1 | UniProt n | Q6P4T2 | 61 | 287401 | 2 | 2 | 2 | 2 | 0.03 | 1 | 0.918 | 1.004 | 0.743 | 0.789 | 0.707 | 0.973 | 0.746 | U5 small nuclear ribonucleoprotein 200 kDa helicase OS=Mus musculus GN=Snrnp200 PE=1 SV=1 |
| 1004 | 1 | UniProt n | Q9D6S7 | 44 | 37338 | 1 | 1 | 1 | 1 | 0.12 | 1 | 1.149 | 0.783 | 0.741 | 0.599 | 0.931 | 0.965 | 0.745 | Ribosome-recycling factor, mitochondrial OS=Mus musculus GN=Mrrf PE=1 SV=1 |
| 350 | 1 | UniProt n | Q9JKB1 | 243 | 31140 | 9 | 9 | 4 | 4 | 0.71 | 1 | 0.963 | 1.002 | 0.812 | 0.733 | 0.694 | 0.988 | 0.745 | Ubiquitin carboxyl-terminal hydrolase isozyme L3 OS=Mus musculus GN=Uchl3 PE=1 SV=2 |
| 514 | 1 | UniProt n | Q61881 | 136 | 92008 | 5 | 5 | 4 | 4 | 0.2 | 1 | 0.793 | 0.907 | 0.776 | 0.778 | 0.683 | 0.896 | 0.744 | DNA replication licensing factor MCM7 OS=Mus musculus GN=Mcm7 PE=1 SV=1 |
| 481 | 1 | UniProt n | O55234 | 155 | 31694 | 8 | 8 | 6 | 6 | 1.21 | 1 | 1.039 | 1.042 | 0.751 | 0.753 | 0.728 | 1.027 | 0.744 | Proteasome subunit beta type-5 OS=Mus musculus GN=Psmb5 PE=1 SV=3 |
| 289 | 2 | UniProt n | Q60973 | 237 | 55641 | 13 | 13 | 7 | 7 | 0.7 | 1 | 1.039 | 1.033 | 0.743 | 0.713 | 0.777 | 1.024 | 0.744 | Histone-binding protein RBBP7 OS=Mus musculus GN=Rbbp7 PE=1 SV=1 |
| 656 | 1 | UniProt n | P47791 | 90 | 64782 | 3 | 3 | 3 | 3 | 0.22 | 1 | 1.019 | 1.008 | 0.724 | 0.742 | 0.766 | 1.009 | 0.744 | Glutathione reductase, mitochondrial OS=Mus musculus GN=Gsr PE=1 SV=3 |
| 148 | 1 | UniProt n | F8WJK8 | 675 | 49169 | 29 | 29 | 8 | 8 | 1.36 | 1 | 1.046 | 1.029 | 0.755 | 0.717 | 0.76 | 1.025 | 0.744 | Hsc70-interacting protein OS=Mus musculus GN=St13 PE=1 SV=1 |
| 437 | 1 | UniProt n | Q8VCR7 | 177 | 24658 | 7 | 7 | 5 | 5 | 1.77 | 1 | 1.021 | 0.967 | 0.704 | 0.773 | 0.755 | 0.996 | 0.743 | Protein ABHD14B OS=Mus musculus GN=Abhd14b PE=1 SV=1 |
| 23 | 2 | UniProt n | P52480-2 | 3606 | 69617 | 139 | 139 | 30 | 30 | 11.1 | 1 | 1.068 | 1.07 | 0.77 | 0.716 | 0.745 | 1.045 | 0.743 | Isoform M1 of Pyruvate kinase PKM OS=Mus musculus GN=Pkm |
| 619 | 1 | UniProt n | P20108 | 100 | 32552 | 5 | 5 | 3 | 3 | 0.47 | 1 | 0.962 | 1.01 | 0.762 | 0.773 | 0.697 | 0.990 | 0.743 | Thioredoxin-dependent peroxide reductase, mitochondrial OS=Mus musculus GN=Prdx3 PE=1 SV=1 |
| 21 | 2 | UniProt n | Q80X90 | 2439 | 332864 | 99 | 99 | 57 | 57 | 1.18 | 1 | 0.956 | 0.985 | 0.751 | 0.73 | 0.748 | 0.980 | 0.743 | Filamin-B OS=Mus musculus GN=Flnb PE=1 SV=3 |
| 175 | 3 | UniProt n | P47754 | 429 | 39777 | 15 | 15 | 6 | 6 | 1.1 | 1 | 0.98 | 0.946 | 0.717 | 0.72 | 0.794 | 0.975 | 0.743 | F-actin-capping protein subunit alpha-2 OS=Mus musculus GN=Capza2 PE=1 SV=3 |
| 313 | 1 | UniProt n | Q3UEB3 | 278 | 70388 | 8 | 8 | 5 | 5 | 0.35 | 1 | 1.009 | 1.069 | 0.77 | 0.715 | 0.744 | 1.026 | 0.743 | Poly(U)-binding-splicing factor PUF60 OS=Mus musculus GN=Puf60 PE=1 SV=2 |
| 100 | 1 | UniProt n | A0A1B0G | 1009 | 50048 | 46 | 46 | 17 | 17 | 4.88 | 1 | 1.063 | 1.049 | 0.74 | 0.73 | 0.757 | 1.037 | 0.742 | L-lactate dehydrogenase OS=Mus musculus GN=Ldha PE=1 SV=1 |
| 193 | 1 | UniProt n | Q923D2 | 469 | 25317 | 16 | 16 | 6 | 6 | 1.7 | 1 | 0.789 | 0.821 | 0.744 | 0.759 | 0.724 | 0.865 | 0.742 | Flavin reductase (NADPH) OS=Mus musculus GN=Blvrb PE=1 SV=3 |
| 540 | 1 | UniProt n | P14685 | 125 | 71420 | 5 | 5 | 3 | 3 | 0.19 | 1 | 0.793 | 0.941 | 0.637 | 0.89 | 0.721 | 0.907 | 0.742 | 26S proteasome non-ATPase regulatory subunit 3 OS=Mus musculus GN=Psmd3 PE=1 SV=3 |
| 854 | 1 | UniProt n | P97946 | 59 | 48907 | 1 | 1 | 1 | 1 | 0.09 | 1 | 1.016 | 1.036 | 0.711 | 0.802 | 0.716 | 1.017 | 0.742 | Vascular endothelial growth factor D OS=Mus musculus GN=Vegfd PE=2 SV=1 |
| 547 | 1 | UniProt n | Q9CYA0 | 123 | 45721 | 4 | 4 | 2 | 2 | 0.2 | 1 | 0.929 | 0.901 | 0.722 | 0.781 | 0.724 | 0.942 | 0.742 | Cysteine-rich with EGF-like domain protein 2 OS=Mus musculus GN=Creld2 PE=1 SV=1 |
| 688 | 1 | UniProt n | H3BJ97 | 85 | 52294 | 2 | 2 | 1 | 1 | 0.08 | 1 | 0.851 | 0.826 | 0.766 | 0.723 | 0.736 | 0.889 | 0.741 | Tubulointerstitial nephritis antigen-like OS=Mus musculus GN=Tinagl1 PE=1 SV=1 |
| 419 | 1 | UniProt n | Q9DAR7 | 194 | 44532 | 10 | 10 | 7 | 7 | 0.94 | 1 | 1.076 | 1.022 | 0.711 | 0.761 | 0.753 | 1.032 | 0.741 | m7GpppX diphosphatase OS=Mus musculus GN=Dcps PE=1 SV=1 |
| 465 | 1 | UniProt n | Q6PDM2 | 162 | 30558 | 9 | 9 | 5 | 5 | 0.99 | 1 | 1.027 | 1.058 | 0.73 | 0.743 | 0.751 | 1.028 | 0.741 | Serine/arginine-rich splicing factor 1 OS=Mus musculus GN=Srsf1 PE=1 SV=3 |
| 701 | 1 | UniProt n | Q8K4Z3 | 82 | 35488 | 2 | 2 | 1 | 1 | 0.13 | 1 | 0.936 | 0.958 | 0.721 | 0.81 | 0.697 | 0.964 | 0.741 | NAD(P)H-hydrate epimerase OS=Mus musculus GN=Naxe PE=1 SV=1 |
| 967 | 1 | UniProt n | Q99LJ6 | 48 | 24836 | 3 | 3 | 2 | 2 | 0.4 | 1 | 1.038 | 1.096 | 0.81 | 0.676 | 0.743 | 1.044 | 0.741 | Glutathione peroxidase 7 OS=Mus musculus GN=Gpx7 PE=1 SV=1 |
| 48 | 1 | UniProt n | A3KGU5 | 2043 | 343597 | 72 | 72 | 48 | 48 | 0.93 | 1 | 0.998 | 1.026 | 0.761 | 0.734 | 0.728 | 1.008 | 0.741 | Spectrin alpha chain, non-erythrocytic 1 OS=Mus musculus GN=Sptan1 PE=1 SV=1 |
| 267 | 1 | UniProt n | Q9JII6 | 328 | 43136 | 19 | 19 | 9 | 9 | 1.66 | 1 | 1.037 | 1.016 | 0.731 | 0.727 | 0.765 | 1.018 | 0.741 | Alcohol dehydrogenase [NADP(+)] OS=Mus musculus GN=Akr1a1 PE=1 SV=3 |
| 92 | 1 | UniProt n | Q99PT1 | 1092 | 29523 | 53 | 53 | 10 | 10 | 5.35 | 1 | 1.034 | 1.058 | 0.757 | 0.715 | 0.751 | 1.030 | 0.741 | Rho GDP-dissociation inhibitor 1 OS=Mus musculus GN=Arhgdia PE=1 SV=3 |
| 347 | 1 | UniProt n | Q8JZQ9 | 246 | 108671 | 10 | 10 | 6 | 6 | 0.26 | 1 | 0.975 | 1.031 | 0.77 | 0.701 | 0.752 | 1.002 | 0.740 | Eukaryotic translation initiation factor 3 subunit B OS=Mus musculus GN=Eif3b PE=1 SV=1 |
| 1156 | 1 | UniProt n | E9PYH2 | 31 | 51076 | 1 | 1 | 1 | 1 | 0.09 | 1 | 0.989 | 0.995 | 0.769 | 0.7 | 0.754 | 0.995 | 0.740 | Cytosolic acyl coenzyme A thioester hydrolase OS=Mus musculus GN=Acot7 PE=1 SV=1 |
| 26 | 1 | UniProt n | P05064 | 3805 | 47913 | 105 | 105 | 16 | 16 | 5.35 | 1 | 1.034 | 1.028 | 0.765 | 0.744 | 0.713 | 1.021 | 0.740 | Fructose-bisphosphate aldolase A OS=Mus musculus GN=Aldoa PE=1 SV=2 |
| 53 | 1 | UniProt n | P08122 | 1726 | 192826 | 61 | 61 | 17 | 17 | 0.52 | 1 | 0.925 | 0.92 | 0.74 | 0.743 | 0.737 | 0.948 | 0.740 | Collagen alpha-2(IV) chain OS=Mus musculus GN=Col4a2 PE=1 SV=4 |
| 306 | 1 | UniProt n | P80313 | 288 | 73109 | 12 | 12 | 9 | 9 | 0.68 | 1 | 0.947 | 0.966 | 0.741 | 0.719 | 0.76 | 0.971 | 0.740 | T-complex protein 1 subunit eta OS=Mus musculus GN=Cct7 PE=1 SV=1 |
| 1136 | 1 | UniProt n | P52293 | 33 | 67903 | 1 | 1 | 1 | 1 | 0.06 | 1 | 0.858 | 0.83 | 0.77 | 0.654 | 0.801 | 0.893 | 0.739 | Importin subunit alpha-1 OS=Mus musculus GN=Kpna2 PE=1 SV=2 |
| 639 | 1 | UniProt n | P70362 | 93 | 41686 | 3 | 3 | 2 | 2 | 0.35 | 1 | 0.943 | 0.934 | 0.753 | 0.724 | 0.737 | 0.959 | 0.738 | Ubiquitin recognition factor in ER-associated degradation protein 1 OS=Mus musculus GN=Ufd1 PE=1 SV=2 |
| 230 | 1 | UniProt n | O70400 | 380 | 43117 | 12 | 12 | 7 | 7 | 0.98 | 1 | 0.984 | 0.958 | 0.787 | 0.708 | 0.721 | 0.981 | 0.738 | PDZ and LIM domain protein 1 OS=Mus musculus GN=Pdlim1 PE=1 SV=4 |
| 705 | 1 | UniProt n | Q8VIJ6 | 81 | 85829 | 6 | 6 | 6 | 6 | 0.34 | 1 | 0.973 | 0.963 | 0.748 | 0.712 | 0.754 | 0.979 | 0.738 | Splicing factor, proline- and glutamine-rich OS=Mus musculus GN=Sfpq PE=1 SV=1 |
| 51 | 1 | UniProt n | Q62261 | 1854 | 329803 | 75 | 75 | 48 | 48 | 0.93 | 1 | 0.997 | 1.052 | 0.768 | 0.734 | 0.712 | 1.016 | 0.738 | Spectrin beta chain, non-erythrocytic 1 OS=Mus musculus GN=Sptbn1 PE=1 SV=2 |
| 293 | 1 | UniProt n | A0A0J9YU | 301 | 30242 | 13 | 13 | 6 | 6 | 1.3 | 1 | 1.128 | 1.148 | 0.755 | 0.72 | 0.738 | 1.090 | 0.738 | High mobility group protein B1 OS=Mus musculus GN=Hmgb1 PE=1 SV=1 |
| 17 | 4 | UniProt n | P17879 | 889 | 84661 | 26 | 26 | 5 | 5 | 0.35 | 1 | 0.945 | 0.97 | 0.783 | 0.707 | 0.724 | 0.971 | 0.737 | Heat shock 70 kDa protein 1B OS=Mus musculus GN=Hspa1b PE=1 SV=3 |
| 1273 | 1 | UniProt n | P61971 | 19 | 16736 | 1 | 1 | 1 | 1 | 0.28 | 1 | 0.844 | 0.847 | 0.724 | 0.84 | 0.659 | 0.894 | 0.737 | Nuclear transport factor 2 OS=Mus musculus GN=Nutf2 PE=1 SV=1 |
| 895 | 1 | UniProt n | Q3TE40 | 55 | 33514 | 4 | 4 | 2 | 2 | 0.29 | 1 | 0.938 | 1.004 | 0.835 | 0.66 | 0.727 | 0.980 | 0.737 | Replication protein A 32 kDa subunit OS=Mus musculus GN=Rpa2 PE=1 SV=1 |
| 86 | 8 | UniProt n | G5E8R1 | 482 | 36191 | 37 | 37 | 11 | 11 | 3.04 | 1 | 1.017 | 0.971 | 0.769 | 0.766 | 0.68 | 0.996 | 0.737 | Tropomyosin 1, alpha, isoform CRA_j OS=Mus musculus GN=Tpm1 PE=1 SV=1 |
| 958 | 1 | UniProt n | P47962 | 48 | 44906 | 1 | 1 | 1 | 1 | 0.1 | 1 | 1.266 | 1.416 | 0.806 | 0.719 | 0.69 | 1.215 | 0.737 | 60S ribosomal protein L5 OS=Mus musculus GN=Rpl5 PE=1 SV=3 |
| 106 | 1 | UniProt n | P35700 | 978 | 28430 | 61 | 61 | 11 | 11 | 4.88 | 1 | 1.05 | 1.041 | 0.739 | 0.728 | 0.743 | 1.030 | 0.737 | Peroxiredoxin-1 OS=Mus musculus GN=Prdx1 PE=1 SV=1 |
| 21 | 3 | UniProt n | Q8VHX6 | 1724 | 341117 | 67 | 67 | 41 | 41 | 0.69 | 1 | 0.98 | 1.028 | 0.749 | 0.726 | 0.734 | 1.002 | 0.736 | Filamin-C OS=Mus musculus GN=Flnc PE=1 SV=3 |
| 77 | 1 | UniProt n | Q61598 | 1276 | 62478 | 56 | 56 | 16 | 16 | 3.14 | 1 | 1.017 | 1.042 | 0.742 | 0.718 | 0.748 | 1.020 | 0.736 | Rab GDP dissociation inhibitor beta OS=Mus musculus GN=Gdi2 PE=1 SV=1 |
| 1073 | 1 | UniProt n | Q00780 | 39 | 88253 | 2 | 2 | 2 | 2 | 0.1 | 1 | 0.889 | 0.905 | 0.706 | 0.738 | 0.763 | 0.930 | 0.735 | Collagen alpha-1(VIII) chain OS=Mus musculus GN=Col8a1 PE=1 SV=3 |
| 862 | 1 | UniProt n | Q9R001 | 58 | 120746 | 1 | 1 | 1 | 1 | 0.04 | 1 | 0.842 | 0.983 | 0.733 | 0.639 | 0.848 | 0.939 | 0.735 | A disintegrin and metalloproteinase with thrombospondin motifs 5 OS=Mus musculus GN=Adamts5 PE=2 SV=2 |
| 450 | 1 | UniProt n | Q3UKN6 | 171 | 62278 | 7 | 7 | 5 | 5 | 0.5 | 1 | 0.836 | 0.81 | 0.73 | 0.794 | 0.685 | 0.878 | 0.735 | Nucleobindin-2 OS=Mus musculus GN=Nucb2 PE=1 SV=1 |
| 381 | 1 | UniProt n | Q9JK81 | 222 | 46972 | 10 | 10 | 5 | 5 | 0.71 | 1 | 0.992 | 1.015 | 0.771 | 0.716 | 0.717 | 1.002 | 0.734 | UPF0160 protein MYG1, mitochondrial OS=Mus musculus GN=Myg1 PE=1 SV=1 |
| 858 | 1 | UniProt n | P50543 | 59 | 13951 | 4 | 4 | 2 | 2 | 1.43 | 1 | 1.047 | 0.956 | 0.744 | 0.751 | 0.708 | 1.000 | 0.734 | Protein S100-A11 OS=Mus musculus GN=S100a11 PE=1 SV=1 |
| 794 | 1 | UniProt n | A2A8E2 | 66 | 20308 | 2 | 2 | 1 | 1 | 0.23 | 1 | 0.973 | 0.937 | 0.727 | 0.744 | 0.731 | 0.970 | 0.734 | RIKEN cDNA 0610037L13 gene OS=Mus musculus GN=0610037L13Rik PE=1 SV=1 |
| 979 | 1 | UniProt n | P83870 | 47 | 16949 | 1 | 1 | 1 | 1 | 0.28 | 1 | 0.997 | 0.846 | 0.776 | 0.683 | 0.746 | 0.945 | 0.734 | PHD finger-like domain-containing protein 5A OS=Mus musculus GN=Phf5a PE=1 SV=1 |
| 403 | 1 | UniProt n | A0A1L1SS | 206 | 16423 | 3 | 3 | 1 | 1 | 0.66 | 1 | 0.741 | 0.783 | 0.782 | 0.71 | 0.712 | 0.834 | 0.734 | Layilin (Fragment) OS=Mus musculus GN=Layn PE=1 SV=1 |
| 233 | 1 | UniProt n | Q11011 | 373 | 121456 | 18 | 18 | 10 | 10 | 0.42 | 1 | 0.957 | 0.919 | 0.769 | 0.694 | 0.74 | 0.958 | 0.734 | Puromycin-sensitive aminopeptidase OS=Mus musculus GN=Npepps PE=1 SV=2 |
| 77 | 2 | UniProt n | P50396 | 734 | 60075 | 37 | 37 | 12 | 12 | 1.88 | 1 | 0.97 | 0.987 | 0.749 | 0.728 | 0.723 | 0.986 | 0.733 | Rab GDP dissociation inhibitor alpha OS=Mus musculus GN=Gdi1 PE=1 SV=3 |
| 353 | 1 | UniProt n | P14069 | 239 | 13437 | 22 | 22 | 4 | 4 | 2.42 | 1 | 1.113 | 1.199 | 0.77 | 0.713 | 0.718 | 1.101 | 0.733 | Protein S100-A6 OS=Mus musculus GN=S100a6 PE=1 SV=3 |
| 186 | 1 | UniProt n | Q9Z2U0 | 493 | 35277 | 21 | 21 | 6 | 6 | 1.3 | 1 | 1.109 | 1.121 | 0.755 | 0.731 | 0.714 | 1.075 | 0.733 | Proteasome subunit alpha type-7 OS=Mus musculus GN=Psma7 PE=1 SV=1 |
| 188 | 1 | UniProt n | Q80XU3 | 486 | 38162 | 13 | 13 | 5 | 5 | 1.16 | 1 | 1.036 | 1.029 | 0.762 | 0.685 | 0.754 | 1.022 | 0.733 | Nuclear ubiquitous casein and cyclin-dependent kinase substrate 1 OS=Mus musculus GN=Nucks1 PE=1 SV=1 |
| 144 | 1 | UniProt n | P24369 | 690 | 31912 | 43 | 43 | 10 | 10 | 5.34 | 1 | 1.041 | 1.011 | 0.737 | 0.715 | 0.744 | 1.017 | 0.732 | Peptidyl-prolyl cis-trans isomerase B OS=Mus musculus GN=Ppib PE=1 SV=2 |
| 915 | 1 | UniProt n | A0A0A6Y\ | 53 | 31577 | 2 | 2 | 1 | 1 | 0.14 | 1 | 0.821 | 0.976 | 0.662 | 0.846 | 0.698 | 0.929 | 0.731 | Actin-like protein 6A (Fragment) OS=Mus musculus GN=Actl6a PE=1 SV=1 |
| 67 | 1 | UniProt n | P70296 | 1477 | 24910 | 26 | 26 | 7 | 7 | 3.55 | 1 | 1.012 | 0.988 | 0.761 | 0.713 | 0.72 | 1.000 | 0.731 | Phosphatidylethanolamine-binding protein 1 OS=Mus musculus GN=Pebp1 PE=1 SV=3 |
| 19 | 12 | UniProt n | P14733 | 480 | 78793 | 25 | 25 | 11 | 11 | 0.81 | 1 | 0.999 | 0.986 | 0.756 | 0.741 | 0.697 | 0.995 | 0.731 | Lamin-B1 OS=Mus musculus GN=Lmnb1 PE=1 SV=3 |
| 259 | 1 | UniProt n | A0A1B0GI | 339 | 53186 | 23 | 23 | 9 | 9 | 1.04 | 1 | 1.017 | 1.019 | 0.756 | 0.746 | 0.692 | 1.012 | 0.731 | Transaldolase OS=Mus musculus GN=Taldo1 PE=1 SV=1 |
| 702 | 1 | UniProt n | Q60692 | 82 | 27067 | 3 | 3 | 2 | 2 | 0.36 | 1 | 0.999 | 0.867 | 0.863 | 0.666 | 0.679 | 0.953 | 0.731 | Proteasome subunit beta type-6 OS=Mus musculus GN=Psmb6 PE=1 SV=3 |
| 768 | 1 | UniProt n | A0A1Y7VL | 71 | 25560 | 4 | 4 | 2 | 2 | 0.39 | 1 | 0.968 | 1.051 | 0.751 | 0.732 | 0.708 | 1.006 | 0.730 | NAD(P)H dehydrogenase, quinone 2, isoform CRA_b OS=Mus musculus GN=Nqo2 PE=4 SV=1 |
| 1055 | 1 | UniProt n | Q8BFW7 | 40 | 76824 | 1 | 1 | 1 | 1 | 0.06 | 1 | 0.857 | 0.998 | 0.658 | 0.857 | 0.69 | 0.949 | 0.730 | Lipoma-preferred partner homolog OS=Mus musculus GN=Lpp PE=1 SV=1 |
| 279 | 1 | UniProt n | Q8BSL7 | 318 | 25038 | 12 | 12 | 5 | 5 | 2.8 | 1 | 0.868 | 0.881 | 0.711 | 0.76 | 0.72 | 0.914 | 0.730 | ADP-ribosylation factor 2 OS=Mus musculus GN=Arf2 PE=1 SV=2 |
| 37 | 1 | UniProt n | E9Q616 | 2649 | 848482 | 139 | 139 | 78 | 78 | 0.55 | 1 | 1.05 | 1.06 | 0.777 | 0.72 | 0.694 | 1.036 | 0.730 | AHNAK nucleoprotein (desmoyokin) OS=Mus musculus GN=Ahnak PE=1 SV=1 |
| 406 | 1 | UniProt n | Q9CPU0 | 204 | 26714 | 12 | 12 | 6 | 6 | 2.5 | 1 | 1.038 | 0.915 | 0.637 | 0.901 | 0.674 | 0.983 | 0.729 | Lactoylglutathione lyase OS=Mus musculus GN=Glo1 PE=1 SV=3 |
| 965 | 1 | UniProt n | G3UW70 | 48 | 17096 | 1 | 1 | 1 | 1 | 0.28 | 1 | 0.902 | 0.837 | 0.791 | 0.68 | 0.717 | 0.911 | 0.728 | MCG21719 OS=Mus musculus GN=Lamtor5 PE=1 SV=1 |
| 645 | 1 | UniProt n | Q8R242 | 91 | 49167 | 2 | 2 | 1 | 1 | 0.09 | 1 | 0.862 | 0.644 | 0.722 | 0.685 | 0.779 | 0.822 | 0.728 | Di-N-acetylchitobiase OS=Mus musculus GN=Ctbs PE=1 SV=2 |
| 1042 | 1 | UniProt n | F8WHR6 | 41 | 149897 | 1 | 1 | 1 | 1 | 0.03 | 1 | 0.75 | 0.809 | 0.861 | 0.749 | 0.597 | 0.847 | 0.727 | Probable ATP-dependent RNA helicase DDX46 OS=Mus musculus GN=Ddx46 PE=1 SV=1 |
| 615 | 1 | UniProt n | A2BGG7 | 101 | 27160 | 2 | 2 | 2 | 2 | 0.36 | 1 | 1.232 | 1.258 | 0.583 | 0.863 | 0.764 | 1.157 | 0.727 | Nuclease-sensitive element-binding protein 1 (Fragment) OS=Mus musculus GN=Ybx1 PE=1 SV=1 |
| 509 | 1 | UniProt n | P57759 | 139 | 37673 | 5 | 5 | 3 | 3 | 0.4 | 1 | 0.995 | 1.018 | 0.709 | 0.754 | 0.719 | 1.004 | 0.727 | Endoplasmic reticulum resident protein 29 OS=Mus musculus GN=Erp29 PE=1 SV=2 |
| 140 | 1 | UniProt n | P14869 | 708 | 40417 | 19 | 19 | 9 | 9 | 1.84 | 1 | 1.115 | 1.137 | 0.693 | 0.763 | 0.726 | 1.082 | 0.727 | 60S acidic ribosomal protein P0 OS=Mus musculus GN=Rplp0 PE=1 SV=3 |
| 1155 | 1 | UniProt n | B1AZ46 | 31 | 70911 | 1 | 1 | 1 | 1 | 0.06 | 1 | 1.051 | 1.107 | 0.545 | 0.79 | 0.889 | 1.052 | 0.726 | Brain-specific angiogenesis inhibitor 1-associated protein 2 OS=Mus musculus GN=Baiap2 PE=1 SV=1 |
| 534 | 1 | UniProt n | E9Q242 | 127 | 60995 | 2 | 2 | 2 | 2 | 0.15 | 1 | 0.961 | 1.038 | 0.691 | 0.7 | 0.79 | 0.999 | 0.726 | Adenylosuccinate lyase OS=Mus musculus GN=Adsl PE=1 SV=1 |
| 971 | 1 | UniProt n | P57784 | 47 | 35125 | 1 | 1 | 1 | 1 | 0.13 | 1 | 1.032 | 1.121 | 0.719 | 0.733 | 0.725 | 1.050 | 0.726 | U2 small nuclear ribonucleoprotein A' OS=Mus musculus GN=Snrpa1 PE=1 SV=2 |
| 920 | 1 | UniProt n | P62309 | 53 | 10666 | 1 | 1 | 1 | 1 | 0.47 | 1 | 0.85 | 0.744 | 0.687 | 0.782 | 0.711 | 0.858 | 0.726 | Small nuclear ribonucleoprotein G OS=Mus musculus GN=Snrpg PE=1 SV=1 |
| 326 | 2 | UniProt n | G3UYJ7 | 200 | 33105 | 7 | 7 | 5 | 5 | 0.89 | 1 | 0.965 | 0.873 | 0.742 | 0.739 | 0.695 | 0.944 | 0.725 | Predicted gene 20441 (Fragment) OS=Mus musculus GN=Gm20441 PE=4 SV=1 |
| 504 | 1 | UniProt n | Q9D0J8 | 143 | 15682 | 5 | 5 | 2 | 2 | 0.7 | 1 | 1.14 | 1.031 | 0.772 | 0.78 | 0.632 | 1.055 | 0.725 | Parathymosin OS=Mus musculus GN=Ptms PE=1 SV=3 |
| 751 | 1 | UniProt n | Q3TPM3 | 73 | 98992 | 1 | 1 | 1 | 1 | 0.04 | 1 | 0.931 | 0.957 | 0.685 | 0.764 | 0.727 | 0.962 | 0.725 | Cullin-1 (Fragment) OS=Mus musculus GN=Cul1 PE=1 SV=1 |
| 956 | 1 | UniProt n | P61924 | 49 | 23882 | 1 | 1 | 1 | 1 | 0.19 | 1 | 0.896 | 0.972 | 0.733 | 0.731 | 0.708 | 0.955 | 0.724 | Coatomer subunit zeta-1 OS=Mus musculus GN=Copz1 PE=1 SV=1 |
| 438 | 1 | UniProt n | G5E866 | 177 | 172193 | 3 | 3 | 2 | 2 | 0.05 | 1 | 0.867 | 0.81 | 0.763 | 0.683 | 0.727 | 0.889 | 0.724 | Splicing factor 3B subunit 1 OS=Mus musculus GN=Sf3b1 PE=1 SV=1 |
| 408 | 1 | UniProt n | P28063 | 203 | 33513 | 10 | 10 | 7 | 7 | 1.41 | 1 | 0.96 | 0.97 | 0.734 | 0.707 | 0.73 | 0.977 | 0.724 | Proteasome subunit beta type-8 OS=Mus musculus GN=Psmb8 PE=1 SV=2 |
| 771 | 1 | UniProt n | Q9CQF3 | 71 | 30833 | 4 | 4 | 3 | 3 | 0.5 | 1 | 0.983 | 0.962 | 0.761 | 0.72 | 0.69 | 0.982 | 0.723 | Cleavage and polyadenylation specificity factor subunit 5 OS=Mus musculus GN=Nudt21 PE=1 SV=1 |
| 171 | 1 | UniProt n | P97927 | 547 | 235491 | 17 | 17 | 11 | 11 | 0.24 | 1 | 0.955 | 0.94 | 0.718 | 0.72 | 0.731 | 0.965 | 0.723 | Laminin subunit alpha-4 OS=Mus musculus GN=Lama4 PE=1 SV=2 |
| 949 | 1 | UniProt n | B1AVK5 | 50 | 191972 | 2 | 2 | 2 | 2 | 0.05 | 1 | 1.115 | 1.133 | 0.827 | 0.762 | 0.599 | 1.081 | 0.723 | Collagen, type IV, alpha 6 OS=Mus musculus GN=Col4a6 PE=1 SV=1 |
| 305 | 1 | UniProt n | P10518 | 289 | 40019 | 9 | 9 | 4 | 4 | 0.88 | 1 | 1.032 | 0.993 | 0.718 | 0.737 | 0.713 | 1.008 | 0.723 | Delta-aminolevulinic acid dehydratase OS=Mus musculus GN=Alad PE=1 SV=1 |
| 1133 | 1 | UniProt n | P62305 | 33 | 12972 | 1 | 1 | 1 | 1 | 0.37 | 1 | 0.736 | 0.872 | 0.662 | 0.824 | 0.691 | 0.863 | 0.722 | Small nuclear ribonucleoprotein E OS=Mus musculus GN=Snrpe PE=1 SV=1 |
| 83 | 1 | UniProt n | O88569 | 1175 | 43510 | 56 | 56 | 12 | 12 | 2.88 | 1 | 1.025 | 0.991 | 0.74 | 0.726 | 0.698 | 1.005 | 0.721 | Heterogeneous nuclear ribonucleoproteins A2/B1 OS=Mus musculus GN=Hnrnpa2b1 PE=1 SV=2 |
| 35 | 2 | UniProt n | E9Q8B5 | 149 | 122376 | 7 | 7 | 3 | 3 | 0.15 | 1 | 0.983 | 0.978 | 0.755 | 0.666 | 0.745 | 0.987 | 0.721 | Predicted gene 4788 OS=Mus musculus GN=Gm4788 PE=1 SV=1 |
| 875 | 1 | UniProt n | P29595 | 57 | 12009 | 2 | 2 | 1 | 1 | 0.41 | 1 | 1.053 | 0.955 | 0.762 | 0.822 | 0.598 | 1.002 | 0.721 | NEDD8 OS=Mus musculus GN=Nedd8 PE=1 SV=2 |
| 1269 | 1 | UniProt n | H3BKY9 | 20 | 12530 | 1 | 1 | 1 | 1 | 0.39 | 1 | 0.811 | 0.649 | 0.852 | 0.702 | 0.626 | 0.807 | 0.721 | Ubiquitin-like modifier-activating enzyme 7 (Fragment) OS=Mus musculus GN=Uba7 PE=4 SV=1 |
| 1012 | 1 | UniProt n | Z4YJL4 | 44 | 125088 | 1 | 1 | 1 | 1 | 0.03 | 1 | 0.886 | 0.855 | 0.779 | 0.651 | 0.738 | 0.912 | 0.721 | 182 kDa tankyrase-1-binding protein OS=Mus musculus GN=Tnks1bp1 PE=1 SV=1 |
| 86 | 5 | UniProt n | P58771-2 | 1008 | 44599 | 61 | 61 | 13 | 13 | 3.13 | 1 | 1.014 | 0.961 | 0.758 | 0.738 | 0.669 | 0.991 | 0.721 | Isoform 2 of Tropomyosin alpha-1 chain OS=Mus musculus GN=Tpm1 |
| 289 | 1 | UniProt n | Q60972 | 303 | 55157 | 20 | 20 | 7 | 7 | 0.85 | 1 | 0.982 | 0.988 | 0.736 | 0.697 | 0.729 | 0.990 | 0.720 | Histone-binding protein RBBP4 OS=Mus musculus GN=Rbbp4 PE=1 SV=5 |
| 770 | 1 | UniProt n | A0A0R4J1 | 71 | 80261 | 3 | 3 | 2 | 2 | 0.11 | 1 | 1.002 | 1.028 | 0.787 | 0.698 | 0.68 | 1.010 | 0.720 | Drebrin OS=Mus musculus GN=Dbn1 PE=1 SV=1 |
| 86 | 1 | UniProt n | P21107-2 | 1147 | 36488 | 66 | 66 | 13 | 13 | 5.33 | 1 | 1.007 | 0.977 | 0.746 | 0.736 | 0.68 | 0.995 | 0.720 | Isoform 2 of Tropomyosin alpha-3 chain OS=Mus musculus GN=Tpm3 |

| | | | | | | | | | | | | | | | | | | | | |
|---|---|---|---|---|---|---|---|---|---|---|---|---|---|---|---|---|---|---|---|---|
| 996 | 1 | UniProt | n | D3YX34 | 45 | 151585 | 2 | 2 | 2 | 2 | 0.06 | 1 | 0.873 | 0.991 | 0.712 | 0.782 | 0.67 | 0.953 | 0.720 | Dynactin subunit 1 OS=Mus musculus GN=Dctn1 PE=1 SV=1 |
| 142 | 1 | UniProt | n | P14152 | 703 | 46056 | 36 | 36 | 11 | 11 | 2 | 1 | 1.031 | 0.998 | 0.719 | 0.698 | 0.743 | 1.010 | 0.720 | Malate dehydrogenase, cytoplasmic OS=Mus musculus GN=Mdh1 PE=1 SV=3 |
| 229 | 1 | UniProt | n | O09061 | 381 | 30798 | 14 | 14 | 6 | 6 | 1.98 | 1 | 1.058 | 1.04 | 0.702 | 0.705 | 0.753 | 1.032 | 0.720 | Proteasome subunit beta type-1 OS=Mus musculus GN=Psmb1 PE=1 SV=1 |
| 151 | 1 | UniProt | n | A0A0R4J1 | 665 | 85560 | 19 | 19 | 9 | 9 | 0.56 | 1 | 0.995 | 1.024 | 0.733 | 0.737 | 0.689 | 1.006 | 0.719 | Elongation factor 1-delta OS=Mus musculus GN=Eef1d PE=1 SV=1 |
| 122 | 1 | UniProt | n | Q99LB4 | 828 | 45971 | 22 | 22 | 7 | 7 | 1.5 | 1 | 0.972 | 0.957 | 0.734 | 0.704 | 0.72 | 0.976 | 0.719 | Capping protein (Actin filament), gelsolin-like OS=Mus musculus GN=Capg PE=1 SV=1 |
| 248 | 1 | UniProt | n | Q6ZQ38 | 354 | 161352 | 23 | 23 | 17 | 17 | 0.61 | 1 | 0.936 | 0.968 | 0.7 | 0.725 | 0.733 | 0.968 | 0.719 | Cullin-associated NEDD8-dissociated protein 1 OS=Mus musculus GN=Cand1 PE=1 SV=2 |
| 646 | 1 | UniProt | n | Q9R1P3 | 91 | 26680 | 2 | 2 | 2 | 2 | 0.37 | 1 | 0.954 | 1.071 | 0.639 | 0.874 | 0.666 | 1.007 | 0.719 | Proteasome subunit beta type-2 OS=Mus musculus GN=Psmb2 PE=1 SV=1 |
| 913 | 1 | UniProt | n | Q8BKC5 | 54 | 142959 | 1 | 1 | 1 | 1 | 0.03 | 1 | 0.885 | 0.945 | 0.732 | 0.731 | 0.695 | 0.942 | 0.719 | Importin-5 OS=Mus musculus GN=Ipo5 PE=1 SV=3 |
| 564 | 1 | UniProt | n | Q8K183 | 118 | 40698 | 3 | 3 | 2 | 2 | 0.23 | 1 | 0.702 | 0.811 | 0.907 | 0.724 | 0.566 | 0.829 | 0.719 | Pyridoxal kinase OS=Mus musculus GN=Pdxk PE=1 SV=1 |
| 118 | 1 | UniProt | n | P27773 | 848 | 72829 | 56 | 56 | 23 | 23 | 3.04 | 1 | 1.035 | 0.991 | 0.712 | 0.725 | 0.719 | 1.008 | 0.719 | Protein disulfide-isomerase A3 OS=Mus musculus GN=Pdia3 PE=1 SV=2 |
| 339 | 1 | UniProt | n | P97315 | 250 | 28560 | 9 | 9 | 7 | 7 | 1.8 | 1 | 0.952 | 0.909 | 0.722 | 0.737 | 0.696 | 0.953 | 0.718 | Cysteine and glycine-rich protein 1 OS=Mus musculus GN=Csrp1 PE=1 SV=3 |
| 388 | 1 | UniProt | n | P40336 | 215 | 47004 | 12 | 12 | 6 | 6 | 0.71 | 1 | 0.921 | 0.933 | 0.716 | 0.723 | 0.715 | 0.951 | 0.718 | Vacuolar protein sorting-associated protein 26A OS=Mus musculus GN=Vps26a PE=1 SV=1 |
| 538 | 2 | UniProt | n | Q8BH64 | 109 | 73442 | 5 | 5 | 3 | 3 | 0.19 | 1 | 0.952 | 0.9 | 0.712 | 0.705 | 0.737 | 0.950 | 0.718 | EH domain-containing protein 2 OS=Mus musculus GN=Ehd2 PE=1 SV=1 |
| 217 | 1 | UniProt | n | Q921M7 | 404 | 43371 | 10 | 10 | 6 | 6 | 0.79 | 1 | 0.992 | 0.96 | 0.734 | 0.735 | 0.685 | 0.984 | 0.718 | Protein FAM49B OS=Mus musculus GN=Fam49b PE=1 SV=1 |
| 429 | 1 | UniProt | n | D3Z0Y2 | 184 | 28960 | 11 | 11 | 4 | 4 | 1.06 | 1 | 1.062 | 1.051 | 0.717 | 0.689 | 0.747 | 1.037 | 0.717 | Peroxiredoxin-6 OS=Mus musculus GN=Prdx6 PE=1 SV=1 |
| 207 | 1 | UniProt | n | O88844 | 429 | 58830 | 25 | 25 | 11 | 11 | 1.2 | 1 | 0.994 | 0.985 | 0.71 | 0.729 | 0.711 | 0.993 | 0.717 | Isocitrate dehydrogenase [NADP] cytoplasmic OS=Mus musculus GN=Idh1 PE=1 SV=2 |
| 756 | 1 | UniProt | n | P62317 | 72 | 19086 | 4 | 4 | 3 | 3 | 0.92 | 1 | 1.019 | 0.992 | 0.738 | 0.707 | 0.704 | 1.004 | 0.716 | Small nuclear ribonucleoprotein Sm D2 OS=Mus musculus GN=Snrpd2 PE=1 SV=1 |
| 1137 | 1 | UniProt | n | A0A0G2JE | 33 | 18433 | 3 | 3 | 1 | 1 | 0.25 | 1 | 0.909 | 0.904 | 0.801 | 0.72 | 0.636 | 0.937 | 0.716 | 2-oxoglutarate and iron-dependent oxygenase domain-containing protein 2 (Fragment) OS=Mus musculus GN=Ogfod2 PE=4 SV=1 |
| 1116 | 1 | UniProt | n | D6REG4 | 34 | 24171 | 1 | 1 | 1 | 1 | 0.19 | 1 | 0.742 | 0.784 | 0.796 | 0.583 | 0.789 | 0.835 | 0.715 | Glyoxylate reductase/hydroxypyruvate reductase OS=Mus musculus GN=Grhpr PE=1 SV=1 |
| 86 | 4 | UniProt | n | P58771 | 1025 | 44875 | 60 | 60 | 12 | 12 | 2.73 | 1 | 1.014 | 0.959 | 0.747 | 0.737 | 0.665 | 0.991 | 0.715 | Tropomyosin alpha-1 chain OS=Mus musculus GN=Tpm1 PE=1 SV=1 |
| 785 | 1 | UniProt | n | A2AUR7 | 68 | 36521 | 4 | 4 | 4 | 4 | 0.59 | 1 | 0.996 | 0.922 | 0.695 | 0.709 | 0.742 | 0.972 | 0.715 | Ras suppressor protein 1 OS=Mus musculus GN=Rsu1 PE=1 SV=1 |
| 327 | 1 | UniProt | n | Q9JLI2 | 264 | 198623 | 10 | 10 | 5 | 5 | 0.16 | 1 | 0.979 | 0.992 | 0.689 | 0.734 | 0.722 | 0.990 | 0.715 | Collagen type V alpha 3 chain OS=Mus musculus GN=Col5a3 PE=1 SV=1 |
| 740 | 1 | UniProt | n | Q8BK64 | 75 | 47099 | 4 | 4 | 3 | 3 | 0.31 | 1 | 0.97 | 0.959 | 0.764 | 0.694 | 0.685 | 0.976 | 0.713 | Activator of 90 kDa heat shock protein ATPase homolog 1 OS=Mus musculus GN=Ahsa1 PE=1 SV=2 |
| 58 | 3 | UniProt | n | P26040 | 288 | 86491 | 21 | 21 | 12 | 12 | 0.99 | 1 | 0.986 | 0.983 | 0.71 | 0.683 | 0.748 | 0.990 | 0.713 | Ezrin OS=Mus musculus GN=Ezr PE=1 SV=3 |
| 989 | 1 | UniProt | n | Q61024 | 46 | 76611 | 2 | 2 | 2 | 2 | 0.12 | 1 | 0.891 | 0.865 | 0.693 | 0.745 | 0.702 | 0.917 | 0.713 | Asparagine synthetase [glutamine-hydrolyzing] OS=Mus musculus GN=Asns PE=1 SV=3 |
| 739 | 1 | UniProt | n | Q9D0W5 | 75 | 21663 | 3 | 3 | 2 | 2 | 0.78 | 1 | 1.003 | 1.008 | 0.69 | 0.783 | 0.669 | 1.004 | 0.712 | Peptidyl-prolyl cis-trans isomerase-like 1 OS=Mus musculus GN=Ppil1 PE=1 SV=1 |
| 231 | 1 | UniProt | n | Q9CQW1 | 376 | 27610 | 7 | 7 | 3 | 3 | 0.58 | 1 | 0.966 | 0.982 | 0.722 | 0.746 | 0.671 | 0.983 | 0.712 | Synaptobrevin homolog YKT6 OS=Mus musculus GN=Ykt6 PE=1 SV=1 |
| 830 | 1 | UniProt | n | Q78PY7 | 62 | 120220 | 3 | 3 | 3 | 3 | 0.11 | 1 | 0.793 | 0.918 | 0.756 | 0.653 | 0.731 | 0.900 | 0.712 | Staphylococcal nuclease domain-containing protein 1 OS=Mus musculus GN=Snd1 PE=1 SV=1 |
| 256 | 1 | UniProt | n | Q924B0 | 347 | 36207 | 14 | 14 | 7 | 7 | 1.26 | 1 | 1.023 | 1.026 | 0.7 | 0.716 | 0.72 | 1.016 | 0.712 | Inositol-1-monophosphatase OS=Mus musculus GN=Impa1 PE=1 SV=1 |
| 669 | 1 | UniProt | n | Q9CQU0 | 89 | 22825 | 2 | 2 | 2 | 2 | 0.44 | 1 | 1.035 | 0.983 | 0.658 | 0.756 | 0.725 | 1.006 | 0.712 | Thioredoxin domain-containing protein 12 OS=Mus musculus GN=Txndc12 PE=1 SV=1 |
| 19 | 14 | UniProt | n | G3X981 | 252 | 64963 | 18 | 18 | 4 | 4 | 0.39 | 1 | 0.929 | 0.844 | 0.785 | 0.741 | 0.62 | 0.922 | 0.712 | Peripherin OS=Mus musculus GN=Prph PE=1 SV=1 |
| 453 | 1 | UniProt | n | A0A0G2JG | 171 | 11840 | 19 | 19 | 4 | 4 | 4.65 | 1 | 1.098 | 0.997 | 0.702 | 0.731 | 0.702 | 1.031 | 0.712 | Protein S100-A4 (Fragment) OS=Mus musculus GN=S100a4 PE=1 SV=1 |
| 3 | 5 | UniProt | n | P28481 | 468 | 162838 | 32 | 32 | 4 | 4 | 0.14 | 1 | 1.182 | 1.109 | 0.676 | 0.718 | 0.74 | 1.094 | 0.711 | Collagen alpha-1(II) chain OS=Mus musculus GN=Col2a1 PE=1 SV=2 |
| 30 | 1 | UniProt | n | Q9QXS1-3 | 3134 | 598449 | 170 | 170 | 99 | 99 | 1.05 | 1 | 0.963 | 0.975 | 0.727 | 0.7 | 0.704 | 0.979 | 0.710 | Isoform PLEC-1A of Plectin OS=Mus musculus GN=Plec |
| 56 | 1 | UniProt | n | Q01853 | 1627 | 104419 | 64 | 64 | 28 | 28 | 2.67 | 1 | 1.034 | 1.063 | 0.733 | 0.691 | 0.706 | 1.032 | 0.710 | Transitional endoplasmic reticulum ATPase OS=Mus musculus GN=Vcp PE=1 SV=4 |
| 58 | 2 | UniProt | n | P26043 | 453 | 88366 | 29 | 29 | 17 | 17 | 1.49 | 1 | 0.99 | 0.989 | 0.714 | 0.684 | 0.732 | 0.993 | 0.710 | Radixin OS=Mus musculus GN=Rdx PE=1 SV=3 |
| 210 | 1 | UniProt | n | Q9R1P0 | 417 | 37591 | 17 | 17 | 6 | 6 | 1.19 | 1 | 1.03 | 1.009 | 0.715 | 0.684 | 0.729 | 1.013 | 0.709 | Proteasome subunit alpha type-4 OS=Mus musculus GN=Psma4 PE=1 SV=1 |
| 1086 | 1 | UniProt | n | A2A699 | 37 | 93793 | 1 | 1 | 1 | 1 | 0.05 | 1 | 0.809 | 0.766 | 0.653 | 0.725 | 0.753 | 0.853 | 0.709 | Protein FAM171A2 OS=Mus musculus GN=Fam171a2 PE=1 SV=1 |
| 801 | 1 | UniProt | n | D3Z0F5 | 63 | 38971 | 2 | 2 | 2 | 2 | 0.24 | 1 | 0.987 | 1.004 | 0.757 | 0.626 | 0.752 | 0.997 | 0.709 | COP9 signalosome complex subunit 6 OS=Mus musculus GN=Cops6 PE=1 SV=1 |
| 754 | 1 | UniProt | n | Q8R326 | 72 | 66418 | 1 | 1 | 1 | 1 | 0.07 | 1 | 0.991 | 0.904 | 0.684 | 0.748 | 0.695 | 0.964 | 0.708 | Paraspeckle component 1 OS=Mus musculus GN=Pspc1 PE=1 SV=1 |
| 552 | 1 | UniProt | n | Q9Z1B5 | 121 | 28026 | 3 | 3 | 2 | 2 | 0.57 | 1 | 0.99 | 0.941 | 0.749 | 0.751 | 0.632 | 0.977 | 0.708 | Mitotic spindle assembly checkpoint protein MAD2A OS=Mus musculus GN=Mad2l1 PE=1 SV=2 |
| 315 | 1 | UniProt | n | P63085 | 275 | 48872 | 13 | 13 | 8 | 8 | 1 | 1 | 0.961 | 0.958 | 0.731 | 0.672 | 0.722 | 0.973 | 0.708 | Mitogen-activated protein kinase 1 OS=Mus musculus GN=Mapk1 PE=1 SV=3 |
| 1087 | 1 | UniProt | n | Q9JMG1 | 37 | 23051 | 2 | 2 | 1 | 1 | 0.44 | 1 | 0.967 | 0.994 | 0.685 | 0.725 | 0.714 | 0.987 | 0.708 | Endothelial differentiation-related factor 1 OS=Mus musculus GN=Edf1 PE=1 SV=1 |
| 535 | 1 | UniProt | n | Q5SUS9 | 127 | 75004 | 9 | 9 | 4 | 4 | 0.25 | 1 | 0.957 | 0.948 | 0.722 | 0.71 | 0.691 | 0.968 | 0.708 | RNA-binding protein EWS OS=Mus musculus GN=Ewsr1 PE=1 SV=1 |
| 1202 | 1 | UniProt | n | A0A140LH | 27 | 42541 | 1 | 1 | 1 | 1 | 0.1 | 1 | 0.846 | 0.926 | 0.671 | 0.668 | 0.79 | 0.922 | 0.707 | Mitotic checkpoint protein BUB3 OS=Mus musculus GN=Bub3 PE=1 SV=1 |
| 607 | 1 | UniProt | n | P23591 | 102 | 42123 | 2 | 2 | 2 | 2 | 0.22 | 1 | 0.977 | 1.024 | 0.607 | 0.688 | 0.847 | 1.000 | 0.707 | GDP-L-fucose synthase OS=Mus musculus GN=Tsta3 PE=1 SV=3 |
| 86 | 3 | UniProt | n | E9Q7Q3 | 1105 | 36672 | 65 | 65 | 10 | 10 | 2.97 | 1 | 1.01 | 0.973 | 0.728 | 0.728 | 0.666 | 0.994 | 0.707 | Tropomyosin alpha-3 chain OS=Mus musculus GN=Tpm3 PE=1 SV=1 |
| 425 | 1 | UniProt | n | P11031 | 186 | 20503 | 14 | 14 | 5 | 5 | 1.76 | 1 | 0.939 | 0.961 | 0.731 | 0.702 | 0.686 | 0.966 | 0.706 | Activated RNA polymerase II transcriptional coactivator p15 OS=Mus musculus GN=Sub1 PE=1 SV=3 |
| 352 | 1 | UniProt | n | Q9CY58-2 | 240 | 53307 | 10 | 10 | 7 | 7 | 0.74 | 1 | 1.098 | 1.057 | 0.723 | 0.699 | 0.694 | 1.051 | 0.705 | Isoform 2 of Plasminogen activator inhibitor 1 RNA-binding protein OS=Mus musculus GN=Serbp1 |
| 326 | 1 | UniProt | n | O35215 | 265 | 14986 | 11 | 11 | 7 | 7 | 5.93 | 1 | 0.991 | 0.881 | 0.727 | 0.714 | 0.675 | 0.956 | 0.705 | D-dopachrome decarboxylase OS=Mus musculus GN=Ddt PE=1 SV=3 |
| 291 | 1 | UniProt | n | Q99KJ8 | 303 | 54220 | 9 | 9 | 5 | 5 | 0.48 | 1 | 0.926 | 0.906 | 0.754 | 0.703 | 0.661 | 0.943 | 0.705 | Dynactin subunit 2 OS=Mus musculus GN=Dctn2 PE=1 SV=3 |
| 480 | 1 | UniProt | n | Q62093 | 155 | 29416 | 4 | 4 | 2 | 2 | 0.33 | 1 | 1.686 | 1.104 | 0.801 | 0.662 | 0.66 | 1.230 | 0.705 | Serine/arginine-rich splicing factor 2 OS=Mus musculus GN=Srsf2 PE=1 SV=4 |
| 1204 | 1 | UniProt | n | O54941 | 27 | 56086 | 1 | 1 | 1 | 1 | 0.08 | 1 | 0.888 | 0.94 | 0.672 | 0.745 | 0.699 | 0.942 | 0.705 | SWI/SNF-related matrix-associated actin-dependent regulator of chromatin subfamily E member 1 OS=Mus musculus GN=Smarce1 PE=1 SV=1 |
| 31 | 1 | UniProt | n | Q61554 | 3066 | 361843 | 122 | 122 | 51 | 51 | 1.02 | 1 | 0.952 | 0.954 | 0.722 | 0.689 | 0.703 | 0.968 | 0.705 | Fibrillin-1 OS=Mus musculus GN=Fbn1 PE=1 SV=2 |
| 1238 | 1 | UniProt | n | Q8BLB0 | 24 | 70740 | 1 | 1 | 1 | 1 | 0.06 | 1 | 0.669 | 0.558 | 0.928 | 0.781 | 0.482 | 0.720 | 0.704 | Zinc finger protein 3 OS=Mus musculus GN=Zfp3 PE=2 SV=1 |
| 273 | 1 | UniProt | n | P61089 | 325 | 20215 | 18 | 18 | 7 | 7 | 3.24 | 1 | 0.928 | 0.842 | 0.71 | 0.732 | 0.672 | 0.921 | 0.704 | Ubiquitin-conjugating enzyme E2 N OS=Mus musculus GN=Ube2n PE=1 SV=1 |
| 270 | 1 | UniProt | n | Q91YR1 | 327 | 48756 | 9 | 9 | 4 | 4 | 0.54 | 1 | 0.978 | 1.027 | 0.772 | 0.67 | 0.675 | 1.001 | 0.704 | Twinfilin-1 OS=Mus musculus GN=Twf1 PE=1 SV=2 |
| 618 | 1 | UniProt | n | Q6ZWX6 | 101 | 43008 | 8 | 8 | 5 | 5 | 0.63 | 1 | 0.962 | 1.003 | 0.747 | 0.694 | 0.673 | 0.988 | 0.704 | Eukaryotic translation initiation factor 2 subunit 1 OS=Mus musculus GN=Eif2s1 PE=1 SV=3 |
| 579 | 1 | UniProt | n | Q9JHU9 | 112 | 66875 | 8 | 8 | 6 | 6 | 0.46 | 1 | 0.963 | 0.916 | 0.695 | 0.708 | 0.709 | 0.959 | 0.704 | Inositol-3-phosphate synthase 1 OS=Mus musculus GN=Isyna1 PE=1 SV=1 |
| 600 | 1 | UniProt | n | P62869 | 103 | 15641 | 5 | 5 | 3 | 3 | 1.89 | 1 | 0.931 | 0.946 | 0.726 | 0.66 | 0.727 | 0.959 | 0.704 | Elongin-B OS=Mus musculus GN=Elob PE=1 SV=1 |
| 1205 | 1 | UniProt | n | Q91WK2 | 27 | 48159 | 1 | 1 | 1 | 1 | 0.09 | 1 | 0.701 | 0.869 | 0.724 | 0.717 | 0.671 | 0.848 | 0.704 | Eukaryotic translation initiation factor 3 subunit H OS=Mus musculus GN=Eif3h PE=1 SV=1 |
| 802 | 1 | UniProt | n | Q8BH40 | 65 | 34723 | 2 | 2 | 1 | 1 | 0.27 | 1 | 0.902 | 1.01 | 0.831 | 0.681 | 0.613 | 0.969 | 0.703 | Syntaxin-7 OS=Mus musculus GN=Stx7 PE=1 SV=1 |
| 1121 | 1 | UniProt | n | D3YVR4 | 34 | 21336 | 2 | 2 | 1 | 1 | 0.22 | 1 | 1.006 | 0.976 | 0.612 | 1.293 | 0.438 | 0.994 | 0.702 | Mesoderm development LRP chaperone (Fragment) OS=Mus musculus GN=Mesd PE=1 SV=1 |
| 886 | 1 | UniProt | n | A2A7S7 | 56 | 82145 | 3 | 3 | 3 | 3 | 0.17 | 1 | 0.832 | 0.902 | 0.747 | 0.707 | 0.656 | 0.909 | 0.702 | Tyrosine--tRNA ligase OS=Mus musculus GN=Yars PE=1 SV=1 |
| 378 | 1 | UniProt | n | A2AVR9 | 224 | 14354 | 7 | 7 | 3 | 3 | 1.38 | 1 | 0.969 | 0.896 | 0.714 | 0.741 | 0.653 | 0.954 | 0.702 | Dynein light chain roadblock OS=Mus musculus GN=Dynlrb1 PE=1 SV=1 |
| 906 | 1 | UniProt | n | Q8BTU1 | 54 | 28043 | 2 | 2 | 2 | 2 | 0.35 | 1 | 0.9 | 0.985 | 0.691 | 0.696 | 0.717 | 0.961 | 0.701 | Cilia- and flagella-associated protein 20 OS=Mus musculus GN=Cfap20 PE=1 SV=1 |
| 81 | 1 | UniProt | n | P08249 | 1199 | 43866 | 38 | 38 | 13 | 13 | 2.85 | 1 | 1.006 | 0.99 | 0.686 | 0.686 | 0.731 | 0.999 | 0.701 | Malate dehydrogenase, mitochondrial OS=Mus musculus GN=Mdh2 PE=1 SV=3 |
| 681 | 1 | UniProt | n | A0A087W | 86 | 9657 | 1 | 1 | 1 | 1 | 0.53 | 1 | 0.952 | 0.894 | 0.742 | 0.656 | 0.705 | 0.948 | 0.700 | COP9 signalosome complex subunit 8 (Fragment) OS=Mus musculus GN=Cops8 PE=1 SV=6 |
| 478 | 1 | UniProt | n | Q8CHP8 | 156 | 39145 | 6 | 6 | 5 | 5 | 0.71 | 1 | 0.956 | 0.987 | 0.744 | 0.702 | 0.653 | 0.981 | 0.699 | Glycerol-3-phosphate phosphatase OS=Mus musculus GN=Pgp PE=1 SV=1 |
| 837 | 1 | UniProt | n | Q9WVF5 | 61 | 85546 | 1 | 1 | 1 | 1 | 0.05 | 1 | 0.963 | 1.049 | 0.606 | 0.735 | 0.765 | 1.003 | 0.698 | Epidermal growth factor receptor OS=Mus musculus GN=Egfr PE=1 SV=1 |
| 1131 | 1 | UniProt | n | E9Q0U7 | 33 | 115171 | 1 | 1 | 1 | 1 | 0.04 | 1 | 0.847 | 0.891 | 0.729 | 0.755 | 0.619 | 0.910 | 0.698 | Heat shock protein 105 kDa OS=Mus musculus GN=Hsph1 PE=1 SV=1 |
| 1265 | 1 | UniProt | n | Q62312 | 20 | 80138 | 1 | 1 | 1 | 1 | 0.05 | 1 | 0.888 | 0.765 | 0.689 | 0.627 | 0.787 | 0.879 | 0.698 | TGF-beta receptor type-2 OS=Mus musculus GN=Tgfbr2 PE=1 SV=1 |
| 456 | 1 | UniProt | n | Q4VWZ5 | 169 | 20436 | 5 | 5 | 2 | 2 | 0.85 | 1 | 0.966 | 0.814 | 0.664 | 0.859 | 0.596 | 0.923 | 0.698 | Acyl-CoA-binding protein OS=Mus musculus GN=Dbi PE=1 SV=1 |
| 455 | 1 | UniProt | n | G3XA48 | 170 | 39213 | 5 | 5 | 4 | 4 | 0.54 | 1 | 0.9 | 0.903 | 0.737 | 0.701 | 0.657 | 0.933 | 0.698 | Isopentenyl-diphosphate Delta-isomerase 1 OS=Mus musculus GN=Idi1 PE=1 SV=1 |
| 1150 | 1 | UniProt | n | Q64442 | 32 | 46594 | 2 | 2 | 1 | 1 | 0.09 | 1 | 0.946 | 0.963 | 0.724 | 0.747 | 0.627 | 0.969 | 0.697 | Sorbitol dehydrogenase OS=Mus musculus GN=Sord PE=1 SV=3 |
| 86 | 7 | UniProt | n | P58774-2 | 786 | 43631 | 35 | 35 | 7 | 7 | 0.97 | 1 | 0.989 | 0.921 | 0.718 | 0.724 | 0.652 | 0.969 | 0.697 | Isoform 2 of Tropomyosin beta chain OS=Mus musculus GN=Tpm2 |
| 86 | 6 | UniProt | n | A2AIM4 | 799 | 44275 | 35 | 35 | 8 | 8 | 1.14 | 1 | 0.99 | 0.922 | 0.719 | 0.724 | 0.651 | 0.970 | 0.697 | Tropomyosin beta chain OS=Mus musculus GN=Tpm2 PE=1 SV=1 |
| 835 | 1 | UniProt | n | A0A1B0GI | 61 | 41920 | 2 | 2 | 1 | 1 | 0.11 | 1 | 1.091 | 1.068 | 0.604 | 0.641 | 0.874 | 1.052 | 0.697 | 3'(2'),5'-bisphosphate nucleotidase 1 OS=Mus musculus GN=Bpnt1 PE=1 SV=1 |
| 935 | 1 | UniProt | n | D5MCW4 | 51 | 18664 | 1 | 1 | 1 | 1 | 0.25 | 1 | 0.811 | 0.856 | 0.732 | 0.724 | 0.638 | 0.885 | 0.697 | Protein CutA OS=Mus musculus GN=Cuta PE=1 SV=1 |
| 281 | 1 | UniProt | n | Q9Z1Q5 | 318 | 32747 | 10 | 10 | 7 | 7 | 1.46 | 1 | 0.897 | 0.892 | 0.675 | 0.678 | 0.738 | 0.928 | 0.696 | Chloride intracellular channel protein 1 OS=Mus musculus GN=Clic1 PE=1 SV=3 |
| 968 | 1 | UniProt | n | A0A1Y7VI | 48 | 13727 | 3 | 3 | 2 | 2 | 0.82 | 1 | 0.964 | 0.942 | 0.682 | 0.738 | 0.67 | 0.968 | 0.696 | Glutaredoxin-related protein 5, mitochondrial OS=Mus musculus GN=Glrx5 PE=4 SV=1 |
| 90 | 1 | UniProt | n | E9PZF0 | 1111 | 37011 | 72 | 72 | 8 | 8 | 2.5 | 1 | 1.131 | 1.047 | 0.678 | 0.725 | 0.685 | 1.058 | 0.696 | Nucleoside diphosphate kinase OS=Mus musculus GN=Gm20390 PE=3 SV=1 |
| 90 | 2 | UniProt | n | P15532 | 940 | 21244 | 61 | 61 | 8 | 8 | 9.63 | 1 | 1.121 | 1.038 | 0.68 | 0.73 | 0.678 | 1.052 | 0.696 | Nucleoside diphosphate kinase A OS=Mus musculus GN=Nme1 PE=1 SV=1 |
| 277 | 1 | UniProt | n | P99026 | 323 | 31015 | 13 | 13 | 3 | 3 | 1.26 | 1 | 0.91 | 0.836 | 0.731 | 0.756 | 0.609 | 0.913 | 0.696 | Proteasome subunit beta type-4 OS=Mus musculus GN=Psmb4 PE=1 SV=1 |
| 391 | 1 | UniProt | n | Q5XJY5 | 212 | 69425 | 11 | 11 | 9 | 9 | 0.73 | 1 | 0.923 | 0.932 | 0.692 | 0.74 | 0.657 | 0.951 | 0.696 | Coatomer subunit delta OS=Mus musculus GN=Arcn1 PE=1 SV=2 |
| 1174 | 1 | UniProt | n | Q6PAJ1 | 30 | 169013 | 1 | 1 | 1 | 1 | 0.03 | 1 | 0.63 | 0.766 | 0.97 | 0.762 | 0.455 | 0.784 | 0.695 | Breakpoint cluster region protein OS=Mus musculus GN=Bcr PE=1 SV=3 |
| 205 | 1 | UniProt | n | Q62465 | 436 | 49337 | 12 | 12 | 6 | 6 | 0.82 | 1 | 1.018 | 0.957 | 0.706 | 0.654 | 0.728 | 0.991 | 0.695 | Synaptic vesicle membrane protein VAT-1 homolog OS=Mus musculus GN=Vat1 PE=1 SV=3 |
| 548 | 1 | UniProt | n | Q810A7 | 123 | 121601 | 3 | 3 | 3 | 3 | 0.11 | 1 | 0.945 | 0.94 | 0.723 | 0.69 | 0.673 | 0.961 | 0.695 | ATP-dependent RNA helicase DDX42 OS=Mus musculus GN=Ddx42 PE=1 SV=3 |
| 820 | 1 | UniProt | n | Q8BGD9 | 63 | 80497 | 3 | 3 | 3 | 3 | 0.17 | 1 | 0.934 | 0.92 | 0.749 | 0.685 | 0.651 | 0.951 | 0.694 | Eukaryotic translation initiation factor 4B OS=Mus musculus GN=Eif4b PE=1 SV=1 |
| 202 | 1 | UniProt | n | P05201 | 445 | 52837 | 21 | 21 | 8 | 8 | 0.9 | 1 | 0.987 | 0.963 | 0.682 | 0.679 | 0.721 | 0.983 | 0.694 | Aspartate aminotransferase, cytoplasmic OS=Mus musculus GN=Got1 PE=1 SV=3 |
| 395 | 1 | UniProt | n | E9Q1G8 | 210 | 65191 | 10 | 10 | 5 | 5 | 0.38 | 1 | 0.967 | 0.982 | 0.708 | 0.707 | 0.667 | 0.983 | 0.694 | Septin-7 OS=Mus musculus GN=Sept7 PE=1 SV=2 |
| 1194 | 1 | UniProt | n | A2AW05 | 29 | 89099 | 1 | 1 | 1 | 1 | 0.05 | 1 | 0.872 | 0.917 | 0.701 | 0.688 | 0.692 | 0.928 | 0.694 | FACT complex subunit SSRP1 (Fragment) OS=Mus musculus GN=Ssrp1 PE=1 SV=1 |
| 1212 | 1 | UniProt | n | Q6GQW0 | 27 | 137465 | 1 | 1 | 1 | 1 | 0.03 | 1 | 0.839 | 0.921 | 0.608 | 0.809 | 0.677 | 0.918 | 0.693 | Ankyrin repeat and BTB/POZ domain-containing protein BTBD11 OS=Mus musculus GN=Btbd11 PE=2 SV=2 |
| 1054 | 1 | UniProt | n | Q9CQI3 | 40 | 20805 | 2 | 2 | 2 | 2 | 0.5 | 1 | 1.011 | 0.909 | 0.7 | 0.727 | 0.654 | 0.972 | 0.693 | Glia maturation factor beta OS=Mus musculus GN=Gmfb PE=1 SV=3 |
| 737 | 1 | UniProt | n | Q6P5E4 | 75 | 206991 | 4 | 4 | 2 | 2 | 0.06 | 1 | 0.758 | 1.175 | 0.649 | 0.791 | 0.648 | 0.962 | 0.693 | UDP-glucose:glycoprotein glucosyltransferase 1 OS=Mus musculus GN=Uggt1 PE=1 SV=4 |
| 763 | 1 | UniProt | n | O88271 | 71 | 44461 | 2 | 2 | 2 | 2 | 0.21 | 1 | 0.739 | 0.864 | 0.724 | 0.691 | 0.664 | 0.861 | 0.693 | Craniofacial development protein 1 OS=Mus musculus GN=Cfdp1 PE=1 SV=1 |
| 417 | 1 | UniProt | n | Q9ESY9 | 197 | 32014 | 4 | 4 | 2 | 2 | 0.48 | 1 | 0.968 | 0.947 | 0.685 | 0.74 | 0.655 | 0.971 | 0.692 | Gamma-interferon-inducible lysosomal thiol reductase OS=Mus musculus GN=Ifi30 PE=1 SV=3 |
| 769 | 1 | UniProt | n | A0A0G2JG | 71 | 25496 | 2 | 2 | 2 | 2 | 0.39 | 1 | 1.019 | 1.023 | 0.696 | 0.661 | 0.721 | 1.014 | 0.692 | Eukaryotic translation initiation factor 4E (Fragment) OS=Mus musculus GN=Eif4e PE=1 SV=1 |
| 176 | 1 | UniProt | n | Q8C253 | 531 | 29877 | 25 | 25 | 6 | 6 | 1.68 | 1 | 0.995 | 1.004 | 0.686 | 0.675 | 0.715 | 1.000 | 0.692 | Galectin OS=Mus musculus GN=Lgals3 PE=1 SV=1 |
| 473 | 1 | UniProt | n | P62320 | 158 | 17346 | 4 | 4 | 2 | 2 | 0.62 | 1 | 0.959 | 0.969 | 0.728 | 0.687 | 0.659 | 0.976 | 0.691 | Small nuclear ribonucleoprotein Sm D3 OS=Mus musculus GN=Snrpd3 PE=1 SV=1 |
| 170 | 1 | UniProt | n | Q8CAY6 | 548 | 48027 | 16 | 16 | 7 | 7 | 1.41 | 1 | 1.078 | 1.028 | 0.689 | 0.693 | 0.69 | 1.035 | 0.691 | Acetyl-CoA acetyltransferase, cytosolic OS=Mus musculus GN=Acat2 PE=1 SV=2 |
| 800 | 1 | UniProt | n | Q3TIX9 | 65 | 80029 | 2 | 2 | 1 | 1 | 0.05 | 1 | 0.936 | 1.095 | 0.738 | 0.599 | 0.741 | 1.008 | 0.689 | U4/U6.U5 tri-snRNP-associated protein 2 OS=Mus musculus GN=Usp39 PE=1 SV=2 |
| 822 | 1 | UniProt | n | A0A0R4J0 | 62 | 70082 | 2 | 2 | 2 | 2 | 0.13 | 1 | 0.632 | 0.634 | 0.707 | 0.717 | 0.645 | 0.737 | 0.689 | UDP-N-acetylhexosamine pyrophosphorylase OS=Mus musculus GN=Uap1 PE=1 SV=1 |
| 684 | 1 | UniProt | n | Q924M7 | 85 | 55614 | 2 | 2 | 1 | 1 | 0.08 | 1 | 0.932 | 0.985 | 0.718 | 0.7 | 0.649 | 0.972 | 0.688 | Mannose-6-phosphate isomerase OS=Mus musculus GN=Mpi PE=1 SV=1 |
| 537 | 1 | UniProt | n | A0A1L1SS | 126 | 14789 | 3 | 3 | 3 | 3 | 1.32 | 1 | 1.04 | 0.966 | 0.712 | 0.753 | 0.608 | 1.002 | 0.688 | Eukaryotic translation initiation factor 1b (Fragment) OS=Mus musculus GN=Eif1b PE=1 SV=1 |
| 604 | 1 | UniProt | n | P62715 | 102 | 39663 | 3 | 3 | 2 | 2 | 0.24 | 1 | 0.93 | 1.05 | 0.704 | 0.649 | 0.713 | 0.992 | 0.688 | Serine/threonine-protein phosphatase 2A catalytic subunit beta isoform OS=Mus musculus GN=Ppp2cb PE=1 SV=1 |
| 1003 | 1 | UniProt | n | A0A087W | 44 | 10006 | 2 | 2 | 1 | 1 | 0.51 | 1 | 0.973 | 0.865 | 0.636 | 0.801 | 0.638 | 0.944 | 0.688 | TAR DNA-binding protein 43 OS=Mus musculus GN=Tardbp PE=1 SV=1 |
| 598 | 1 | UniProt | n | O35841 | 104 | 70411 | 9 | 9 | 5 | 5 | 0.35 | 1 | 0.899 | 0.932 | 0.705 | 0.685 | 0.673 | 0.943 | 0.688 | Apoptosis inhibitor 5 OS=Mus musculus GN=Api5 PE=1 SV=2 |
| 992 | 1 | UniProt | n | Q8R0H9 | 46 | 81459 | 2 | 2 | 1 | 1 | 0.05 | 1 | 0.882 | 0.89 | 0.676 | 0.736 | 0.652 | 0.922 | 0.687 | ADP-ribosylation factor-binding protein GGA1 OS=Mus musculus GN=Gga1 PE=1 SV=1 |
| 950 | 1 | UniProt | n | F8WJ30 | 50 | 17177 | 1 | 1 | 1 | 1 | 0.27 | 1 | 1.068 | 0.988 | 0.607 | 0.739 | 0.723 | 1.018 | 0.687 | Prefoldin subunit 2 (Fragment) OS=Mus musculus GN=Pfdn2 PE=1 SV=1 |
| 697 | 1 | UniProt | n | Q9D967 | 83 | 20746 | 3 | 3 | 2 | 2 | 0.5 | 1 | 0.714 | 0.774 | 0.686 | 0.738 | 0.638 | 0.821 | 0.686 | Magnesium-dependent phosphatase 1 OS=Mus musculus GN=Mdp1 PE=1 SV=1 |
| 374 | 1 | UniProt | n | Q9D0I9 | 228 | 93913 | 8 | 8 | 6 | 6 | 0.31 | 1 | 0.927 | 0.967 | 0.685 | 0.701 | 0.672 | 0.964 | 0.686 | Arginine--tRNA ligase, cytoplasmic OS=Mus musculus GN=Rars PE=1 SV=2 |
| 1034 | 1 | UniProt | n | Q62318 | 42 | 103297 | 1 | 1 | 1 | 1 | 0.04 | 1 | 0.945 | 1.027 | 0.756 | 0.658 | 0.648 | 0.990 | 0.686 | Transcription intermediary factor 1-beta OS=Mus musculus GN=Trim28 PE=1 SV=3 |
| 164 | 1 | UniProt | n | Q922R8 | 589 | 58735 | 19 | 19 | 10 | 10 | 1.05 | 1 | 0.902 | 0.899 | 0.693 | 0.686 | 0.677 | 0.933 | 0.685 | Protein disulfide-isomerase A6 OS=Mus musculus GN=Pdia6 PE=1 SV=3 |
| 91 | 1 | UniProt | n | P14211 | 1096 | 61488 | 46 | 46 | 14 | 14 | 2.22 | 1 | 1.019 | 1.003 | 0.699 | 0.678 | 0.679 | 1.007 | 0.685 | Calreticulin OS=Mus musculus GN=Calr PE=1 SV=1 |
| 19 | 16 | UniProt | n | A0A0R4J0 | 82 | 80086 | 7 | 7 | 3 | 3 | 0.17 | 1 | 0.996 | 0.939 | 0.727 | 0.685 | 0.645 | 0.978 | 0.685 | Lamin-B2 OS=Mus musculus GN=Lmnb2 PE=1 SV=1 |
| 700 | 1 | UniProt | n | A0A0A6Y\ | 82 | 6133 | 2 | 2 | 1 | 1 | 0.92 | 1 | 1.147 | 1.121 | 0.704 | 0.563 | 0.808 | 1.087 | 0.684 | BRO1 domain-containing protein BROX OS=Mus musculus GN=Brox PE=1 SV=1 |
| 602 | 1 | UniProt | n | Q9QZD9 | 103 | 42235 | 4 | 4 | 4 | 4 | 0.49 | 1 | 0.86 | 0.98 | 0.675 | 0.677 | 0.699 | 0.945 | 0.684 | Eukaryotic translation initiation factor 3 subunit I OS=Mus musculus GN=Eif3i PE=1 SV=1 |
| 1219 | 1 | UniProt | n | J3QNW0 | 26 | 206885 | 1 | 1 | 1 | 1 | 0.02 | 1 | 0.68 | 0.704 | 0.828 | 0.695 | 0.555 | 0.782 | 0.684 | DNA (cytosine-5)-methyltransferase OS=Mus musculus GN=Dnmt1 PE=1 SV=1 |
| 261 | 1 | UniProt | n | P52760 | 336 | 17030 | 10 | 10 | 5 | 5 | 3.34 | 1 | 0.979 | 0.89 | 0.679 | 0.667 | 0.705 | 0.955 | 0.683 | 2-iminobutanoate/2-iminopropanoate deaminase OS=Mus musculus GN=Rida PE=1 SV=3 |
| 921 | 1 | UniProt | n | Q9CT10 | 53 | 60984 | 2 | 2 | 2 | 2 | 0.15 | 1 | 0.901 | 0.814 | 0.751 | 0.613 | 0.693 | 0.902 | 0.683 | Ran-binding protein 3 OS=Mus musculus GN=Ranbp3 PE=1 SV=2 |
| 54 | 1 | UniProt | n | P26645 | 1695 | 36991 | 53 | 53 | 10 | 10 | 2.92 | 1 | 1.062 | 1.095 | 0.694 | 0.749 | 0.61 | 1.052 | 0.682 | Myristoylated alanine-rich C-kinase substrate OS=Mus musculus GN=Marcks PE=1 SV=2 |
| 361 | 2 | UniProt | n | G3UZR0 | 49 | 18195 | 3 | 3 | 2 | 2 | 0.58 | 1 | 0.9 | 0.843 | 0.655 | 0.672 | 0.72 | 0.912 | 0.682 | N(G),N(G)-dimethylarginine dimethylaminohydrolase 2 (Fragment) OS=Mus musculus GN=Ddah2 PE=1 SV=1 |
| 1069 | 1 | UniProt | n | A0A087W | 39 | 19888 | 2 | 2 | 1 | 1 | 0.23 | 1 | 1.065 | 1.036 | 0.63 | 0.723 | 0.695 | 1.033 | 0.682 | Predicted gene, 20379 (Fragment) OS=Mus musculus GN=Gm20379 PE=4 SV=1 |
| 154 | 1 | UniProt | n | Q61029 | 645 | 61644 | 20 | 20 | 8 | 8 | 0.73 | 1 | 0.979 | 1.031 | 0.719 | 0.677 | 0.65 | 1.003 | 0.681 | Lamina-associated polypeptide 2, isoforms beta/delta/epsilon/gamma OS=Mus musculus GN=Tmpo PE=1 SV=4 |
| 728 | 1 | UniProt | n | Q9Z2X2 | 77 | 30469 | 2 | 2 | 1 | 1 | 0.32 | 1 | 0.799 | 0.848 | 0.724 | 0.641 | 0.678 | 0.878 | 0.680 | 26S proteasome non-ATPase regulatory subunit 10 OS=Mus musculus GN=Psmd10 PE=1 SV=3 |
| 1103 | 1 | UniProt | n | Q5M8M8 | 35 | 27661 | 1 | 1 | 1 | 1 | 0.16 | 1 | 0.969 | 1.159 | 0.609 | 0.899 | 0.574 | 1.039 | 0.680 | 60S ribosomal protein L29 OS=Mus musculus GN=Rpl29 PE=1 SV=1 |
| 819 | 1 | UniProt | n | F6TMZ3 | 63 | 60880 | 2 | 2 | 1 | 1 | 0.07 | 1 | 0.86 | 0.926 | 0.565 | 0.726 | 0.764 | 0.927 | 0.679 | Epididymis-specific alpha-mannosidase (Fragment) OS=Mus musculus GN=Man2b2 PE=1 SV=1 |
| 813 | 1 | UniProt | n | Q64012 | 64 | 40303 | 2 | 2 | 2 | 2 | 0.23 | 1 | 0.864 | 0.932 | 0.656 | 0.657 | 0.727 | 0.930 | 0.679 | RNA-binding protein Raly OS=Mus musculus GN=Raly PE=1 SV=3 |
| 442 | 1 | UniProt | n | Q9DCC5 | 176 | 28846 | 7 | 7 | 3 | 3 | 0.55 | 1 | 0.846 | 0.904 | 0.74 | 0.708 | 0.597 | 0.914 | 0.679 | Cbx3 protein OS=Mus musculus GN=Cbx3 PE=1 SV=1 |
| 677 | 1 | UniProt | n | P62315 | 87 | 16620 | 3 | 3 | 2 | 2 | 0.65 | 1 | 1.044 | 1.057 | 0.682 | 0.713 | 0.643 | 1.033 | 0.679 | Small nuclear ribonucleoprotein Sm D1 OS=Mus musculus GN=Snrpd1 PE=1 SV=1 |
| 45 | 1 | UniProt | n | O88322 | 2186 | 163979 | 92 | 92 | 34 | 34 | 1.48 | 1 | 0.979 | 0.982 | 0.683 | 0.674 | 0.679 | 0.987 | 0.679 | Nidogen-2 OS=Mus musculus GN=Nid2 PE=1 SV=2 |
| 244 | 1 | UniProt | n | A0A1W2P | 357 | 20174 | 13 | 13 | 6 | 6 | 3.24 | 1 | 0.986 | 0.91 | 0.675 | 0.735 | 0.63 | 0.965 | 0.679 | Myosin light polypeptide 6 OS=Mus musculus GN=Myl6 PE=1 SV=1 |
| 206 | 1 | UniProt | n | P30416 | 430 | 64030 | 18 | 18 | 7 | 7 | 0.81 | 1 | 0.777 | 0.838 | 0.699 | 0.724 | 0.617 | 0.867 | 0.678 | Peptidyl-prolyl cis-trans isomerase FKBP4 OS=Mus musculus GN=Fkbp4 PE=1 SV=5 |
| 588 | 1 | UniProt | n | D3Z315 | 108 | 28839 | 4 | 4 | 2 | 2 | 0.34 | 1 | 0.986 | 0.992 | 0.734 | 0.677 | 0.627 | 0.993 | 0.678 | Coatomer subunit epsilon (Fragment) OS=Mus musculus GN=Cope PE=1 SV=1 |
| 418 | 1 | UniProt | n | Q91WN1 | 196 | 39304 | 5 | 5 | 4 | 4 | 0.54 | 1 | 1.086 | 1.06 | 0.71 | 0.635 | 0.69 | 1.048 | 0.678 | DnaJ homolog subfamily C member 9 OS=Mus musculus GN=Dnajc9 PE=1 SV=2 |
| 268 | 1 | UniProt | n | P45376 | 328 | 44199 | 18 | 18 | 6 | 6 | 0.95 | 1 | 1.007 | 1.006 | 0.681 | 0.664 | 0.686 | 1.004 | 0.677 | Aldose reductase OS=Mus musculus GN=Akr1b1 PE=1 SV=3 |
| 1244 | 1 | UniProt | n | A0A0N4S\ | 24 | 50865 | 1 | 1 | 1 | 1 | 0.09 | 1 | 0.281 | 0.863 | 1.32 | 0.833 | 0.281 | 0.624 | 0.676 | KICSTOR complex protein ITFG2 OS=Mus musculus GN=Itfg2 PE=1 SV=1 |
| 561 | 1 | UniProt | n | Q8BH78 | 119 | 43856 | 5 | 5 | 4 | 4 | 0.62 | 1 | 0.816 | 0.916 | 0.704 | 0.701 | 0.623 | 0.908 | 0.675 | Reticulon OS=Mus musculus GN=Rtn4 PE=1 SV=1 |
| 364 | 1 | UniProt | n | Q9DCL9 | 232 | 58221 | 8 | 8 | 4 | 4 | 0.55 | 1 | 0.941 | 0.969 | 0.732 | 0.65 | 0.646 | 0.970 | 0.675 | Multifunctional protein ADE2 OS=Mus musculus GN=Paics PE=1 SV=4 |
| 239 | 1 | UniProt | n | Q8BPB5 | 367 | 60102 | 15 | 15 | 9 | 9 | 0.88 | 1 | 0.93 | 0.936 | 0.688 | 0.654 | 0.683 | 0.955 | 0.675 | EGF-containing fibulin-like extracellular matrix protein 1 OS=Mus musculus GN=Efemp1 PE=1 SV=1 |
| 566 | 1 | UniProt | n | O08529 | 117 | 93288 | 4 | 4 | 3 | 3 | 0.15 | 1 | 0.818 | 0.861 | 0.658 | 0.716 | 0.651 | 0.890 | 0.674 | Calpain-2 catalytic subunit OS=Mus musculus GN=Capn2 PE=1 SV=4 |
| 963 | 1 | UniProt | n | Q99JT9 | 48 | 25814 | 2 | 2 | 1 | 1 | 0.18 | 1 | 0.991 | 1.018 | 0.729 | 0.633 | 0.661 | 1.003 | 0.673 | 1,2-dihydroxy-3-keto-5-methylthiopentene dioxygenase OS=Mus musculus GN=Adi1 PE=1 SV=1 |
| 271 | 1 | UniProt | n | Q3TLP8 | 325 | 30173 | 18 | 18 | 5 | 5 | 1.3 | 1 | 0.982 | 0.974 | 0.677 | 0.648 | 0.691 | 0.985 | 0.672 | RAS-related C3 botulinum substrate 1, isoform CRA a OS=Mus musculus GN=Rac1 PE=1 SV=1 |
| 825 | 1 | UniProt | n | P45377 | 62 | 45500 | 2 | 2 | 2 | 2 | 0.2 | 1 | 0.843 | 0.771 | 0.657 | 0.73 | 0.632 | 0.866 | 0.672 | Aldose reductase-related protein 2 OS=Mus musculus GN=Akr1b8 PE=1 SV=2 |
| 592 | 1 | UniProt | n | P28076 | 106 | 25299 | 4 | 4 | 3 | 3 | 0.64 | 1 | 0.937 | 0.943 | 0.658 | 0.697 | 0.66 | 0.960 | 0.671 | Proteasome subunit beta type-9 OS=Mus musculus GN=Psmb9 PE=1 SV=1 |
| 570 | 1 | UniProt | n | Q8BH69 | 116 | 49073 | 2 | 2 | 2 | 2 | 0.19 | 1 | 0.88 | 0.912 | 0.632 | 0.698 | 0.686 | 0.929 | 0.671 | Selenide, water dikinase 1 OS=Mus musculus GN=Sephs1 PE=1 SV=1 |
| 377 | 1 | UniProt | n | F7DBB3 | 225 | 202440 | 8 | 8 | 5 | 5 | 0.11 | 1 | 1.023 | 1.147 | 0.745 | 0.663 | 0.609 | 1.055 | 0.670 | AHNAK nucleoprotein 2 (Fragment) OS=Mus musculus GN=Ahnak2 PE=1 SV=1 |
| 24 | 1 | UniProt | n | P13020-2 | 3865 | 93719 | 114 | 114 | 24 | 24 | 2.89 | 1 | 1.017 | 1.006 | 0.687 | 0.652 | 0.666 | 1.008 | 0.668 | Isoform 2 of Gelsolin OS=Mus musculus GN=Gsn |
| 57 | 1 | UniProt | n | P47880 | 1585 | 27891 | 46 | 46 | 10 | 10 | 6.08 | 1 | 1.063 | 0.946 | 0.617 | 0.712 | 0.678 | 1.002 | 0.668 | Insulin-like growth factor-binding protein 6 OS=Mus musculus GN=Igfbp6 PE=2 SV=2 |
| 1018 | 1 | UniProt | n | D3YWS3 | 43 | 13870 | 1 | 1 | 1 | 1 | 0.35 | 1 | 0.855 | 0.883 | 0.803 | 0.613 | 0.605 | 0.911 | 0.668 | Profilin OS=Mus musculus GN=Pfn2 PE=1 SV=1 |
| 708 | 1 | UniProt | n | Q923F1 | 80 | 28423 | 1 | 1 | 1 | 1 | 0.16 | 1 | 0.885 | 0.829 | 0.639 | 0.704 | 0.66 | 0.902 | 0.667 | Chloride channel, nucleotide-sensitive, 1A OS=Mus musculus GN=Clns1a PE=1 SV=1 |
| 670 | 1 | UniProt | n | Q3B7Z2 | 88 | 106771 | 4 | 4 | 2 | 2 | 0.08 | 1 | 0.982 | 0.977 | 0.633 | 0.653 | 0.717 | 0.986 | 0.667 | Oxysterol-binding protein 1 OS=Mus musculus GN=Osbp PE=1 SV=3 |
| 1129 | 1 | UniProt | n | A0A0R4J0 | 33 | 117154 | 1 | 1 | 1 | 1 | 0.04 | 1 | 0.862 | 1.13 | 0.671 | 0.578 | 0.757 | 0.991 | 0.665 | Beta-mannosidase OS=Mus musculus GN=Manba PE=1 SV=1 |
| 214 | 1 | UniProt | n | O08795 | 407 | 69272 | 14 | 14 | 9 | 9 | 0.73 | 1 | 0.861 | 0.855 | 0.676 | 0.677 | 0.64 | 0.903 | 0.664 | Glucosidase 2 subunit beta OS=Mus musculus GN=Prkcsh PE=1 SV=1 |
| 485 | 1 | UniProt | n | D3Z7M9 | 153 | 32762 | 8 | 8 | 4 | 4 | 0.67 | 1 | 1.024 | 0.901 | 0.707 | 0.68 | 0.608 | 0.974 | 0.664 | Acidic leucine-rich nuclear phosphoprotein 32 family member A OS=Mus musculus GN=Anp32a PE=1 SV=1 |
| 38 | 2 | UniProt | n | P62259 | 1759 | 35073 | 63 | 63 | 16 | 16 | 11.42 | 1 | 0.899 | 0.634 | 0.598 | 0.848 | 0.574 | 0.829 | 0.663 | 14-3-3 protein epsilon OS=Mus musculus GN=Ywhae PE=1 SV=1 |
| 1104 | 1 | UniProt | n | Q8VGI4 | 35 | 37460 | 1 | 1 | 1 | 1 | 0.12 | 1 | 1.07 | 0.976 | 0.604 | 0.638 | 0.754 | 1.015 | 0.662 | Olfactory receptor 476 OS=Mus musculus GN=Olfr476 PE=3 SV=1 |
| 157 | 1 | UniProt | n | Q9Z2U1 | 622 | 30791 | 20 | 20 | 6 | 6 | 2.42 | 1 | 0.947 | 0.996 | 0.658 | 0.684 | 0.643 | 0.981 | 0.661 | Proteasome subunit alpha type-5 OS=Mus musculus GN=Psma5 PE=1 SV=1 |
| 101 | 1 | UniProt | n | A2AE89 | 1009 | 34760 | 63 | 63 | 10 | 10 | 3.83 | 1 | 0.999 | 0.935 | 0.655 | 0.687 | 0.643 | 0.978 | 0.661 | Glutathione S-transferase Mu 1 OS=Mus musculus GN=Gstm1 PE=1 SV=2 |
| 959 | 1 | UniProt | n | Q62348 | 48 | 30840 | 1 | 1 | 1 | 1 | 0.15 | 1 | 0.867 | 0.97 | 0.817 | 0.671 | 0.526 | 0.944 | 0.661 | Translin OS=Mus musculus GN=Tsn PE=1 SV=1 |
| 574 | 1 | UniProt | n | P63328 | 115 | 68588 | 6 | 6 | 4 | 4 | 0.36 | 1 | 0.971 | 0.955 | 0.656 | 0.666 | 0.657 | 0.975 | 0.660 | Serine/threonine-protein phosphatase 2B catalytic subunit alpha isoform OS=Mus musculus GN=Ppp3ca PE=1 SV=1 |
| 930 | 1 | UniProt | n | A0A0R4J0 | 51 | 69759 | 1 | 1 | 1 | 1 | 0.06 | 1 | 0.786 | 0.778 | 0.66 | 0.626 | 0.694 | 0.849 | 0.659 | Acyl-Coenzyme A binding domain containing 3, isoform CRA b OS=Mus musculus GN=Acbd3 PE=1 SV=1 |
| 415 | 1 | UniProt | n | Q9QZE5 | 198 | 115193 | 6 | 6 | 5 | 5 | 0.2 | 1 | 0.868 | 0.838 | 0.632 | 0.66 | 0.686 | 0.899 | 0.659 | Coatomer subunit gamma-1 OS=Mus musculus GN=Copg1 PE=1 SV=1 |
| 286 | 1 | UniProt | n | O08583 | 309 | 30574 | 8 | 8 | 3 | 3 | 0.51 | 1 | 0.894 | 0.877 | 0.724 | 0.64 | 0.61 | 0.922 | 0.656 | THO complex subunit 4 OS=Mus musculus GN=Alyref PE=1 SV=3 |
| 311 | 1 | UniProt | n | D3Z607 | 282 | 14804 | 8 | 8 | 2 | 2 | 2.07 | 1 | 0.952 | 0.909 | 0.644 | 0.734 | 0.597 | 0.953 | 0.656 | Non-histone chromosomal protein HMG-14 OS=Mus musculus GN=Hmgn1 PE=1 SV=1 |
| 457 | 1 | UniProt | n | Q9CR57 | 169 | 35505 | 4 | 4 | 1 | 1 | 0.13 | 1 | 0.946 | 0.872 | 0.622 | 0.738 | 0.614 | 0.938 | 0.656 | 60S ribosomal protein L14 OS=Mus musculus GN=Rpl14 PE=1 SV=3 |

| | | | | | | | | | | | | | | | | | | | |
|---|---|---|---|---|---|---|---|---|---|---|---|---|---|---|---|---|---|---|---|
| 730 | 1 | UniProt n | P03958 | 77 | 49018 | 2 | 2 | 1 | 1 | 0.09 | 1 | 0.983 | 0.916 | 0.638 | 0.636 | 0.693 | 0.966 | 0.655 | Adenosine deaminase OS=Mus musculus GN=Ada PE=1 SV=3 |
| 10 | 1 | UniProt n | P16045 | 5624 | 18174 | 244 | 244 | 10 | 10 | 95.93 | 1 | 1.071 | 0.986 | 0.656 | 0.651 | 0.658 | 1.018 | 0.655 | Galectin-1 OS=Mus musculus GN=Lgals1 PE=1 SV=3 |
| 1050 | 1 | UniProt n | P62311 | 40 | 13055 | 1 | 1 | 1 | 1 | 0.37 | 1 | 0.785 | 0.846 | 0.573 | 0.75 | 0.653 | 0.872 | 0.655 | U6 snRNA-associated Sm-like protein LSm3 OS=Mus musculus GN=Lsm3 PE=1 SV=2 |
| 340 | 1 | UniProt n | Q9CQ60 | 249 | 30159 | 14 | 14 | 6 | 6 | 1.65 | 1 | 0.944 | 0.861 | 0.681 | 0.663 | 0.62 | 0.933 | 0.654 | 6-phosphogluconolactonase OS=Mus musculus GN=Pgls PE=1 SV=1 |
| 642 | 1 | UniProt n | Q3UJB0 | 93 | 123174 | 2 | 2 | 1 | 1 | 0.04 | 1 | 0.863 | 0.899 | 0.699 | 0.65 | 0.614 | 0.919 | 0.653 | Splicing factor 3b, subunit 2 OS=Mus musculus GN=Sf3b2 PE=1 SV=1 |
| 628 | 1 | UniProt n | A0A0R4J1 | 97 | 67428 | 4 | 4 | 3 | 3 | 0.21 | 1 | 0.849 | 0.841 | 0.647 | 0.647 | 0.664 | 0.894 | 0.653 | Arylsulfatase B OS=Mus musculus GN=Arsb PE=1 SV=1 |
| 745 | 1 | UniProt n | H3BJT2 | 74 | 17049 | 3 | 3 | 1 | 1 | 0.28 | 1 | 0.955 | 1.05 | 0.678 | 0.724 | 0.565 | 1.001 | 0.652 | Insulin-like growth factor-binding protein 4 (Fragment) OS=Mus musculus GN=Igfbp4 PE=1 SV=1 |
| 486 | 1 | UniProt n | A0A1W2P | 152 | 17574 | 5 | 5 | 4 | 4 | 1.58 | 1 | 0.919 | 0.935 | 0.704 | 0.657 | 0.591 | 0.951 | 0.649 | Protein canopy homolog 2 (Fragment) OS=Mus musculus GN=Cnpy2 PE=1 SV=1 |
| 871 | 1 | UniProt n | G3UZ63 | 57 | 76669 | 1 | 1 | 1 | 1 | 0.06 | 1 | 0.871 | 0.807 | 0.705 | 0.617 | 0.626 | 0.889 | 0.648 | Kelch domain-containing protein 4 OS=Mus musculus GN=Klhdc4 PE=1 SV=1 |
| 1206 | 1 | UniProt n | Q8BWP8 | 27 | 51981 | 1 | 1 | 1 | 1 | 0.08 | 1 | 0.712 | 0.586 | 0.436 | 0.943 | 0.661 | 0.747 | 0.648 | Beta-1,4-glucuronyltransferase 1 OS=Mus musculus GN=B4gat1 PE=1 SV=1 |
| 1256 | 1 | UniProt n | A0A1D5RI | 21 | 18715 | 1 | 1 | 1 | 1 | 0.25 | 1 | 1.085 | 0.867 | 0.68 | 0.611 | 0.653 | 0.980 | 0.647 | Metallothionein-1 OS=Mus musculus GN=Mt1 PE=1 SV=1 |
| 1122 | 1 | UniProt n | Q0VF55 | 34 | 158056 | 1 | 1 | 1 | 1 | 0.03 | 1 | 0.275 | 0.468 | 0.634 | 0.934 | 0.457 | 0.505 | 0.647 | Calcium-transporting ATPase OS=Mus musculus GN=Atp2b3 PE=1 SV=1 |
| 749 | 2 | UniProt n | P54726 | 65 | 44899 | 3 | 3 | 3 | 3 | 0.33 | 1 | 0.839 | 0.888 | 0.704 | 0.612 | 0.625 | 0.907 | 0.646 | UV excision repair protein RAD23 homolog A OS=Mus musculus GN=Rad23a PE=1 SV=2 |
| 832 | 1 | UniProt n | O08797 | 62 | 50859 | 2 | 2 | 1 | 1 | 0.09 | 1 | 0.965 | 0.996 | 0.695 | 0.6 | 0.645 | 0.987 | 0.646 | SPI6 OS=Mus musculus GN=Serpinb9 PE=1 SV=1 |
| 1070 | 1 | UniProt n | Q6PDY2 | 39 | 31110 | 1 | 1 | 1 | 1 | 0.14 | 1 | 0.638 | 0.607 | 0.646 | 0.822 | 0.506 | 0.729 | 0.645 | 2-aminoethanethiol dioxygenase OS=Mus musculus GN=Ado PE=1 SV=2 |
| 995 | 1 | UniProt n | B7ZNL2 | 45 | 54739 | 2 | 2 | 2 | 2 | 0.17 | 1 | 0.809 | 0.794 | 0.614 | 0.68 | 0.641 | 0.863 | 0.644 | Nap1l4 protein OS=Mus musculus GN=Nap1l4 PE=1 SV=1 |
| 38 | 3 | UniProt n | Q9CQV8 | 1089 | 34549 | 65 | 65 | 12 | 12 | 6.03 | 1 | 0.919 | 0.666 | 0.599 | 0.777 | 0.574 | 0.849 | 0.644 | 14-3-3 protein beta/alpha OS=Mus musculus GN=Ywhab PE=1 SV=3 |
| 718 | 1 | UniProt n | G3UY29 | 78 | 19652 | 5 | 5 | 3 | 3 | 0.89 | 1 | 0.906 | 0.951 | 0.601 | 0.752 | 0.591 | 0.952 | 0.644 | MCG22989, isoform CRA a OS=Mus musculus GN=Rab11b PE=4 SV=1 |
| 1243 | 1 | UniProt n | Q5NC05 | 24 | 154755 | 1 | 1 | 1 | 1 | 0.03 | 1 | 0.845 | 0.661 | 0.602 | 0.691 | 0.641 | 0.824 | 0.644 | Transcription termination factor 2 OS=Mus musculus GN=Ttf2 PE=1 SV=2 |
| 89 | 1 | UniProt n | P08228 | 1112 | 19113 | 46 | 46 | 8 | 8 | 4.73 | 1 | 0.884 | 0.831 | 0.671 | 0.661 | 0.601 | 0.902 | 0.644 | Superoxide dismutase [Cu-Zn] OS=Mus musculus GN=Sod1 PE=1 SV=2 |
| 610 | 1 | UniProt n | Q62446 | 102 | 36130 | 6 | 6 | 5 | 5 | 0.79 | 1 | 0.999 | 0.958 | 0.62 | 0.661 | 0.648 | 0.985 | 0.643 | Peptidyl-prolyl cis-trans isomerase FKBP3 OS=Mus musculus GN=Fkbp3 PE=1 SV=2 |
| 766 | 1 | UniProt n | P09055 | 71 | 108134 | 2 | 2 | 2 | 2 | 0.08 | 1 | 0.383 | 0.516 | 0.599 | 0.735 | 0.603 | 0.582 | 0.643 | Integrin beta-1 OS=Mus musculus GN=Itgb1 PE=1 SV=1 |
| 38 | 5 | UniProt n | P68510 | 648 | 34720 | 36 | 36 | 12 | 12 | 3.83 | 1 | 0.913 | 0.647 | 0.584 | 0.801 | 0.566 | 0.839 | 0.642 | 14-3-3 protein eta OS=Mus musculus GN=Ywhah PE=1 SV=2 |
| 380 | 1 | UniProt n | Q9D154 | 224 | 52724 | 8 | 8 | 4 | 4 | 0.49 | 1 | 0.969 | 0.958 | 0.641 | 0.637 | 0.647 | 0.976 | 0.642 | Leukocyte elastase inhibitor A OS=Mus musculus GN=Serpinb1a PE=1 SV=1 |
| 38 | 6 | UniProt n | P61982 | 629 | 33899 | 44 | 44 | 13 | 13 | 5.45 | 1 | 0.921 | 0.665 | 0.594 | 0.778 | 0.569 | 0.849 | 0.641 | 14-3-3 protein gamma OS=Mus musculus GN=Ywhag PE=1 SV=2 |
| 582 | 1 | UniProt n | A0A1L1SV | 111 | 25785 | 2 | 2 | 2 | 2 | 0.38 | 1 | 0.913 | 0.838 | 0.641 | 0.643 | 0.632 | 0.915 | 0.639 | Platelet-activating factor acetylhydrolase IB subunit beta (Fragment) OS=Mus musculus GN=Pafah1b2 PE=1 SV=1 |
| 38 | 4 | UniProt n | F6VW30 | 710 | 41250 | 35 | 35 | 10 | 10 | 2.4 | 1 | 0.914 | 0.649 | 0.581 | 0.786 | 0.568 | 0.840 | 0.638 | 14-3-3 protein theta (Fragment) OS=Mus musculus GN=Ywhaq PE=1 SV=1 |
| 1260 | 1 | UniProt n | Q7TSV4 | 21 | 82245 | 1 | 1 | 1 | 1 | 0.05 | 1 | 0.809 | 0.828 | 0.656 | 0.582 | 0.677 | 0.875 | 0.637 | Phosphoglucomutase-2 OS=Mus musculus GN=Pgm2 PE=1 SV=1 |
| 673 | 1 | UniProt n | Q8BMK4 | 87 | 75564 | 2 | 2 | 1 | 1 | 0.12 | 1 | 1.006 | 0.937 | 0.658 | 0.637 | 0.614 | 0.980 | 0.636 | Cytoskeleton-associated protein 4 OS=Mus musculus GN=Ckap4 PE=1 SV=2 |
| 38 | 7 | UniProt n | O70456 | 200 | 33561 | 17 | 17 | 5 | 5 | 1.12 | 1 | 0.918 | 0.635 | 0.567 | 0.79 | 0.567 | 0.835 | 0.633 | 14-3-3 protein sigma OS=Mus musculus GN=Sfn PE=1 SV=2 |
| 163 | 1 | UniProt n | P99027 | 590 | 14990 | 14 | 14 | 5 | 5 | 4.25 | 1 | 1.003 | 0.893 | 0.653 | 0.686 | 0.564 | 0.964 | 0.632 | 60S acidic ribosomal protein P2 OS=Mus musculus GN=Rplp2 PE=1 SV=3 |
| 321 | 1 | UniProt n | P48036 | 269 | 42469 | 8 | 8 | 7 | 7 | 1 | 1 | 0.769 | 0.61 | 0.558 | 0.829 | 0.545 | 0.777 | 0.632 | Annexin A5 OS=Mus musculus GN=Anxa5 PE=1 SV=1 |
| 38 | 1 | UniProt n | P63101 | 2557 | 34280 | 101 | 101 | 15 | 15 | 12.14 | 1 | 0.914 | 0.645 | 0.588 | 0.772 | 0.554 | 0.838 | 0.631 | 14-3-3 protein zeta/delta OS=Mus musculus GN=Ywhaz PE=1 SV=1 |
| 379 | 1 | UniProt n | G3UXZ5 | 224 | 36354 | 13 | 13 | 7 | 7 | 1.25 | 1 | 1.015 | 1.032 | 0.632 | 0.641 | 0.618 | 1.016 | 0.630 | Proteasome activator complex subunit 1 (Fragment) OS=Mus musculus GN=Psme1 PE=1 SV=1 |
| 101 | 2 | UniProt n | P15626 | 488 | 32530 | 25 | 25 | 8 | 8 | 2.64 | 1 | 0.935 | 0.778 | 0.616 | 0.717 | 0.564 | 0.899 | 0.629 | Glutathione S-transferase Mu 2 OS=Mus musculus GN=Gstm2 PE=1 SV=2 |
| 1147 | 1 | UniProt n | Q8BK67 | 32 | 65976 | 2 | 2 | 1 | 1 | 0.07 | 1 | 0.84 | 0.908 | 0.674 | 0.582 | 0.629 | 0.914 | 0.627 | Protein RCC2 OS=Mus musculus GN=Rcc2 PE=1 SV=1 |
| 879 | 1 | UniProt n | A0JNY3 | 57 | 95420 | 1 | 1 | 1 | 1 | 0.05 | 1 | 0.83 | 0.865 | 0.625 | 0.585 | 0.673 | 0.895 | 0.627 | Gephyrin OS=Mus musculus GN=Gphn PE=1 SV=1 |
| 778 | 1 | UniProt n | P08207 | 69 | 14617 | 5 | 5 | 2 | 2 | 0.76 | 1 | 1.112 | 1.016 | 0.606 | 0.633 | 0.639 | 1.042 | 0.626 | Protein S100-A10 OS=Mus musculus GN=S100a10 PE=1 SV=2 |
| 312 | 1 | UniProt n | O88792 | 279 | 37805 | 8 | 8 | 4 | 4 | 0.56 | 1 | 1.068 | 1.061 | 0.614 | 0.66 | 0.602 | 1.043 | 0.625 | Junctional adhesion molecule A OS=Mus musculus GN=F11r PE=1 SV=2 |
| 301 | 2 | UniProt n | Q9CQ19 | 165 | 24451 | 8 | 8 | 4 | 4 | 0.99 | 1 | 1.054 | 0.961 | 0.57 | 0.674 | 0.629 | 1.004 | 0.623 | Myosin regulatory light polypeptide 9 OS=Mus musculus GN=Myl9 PE=1 SV=3 |
| 683 | 1 | UniProt n | E9QJT5 | 86 | 20817 | 5 | 5 | 2 | 2 | 0.49 | 1 | 0.806 | 0.728 | 0.615 | 0.69 | 0.567 | 0.837 | 0.622 | Acylphosphatase OS=Mus musculus GN=Acyp1 PE=1 SV=1 |
| 207 | 2 | UniProt n | A0A0U1RI | 92 | 21037 | 4 | 4 | 3 | 3 | 0.81 | 1 | 0.926 | 0.798 | 0.587 | 0.618 | 0.658 | 0.904 | 0.620 | Isocitrate dehydrogenase [NADP], mitochondrial (Fragment) OS=Mus musculus GN=Idh2 PE=1 SV=1 |
| 753 | 1 | UniProt n | D3YVH4 | 73 | 36983 | 1 | 1 | 1 | 1 | 0.12 | 1 | 0.915 | 0.841 | 0.623 | 0.644 | 0.593 | 0.916 | 0.620 | Splicing factor 1 (Fragment) OS=Mus musculus GN=Sf1 PE=1 SV=1 |
| 129 | 1 | UniProt n | Q62000 | 785 | 42784 | 55 | 55 | 14 | 14 | 4.35 | 1 | 1.107 | 1.078 | 0.624 | 0.616 | 0.61 | 1.061 | 0.617 | Mimecan OS=Mus musculus GN=Ogn PE=1 SV=1 |
| 461 | 1 | UniProt n | A0A1L1SU | 167 | 30832 | 5 | 5 | 3 | 3 | 0.5 | 1 | 0.706 | 0.786 | 0.637 | 0.604 | 0.603 | 0.822 | 0.614 | Twinfilin-2 (Fragment) OS=Mus musculus GN=Twf2 PE=1 SV=1 |
| 1014 | 1 | UniProt n | A0A087W | 44 | 190975 | 1 | 1 | 1 | 1 | 0.02 | 1 | 0.844 | 0.825 | 0.549 | 0.596 | 0.709 | 0.886 | 0.614 | Kinectin OS=Mus musculus GN=Ktn1 PE=1 SV=1 |
| 712 | 1 | UniProt n | P63325 | 80 | 23163 | 1 | 1 | 1 | 1 | 0.2 | 1 | 0.89 | 0.963 | 0.567 | 0.608 | 0.671 | 0.950 | 0.614 | 40S ribosomal protein S10 OS=Mus musculus GN=Rps10 PE=1 SV=1 |
| 117 | 1 | UniProt n | O35887 | 855 | 46259 | 27 | 27 | 10 | 10 | 1.99 | 1 | 1.047 | 1.089 | 0.636 | 0.626 | 0.579 | 1.045 | 0.613 | Calumenin OS=Mus musculus GN=Calu PE=1 SV=1 |
| 463 | 1 | UniProt n | Q61468 | 164 | 80062 | 8 | 8 | 3 | 3 | 0.17 | 1 | 0.876 | 0.837 | 0.658 | 0.599 | 0.582 | 0.902 | 0.612 | Mesothelin OS=Mus musculus GN=Msln PE=1 SV=1 |
| 117 | 2 | UniProt n | Q6XLQ8 | 807 | 45704 | 27 | 27 | 9 | 9 | 1.76 | 1 | 1.033 | 1.088 | 0.653 | 0.617 | 0.567 | 1.040 | 0.611 | Calumenin OS=Mus musculus GN=Calu PE=1 SV=1 |
| 1262 | 1 | UniProt n | A0A0J9YU | 20 | 109368 | 1 | 1 | 1 | 1 | 0.04 | 1 | 0.703 | 0.7 | 0.695 | 0.601 | 0.544 | 0.789 | 0.610 | Dynamin-1 OS=Mus musculus GN=Dnm1 PE=1 SV=1 |
| 523 | 1 | UniProt n | B1AQR8 | 131 | 44767 | 4 | 4 | 3 | 3 | 0.33 | 1 | 0.943 | 0.972 | 0.556 | 0.584 | 0.697 | 0.971 | 0.609 | Galectin OS=Mus musculus GN=Lgals9 PE=1 SV=1 |
| 826 | 1 | UniProt n | O08599 | 62 | 83971 | 1 | 1 | 1 | 1 | 0.05 | 1 | 0.92 | 0.875 | 0.674 | 0.617 | 0.54 | 0.930 | 0.608 | Syntaxin-binding protein 1 OS=Mus musculus GN=Stxbp1 PE=1 SV=2 |
| 398 | 1 | UniProt n | P51855 | 208 | 59699 | 8 | 8 | 5 | 5 | 0.53 | 1 | 0.674 | 0.746 | 0.623 | 0.573 | 0.623 | 0.795 | 0.606 | Glutathione synthetase OS=Mus musculus GN=Gss PE=1 SV=1 |
| 1227 | 1 | UniProt n | A0A0R4IZ' | 26 | 32130 | 1 | 1 | 1 | 1 | 0.14 | 1 | 0.913 | 0.964 | 0.651 | 0.621 | 0.55 | 0.958 | 0.606 | Calpain small subunit 1 OS=Mus musculus GN=Capns1 PE=1 SV=1 |
| 919 | 1 | UniProt n | Q8K0Q5 | 53 | 93016 | 2 | 2 | 2 | 2 | 0.1 | 1 | 0.914 | 1.024 | 0.612 | 0.596 | 0.606 | 0.978 | 0.605 | Rho GTPase-activating protein 18 OS=Mus musculus GN=Arhgap18 PE=1 SV=1 |
| 1125 | 1 | UniProt n | B8JJG6 | 34 | 19346 | 1 | 1 | 1 | 1 | 0.24 | 1 | 0.824 | 0.735 | 0.629 | 0.666 | 0.52 | 0.846 | 0.602 | Trk-fused OS=Mus musculus GN=Tfg PE=1 SV=1 |
| 1095 | 1 | UniProt n | B8JJ89 | 36 | 36447 | 1 | 1 | 1 | 1 | 0.12 | 1 | 0.737 | 0.814 | 0.634 | 0.559 | 0.608 | 0.843 | 0.600 | Apoptotic chromatin condensation inducer in the nucleus (Fragment) OS=Mus musculus GN=Acin1 PE=1 SV=8 |
| 585 | 1 | UniProt n | Q501P1 | 111 | 54097 | 4 | 4 | 3 | 3 | 0.26 | 1 | 0.925 | 1.022 | 0.647 | 0.523 | 0.634 | 0.981 | 0.599 | Fibulin-7 OS=Mus musculus GN=Fbln7 PE=1 SV=1 |
| 816 | 1 | UniProt n | P0DOV1 | 63 | 59880 | 2 | 2 | 2 | 2 | 0.15 | 1 | 0.841 | 1.013 | 0.622 | 0.595 | 0.577 | 0.948 | 0.598 | Interferon-activable protein 205-B OS=Mus musculus GN=Mnda PE=1 SV=1 |
| 334 | 1 | UniProt n | A0A087W | 260 | 8617 | 14 | 14 | 1 | 1 | 1.56 | 1 | 1.136 | 0.943 | 0.564 | 0.675 | 0.559 | 1.023 | 0.597 | Prothymosin alpha OS=Mus musculus GN=Ptma PE=1 SV=1 |
| 1025 | 1 | UniProt n | A2A6U3 | 43 | 76481 | 2 | 2 | 1 | 1 | 0.06 | 1 | 0.956 | 0.951 | 0.604 | 0.469 | 0.746 | 0.969 | 0.596 | Septin-9 OS=Mus musculus GN=Sept9 PE=1 SV=1 |
| 1221 | 1 | UniProt n | Q8CHY3 | 26 | 84778 | 1 | 1 | 1 | 1 | 0.05 | 1 | 1.126 | 1.439 | 0.334 | 0.721 | 0.877 | 1.175 | 0.596 | Dymeclin OS=Mus musculus GN=Dym PE=1 SV=1 |
| 847 | 1 | UniProt n | Q99NF7 | 60 | 59276 | 1 | 1 | 1 | 1 | 0.07 | 1 | 0.976 | 1.217 | 0.526 | 0.594 | 0.673 | 1.059 | 0.595 | Ppm1b protein OS=Mus musculus GN=Ppm1b PE=1 SV=1 |
| 351 | 1 | UniProt n | Q61576 | 242 | 71154 | 13 | 13 | 9 | 9 | 0.81 | 1 | 1.004 | 0.983 | 0.591 | 0.604 | 0.586 | 0.996 | 0.594 | Peptidyl-prolyl cis-trans isomerase FKBP10 OS=Mus musculus GN=Fkbp10 PE=1 SV=2 |
| 838 | 1 | UniProt n | G5E8W1 | 61 | 46830 | 1 | 1 | 1 | 1 | 0.09 | 1 | 0.869 | 0.874 | 0.629 | 0.586 | 0.567 | 0.912 | 0.593 | TRIO and F-actin-binding protein OS=Mus musculus GN=Triobp PE=1 SV=1 |
| 101 | 3 | UniProt n | Q8R5I6 | 115 | 30508 | 7 | 7 | 4 | 4 | 0.99 | 1 | 0.87 | 0.714 | 0.565 | 0.658 | 0.561 | 0.853 | 0.593 | Glutathione S-transferase mu 4 OS=Mus musculus GN=Gstm4 PE=1 SV=1 |
| 1232 | 1 | UniProt n | Q8BI74 | 25 | 28592 | 1 | 1 | 1 | 1 | 0.16 | 1 | 0.886 | 0.924 | 0.637 | 0.565 | 0.567 | 0.935 | 0.589 | VPS10 domain-containing receptor SorCS2 OS=Mus musculus GN=Sorcs2 PE=1 SV=1 |
| 654 | 1 | UniProt n | Q9CR26 | 90 | 39902 | 2 | 2 | 1 | 1 | 0.24 | 1 | 0.565 | 0.625 | 0.539 | 0.771 | 0.489 | 0.707 | 0.588 | Vacuolar protein sorting-associated protein VTA1 homolog OS=Mus musculus GN=Vta1 PE=1 SV=1 |
| 892 | 1 | UniProt n | Q9Z1D1 | 55 | 44972 | 2 | 2 | 2 | 2 | 0.21 | 1 | 0.923 | 0.936 | 0.614 | 0.478 | 0.69 | 0.952 | 0.587 | Eukaryotic translation initiation factor 3 subunit G OS=Mus musculus GN=Eif3g PE=1 SV=2 |
| 925 | 1 | UniProt n | O35955 | 53 | 31709 | 2 | 2 | 2 | 2 | 0.3 | 1 | 0.999 | 0.875 | 0.58 | 0.612 | 0.55 | 0.956 | 0.580 | Proteasome subunit beta type-10 OS=Mus musculus GN=Psmb10 PE=1 SV=1 |
| 385 | 1 | UniProt n | G3UYV7 | 219 | 7299 | 8 | 8 | 3 | 3 | 4.21 | 1 | 0.908 | 0.822 | 0.587 | 0.633 | 0.519 | 0.907 | 0.578 | 40S ribosomal protein S28 (Fragment) OS=Mus musculus GN=Rps28 PE=1 SV=1 |
| 1126 | 1 | UniProt n | A0A1L1SR | 34 | 144311 | 1 | 1 | 1 | 1 | 0.03 | 1 | 0.983 | 0.862 | 0.701 | 0.491 | 0.552 | 0.946 | 0.575 | WD repeat-containing protein 72 OS=Mus musculus GN=Wdr72 PE=1 SV=1 |
| 393 | 1 | UniProt n | Q91XV3 | 210 | 30896 | 7 | 7 | 5 | 5 | 0.98 | 1 | 0.848 | 0.801 | 0.614 | 0.567 | 0.532 | 0.879 | 0.570 | Brain acid soluble protein 1 OS=Mus musculus GN=Basp1 PE=1 SV=3 |
| 1057 | 1 | UniProt n | Q8VCT3 | 40 | 82657 | 1 | 1 | 1 | 1 | 0.05 | 1 | 0.948 | 1.077 | 0.519 | 0.622 | 0.567 | 1.007 | 0.568 | Aminopeptidase B OS=Mus musculus GN=Rnpep PE=1 SV=2 |
| 1008 | 1 | UniProt n | Q64339 | 44 | 21675 | 3 | 3 | 2 | 2 | 0.78 | 1 | 1.081 | 0.981 | 0.494 | 0.668 | 0.552 | 1.020 | 0.567 | Ubiquitin-like protein ISG15 OS=Mus musculus GN=Isg15 PE=1 SV=4 |
| 1240 | 1 | UniProt n | E9Q4T8 | 24 | 98297 | 1 | 1 | 1 | 1 | 0.04 | 1 | 0.817 | 0.891 | 0.538 | 0.607 | 0.55 | 0.900 | 0.564 | Cullin-3 OS=Mus musculus GN=Cul3 PE=1 SV=1 |
| 1196 | 1 | UniProt n | A0A1B0G( | 29 | 14078 | 1 | 1 | 1 | 1 | 0.34 | 1 | 1.054 | 1.103 | 0.501 | 0.608 | 0.587 | 1.051 | 0.563 | U6 snRNA-associated Sm-like protein LSm4 OS=Mus musculus GN=Lsm4 PE=1 SV=1 |
| 1019 | 1 | UniProt n | Q8R1G6 | 43 | 41136 | 2 | 2 | 1 | 1 | 0.11 | 1 | 0.905 | 0.921 | 0.562 | 0.548 | 0.575 | 0.941 | 0.562 | PDZ and LIM domain protein 2 OS=Mus musculus GN=Pdlim2 PE=1 SV=1 |
| 1144 | 1 | UniProt n | A0A0N4S\ | 32 | 22794 | 2 | 2 | 1 | 1 | 0.2 | 1 | 0.843 | 0.875 | 0.567 | 0.572 | 0.545 | 0.904 | 0.561 | Secernin-1 (Fragment) OS=Mus musculus GN=Scrn1 PE=1 SV=1 |
| 983 | 1 | UniProt n | A0A0G2JF | 47 | 12107 | 1 | 1 | 1 | 1 | 0.41 | 1 | 1.473 | 0.917 | 0.511 | 0.549 | 0.624 | 1.105 | 0.559 | Cysteine-rich protein 2 OS=Mus musculus GN=Crip2 PE=1 SV=1 |
| 83 | 3 | UniProt n | A2AL12 | 202 | 41331 | 8 | 8 | 4 | 4 | 0.5 | 1 | 0.72 | 0.717 | 0.555 | 0.549 | 0.565 | 0.802 | 0.556 | Heterogeneous nuclear ribonucleoprotein A3 OS=Mus musculus GN=Hnrnpa3 PE=1 SV=1 |
| 962 | 1 | UniProt n | E9Q514 | 48 | 18141 | 1 | 1 | 1 | 1 | 0.26 | 1 | 0.664 | 0.759 | 0.523 | 0.614 | 0.53 | 0.796 | 0.554 | Protein NDRG1 (Fragment) OS=Mus musculus GN=Ndrg1 PE=1 SV=1 |
| 301 | 1 | UniProt n | Q3THE2 | 292 | 24680 | 11 | 11 | 5 | 5 | 1.34 | 1 | 1.06 | 0.934 | 0.517 | 0.606 | 0.542 | 0.997 | 0.554 | Myosin regulatory light chain 12B OS=Mus musculus GN=Myl12b PE=1 SV=2 |
| 998 | 1 | UniProt n | O88653 | 45 | 16632 | 2 | 2 | 2 | 2 | 0.65 | 1 | 1.024 | 0.956 | 0.499 | 0.461 | 0.732 | 0.993 | 0.552 | Ragulator complex protein LAMTOR3 OS=Mus musculus GN=Lamtor3 PE=1 SV=1 |
| 203 | 1 | UniProt n | Q99K85 | 444 | 48282 | 29 | 29 | 12 | 12 | 2.12 | 1 | 0.98 | 0.974 | 0.595 | 0.512 | 0.534 | 0.985 | 0.546 | Phosphoserine aminotransferase OS=Mus musculus GN=Psat1 PE=1 SV=1 |
| 982 | 1 | UniProt n | A0A140LII | 47 | 6171 | 1 | 1 | 1 | 1 | 0.9 | 1 | 0.884 | 0.787 | 0.503 | 0.642 | 0.496 | 0.886 | 0.543 | Cytochrome c oxidase subunit 6B1 OS=Mus musculus GN=Cox6b1 PE=1 SV=1 |
| 102 | 1 | UniProt n | A0A0N4S\ | 1008 | 17302 | 13 | 13 | 2 | 2 | 1.61 | 1 | 1.16 | 1.032 | 0.5 | 0.555 | 0.57 | 1.062 | 0.541 | Histone H2A OS=Mus musculus GN=H2afj PE=1 SV=1 |
| 981 | 1 | UniProt n | A0A0G2JD | 47 | 17194 | 1 | 1 | 1 | 1 | 0.27 | 1 | 0.85 | 0.925 | 0.591 | 0.422 | 0.634 | 0.923 | 0.541 | Mevalonate kinase (Fragment) OS=Mus musculus GN=Mvk PE=1 SV=1 |
| 101 | 4 | UniProt n | E9PVM7 | 75 | 30585 | 5 | 5 | 2 | 2 | 0.32 | 1 | 0.857 | 0.683 | 0.519 | 0.575 | 0.504 | 0.837 | 0.532 | Glutathione S-transferase (Fragment) OS=Mus musculus GN=Gstm5 PE=1 SV=1 |
| 102 | 2 | UniProt n | Q64523 | 855 | 18543 | 11 | 11 | 2 | 2 | 0.96 | 1 | 1.165 | 1.032 | 0.478 | 0.541 | 0.57 | 1.063 | 0.528 | Histone H2A type 2-C OS=Mus musculus GN=Hist2h2ac PE=1 SV=3 |
| 899 | 1 | UniProt n | Q3UHD3 | 55 | 175196 | 4 | 4 | 2 | 2 | 0.05 | 1 | 1.25 | 1.183 | 0.497 | 0.608 | 0.483 | 1.139 | 0.527 | Microtubule-associated tumor suppressor candidate 2 homolog OS=Mus musculus GN=Mtus2 PE=1 SV=1 |
| 1085 | 1 | UniProt n | Q9CSH3 | 37 | 129933 | 1 | 1 | 1 | 1 | 0.03 | 1 | 0.941 | 1.016 | 0.399 | 0.532 | 0.655 | 0.985 | 0.518 | Exosome complex exonuclease RRP44 OS=Mus musculus GN=Dis3 PE=1 SV=4 |
| 696 | 1 | UniProt n | Q9DBR7 | 83 | 141457 | 2 | 2 | 2 | 2 | 0.06 | 1 | 0.752 | 0.748 | 0.574 | 0.428 | 0.548 | 0.825 | 0.513 | Protein phosphatase 1 regulatory subunit 12A OS=Mus musculus GN=Ppp1r12a PE=1 SV=2 |
| 852 | 1 | UniProt n | P51885 | 60 | 47080 | 3 | 3 | 2 | 2 | 0.2 | 1 | 0.928 | 0.947 | 0.439 | 0.52 | 0.587 | 0.958 | 0.512 | Lumican OS=Mus musculus GN=Lum PE=1 SV=2 |
| 947 | 1 | UniProt n | Q8BQ47 | 50 | 33569 | 1 | 1 | 1 | 1 | 0.13 | 1 | 0.91 | 0.795 | 0.459 | 0.545 | 0.527 | 0.898 | 0.509 | Protein canopy homolog 4 OS=Mus musculus GN=Cnpy4 PE=1 SV=1 |
| 663 | 1 | UniProt n | Q9CPY7 | 89 | 67075 | 5 | 5 | 4 | 4 | 0.29 | 1 | 0.972 | 0.89 | 0.474 | 0.544 | 0.506 | 0.953 | 0.507 | Cytosol aminopeptidase OS=Mus musculus GN=Lap3 PE=1 SV=3 |
| 1143 | 1 | UniProt n | Q6PDH4 | 32 | 64440 | 2 | 2 | 2 | 2 | 0.14 | 1 | 0.678 | 0.696 | 0.519 | 0.511 | 0.46 | 0.779 | 0.496 | Ran-binding protein 3-like OS=Mus musculus GN=Ranbp3l PE=1 SV=1 |
| 130 | 1 | UniProt n | Q6ZWY9 | 780 | 20286 | 32 | 32 | 8 | 8 | 8.58 | 1 | 1.304 | 1.089 | 0.456 | 0.482 | 0.528 | 1.124 | 0.488 | Histone H2B type 1-C/E/G OS=Mus musculus GN=Hist1h2bc PE=1 SV=3 |
| 189 | 1 | UniProt n | P62806 | 482 | 15011 | 22 | 22 | 6 | 6 | 5.93 | 1 | 1.235 | 0.819 | 0.438 | 0.494 | 0.513 | 1.004 | 0.481 | Histone H4 OS=Mus musculus GN=Hist1h4a PE=1 SV=2 |
| 290 | 1 | UniProt n | P19324 | 303 | 57548 | 14 | 14 | 11 | 11 | 1.25 | 1 | 1.051 | 1.052 | 0.474 | 0.499 | 0.46 | 1.034 | 0.477 | Serpin H1 OS=Mus musculus GN=Serpinh1 PE=1 SV=3 |
| 1088 | 1 | UniProt n | D3Z7F7 | 37 | 31429 | 1 | 1 | 1 | 1 | 0.14 | 1 | 0.732 | 0.835 | 0.532 | 0.438 | 0.436 | 0.849 | 0.467 | Serine/threonine-protein phosphatase CPPED1 (Fragment) OS=Mus musculus GN=Cpped1 PE=1 SV=1 |
| 516 | 1 | UniProt n | P97857 | 135 | 125934 | 3 | 3 | 2 | 2 | 0.11 | 1 | 0.992 | 0.902 | 0.462 | 0.452 | 0.466 | 0.964 | 0.460 | A disintegrin and metalloproteinase with thrombospondin motifs 1 OS=Mus musculus GN=Adamts1 PE=1 SV=4 |
| 861 | 1 | UniProt n | A0A1W2P | 59 | 25193 | 5 | 5 | 2 | 2 | 0.39 | 1 | 1.379 | 1.086 | 0.434 | 0.379 | 0.474 | 1.144 | 0.427 | Histone H3.2 OS=Mus musculus GN=Hist2h3c1 PE=1 SV=1 |
| 1228 | 1 | UniProt n | O89023 | 26 | 65931 | 1 | 1 | 1 | 1 | 0.07 | 1 | 0.972 | 0.806 | 0.277 | 0.426 | 0.61 | 0.922 | 0.416 | Tripeptidyl-peptidase 1 OS=Mus musculus GN=Tpp1 PE=1 SV=2 |
| 216 | 1 | UniProt n | Q9QZZ6 | 404 | 26264 | 21 | 21 | 6 | 6 | 3.2 | 1 | 1.016 | 0.986 | 0.421 | 0.402 | 0.4 | 1.001 | 0.408 | Dermatopontin OS=Mus musculus GN=Dpt PE=1 SV=1 |
| 924 | 1 | UniProt n | P17095 | 53 | 16779 | 1 | 1 | 1 | 1 | 0.28 | 1 | 1.011 | 1.098 | 0.307 | 0.332 | 0.627 | 1.035 | 0.400 | High mobility group protein HMG-I/HMG-Y OS=Mus musculus GN=Hmga1 PE=1 SV=4 |
| 223 | 1 | UniProt n | A2BE93 | 390 | 31905 | 13 | 13 | 5 | 5 | 0.93 | 1 | 1.098 | 1.102 | 0.402 | 0.41 | 0.385 | 1.066 | 0.399 | Protein SET (Fragment) OS=Mus musculus GN=Set PE=1 SV=1 |
| 611 | 1 | UniProt n | P51910 | 102 | 25654 | 2 | 2 | 2 | 2 | 0.39 | 1 | 0.894 | 0.925 | 0.385 | 0.393 | 0.348 | 0.939 | 0.375 | Apolipoprotein D OS=Mus musculus GN=Apod PE=1 SV=1 |
| 1141 | 1 | UniProt n | Q05CL8 | 32 | 92204 | 1 | 1 | 1 | 1 | 0.05 | 1 | 0.594 | 0.553 | 0.254 | 0.426 | 0.42 | 0.690 | 0.357 | La-related protein 7 OS=Mus musculus GN=Larp7 PE=1 SV=2 |
| 562 | 1 | UniProt n | D3Z7M4 | 118 | 18734 | 3 | 3 | 2 | 2 | 0.56 | 1 | 0.816 | 0.759 | 0.305 | 0.326 | 0.321 | 0.852 | 0.317 | Insulin-like growth factor I (Fragment) OS=Mus musculus GN=Igf1 PE=1 SV=1 |
| 1142 | 1 | UniProt n | Q7M6Y3 | 27 | 84109 | 2 | 2 | 2 | 2 | 0.11 | 1 | 0.421 | 0.429 | 0.288 | 0.346 | 0.284 | 0.565 | 0.305 | Phosphatidylinositol-binding clathrin assembly protein OS=Mus musculus GN=Picalm PE=1 SV=1 |
| 1251 | 1 | UniProt n | D3YYA0 | 23 | 69566 | 1 | 1 | 1 | 1 | 0.06 | 1 | 0.382 | 0.29 | 0.352 | 0.215 | 0.29 | 0.480 | 0.280 | Set1/Ash2 histone methyltransferase complex subunit ASH2 OS=Mus musculus GN=Ash2l PE=1 SV=1 |
| 1220 | 1 | UniProt_n | P97333 | 26 | 121286 | 1 | 1 | 1 | 1 | 0.04 | 1 | 0.173 | 0.166 | 0.158 | 0.183 | 0.212 | 0.306 | 0.183 | Neuropilin-1 OS=Mus musculus GN=Nrp1 PE=1 SV=2 |

#### Supplementary Table 2. Reagent List

| Reagent | Source | Catalogue Number/Identifier |
|---|---|---|
| **Antibodies** | | |
| Tenascin C | Millipore Sigma | AB19011; RRID:AB_2063439 |
| Phospho-Smad1/5(Ser463/465) | Cell Signaling | 9516; RRID:AB_491015 |
| Smad1 | Cell Signaling | 9743; RRID:AB_2107780 |
| Tenascin C antibody (BC-24) | Millipore Sigma | SAB4200782 |
| Cytokeratin antibody AE1/AE3 | Dako Omnis | M3515; RRID:AB_2132885) |
| Integrin α5β1 antibody M200 (Volociximab) | Novus Biologicals | NBP2-52680 |
| ChromPure Rabbit IgG | Jackson Immuno Research Laboratories | 011-000-003; RRID:AB_2337118 |
| GAPDH | Cell Signaling | 97166; RRID:AB_2756824 |
| Vinculin(E1E9V) | Cell Signaling | 13901; RRID:AB_2728768 |
| Hey1 (WB) | Abcam | Ab154077 |
| Hey1 (IP) | Proteintech | 19929-1-AP; RRID:AB_10646438 |
| YAP-1 | Cell Signaling | 4922; RRID:AB_2218911 |
| Androgen receptor | Abcam | Ab133273; RRID:AB_11156085 |
| FAK | Cell Signaling | 3285; RRID:AB_2269034 |
| pY397-FAK | Thermo Fisher Scientific | 44-624G; RRID:AB_2533701 |
| Src | Cell Technology | 2110; RRID:AB_10691385 |
| pY416-Src | Cell Signaling | 2101; RRID:AB_331697 |
| **Reagents** | | |
| LDN193189 | Axon Medchem | Axon 1509 |
| Recombinant human BMP4 | R&D Systems | 314-BP |
| Recombinant human Tenascin C | R&D Systems | 3358-TC |
| LightSwitchTM promoter Vector | Active Motif | 32021 ProductID: S790005 |
| LightSwitch$^{TM}$ promoter GoClone | Active Motif | 32001 ProductID: S722838 |
| LightSwitch Luciferase Assay Kit | Active Motif | 32031 ProductID: LS010 |
| Mouse TNC Elisa Kit | MyBioSource | MBS9627232 |
| alkaline phosphatase kit | Thermo Fisher Scientific | 37620 |
| SYBR Green | Thermo Fisher Scientific | 4309155 |
| Lipofectamin 2000 | Thermo Fisher Scientific | 11668-019 |

| | | |
|---|---|---|
| RNeasy mini Kit | Qiagen | 74106 |
| DNeasy kit | Qiagen | 69506 |
| PNPP Substrate kit | Thermo Fisher Scientific | 37620 |
| Human Tenascin C Elisa kit | Ray Biotech | ELH-TNC |
| Dual-luciferase Reporter assay kit | Promega | E1960 |
| 8xGTIIC (8xTEAD TIK Luc) | Addgene | RRID:Addgene_115798 |
| **Cell line** | **Source** | **Catalogue Number/Identifier** |
| LNCaP C4-2B | ATCC | CRL-3315; RRID:CVCL_4784 |
| LNCaP | CLS | 300265/p761_LNCaP; RRID:CVCL_0395 |
| PC3 | NCI-DTP | PC-3; RRID: CVCL_0035 |
| PC3-MM2 | KCLB | 80021; RRID:CVCL_4885 |
| DU145 | CLS | 300168/p708_DU-145; RRID:CVCL_0105 |
| 22RV1 | BCRC | 60545; RRID:CVCL_1045 |
| 2H11 | ATCC | CRL-2163; RRID:CVCL_6762 |
| 293T | ATCC | CRL-321; RRID:CVCL_LF41 |

#### Supplementary Table 3. Oligonucleotide sequences of primers.

| primers | Sequence (5'-3') |
|---|---|
| mALP-F | CTCCTCCATCCCTTCCCTTC |
| mALP-R | CCCTGGGTAGACAGCCAAC |
| GAPDH-F | CCCAGAAGACTGTGGATG |
| GAPDH-R | GCAGGGATGATGTTCTGG |
| mOC-F | GCTCTGTCTCTCTGACCTCA |
| mOC-R | TGGACATGAAGGCTTTGTCA |
| hTNC-F1 | CGGTGGTGGATTCTGGGAAG |
| hTNC-R1 | TCCGGTTCGGCTTCTGTAAC |
| hTNC-F2 | GCCATGACTCAGCTGTTG |
| hTNC-R2 | CACTGGCAGCTTGATGTTGT |
| mHey1-F | AAGACGGAGAGGCATCATCGAG |
| mHey1-R | CAGATCCCTGCTTCTCAAAGGCAC |
| mVcan-F | CCCAGTGTGGAGGAGGTCTA |
| mVcan-R | CACTCAATCGACGTTTAAAGCA |
| mCTGF-F | AGAACTGTGTACGGAGCGTG |
| mCTGF-R | GTGCACCATCTTTGGCAGTG |
| mTNC-F | CTCTGGAATTGCTCCCAGCAT |
| mTNC-R | TTCCGGTTCAGCTTCTGTGGTAG |

| | |
|---|---|
| h-Alu-F | CATGGTGAAACCCCGTCTCTA |
| h-Alu-R | GCCTCAGCCTCCCGAGTAG |
| hCOL8A1-F | CAGAAACCAGCCCCAGAGGTGTCAC |
| hCOL8A1-R | GAAATG GTAAGCAGCACTCCCAGCAG |
| hCTGF-F | GCAGAGCCGC CTGTGCATGG |
| hCTGF-R | GGTATGTCTTCATGCTGG |
| hCYR61-F | CACACCAAGGGGCTGGAATG |
| hCYR61-R | CCCGTTTTGGTAGATTCTGG |

| NameSequence (5'-3') | | Product length (bp) |
|---|---|---|
| **Sequences for cDNA overexpression clones** | | |
| hTNC-ECoR1-F | GAATTC ATGGGGGCCATGACTCAGCTGTTG | 1878 |
| hTNC-NotI-R | GCGGCCGC CTAGGGAGGAGACACCTCTGAGCA | |
| Hey1-BamH1-F | GGATCCATGAAGAGAGCTCACCCAGAC | 901 |
| Hey1-EcoRI-R | GAATTCTTTAGAAAGCTCCGATCTCTGT | |

**Sequences for shRNA knockdown clones**

| Gene | Accession | Clone ID | Mature Antisense (5'-3') | Target |
|---|---|---|---|---|
| shTNC | NM_002160 | V2LHS_133230 (#2) | ATTCCGGAAGATGATCTC | ORF |
| | NM_002160 | V3LHS_321947 (#5) | TGCTTGACATGTCACCTCT | ORF |

| | | | | |
|---|---|---|---|---|
| | NM_002160 | V3LHS_321952 (#8) | TGACATGTCACCTCTGCGG | ORF |
| shYAP1 | NM_006106 | TRCN0000107265 (#265) | CCGGCCCAGTTAAATGTTCACCAATCTC GAGATTGGTGAACATTTAACTGGGTTTTTG | 3'UTR |
| | NM_006106 | TRCN0000300325 (#325) | CCGGGCCACCAAGCTAGATAAAGAACTCG AGTTCTTTATCTAGCTTGGTGGCTTTTTG | CDS |
| shHey1 | NM_010423 | V3LMM_516064 (#2) | GCATTAAAGCAGCGTATC | 3'UTR |
| | NM_010423 | V3LMM_73187 (#5) | TAACATTGAGCATAGTGGC | 3'UTR |
| shSmad1 | NM_008539 | V2LMM_30824 (#1) | TAAGACACGGATGAAATAG | ORF |
| | NM_008539 | V3LMM_32418 (#3) | ATAGTCTACATTTGCAGCC | 3'UTR |
| | NM_008539 | V3LMM_457232 (#5) | AGCTGAGCAAACTCTTGGT | ORF |
